# Tumor to normal single cell mRNA comparisons reveal a pan-neuroblastoma cancer cell

**DOI:** 10.1101/2020.06.22.164301

**Authors:** Gerda Kildisiute, Waleed M. Kholosy, Matthew D. Young, Kenny Roberts, Rasa Elmentaite, Sander R. van Hooff, Eleonora Khabirova, Alice Piapi, Christine Thevanesan, Eva Bugallo Blanco, Christina Burke, Lira Mamanova, Philip Lijnzaad, Thanasis Margaritis, Frank C.P. Holstege, Michelle L. Tas, Marc H.W.A. Wijnen, Max M. van Noesel, Ignacio del Valle, Giuseppe Barone, Reinier van der Linden, Catriona Duncan, John Anderson, John C. Achermann, Muzlifah Haniffa, Sarah A. Teichmann, Dyanne Rampling, Neil J. Sebire, Xiaoling He, Ronald R. de Krijger, Roger A. Barker, Kerstin B. Meyer, Omer Bayraktar, Karin Straathof, Jan J. Molenaar, Sam Behjati

## Abstract

Neuroblastoma is an embryonal childhood cancer that arises from aberrant development of the neural crest, mostly within the fetal adrenal medulla. It is not established what developmental processes neuroblastoma cancer cells represent. Here, we sought to reveal the phenotype of neuroblastoma cancer cells by comparing cancer (n=16,591) with fetal adrenal single cell transcriptomes (n=57,972). Our principal finding was that the neuroblastoma cancer cell resembled fetal sympathoblasts, but no other fetal adrenal cell type. The sympathoblastic state was a universal feature of neuroblastoma cells, transcending cell cluster diversity, individual patients and clinical phenotypes. We substantiated our findings in 652 neuroblastoma bulk transcriptomes and by integrating canonical features of the neuroblastoma genome with transcriptional signals. Overall, our observations indicate that there exists a pan-neuroblastoma cancer cell state which may be an attractive target for novel therapeutic avenues.

---

Neuroblastoma is a childhood cancer that exhibits a diverse pattern of disease (*1*), from spontaneously resolving tumors to a highly aggressive cancer. Neuroblastoma arises from aberrant differentiation of the neural crest, mostly within the fetal adrenal medulla, via the sympathetic lineage. Different stages of adrenal medullary cell differentiation have been implicated as the cell of origin of neuroblastoma (*2–4*) and proposed to underlie the diversity of clinical phenotypes (*2,5*).

Advances in single cell transcriptomics have enabled the direct comparison of cancer and normal reference cells. Applied to childhood cancer, such analyses have revealed specific cell types or developmental processes that tumors adopt and adapt (*6*). Here, we sought to establish, through cancer to normal single cell mRNA comparisons, which developmental processes and cell types neuroblastoma recapitulates and correlate our findings with disease phenotypes.

In the first instance, we built a normal cell reference for our analyses, by defining the transcriptional changes underpinning human adrenal development. We subjected to single cell mRNA sequencing (10x Genomics Chromium platform) seven adrenal glands obtained from first and second trimester human fetuses (**table S1**). To verify technical and biological replicability we included bilateral adrenal glands in two cases. Following stringent data filtering, including removal of doublets and ambient mRNAs, we obtained count tables of gene expression from a total of 57,972 cells. These cells could broadly be divided into cortical, medullary, mesenchymal cells, and leukocytes (**Fig. 1A, fig. S1**), using canonical human adrenal markers (**table S2**). We focused our analyses on medullary cells (n=6,451), from which most cases of neuroblastoma arise. Within medullary cells (**Fig. 1B**) we defined developmental trajectories as a reference for subsequent analyses, using two independent, widely adopted methods (trajectory analysis (*7*) and RNA velocity (*8*))(**Fig. 1C**). Overall the developmental sequence of human medullary cells mirrored trajectories recently described in murine adrenal glands (*9*) although we observed inter-species variation in expression of some medullary marker genes (**fig. S2-3**). Human medullary development emanated from a population of neural crest like cells (Schwann cell precursors, SCP). The developmental trajectory then continued via a common root of so called bridge cells and bifurcated into sympathoblastic and chromaffin cell lineages (**Fig. 1B to C**), each associated with distinct expression patterns of transcription factors (**Fig. 1D, fig. S4**).

**Figure 1.**
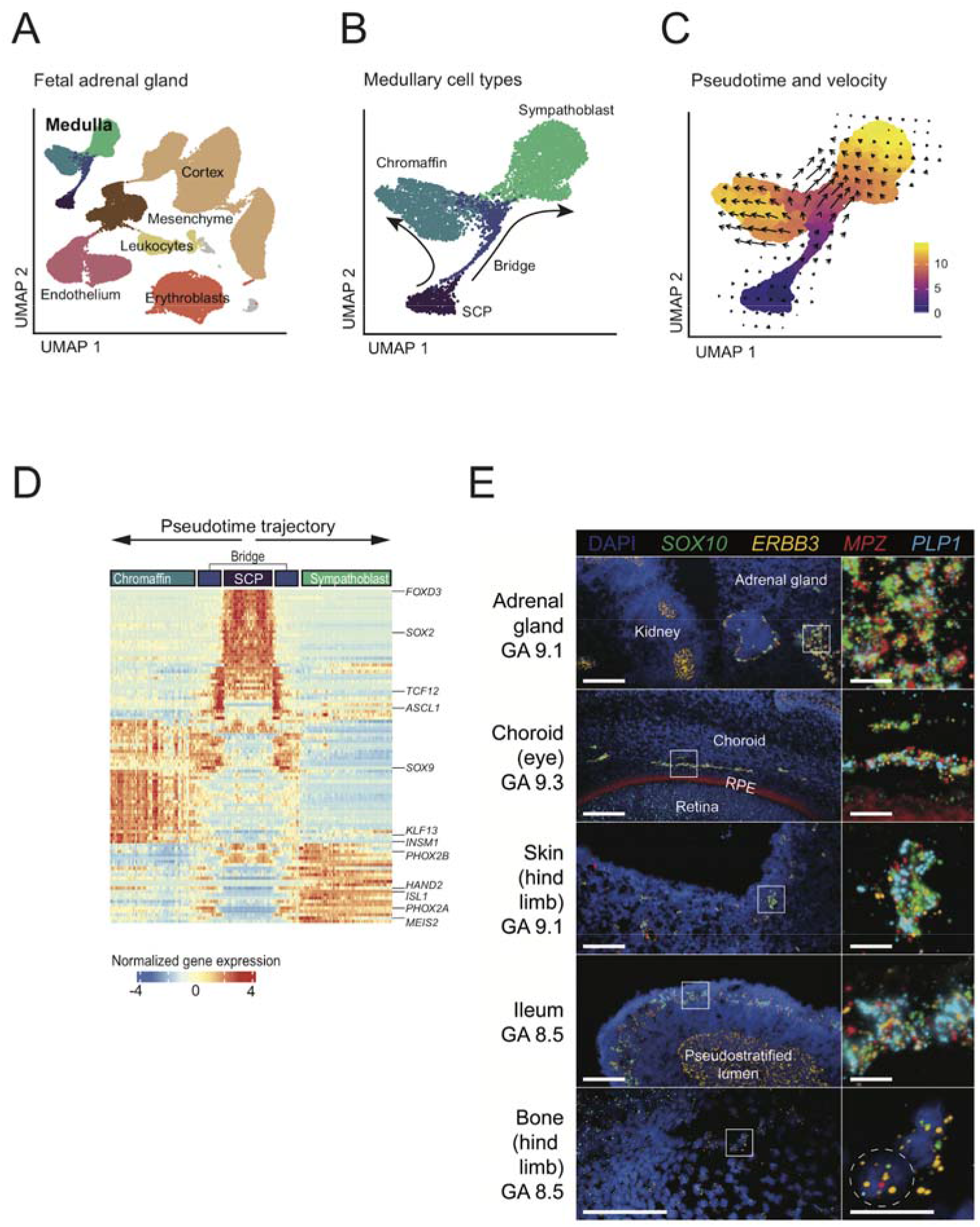
Single cell reference data from seven human fetal adrenal glands. **(A)** UMAP (Uniform Manifold Approximation and Projection) representation of 57,972 fetal adrenal gland cells. Cell types (determined by marker genes, **fig. S1**) are labelled and indicated by cell colour. **(B)** Zoom in of 6,451 fetal medulla cells with detailed cell annotation and arrows indicating developmental trajectories. **(C)** Developmental trajectory of fetal medulla. Colours represent pseudotime direction, while arrows show the cellular state progression (RNA Velocity). **(D)** Top 100 most significant transcription factors that vary with medulla development as defined in (B,C). The center of the heatmap corresponds to SCP, and proceeds along arrows towards chromaffin and sympathoblastic destinations, via bridge cells. Widely known genes of adrenal development are highlighted (full list in **fig. S4**). **(E)** RNAscope single-molecule fluorescent *in situ* hybridisation staining of SCP marker genes in human fetal tissues. Scale bars: left, 100 μm; insets 20 μm.

Of particular note were the neural crest like SCP cells at the root of the trajectory, identified previously in murine (*9–11*), but not human fetal tissues. We searched for these SCPs in several human fetal tissues using single cell mRNA data, and validated these by multiplexed single molecule FISH of SCP defining markers, *SOX10, ERBB3, MPZ*, and *PLP1*. Accordingly, we identified SCPs in fetal gut, eye, skin, and bone, (**Fig. 1E**), indicating that SCPs may contribute to human development across several different fetal tissues.

We next sought to establish the relationship between neuroblastoma cells and human fetal medullary development. We generated single cell mRNA readouts from 18 fresh neuroblastoma specimens from two different centres, using two platforms (Chromium 10x, CEL-seq2). We obtained tissue from untreated patients at diagnosis (n=4), or following treatment with cytotoxic agents at resection (n=14)(**table S3**). In a subset of these samples (n=8), we detected cells carrying the tumour’s genotype, while in some samples (n=6) we detected only stromal or unverified tumour cells. As pre-treated neuroblastomas tend to be largely necrotic at surgical resection, we guided sampling to viable tumor areas through pre-operative metabolic cross-sectional imaging in these specimens (meta-iodobenzylguanidine scan), coupled with morphological assessment of frozen sections in 12/14 pretreated cases.

Our study cohort represented the three principal prognostic categories of neuroblastoma: low (n=3), intermediate (n=5), and high risk (n=10) cases. In total we obtained 16,591 cells, with variable contribution from each tumor (**Fig. 2A to B; table S4**), which segregated into four main cell types (**Fig. 2A to B; fig. S5 to 6**): leukocytes (n=9,341), mesenchymal cells (n=4,073), Schwannian stroma cells (n=251), and putative tumor cells exhibiting adrenal medullary-like features (n=2,296).

**Figure 2.**
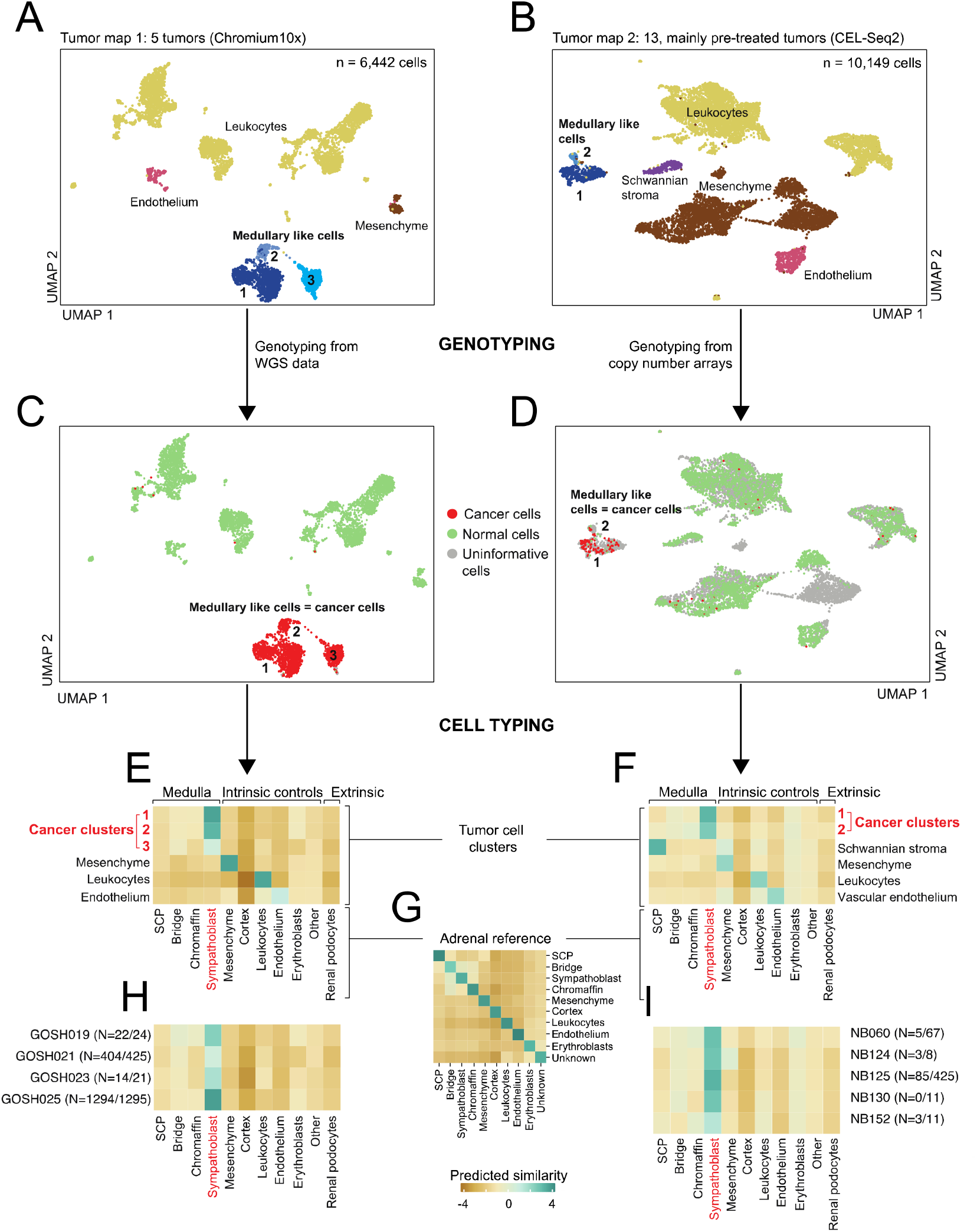
Neuroblastoma single cell transcriptomes. **(A**,**B)** UMAP representation of 6,442 **(A)** and 10,149 **(B)** neuroblastoma cells. Colours and labels indicate different cell types defined using marker gene expression (**fig. S5-6**). **(C**,**D)** The same UMAP representation as (A,B) but here cell colour represents each cell’s genotype, with red indicating the cancer genotype, green the wild type genotype and grey insufficient information. **(E**,**F)** Logit of normal to tumor cell similarity score (logistic regression), comparing the reference fetal adrenal gland (x-axis) to neuroblastoma cells (y-axis) containing intrinsic (non-cancer cells) and an extrinsic (kidney podocytes) control cell population. **(G)** Confusion matrix of fetal adrenal gland model used in (E,F) to predict similarity to normal populations. Colours indicate logit transformed similarity scores: 0 by default, positive (negative) indicating strong similarity (dissimilarity). **(H**,**I)** Similarity of cells from the cancer clusters in (E,F) grouped by patient. Numbers in parenthesis indicate number of tumour cells with the validated genotype/ total number of tumour cells. Only patients with at least 10 tumour cells, or at least one tumour cell with validated genotype were included.

The first challenge was to identify *bonafide* cancer cells amongst tumor derived cells. Marker based cell typing alone was of limited value as controversy exists as to which cell types in neuroblastoma represent cancer cells. Although there is a general consensus that neuroblastoma cancer cells exhibit medullary-like features, it has also been suggested that interstitial (mesenchymal) and Schwannian stroma cells, commonly found in neuroblastoma, may be cancerous (*12–14*). We therefore used somatic copy number changes to identify cancer cells, by interrogating each cell’s mRNA sequence for evidence of the somatic copy number changes underpinning each tumor. Extending a previously developed method (*6*), we integrated single cell tumor RNA information with single nucleotide polymorphisms (SNP) arrays or whole genome sequencing of the tumor DNA. This allowed us to assess the allelic imbalance of SNPs in each cell’s mRNA sequence across patient specific copy number segments. This analysis revealed that in our cohort only adrenal medullary-like cells were cancerous, but not interstitial or Schwannian stroma cells (**Fig. 2C to D**).

Next, we investigated which stage of adrenal medullary development cancer cells recapitulate by performing cancer to normal cell comparisons. We found that neuroblastoma cancer cells did not recapitulate adrenal development, but had assumed the state of sympathoblasts only (**Fig. 2E to G**). Importantly, although cancer cells formed discrete clusters within, and across patients (**Fig. 2A to B**), the cancer to normal cell comparison resolved this diversity into a common sympathoblast-like phenotype. The sympathoblast state was also detected in all patients which contribute cells confirmed to carry the genotype to the tumour cell clusters (**Fig. 2H to I**).

To validate that the neuroblastoma cancer cells principally resemble sympathoblasts, we extended our analyses using neuroblastoma bulk transcriptomes. We curated gene expression profiles of 652 neuroblastomas from two different clinically annotated cohorts that represent the entire clinical spectrum of neuroblastoma (TARGET (n=154) (*15*) and SEQC (n=498) (*16*) cohorts). We probed these expression data for transcripts that define the four populations of the normal human fetal medulla: SCPs, bridge cells, sympathoblasts, and chromaffin cells (**Fig 1B**). Across cohorts we found a clear signal of sympathoblast mRNAs (**Fig. 3A**), thus verifying the sympathoblastic state of neuroblastoma. Neuroblastomas that arose outside the adrenal gland also exhibited sympathoblast signals (**Fig. 3B**). The sympathoblast signal was present across the principal risk groups of neuroblastoma, suggesting that it transcends the clinical diversity of neuroblastoma (**Fig. 3C**). Interestingly, compared to lower risk tumors, the sympathoblast signal was less pronounced in high risk tumors in both cohorts, which may be driven by differences in tissue composition or in cancer cells themselves.

**Figure 3.**
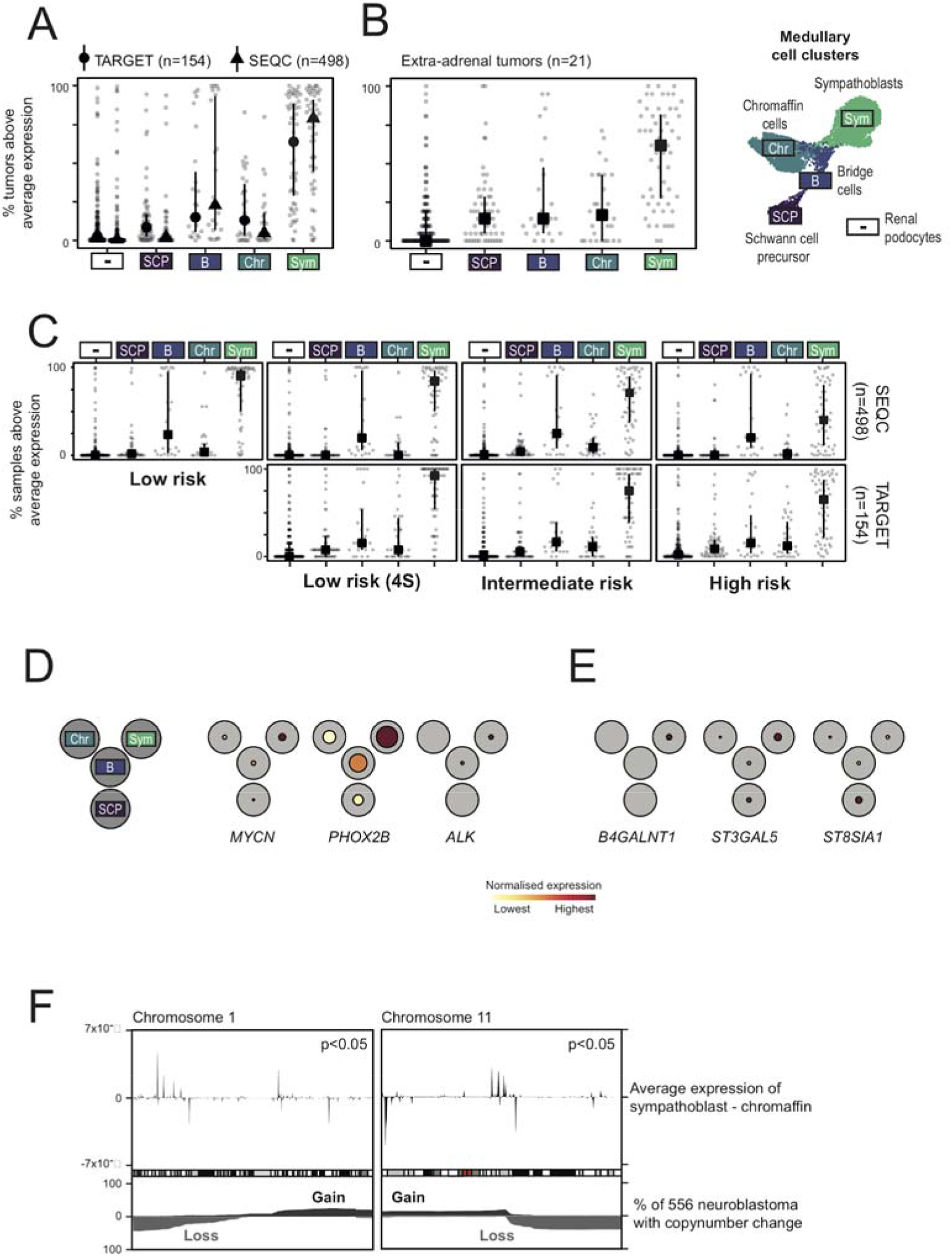
Validation in clinically annotated neuroblastoma cohorts. **(A)** Fraction of bulk neuroblastoma samples (in TARGET and SEQC) where the expression of fetal medulla (plus podocyte negative control) marker genes are above average. For each dataset/cell-type combination points indicate individual marker genes, lines indicate interquartile ranges, and symbols the median. **(B)** The same as (A) but limited to samples taken outside the adrenal gland. **(C)** The same as in (A) but with samples split by clinical risk group (horizontal facets) and datasets (vertical facets). Note there are no non-4S, low risk samples in the TARGET dataset. **(D)** Expression of the principal somatic and germline predisposition neuroblastoma genes in the fetal medulla. The black circles show a reduced resolution representation of the trajectory in **Fig. 1C** with the radius of the coloured circle relative to the grey circle indicating the fraction of cells expressing each gene in that region of the trajectory and the colour indicating average expression, normalised for each gene. Further genes are shown in **fig. S7.** **(E)** As in (D), but for GD2 synthesis genes **(F)** Average difference in expression between sympathoblastic and chromaffin cells as a function of genomic position on chromosomes 1 and 11 (top panel), with positive (negative) values indicating higher expression in smypathoblasts (chromaffin) cells. The bottom panel shows the fraction of samples with copy number changes in 556 neuroblastomas, with grey (black) indicating copy number loss (gain).

To further corroborate the transcriptional evidence that neuroblastoma cells resemble sympathoblasts, we overlaid features of the neuroblastoma cancer genome with transcriptional developmental pathways. We first analysed how somatic cancer (*15*) and germline predisposition (*17*) genes of neuroblastoma are utilised by fetal medullary cells. We found that the majority of these genes were most highly expressed along the trajectory of sympathoblast differentiation (**Fig. 3D; fig. S7**), in particular the two most common neuroblastoma predisposition genes, *ALK* and *PHOX2B*, and the cancer gene, *MYCN*, somatic amplification of which is a key adverse marker used clinically for neuroblastoma risk stratification. Thus, the same genes that operate as cancer genes in neuroblastoma are utilised in normal development predominantly by sympathoblasts. Similarly, genes encoding the synthases of disialoganglioside (GD2), which is a near ubiquitous marker of neuroblastoma that is therapeutically targeted by antibody treatment (*18*), were predominantly expressed in sympathoblasts (**Fig. 3E**).

A variety of copy number changes have been described in neuroblastoma, the most recurrent and pertinent of which occur on chromosomes 1, 11, and 17 (*19*). The presence of these copy number changes carries fundamental prognostic significance and determines treatment intensity in most clinical contexts and treatment protocols. Given the sympathoblastic state of neuroblastoma, it seemed conceivable that somatic copy number changes may impact on expression of genes that define sympathoblasts. We were able to directly test this hypothesis. For example, we compared gene expression of cancer cells that harbour loss of chromosomes 1 and 11 with cancer cells that do not carry these changes (**table S5**). We found that the resulting differentially expressed genes were significantly enriched for high confidence sympathoblast marker genes (tf-idf>1, 283 out of 329 markers, p<0.0001, hypergeometric test)(**table S6**). Furthermore, the majority of these genes (254/329) exhibited lower expression in cells with chromosome 1 and 11 loss, even if those genes did not reside on chromosomes 1 or 11. Following on from this, we correlated genomic regions of recurrent chromosome 1 or 11 loss with the genomic localisation of gene expression in fetal medullary cells. The localisation of the boundaries of altered copy number segments is associated with gene expression in human cancer (*20*). Thus, the genomic position of these boundaries may encode information about the expression profile of the cancer cell of origin. We found regions of sympathoblast specific gene expression that statistically significantly overlapped with the most common boundaries of chromosome 11q loss (p<0.05; permutation test)(**Fig. 3F; fig. S8**) and the most recurrent region of chromosome 1p loss (p<0.05; permutation test)(**Fig. 3F; fig. S9**), thus further corroborating the sympathoblast state of neuroblastoma.

Having established the similarities between neuroblastoma cancer cells and fetal medullary development, we now looked for differences. Previous bulk mRNA analyses have described gene expression changes that characterise neuroblastoma tissues (*3, 21*). Our ability to directly compare cancer cells with their normal counterpart, fetal medullary cells, enabled us to distil the transcriptional essence of neuroblastoma cells, which comprises 106 differentially expressed genes (tf-idf > 0.85, **table S5**). To verify, and to correlate these findings with disease phenotypes, we measured the mRNA levels of these differentially expressed genes in bulk neuroblastoma transcriptomes. A sizable number of these differentially expressed genes (n=38/106) was present in more than half of neuroblastoma bulk samples (**fig. S10**). The expression of 13 of these varied significantly by clinical risk group, even after controlling for key determinants of clinical risk, age and *MYCN* amplification (negative binomial generalised linear model) (**Fig. 4A to B**). In addition to known markers of clinical risk (eg, *PRAME*), we identified novel markers that have not previously been described and could in theory be used to improve risk stratification in clinical practice in the future. Biologically, the transcription factor gene, *SIX3*, may be one of the most interesting findings amongst differentially expressed genes, representing an exclusively embryonal gene that normally regulates forebrain and eye development (*22*).

**Figure 4.**
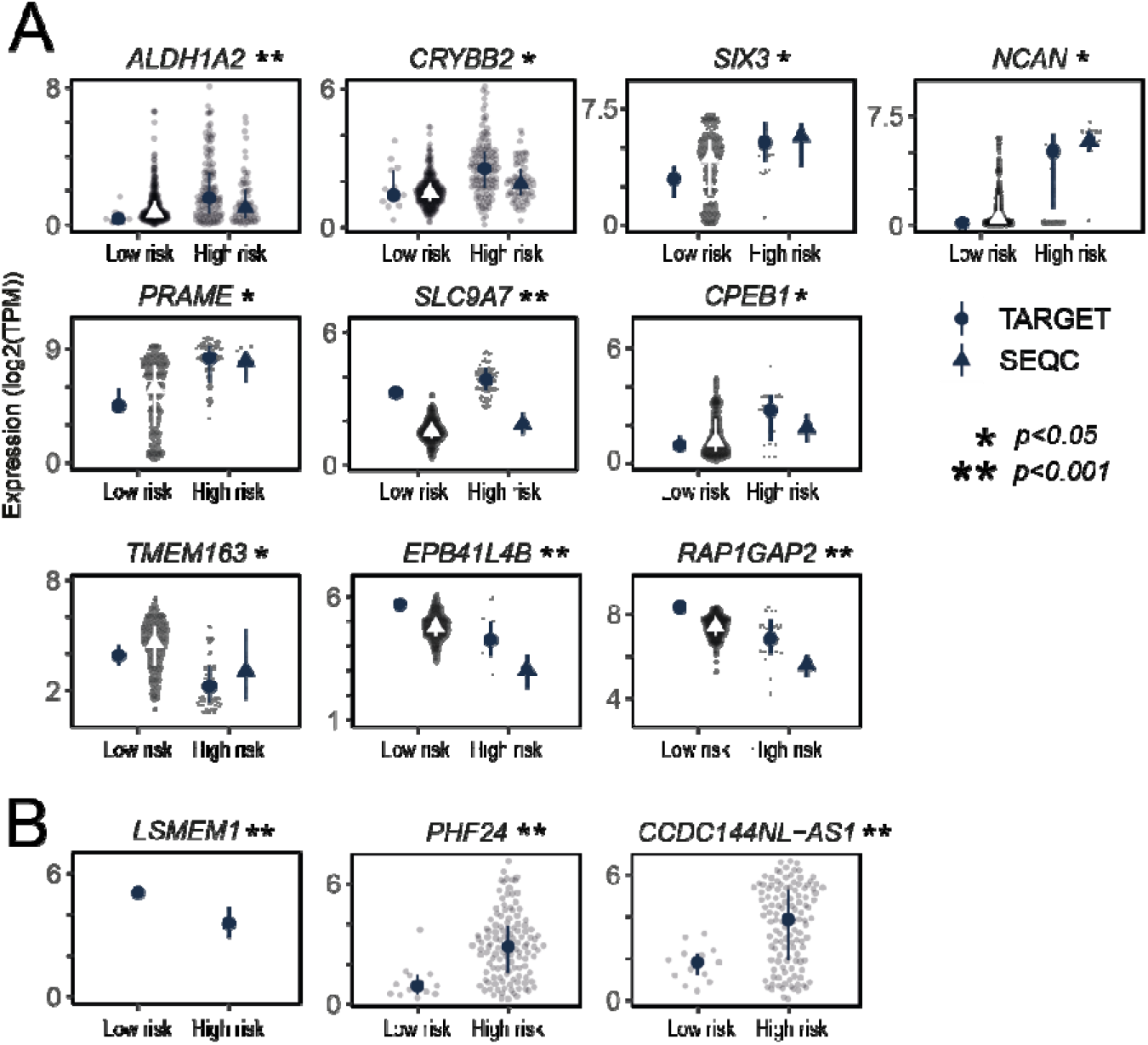
Integration with the neuroblastoma genome and differential gene expression. **(A)** Genes differentially expressed between neuroblastoma tumour cells and foetal medulla that exhibit significant risk group dependence after controlling for MYCN status and age (negative-binomial test, log_2_FC>1, FDR<0.05). Genes which showed an inconsistent risk group dependence in SEQC were removed. For the gene listed above each panel, dots indicate expression (log_2_(TPM)) in individual samples, lines the interquartile range, and symbols the median. **(B)** Genes not present in annotation used by SEQC.

The principal finding of our investigation was that the neuroblastoma cancer cell represented an aberrant fetal sympathoblast. Our expectation might have been that neuroblastoma exhibits a cellular hierarchy adopted from adrenal development, similar to, for example, the childhood kidney cancer, Wilms tumor, recapitulating fetal nephrogenesis. However, neither our direct analyses of single cancer cells, nor indirect insights from bulk cancer transcriptomes would corroborate such a hypothesis. Whilst other neuroblastoma cell types may exist, the sympathoblastic state predominates across the entire spectrum of this enigmatic cancer. In the context of efforts to derive novel treatments for neuroblastoma, such as differentiation agents, the sympathoblastic state may thus lend itself as a pan-neuroblastoma target.

## Supporting information

supplementary scripts

## Acknowledgements

The human embryonic and fetal material was provided by the Joint MRC / Wellcome (MR/R006237/1) Human Developmental Biology Resource (www.hdbr.org). All research at Great Ormond Street Hospital NHS Foundation Trust and UCL Great Ormond Street Institute of Child Health is made possible by the NIHR Great Ormond Street Hospital Biomedical Research Centre. The views expressed are those of the author(s) and not necessarily those of the NHS, the NIHR or the Department of Health and Social Care. We are indebted to the children and families who participated in our research.

## Author contributions

S.B., J.J.M., and K.S. conceived of the experiment. G.K., M.D.Y. and S.B. analyzed data, aided by R.E., S.R.v.H., E.K., I.d.V.T.,J.Ac. Statistical expertise was provided by M.D.Y. K.R. and O.B. performed smFISH experiments. S.A.T. contributed fetal single cell data, together with M.H., K.B.M., R.A.B., X.H., L.M. Tumor samples were curated and/or experiments were performed by W.M.K., A.P., C.T., E.B.B., C.B., G.B., C.D., J.A., P.L., T.M., F.C.P.H., M.L.T., M.H.W.A.W., M.M.v.N., R.v.d.L. Pathological expertise was provided by D.R., N.J.S., and R.R.d.K. G.K., M.D.Y., and S.B. wrote the manuscript. M.D.Y. directed analytical method development and statistical analyses. K.S., J.J.M., and S.B. co-directed the study.

## Data and materials availability

Raw sequencing data have been deposited in EGA. Bulk RNA-seq data were obtained from (*15*) and (*16*).

