## supplementary materials for "Tumor to normal single cell mRNA comparisons reveal a pan-neuroblastoma cancer cell"

**Materials and Methods**

Ethics statement

Informed consent for research was obtained from participants (or their carers). Studies underlying this paper had received appropriate approval by ethics review boards as per national legislation. Dutch tumor samples were obtained through an institutionally approved research study. UK tumor samples were collected under the following studies: NHS National Research Ethics Service reference 16/EE/0394 (tumor samples) and NHS National Research Ethics Service reference 96/085 (fetal tissues). Additional fetal tissue was provided by the Joint MRC / Wellcome Trust-funded (grant # 099175/Z/12/Z) Human Developmental BiologyResource (HBDR, [http://www.hdbr.org](http://www.hdbr.org/); (10)), with appropriate maternal written consent and approval from the Newcastle and North Tyneside NHS Health Authority Joint Ethics Committee. HDBR is regulated by the UK Human Tissue Authority (HTA;[www.hta.gov.](http://www.hta.gov.uk/)uk) and operates in accordance with the relevant HTA Codes of Practice.

Fetal adrenal tissue processing

Tissue was minced with a scalpel and then placed in 15ml falcon tube filled with 5ml Digestion solution (5ml, dilute Liberase TH stock solution (1 vial of 5mg Liberase TH (Sigma 5401135001) powder in 2ml of 1X PBS with final concentration is 2.5 mg/ml.) in 1X PBS (100ul enzyme stock + 4.9ml 1x PBS). Tissue was incubated in 37’C for 30min (water bath) and mixed with 1ml pipette after 15 minutes to facilitate the digestion. 5ml STOP solution (40ml, 2% FBS in 1x PBS (40ml 1x PBS + 800ul FBS)) was added. The mixture was filtered through a 30um strainer and spun down, 750g, 5min at 4’C. If the pellet was red 1ml of 1X RBC (00-4300-54) lysis buffer was added and incubated for 3min at RT. 10ml of STOP solution was added and cells were gently mixed to wash, then centrifuged, 750g, 5min, 4’C. Pellets were resuspended into appropriate amount of 1x PBS or STOP solution, counted and processed on 10x Chromium platform.

Neuroblastoma tissue processing - GOSH pipeline

Surplus tumour tissue obtained at diagnostic biopsy or tumour resection was processed immediately after receipt in the histopathology laboratory (< 1 hour after interventional radiology/surgical procedure). Tissue was minced using a scalpel, and then incubated in RPMI, supplemented with 10% FCS, 1% L-glutamine and 1% Penicillin/Streptomycin, with Collagenase IV (1.6 mg/ml; cat. No. 11410982; MP Biomedicals), for 30 minutes at 37°C, inverting the tube every 10 minutes. The digested tissue was passed through a 70 μm filter, and incubated in 1x Red Blood Cell lysis buffer (cat. No. 420301; Biolegend) for 10 minutes, at room temperature. The obtained single cell suspension was used for downstream processing. Part of the single cell suspension was depleted of CD45^+^ cells to enrich for tumour cells using a CD45 MicroBeads kit (cat. No. 130-045-801, Miltenyi Biotec), following manufacturer’s protocol. Both CD45 non-depleted and CD45-depleted single cell suspensions were depleted of dead cells using a Dead Cell Removal Kit (cat. No. 130-090-101, Miltenyi Biotec), following manufacturer’s protocol. Obtained viable single cell suspensions (CD45-depleted: 2 channels and CD45-non depleted (1 channel) were processed on the 10x Chromium platform.

Neuroblastoma tissue processing - PMC pipeline

Tumour tissue was collected, after obtaining informed consent of the parents of neuroblastoma patients and after the study was approved by the ethical committee at the Princess Maxima Centrum (PMC). MIBG-positive neuroblastoma samples were included if they were diagnosed with histologically proven viable tumor cells. Most samples were debulking neuroblastomas (patients were treated with debulking type surgical procedure) and preoperative radiation and chemotherapy have been received (**table S3**). We extensively optimized the workup protocols and samples are now mildly dissociated to the single-cell level and sorted by flow cytometry. Single-cell suspensions were subsequently subjected to single-cell RNA-Seq using the CEL-seq2 platform. In brief, freshly resected human neuroblastoma specimens were collected and processed immediately following histological confirmation of the presence of viable tumor tissue. Tumor tissue was minced into 3-4 mm pieces and digested with collagenase type I, II and IV (2 mg/ml) in the optimized medium at 37°C, with agitation for 2 hours. The resulting suspension containing small tumor fragments was passed through a 70μm cells strainer, washed with a cold organoid medium. Half of the tumor fragments were used to generate tumor organoids while the other half were fragments were further dissociated with NeuroCult kit until the single-cell suspension was obtained. Cells were washed with cold organoid medium and stained with 7AAD or DAPI to enrich for live cells while others were stained with FITC-GD2 or PE-GD2 antibodies to enrich for tumor and PE-CD3 antibodies to enrich for T cells and to exclude erythrocytes. Cells were stored on ice until sorting using a FACS Jazz or FACS AriaII. Single live cells were sorted into 384-well hard-shell plates with 10 μl mineral oil, 50 nl of RT primers, dNTPs and synthetic mRNA Spike-Ins and immediately spun down, snap-frozen on dry ice, and stored in -80C to proceed with the total transcriptome amplification, library preparation and sequencing.

10X library preparation and sequencing

The concentration of single cell suspensions were manually counted using a hemocytometer and adjusted to 1000 cells/ul or counted by flow cytometry. Cells were loaded according to standard protocol of the Chromium single cell 3’ kit (v2 and v3 chemistry). All the following steps were performed according to the standard manufacturer protocol. We used one lane of an Illumina Hiseq 4000 per 10x chip position.

CEL-seq2 library preparation and sequencing

All samples that were processed proceeded with the total transcriptome amplification, library preparation and sequencing into Illumina sequencing libraries as previously described [(*23*)](http://f1000.com/work/citation?ids=2450417&pre=&suf=&sa=0). Paired-end 2X75 bp sequencing read length was used to sequence the prepared libraries using the Illumina NextSeq sequencer.

Bulk DNA processing of GOSH samples

DNA was extracted from fresh frozen tissue. Peripheral blood DNA was used as a matched normal. Short insert (500bp) genomic libraries were constructed, flowcells prepared and 150 base pair paired-end sequencing clusters generated on the Illumina HiSeq X platform according to Illumina no-PCR library protocols [(*24*)](http://f1000.com/work/citation?ids=426293&pre=&suf=&sa=0). The average sequence coverage was 43X and 42X for tumour and matched peripheral blood samples, respectively.

Bulk DNA processing of PMC samples

DNA was extracted from fresh tissue via AllPrep DNA/RNA/Protein Mini Kit (QIAGEN) according to standard protocol.

Bulk DNA alignment

DNA sequencing reads were aligned to the GRCh 37d5 reference genome using the Burrows-Wheeler transform (BWA-MEM) [(*25*)](http://f1000.com/work/citation?ids=49016&pre=&suf=&sa=0). Sequencing depth at each base was assessed using Bedtools coverage v2.24.0.

Single molecule fluorescent in situ hybridisation

Fresh tissue samples were embedded in OCT and frozen at -80°C on an isopentane-dry ice slurry. Cryosections were cut at a thickness of 16 μm using a Leica CM3050 S cryostat and placed onto SuperFrost Plus slides (VWR). Prior to staining, tissue sections were post-fixed in 4% paraformaldehyde in PBS for 15 minutes at 4°C, then dehydrated through a series of 50%, 70%, 100%, and 100% ethanol, for 5 minutes each. Tissue sections were then processed using a Leica BOND RX to automate staining with the RNAscope Multiplex Fluorescent Reagent Kit v2 Assay and RNAscope 4-plex Ancillary Kit for Multiplex Fluorescent Reagent Kit v2 (Advanced Cell Diagnostics, Bio-Techne), according to the manufacturers’ instructions. Automated processing included pre-treatment with Protease IV for 30 minutes, but no heat treatment. Tyramide signal amplification with Opal 520, Opal 570, and Opal 650 (Akoya Biosciences) was used to develop three probe channels. The fourth was developed using TSA-biotin (TSA Plus Biotin Kit, Perkin Elmer) and streptavidin-conjugated Atto 425 (Sigma Aldrich). Stained sections were imaged with a Perkin Elmer Opera® Phenix™ High-Content Screening System, in confocal mode with 1 μm z-step size, using 20× (NA 0.16, 0.299 μm/pixel) and 40× (NA 1.1, 0.149 μm/pixel) water-immersion objectives.

Copy number detection from DNA data

The ascatNGS algorithm [(*26*)](http://f1000.com/work/citation?ids=8685620&pre=&suf=&sa=0) (v4.0.1) was used to estimate tumor purity and ploidy and to construct copy number profiles prior to running the Battenberg algorithm (v2.2.5) (github.com/cancerit/cgpBattenberg) to allow for tumor sub-clonality in bulk DNA sequencing data. For DNA data measured with Illumina SNP arrays, copy number segments were defined by manual inspection of logR and B allele frequency data.

Mapping and quantification of scRNAseq - 10X

Single cell RNA-seq data were mapped and counts of molecules per barcode quantified using the 10X software package cellranger (versions 2.0.2 and 3.0.2) to map sequencing data to version 2.1.0 of the build of the GRCh38 reference genome supplied by 10X.

Mapping and quantification of scRNAseq - CEL-seq2

Sharq preprocessing and QC pipeline has been applied to process the single-cell RNA-seq data as described [(*27*)](http://f1000.com/work/citation?ids=6727911&pre=&suf=&sa=0). Read mapping has been done using STAR version 2.6.1 on the hg38 Patch 10 human genome. The featureCounts function of the subread package (version 1.5.2) was used to assign reads based on GENCODE version 26.

Quality control of fetal adrenal single cell data

Ambient mRNA contamination was removed from each channel individually using SoupX [(*28*)](http://f1000.com/work/citation?ids=5172334&pre=&suf=&sa=0). Cleaned count data was then normalised, scaled to have mean 0 and standard deviation 1 for each gene, principal component analysis was performed using highly variable genes, and grouped into clusters using a community detection finding algorithm taking the first 75 principal components as inputs and a resolution parameter of 10. This was performed using the Seurat package in R [(*29*, *30*)](http://f1000.com/work/citation?ids=5027067,7035390&pre=&pre=&suf=&suf=&sa=0,0). The clustering parameter was deliberately set to this extremely high value to produce a very large number of small clusters.

Following clustering, we designated each cell as either passing or failing a series of quality control metrics. We marked as failing QC any cell with >30% expression due to mitochondrial genes, fewer than 300 genes detected, fewer than 1000 UMIs detected, or a doublet score greater than 0.2 as determined by Scrublet [(*31*)](http://f1000.com/work/citation?ids=6776726&pre=&suf=&sa=0). In addition to excluding these individual cells, we also marked as failing QC any cell belonging to a cluster containing more than 50% of cells that fail QC due to one of the above filters. Finally, we excluded from all further analysis any cell marked as QC failed for any of the above reasons (**fig. S11 to S15**).

Next we calculated a S and G2M phase score for each cell using Seurat [(*29*, *30*)](http://f1000.com/work/citation?ids=5027067,7035390&pre=&pre=&suf=&suf=&sa=0,0) and excluded any cell where either score was greater than 0. This removed any cells that showed evidence of being in a phase of the cell cycle other than G0/G1. We excluded these cells from our reference map as cells in the S/G2M phase tend to cluster together based on their phase of cell cycle and not their underlying cell type, reducing the utility of the data as a reference map of cell types present in the developing adrenal gland.

As before, data was normalised by sequencing depth, scaled to 10,000 counts, log transformed, and genes were scaled to have mean 0 and standard deviation 1. Principal component analysis was performed on the scaled data using the most highly variable genes. We selected the number of principal components to use for clustering and visualisation to minimise the molecular cross validated error [(*32*)](http://f1000.com/work/citation?ids=7554051&pre=&suf=&sa=0) (<https://github.com/constantAmateur/MCVR>). Using these principal components, we calculated a uniform manifold approximation and projection [(*33*)](http://f1000.com/work/citation?ids=8688127&pre=&suf=&sa=0) two dimensional representation of the data for visualisation and calculated clusters using a k-nearest neighbours algorithm with resolution parameter set to 1.

Quality control of 10x tumour data

We excluded nuclear mitochondrial genes, heat shock proteins and ribosomal genes from our analysis. To filter lower quality cells, we removed any cells that had greater than 20% expression originating from mitochondrial genes, expressed fewer than 300 distinct genes, or contained fewer than 1000 UMIs (**fig. S16 to S17**). Data was normalised by sequencing depth, scaled to 10,000 counts, log transformed, and genes were scaled to have mean 0 and standard deviation 1 using the Seurat ‘NormalizeData’ function. Principal component analysis was performed on the scaled data using 2,000 most variable genes. Using 75 first principal components, we calculated a uniform manifold approximation and projection (UMAP, [(*33*)](http://f1000.com/work/citation?ids=8688127&pre=&suf=&sa=0) ) two dimensional representation of the data for visualisation and calculated clusters using a community detection algorithm with resolution parameter set to 1. We performed these steps using the Seurat package in R [(*29*, *30*)](http://f1000.com/work/citation?ids=5027067,7035390&pre=&pre=&suf=&suf=&sa=0,0).

Quality control of CEL-seq2 tumour data

We removed RNA spike-ins and genes that were not present in the 10X gene list. We excluded nuclear mitochondrial genes, heat shock proteins and ribosomal genes from our analysis. To filter lower quality cells, we removed any cells that had greater than 20% expression originating from mitochondrial genes, expressed fewer than 200 distinct genes, and contained fewer than 500 UMIs (**fig. S18 to S19**). We used less stringent thresholds for CEL-seq data than for 10X data due to data quality. Data was normalised by sequencing depth, scaled to 10,000 counts, log transformed, and genes were scaled to have mean 0 and standard deviation 1 using the Seurat ‘NormalizeData’ function. Principal component analysis was performed on the scaled data using 2,000 most variable genes. Using 50 first principal components, we calculated a uniform manifold approximation and projection (UMAP, [(*33*)](http://f1000.com/work/citation?ids=8688127&pre=&suf=&sa=0) ) two dimensional representation of the data for visualisation and calculated clusters using a community finding algorithm with resolution parameter set to 1. We performed these steps using the Seurat package in R [(*29*, *30*)](http://f1000.com/work/citation?ids=5027067,7035390&pre=&pre=&suf=&suf=&sa=0,0).

Cluster annotation

Using well established marker genes of different cell types curated from the literature (**table S2**), we assigned a cell type to each cluster . Where two clusters were annotated as the same cell type, we merged them together. As further confirmation of our annotation, we next identified marker genes for each population algorithmically (**tables S6, S8, S9**). To do this we used a method which uses the tf-idf metric to identify genes specific to each population, as implemented in the “quickMarkers” function in the SoupX R package[(*28*)](http://f1000.com/work/citation?ids=5172334&pre=&suf=&sa=0). We further filtered genes to include only those genes with p-value less than 0.01 after multiple hypothesis correction (hypergeometric test).

Pseudotime analysis

We ordered cells into a trajectory based on the similarity of their transcriptomes to create a pseudotime ordering of the medulla using monocle3 [(*7*, *34*, *35*)](http://f1000.com/work/citation?ids=17045,3053962,4249067&pre=&pre=&pre=&suf=&suf=&suf=&sa=0,0,0). We constructed a monocle3 object using data extracted from a Seurat object, rather than re-processing the data in monocle3. We used the “learn_graph” function to build a principal graph of the data. We then used “order_cells” function to place cells on a pseudotime trajectory, selecting a node (as defined in “learn_graph”) as a starting point of the trajectory. We used an SCP node based on prior knowledge from murine data that medullary cells originate from SCPs [(*9*)](http://f1000.com/work/citation?ids=3914331&pre=&suf=&sa=0).

Differential expression of transcription factors along pseudotime trajectory

To identify transcription factors that were differentially expressed along the two branches of the medulla ( SCP → sympathoblast and SCP → chromaffin), we performed Moran’s I test using “graph_test” function in monocle3. We limited our gene set to the 1,665 transcription factors [(*36*)](http://f1000.com/work/citation?ids=6586802&pre=&suf=&sa=0), and identified all genes that passed 0.001 p-value threshold after multiple hypothesis correction (**table S10**).

RNA Velocity

RNA velocity was calculated on the foetal adrenal reference map data using velocyto [(*8*)](http://f1000.com/work/citation?ids=5636602&pre=&suf=&sa=0) to calculate the number of reads supporting spliced/unspliced isoforms of each gene in each cell. The velocity field projection to the UMAP embedding was then calculated using the velocyto.R R package, pooling 20 k nearest neighbours and performing a gamma fit to the top/bottom 2% expression quantiles.

Cell similarity calculation

To measure the similarity of a target single cell transcriptome to a reference single cell data-set we used the methodology based on logistic regression outlined in detail in [(*6*)](http://f1000.com/work/citation?ids=5654099&pre=&suf=&sa=0). Briefly, we train a logistic regression model with elastic net regularization (alpha=0.99) on the reference training set. We then use this trained model to infer a similarity score for each cell in the query data set for each cell type in the reference data.

Softmax normalization was not used to allow for the possibility that some cells in the query data set do not resemble any of the cell types in the reference data set. Predicted logits were averaged within each cluster in the query dataset. This approach was implemented using the “glmnet” package in R [(*37*)](http://f1000.com/work/citation?ids=171561&pre=&suf=&sa=0).

Genotyping individual cells in single cell RNA-seq data

**10X data**

For 10X data, bulk DNA sequencing of tumour and normal tissue was available for all samples. For these samples, we first identified all heterozygous single nucleotide polymorphisms (SNPs) using bcftools [(*38*)](http://f1000.com/work/citation?ids=396559&pre=&suf=&sa=0) to find all locations with a coverage of at least 30 reads and between 20% to 80% of reads differing from the reference genome in the normal DNA sequencing. These locations were further filtered to keep only those sites where the null hypothesis of a homozygous genotype and sequencing error rate of 20% modelled with a binomial distribution could be rejected at a false discovery rate of 0.01 (BH [(*39*)](http://f1000.com/work/citation?ids=6279401&pre=&suf=&sa=0) ). We also exclude any sites with evidence of more than 2 genotypes.

Having found a list of heterozygous SNPs, we then identified regions of copy number (CN) change resulting in allelic imbalance (**table S11**). For each heterozygous SNP in each such CN region we calculated the B allele frequency (BAF) in the bulk Tumour data (defined as the fraction of reads mapping to the non-reference allele). We then performed a binomial test and marked as usable for phasing all SNPs with significant deviation from a BAF of .5 (either higher or lower) with false discovery rate 0.05. These “phasable SNPs” were then phased by designating the reference base as belonging to the minor allele (meaning the allele with lower copy number) for SNPs with BAF less than 0.5 and the alternate base as belonging to the minor allele for SNPs with BAF greater than 0.5.

In the single cell RNA-seq data we calculated the counts supporting the reference and alternate base at each SNP in each copy number region in each cell using allele counter [(*40*)](http://f1000.com/work/citation?ids=6684326&pre=&suf=&sa=0) (<https://github.com/cancerit/alleleCount>). We excluded any SNPs that fell outside of a gene and then used the phasing of SNPs to produce an aggregate count of reads supporting the major/minor allele in each cell in each copy number region.

The evidence in favour of a copy number change being present in each cell is then assessed by calculating the posterior probability of the altered CN state and an unaltered diploid configuration using a beta-binomial likelihood and uniform prior distribution. We assume an error rate of 0.01% in sequencing reads, which modifies the expected number of reads supporting the minor allele away from “number of copies of minor allele”/ploidy and closer to 0.5. We set the over-dispersion parameter of the beta-binomial likelihood by fitting a beta-binomial model to either the entire data set, or a subset of cells we strongly expect to have the normal genotype (e.g. leukocytes).

The final posterior probability of the cancer genotype for each cell is taken by calculating a combined likelihood across all copy number regions in the genome for both the cancer and wild type copy number configuration. Any cell with a posterior probability of the cancer genotype greater than 99% is designated as a tumour cell, any cell with less than 1% chance (i.e. 99% chance of the wild type, diploid genotype) is designated as normal, and any cell with intermediate posterior probability is labelled as ambiguous.

There is one exception to this approach, which was used for GOSH014 for which no bulk DNA assay was available. However, the clinical cytogenetic report revealed this sample to have a loss of chromosome 11p. To determine the genotype of cells based on just this information, we made the further assumption that the genotype of cells was common between cells within the same cluster (where clusters were determined using expression data). We also assume that at least 1% of cells have the normal genotype.

Using these assumptions we counted the number of reads supporting the reference and alternate allele at sites of common human variation (>10% SNP prevalence) in chromosome 11p in the single cell data [(*41*)](http://f1000.com/work/citation?ids=790619&pre=&suf=&sa=0). We calculated the binomial likelihood with p = 0.01 (error rate) and p = 0.5 (heterozygosity) for the number of reads supporting the alternate allele at each SNP aggregating across all cells and designated at heterozygous any SNP with a higher likelihood of the heterozygous model (p=0.5).

Finally, for each cluster of cells, we calculated the joint likelihood of all heterozygous SNPs on 11p appearing heterozygous and homozygous in this set of cells. These likelihoods are then compared to calculate a posterior probability of that cluster of cells containing the tumour genotype. This calculation is based on the observation that if the 11p loss is present in all cells in the cluster, then all heterozygous SNPs should appear homozygous due to one allele being lost. We then designate each cluster as having a tumour, normal, or ambiguous genotype as with other samples (i.e, 99% and 1% posterior probability cut-offs).

**CEL-Seq2 data**

For the CEL-Seq2 data, the only additional DNA information available was via Illumina CytoSNP 850k v1.1 arrays for each Tumour sample. To use this data to genotype cells, we identified regions of CN alteration as described above (**table S12**) and counted the reads supporting the reference and alternate allele at each SNP and each cell in the single cell RNA-seq data. We then aggregated counts across all cells and marked as heterozygous any sample that has evidence for both reference and alternate alleles across the array and single cell data. That is, we marked a SNP as heterozygous if it had significant expression of both ref and alt alleles in the single cell data (FDR<0.05, binomial test with null of homozygosity and error rate of 1%). Additionally, we also marked as heterozygous any SNP with evidence for one allele in the single cell data (same test) and the other allele in the tumour array data.

Having identified heterozygous SNPs in CN regions, we then phased these SNPs and calculated each cell’s posterior probability of the normal and cancer genotype as with the 10X data. The lower resolution and reliability (compared to whole genome sequencing) of the SNP array data lead to a far greater number of ambiguous genotype calls. Nevertheless, this data still enabled definitive genotyping for many cells in the single cell RNA-seq data.

Identification of cell-type specific genes

To identify genes that were specific to one cell type and no other in the adrenal medulla, we regenerated algorithmically defined markers as above, but with podocyte cells [(*6*)](http://f1000.com/work/citation?ids=5654099&pre=&suf=&sa=0) included as a negative control. To use only the most specific markers, we applied a tfidf cut-off of 1 to select genes that were specific to each medullary cell type relative to all other cells in the dataset. Note that a tf-idf cut-off of t implies that the global rate at which a gene occurs in cells is less than exp(-t/rate_local), where rate_local is the rate at which the gene occurs in the cells in the target cluster.

We also removed genes that were expressed in more than 20% of cells in any other single cluster in the dataset. The resulting gene list consisted of genes specific to each cell type in developing adrenal medulla (**table S13**).

Adrenal gland cell type signal in bulk RNA-seq data

To identify which adrenal cell type bulk RNA-seq Neuroblastomas resembled most, we measured the expression levels of genes defined in **table S13** in two bulk RNA-seq Neuroblastoma datasets, TARGET [(*15*)](http://f1000.com/work/citation?ids=58383&pre=&suf=&sa=0) and SEQC [(*16*)](http://f1000.com/work/citation?ids=112062&pre=&suf=&sa=0). For both data sets, the transcripts per million reads (TPM) value was calculated for each gene in each sample. We defined a gene as being “present” in a sample when its log_2_(TPM) expression value exceeded that of the peak of the log_2_(TPM) distribution across all genes across all samples. For each gene, we then calculated the fraction of samples in which it was present, both globally and stratified by risk group.

Defining neuroblastoma samples outside the adrenal gland

To define which neuroblastoma samples in TARGET [(*15*)](http://f1000.com/work/citation?ids=58383&pre=&suf=&sa=0) arose outside the adrenal gland, we filtered ICD-O [(*42*)](http://f1000.com/work/citation?ids=8693147&pre=&suf=&sa=0) description of each tumour by sites of occurrence. We excluded all samples where descriptions were absent or contained words “kidney”, “adrenal” , “abdominal”, “abdomen”, “unknown”, “retroperitoneum”, “other”, identifying 21 samples where tumours occured outside the adrenal gland.

Differential expression between fetal adrenal medulla and tumour cells

To identify the key transcriptomic differences between Neuroblastoma tumour cells and the normal foetal medulla, we performed differential expression analysis between fetal medulla single cells and tumour cells from CEL-seq2 and 10X tumour datasets. For each dataset, we extracted tumour clusters and merged them with fetal adrenal medullary cells. We labelled all tumour cells as “tumour”, and used the “quickMarkers” function from SoupX R package [(*28*)](http://f1000.com/work/citation?ids=5172334&pre=&suf=&sa=0) to identify tumour-specific genes for each dataset. We used a cut-off of 0.01 p-value after multiple hypothesis correction (hypergeometric test), and genes that were expressed in more than 20% of cells in any other single cluster in the dataset, to define a set of tumour markers.

We merged the two marker lists and calculated an average tfi-df value for each gene present in either or both lists, and used a tf-idf cut off of 0.85 to define a single list of tumour markers. We then filtered the list to exclude genes expressed by more than 25% of leukocytes in each tumour dataset (**table S7**).

Differential expression between low risk and high risk Neuroblastomas in TARGET

To test if any of the genes defined in **table S7** displayed risk group dependence, we compared their expression levels between 4S and high risk bulk Neuroblastomas in the TARGET data [(*15*)](http://f1000.com/work/citation?ids=58383&pre=&suf=&sa=0). To do this we used a negative binomial model, where we set the square root of the over-dispersion value to 0.4 (as genuine biological replicates were not available). We constructed a generalised linear model, with age, MYCN status, and risk group as covariates then used a quasi-likelihood F-test to calculate genes for which the risk group coefficient was significantly non-zero (0.05 FDR, edgeR [(*43*)](http://f1000.com/work/citation?ids=673952&pre=&suf=&sa=0) package) (**table S14**). We did not perform equivalent analysis in SEQC data as raw counts are not available for this dataset.

Association between recurrent CN changes and sympathoblastic expression

To compare regions of recurrent CN changes in neuroblastoma to the genomic pattern of expression in reference cell populations we first defined break-point regions and recurrently changed regions. We defined a recurrently changed region as one which had a gain (or a loss) in 200 or more samples (out of 556) [(*20*)](http://f1000.com/work/citation?ids=8688170&pre=&suf=&sa=0). For regions with a well defined transition region (e.g. chromosome 11 change between recurrent gain and loss), we defined the break-point region to be 2Mb around the transition point.

For sympathoblastic, erythrocytes and chromaffin cells, we then calculated the average expression as a function of genomic position. To do this, we calculated gaussian kernel smoothed density with bandwidth of 200,000 bp and weights equal to the library size normalised expression in each cell type placed at the transcription start site of each gene. This is roughly equivalent to a running mean of normalised expression of each cell type calculated across the genome.

This normalised, smoothed expression is what is plotted in **Fig. 3F, fig. S20-22**. To calculate the significance of the relative abundance of the sympathoblastic and chromaffin we calculate the average difference in the smoothed expression between the two cell types in the regions of interest. To calculate a p-value for the significance of the observed difference in smoothed expression, we compared the observed value to the value in 1000 randomly selected regions of the genome of the same size. We then define the p-value to be the fraction of the randomly selected regions with a difference greater than or equal to the observed one (**fig. S8 and S9**).

Differential expression of cells carrying prognostic CN change

In our 10X data, we identified one sample (GOSH021) that had loss of both chromosome 1p and 11q, which was not present in any of the other tumour samples. To determine the transcriptomic consequences of these prognostically important copy number changes we calculated the differentially expressed genes between those cells with and without the copy number changes.

We limited this analysis to only those cells in the G1 phase of the cell cycle to exclude confounding with differences in proliferation. We then performed a quasi-likelihood F-test using the differential expression testing package edgeR [(*43*)](http://f1000.com/work/citation?ids=673952&pre=&suf=&sa=0) comparing the cells with/without the CN change. This test was performed with default parameters except for estimating the over-dispersion parameter of the negative binomial distribution where we set prior.df=0 to prevent information sharing between genes. This was done as there were sufficiently many cells in each group to robustly estimate genewise over-dispersion without sharing information.

To test if this list of differentially expressed genes related to sympathoblastic cells we identified all markers of sympathoblastic genes with a tf-idf cut-off of 1 [(*28*)](http://f1000.com/work/citation?ids=5172334&pre=&suf=&sa=0). We then performed a hypergeometric test to measure if this list of sympathoblastic marker genes was over-represented in the genes differentially expressed between tumour cells with and without the prognostic CN changes.

Risk group stratification for SEQC data

As COG risk group information was not available for the SEQC dataset, we stratified samples into risk groups based on *MYCN* amplification status, age at diagnosis, and disease stage. We defined samples as low risk if age at diagnosis was <18 months and *MYCN* amplification status was negative, excluding 4s stage samples. We defined samples as high risk if age at diagnosis was >18 months and *MYCN* amplification status was positive, excluding 4s stage samples.

Code Availability

We have included the source code used to generate the Figures and Tables presented in this analysis as **Data S1**. The purpose of this code is to provide additional explanation of the analyses described in **Methods**, such as the precise function call and parameter values used.

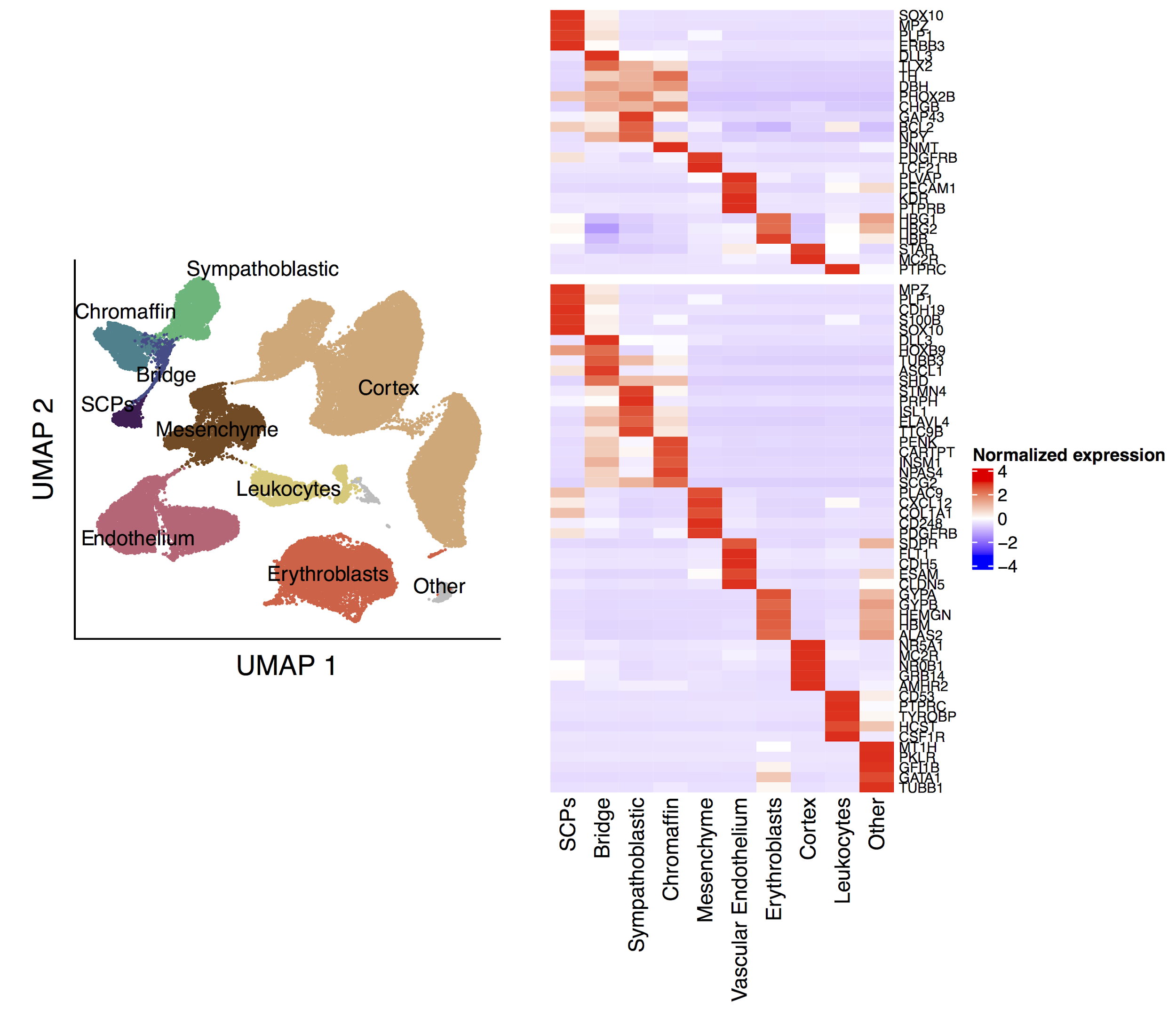

**Figure S1. Marker expression in fetal adrenal gland**

The left panel shows a UMAP ( uniform manifold approximation and projection) of 57,972 cells from the fetal adrenal gland, annotated by cell type. The top right panel shows the expression of canonical markers curated from the literature (see **table S2**) and the bottom right panel shows the expression of the top 5 algorithmically selected marker genes for each cluster (see Methods) in each cell type. Mean expression per cluster is normalized to have a mean of 0 and a standard deviation of 1 for each gene.

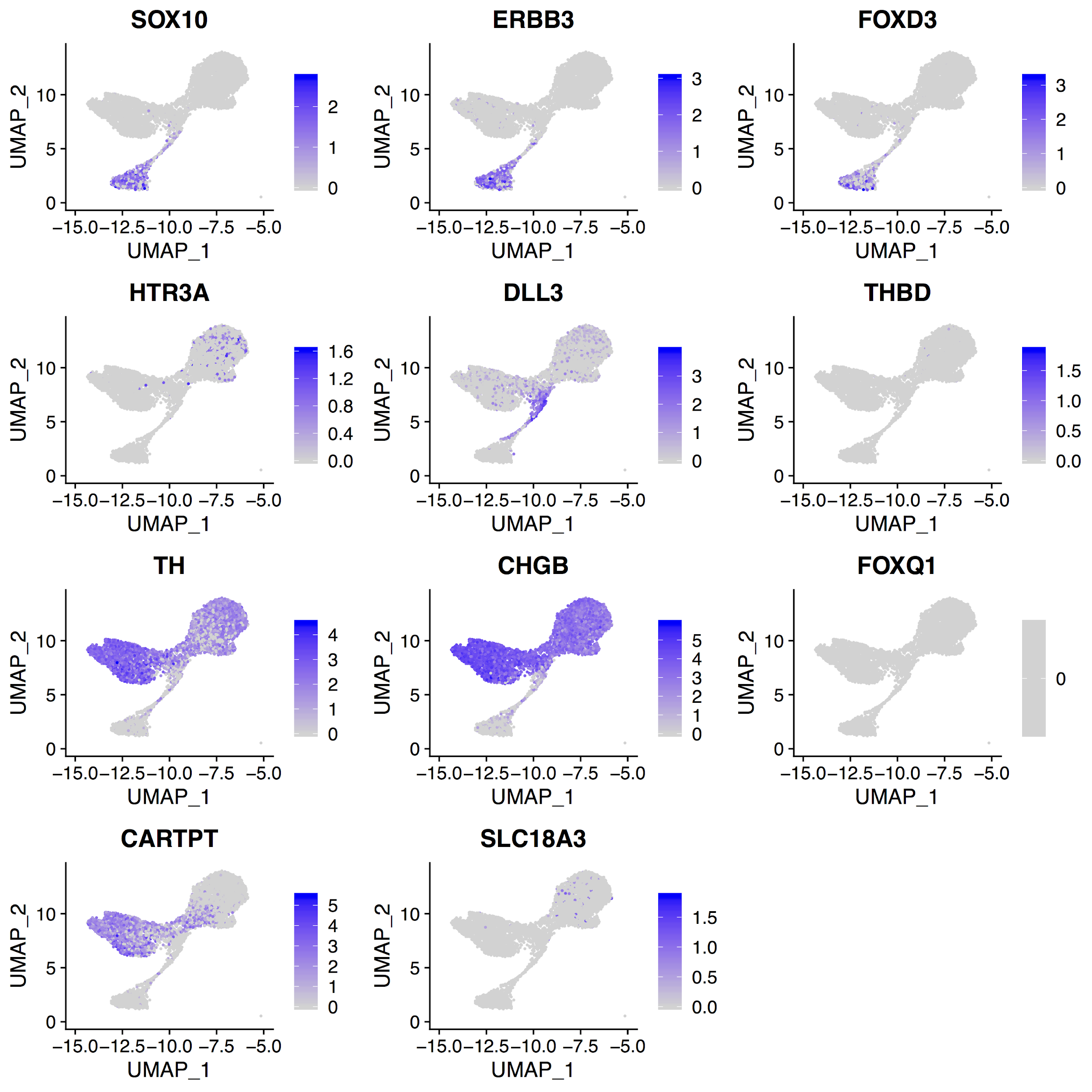

**Figure S2. Murine markers of adrenal medulla in human fetal medulla**

Each panel shows log normalized expression of a murine medullary marker [(*9*)](http://f1000.com/work/citation?ids=3914331&pre=&suf=&sa=0) in human fetal adrenal medulla. In mice, *SOX10*, *ERBB3* and *FOXD3* are SCP marker genes, *HTR3A*, *DLL3* and *THBD* are bridge markers, *TH*, *CHGB* and *FOXQ1* are chromaffin cell markers, and *CARTPT* and *SLC18A3* are sympathoblastic markers.

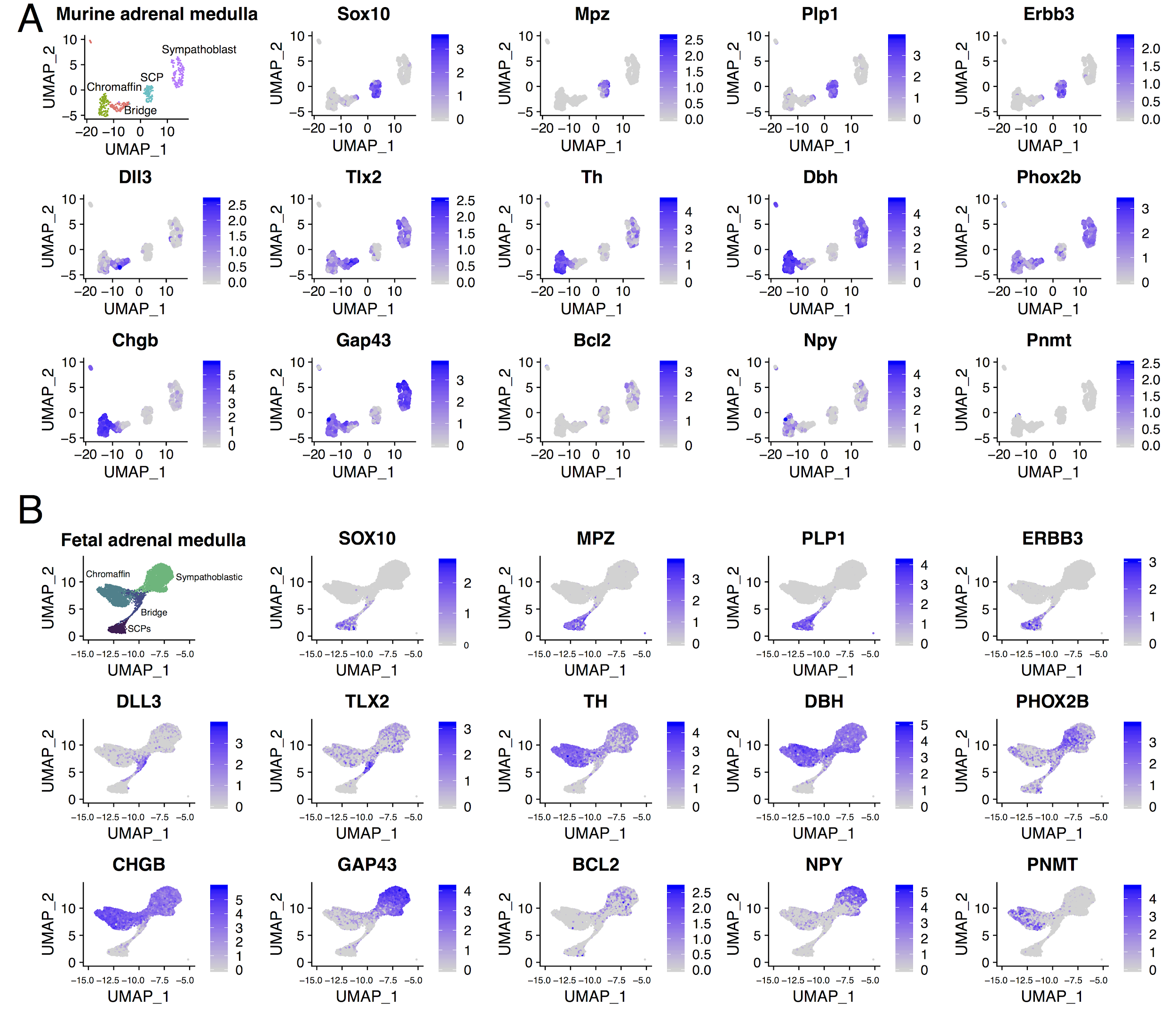

**Figure S3. Literature-defined adrenal medulla markers in murine and human medulla**

Each panel shows a UMAP representation of the log normalised expression of medullary markers curated form literature (**table S2**) in murine (A) and human (B) fetal medullary cells. The first plot in each panel shows the annotation of each cell.

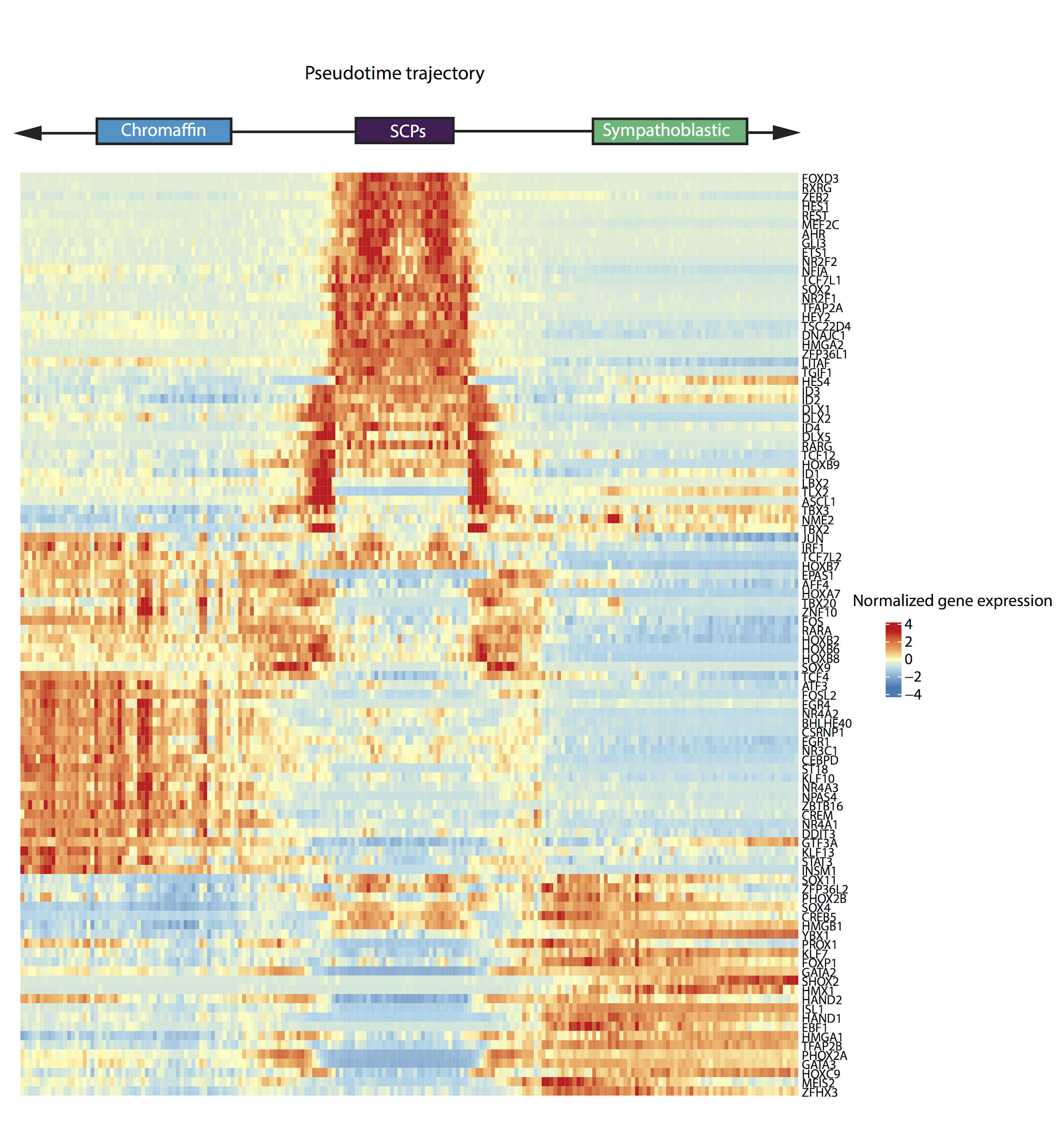

**Figure S4. Differential expression of transcription factors in fetal medulla**

Top 100 differentially expressed genes along single-cell trajectories in the medulla. Cells are ordered by pseudotime and expression is averaged in bins. The center of the heatmap corresponds to early pseudotime (SCPs), and proceeds left (chromaffin) and right (sympathoblast). The expression of each gene along pseudotime was aggregated separately for each trajectory (SCP → Chromaffin and SCP → Sympathoblast), merged, and row normalized to have a mean of 0 and a standard deviation of 1.

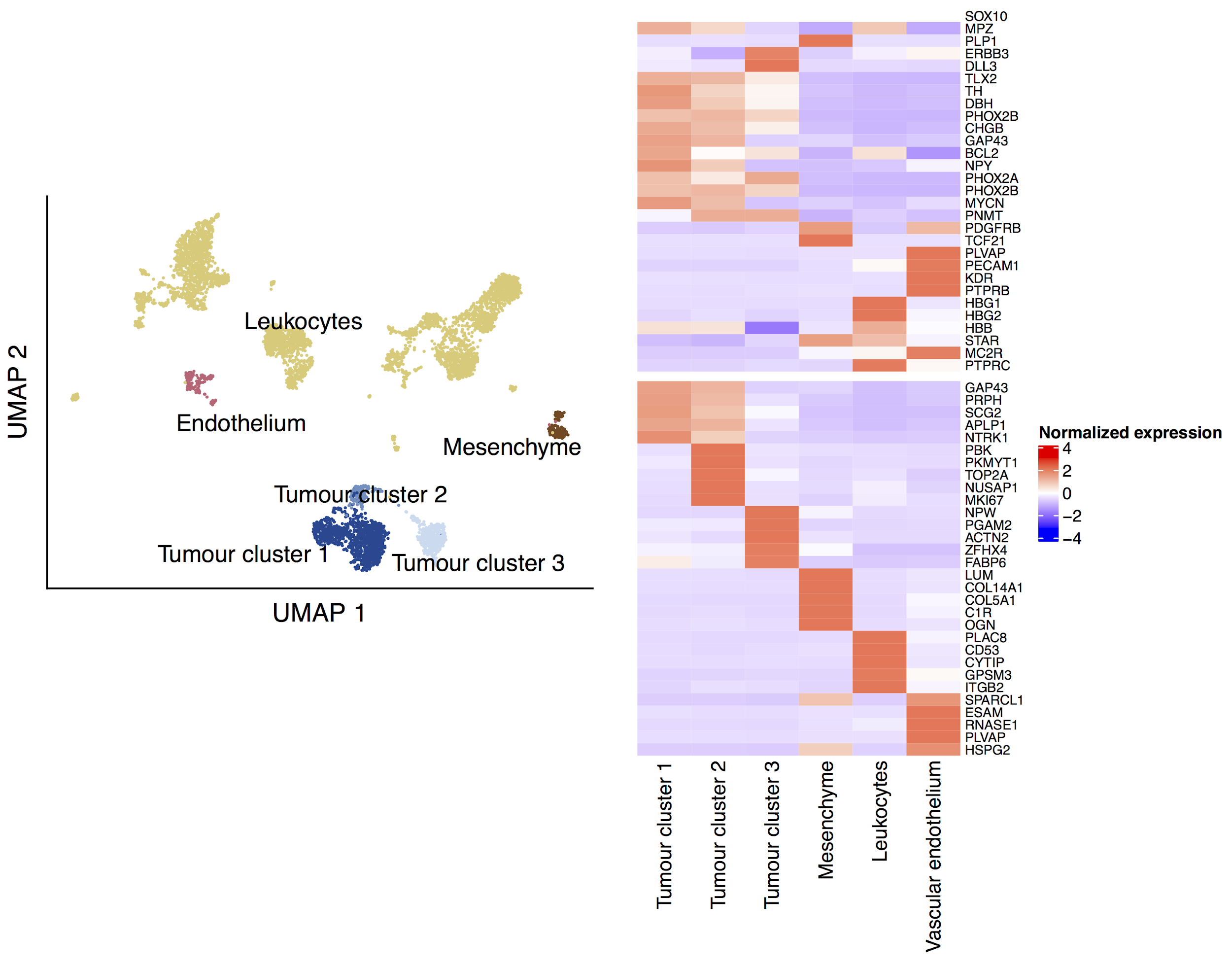

**Figure S5. Marker expression in 10x tumour data**

The left panel shows a UMAP ( uniform manifold approximation and projection) of the 6,442 cells in the neuroblastoma 10X dataset, annotated by cell type. The top right panel shows the expression of canonical markers curated from the literature (see **table S2**) and the bottom right panel shows the expression of the top 5 algorithmically selected marker genes for each cluster (see Methods) in each cell type. Mean expression per cluster is normalized to have a mean of 0 and a standard deviation of 1 for each gene.

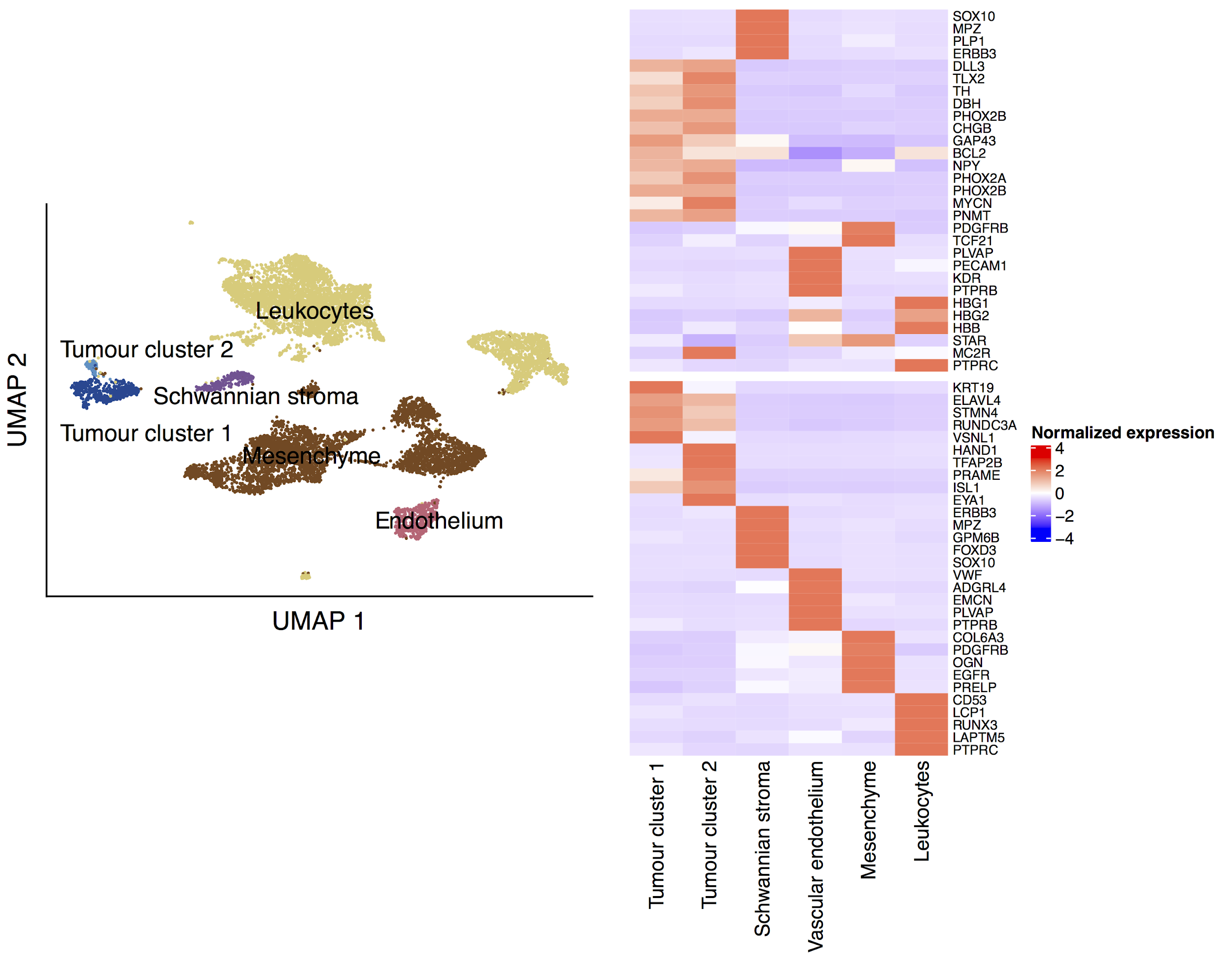

**Figure S6. Marker expression in CEL-seq2 tumour data**

The left panel shows a UMAP ( uniform manifold approximation and projection) of the 10,149 cells in the neuroblastoma CEL-seq2 dataset, annotated by cell type. The top right panel shows the expression of canonical markers curated from the literature (see **table S2**) and the bottom right panel shows the expression of the top 5 algorithmically selected marker genes for each cluster (see Methods) in each cell type. Mean expression per cluster is normalized to have a mean of 0 and a standard deviation of 1 for each gene.

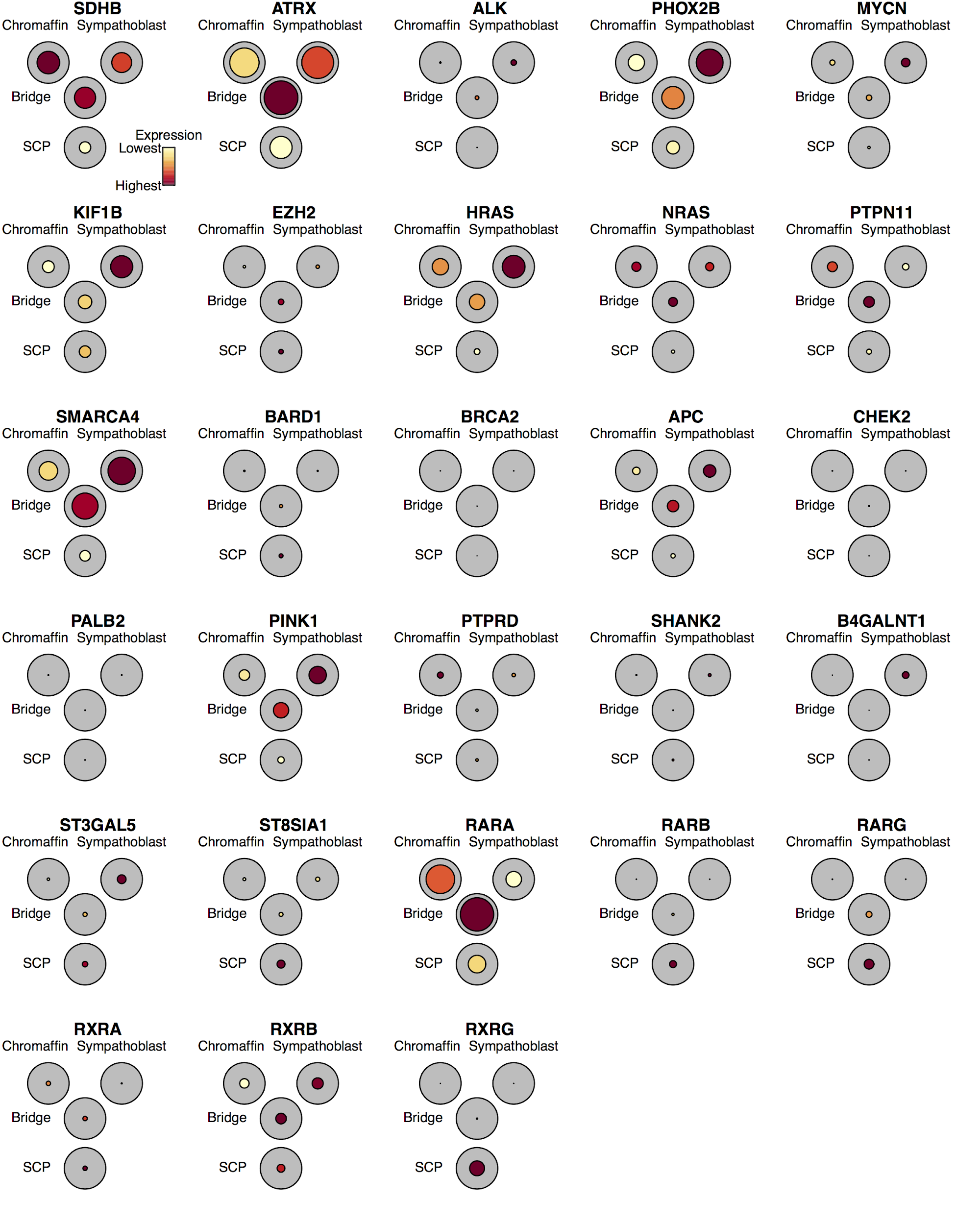

**Figure S7. Neuroblastoma genes in fetal adrenal medulla**

The black circles show a reduced resolution representation of the trajectory in **Fig. 1C** with the size of the coloured circle indicating the fraction of cells expressing each gene in that region of the trajectory and the colour indicating average expression, normalised so that all genes have the same maximum and minimum.

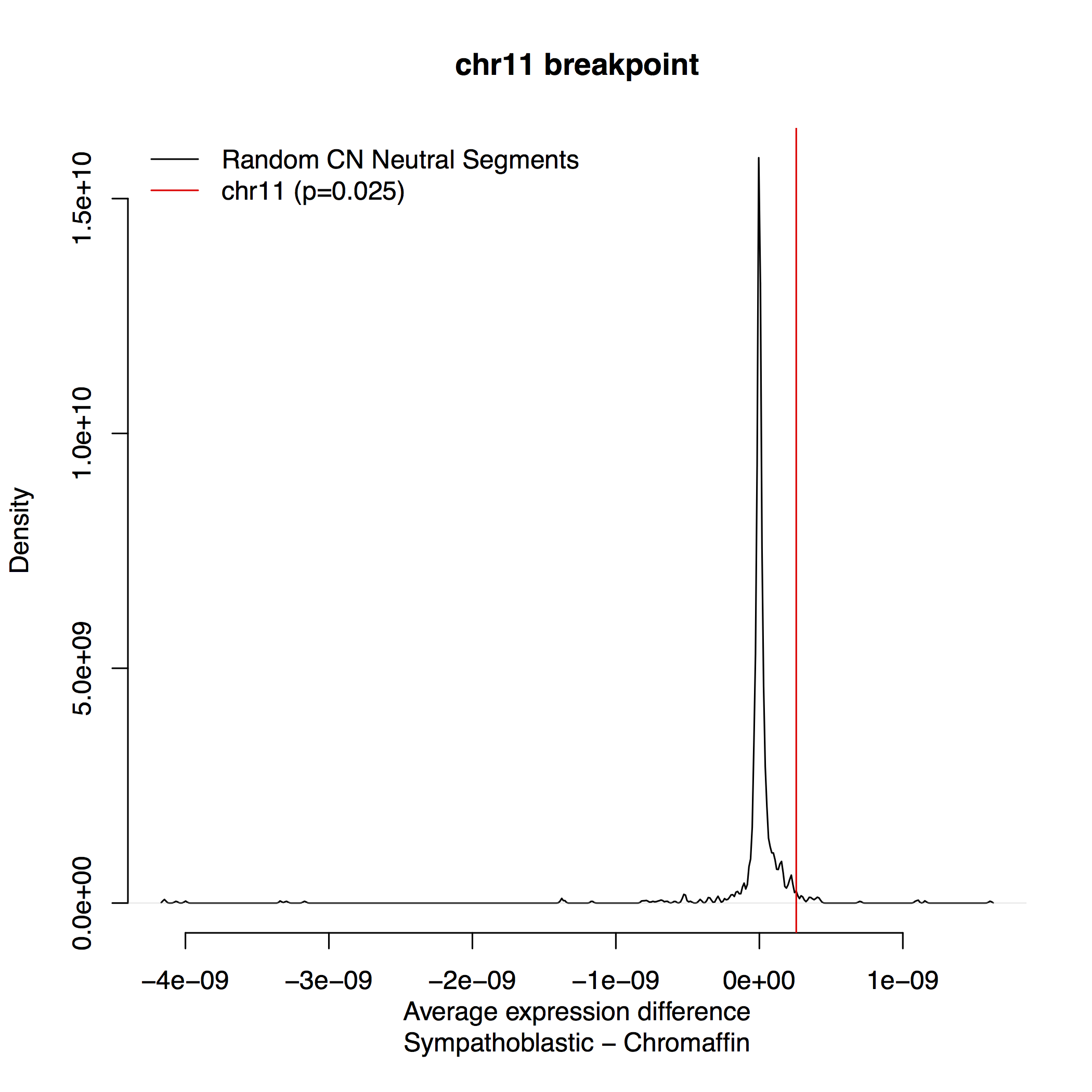

**Figure S8.**

Frequency of different average expression differences between sympathoblastic and chromaffin cells in randomly chosen genomic regions of the same size as the region surrounding the recurrent copy number boundary on chr 11 (**Fig. 3F**). The x-axis shows the expression difference and the y-axis shows the frequency of each expression difference, calculated using kernel density smoothing with automatically determined bandwidth. The red line shows the observed expression difference in the region on chr11 and the fraction of regions with a value more extreme than this is shown in parenthesis.

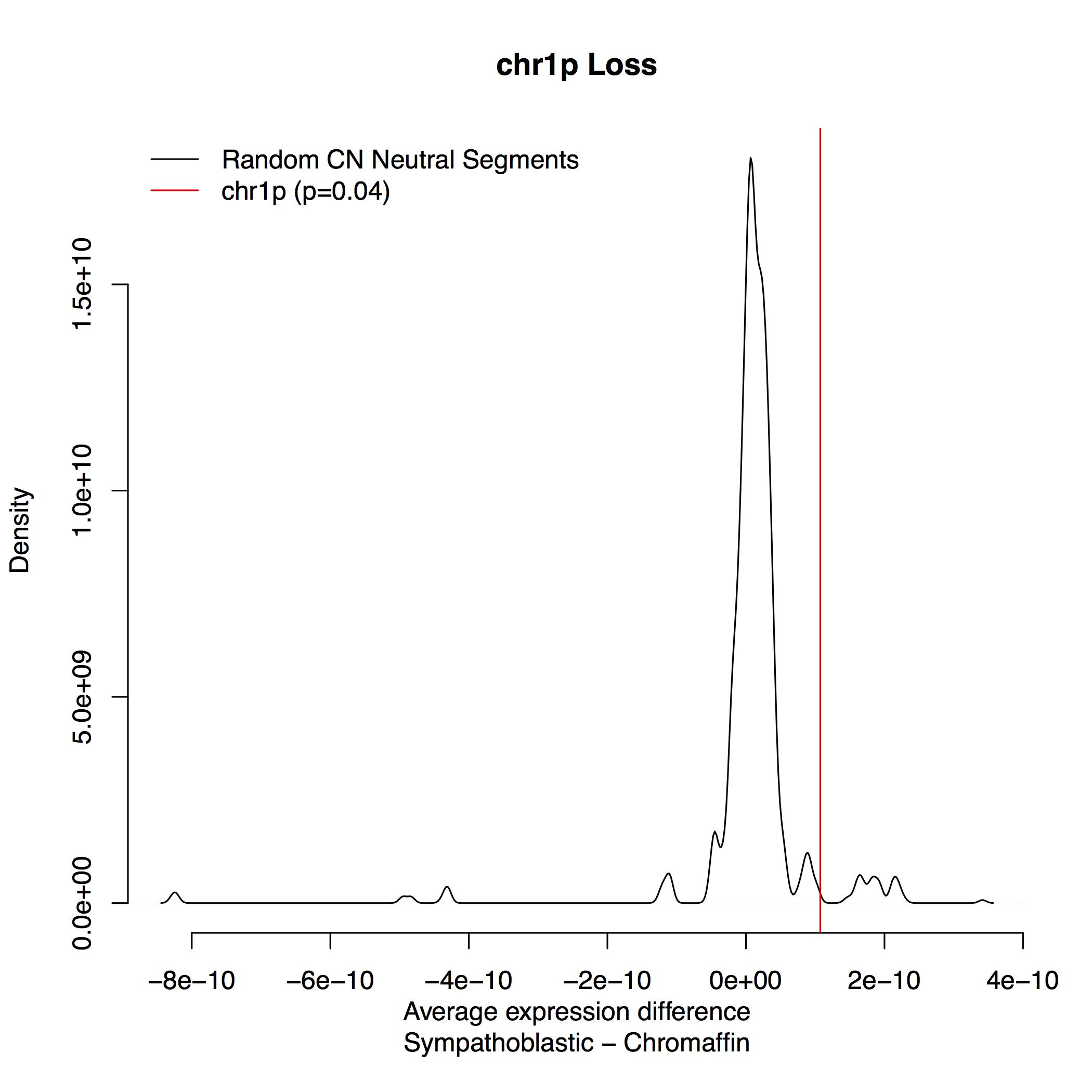

**Figure S9.**

Frequency of different average expression differences between sympathoblastic and chromaffin cells in randomly chosen genomic regions of the same size as the recurrent copy number loss on chr1 (**Fig. 3F**). The x-axis shows the expression difference and the y-axis shows the frequency of each expression difference, calculated using kernel density smoothing with automatically determined bandwidth. The red line shows the observed expression difference in the region on chr1 and the fraction of regions with a value more extreme than this is shown in parenthesis.

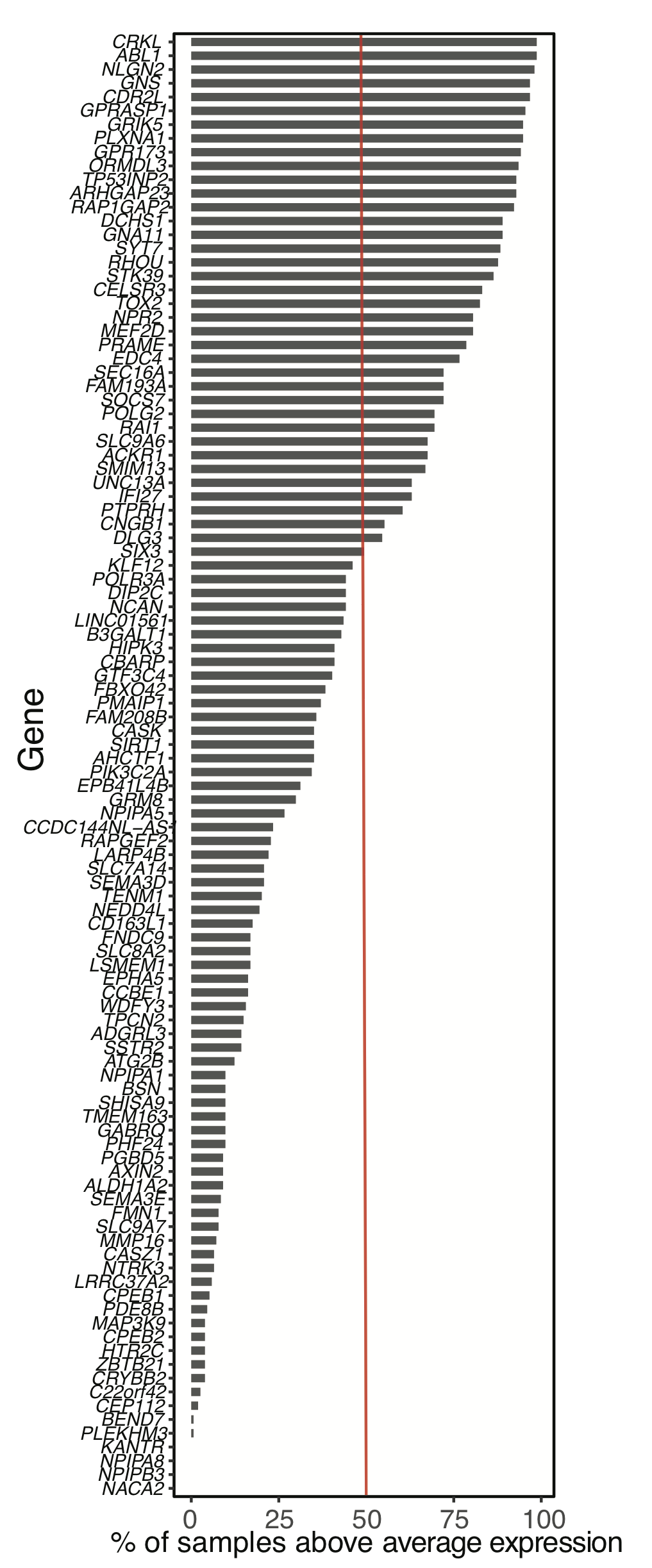

**Figure S10. Table S5 genes in TARGET**

Fraction of sample with above average expression (see Methods) of genes differentially expressed between adrenal medulla and tumour cells (**table S5**). Genes with bars to the right of the red line are expressed in the majority of samples in the TARGET dataset.

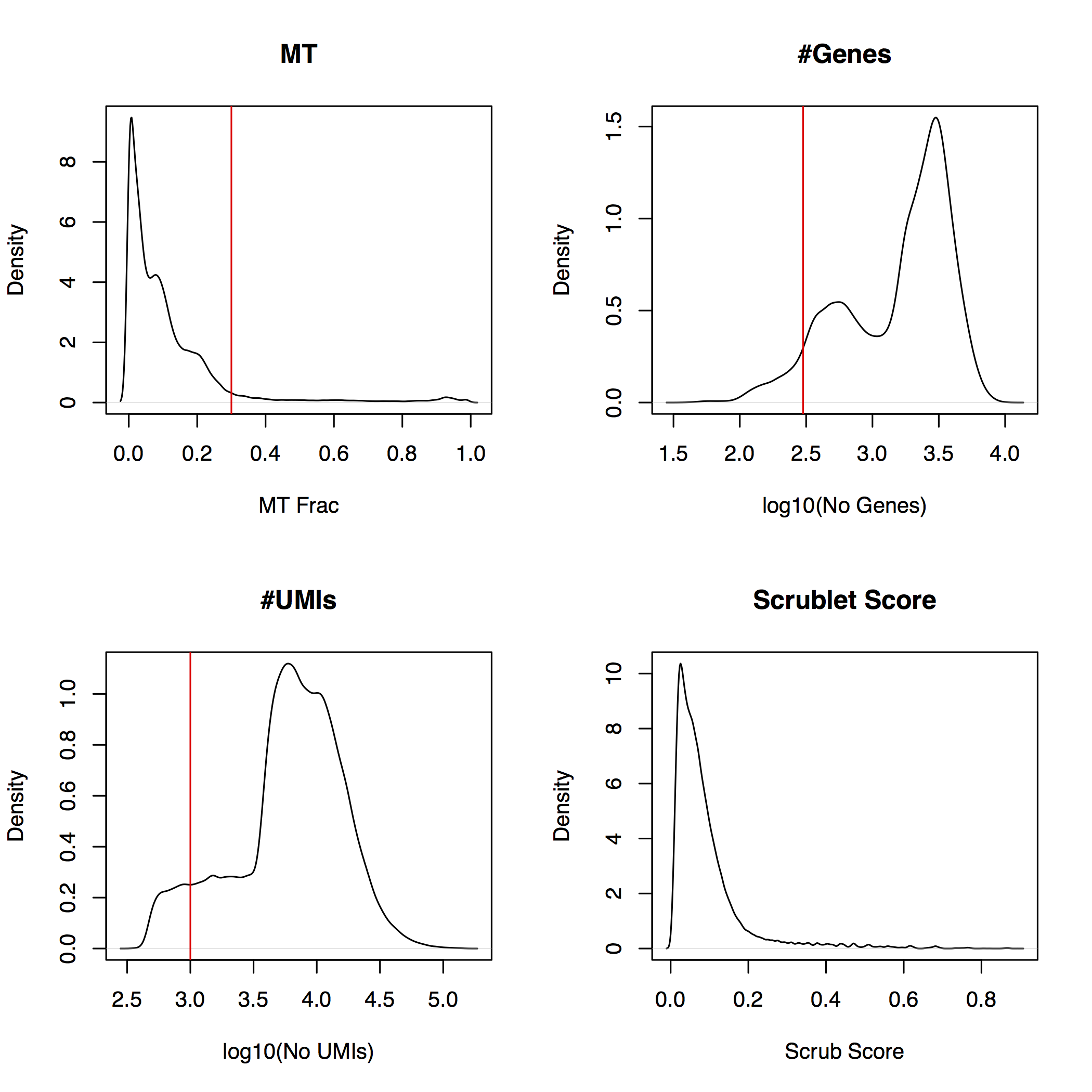
 **Figure S11. QC plots for fetal adrenal data**

Density plots of mitochondrial fraction, no. of Genes, no. of UMIs and scrublet score for all cells in the fetal adrenal dataset. Red lines indicate the cut-off used to filter the data.

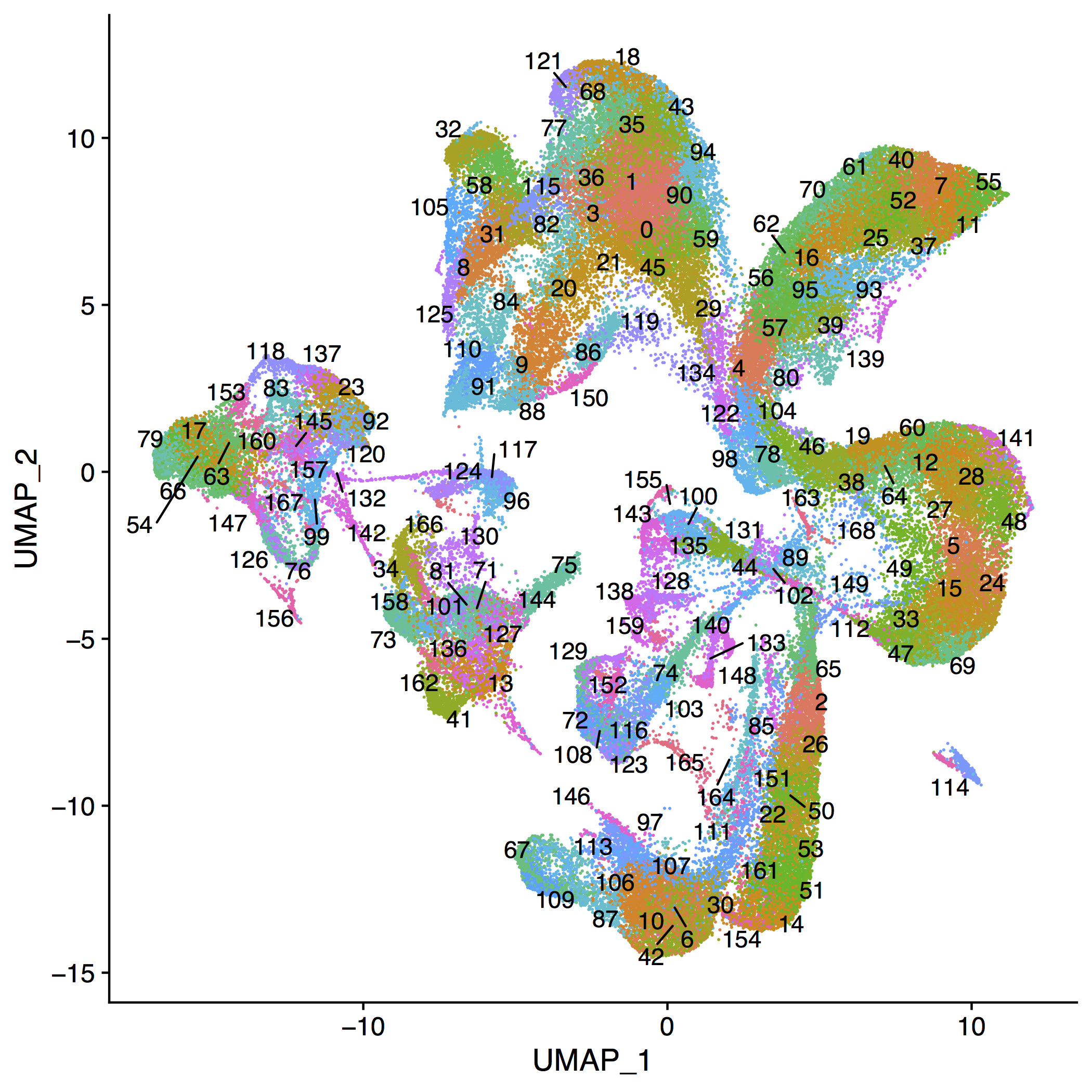

**Figure S12. - High resolution clustering of pre-QC fetal adrenal**

UMAP representation of the fetal adrenal cells before QC filters showing the high resolution clustering. Each cluster is coloured differently and a numeric label indicates the mean UMAP position.

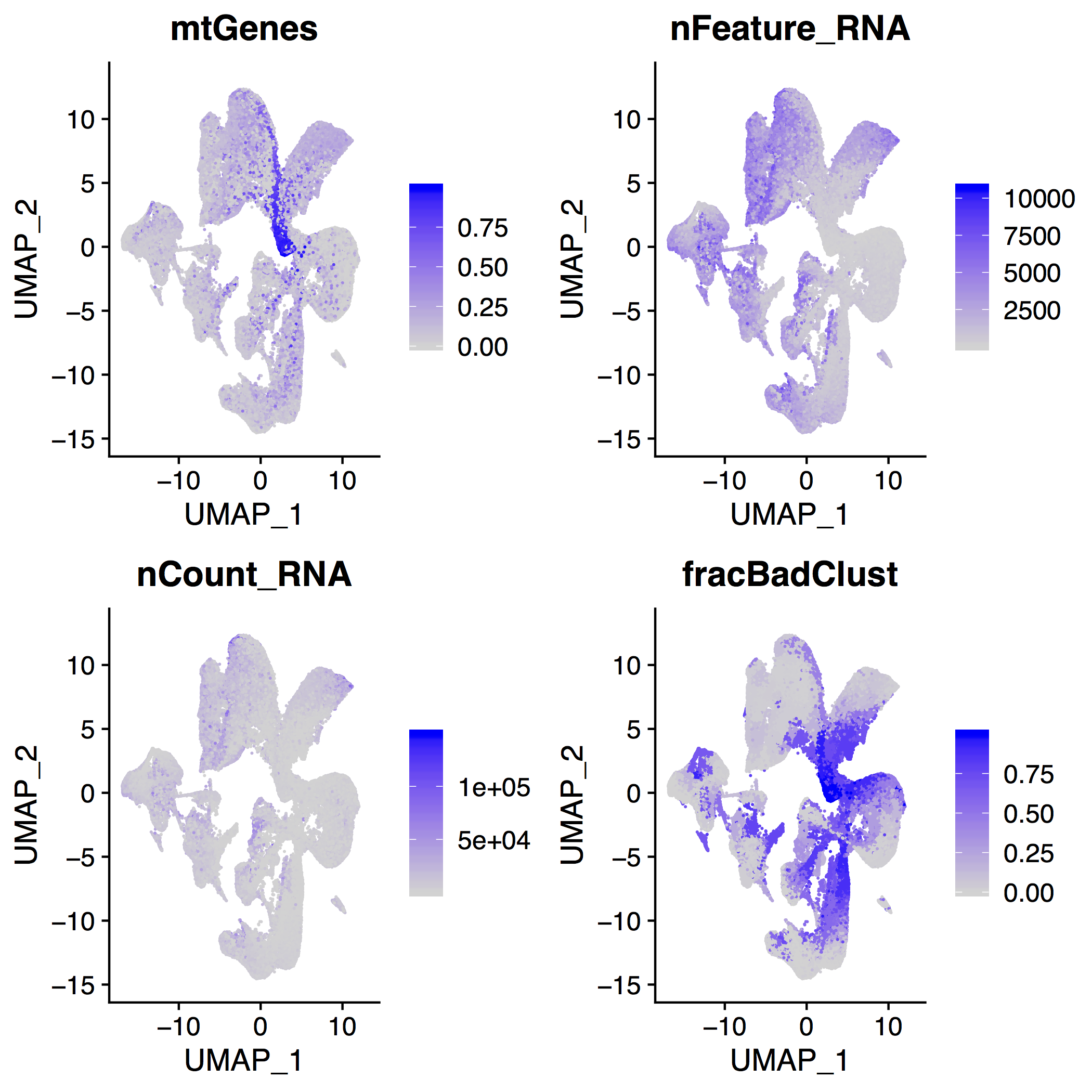

**Figure S13. - Distribution of key QC values in fetal adrenal**

UMAP representation of the fetal adrenal cells before QC filters showing the values of key QC values. Each cell is coloured by the value of the QC property indicated in the panel title, which are MT gene fraction for “mtGenes”, number of genes detected for “nFeature_RNA”, number of UMIs detected for “nCount_RNA”, and fraction of cells in the clusters shown in **fig. S12** that fail QC for “fracBadClust”.

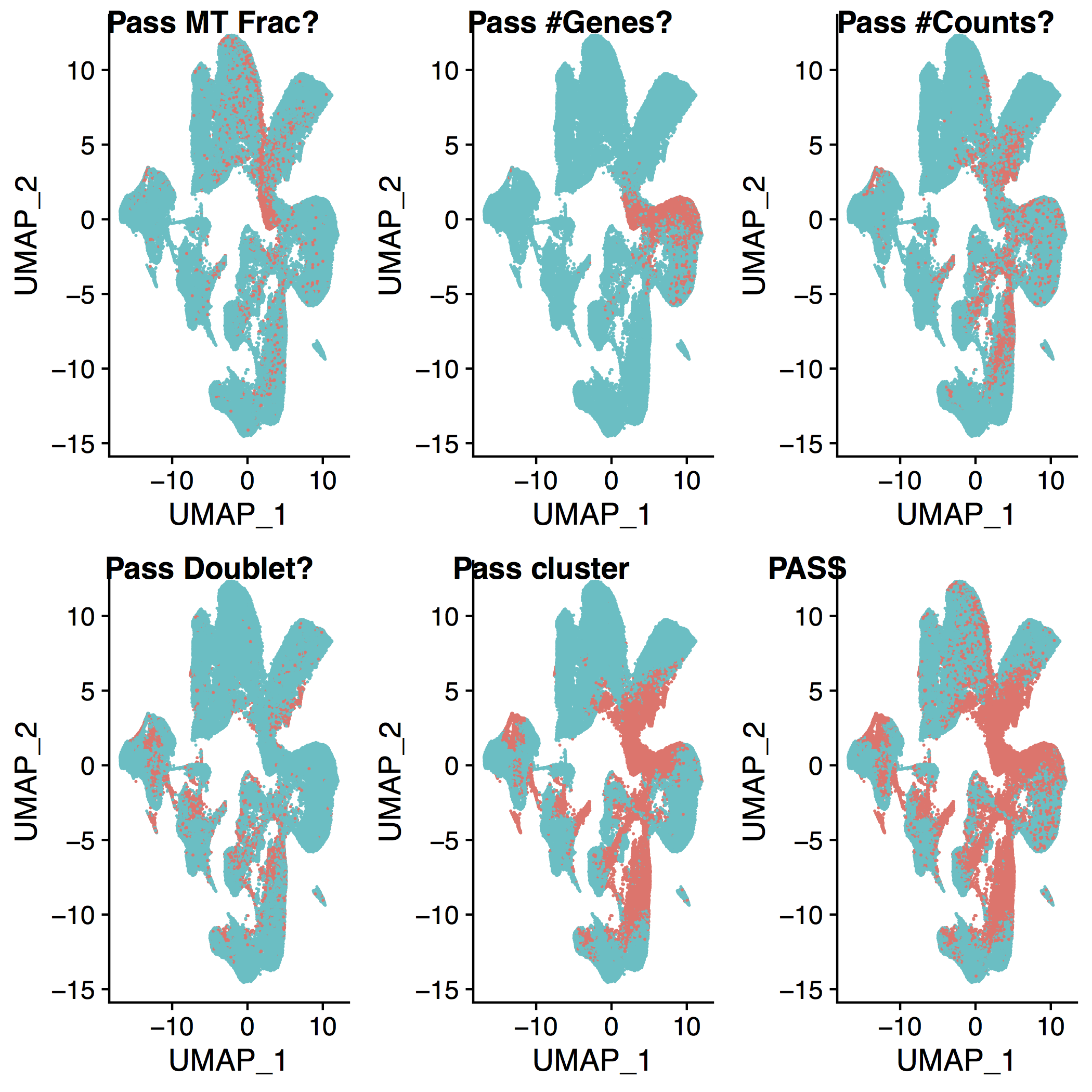

**Figure S14. - Which cells pass QC filters for fetal adrenal.**

UMAP representation of the fetal adrenal cells before QC filters showing which cells pass which QC filters. Cells that pass (fail) QC filters are shown in green (red). Panels show cells that pass/fail from left to right, top to bottom: MT fraction, minimum number of genes, minimum number of counts, doublet detection, do not belong to a bad cluster, and all filters.

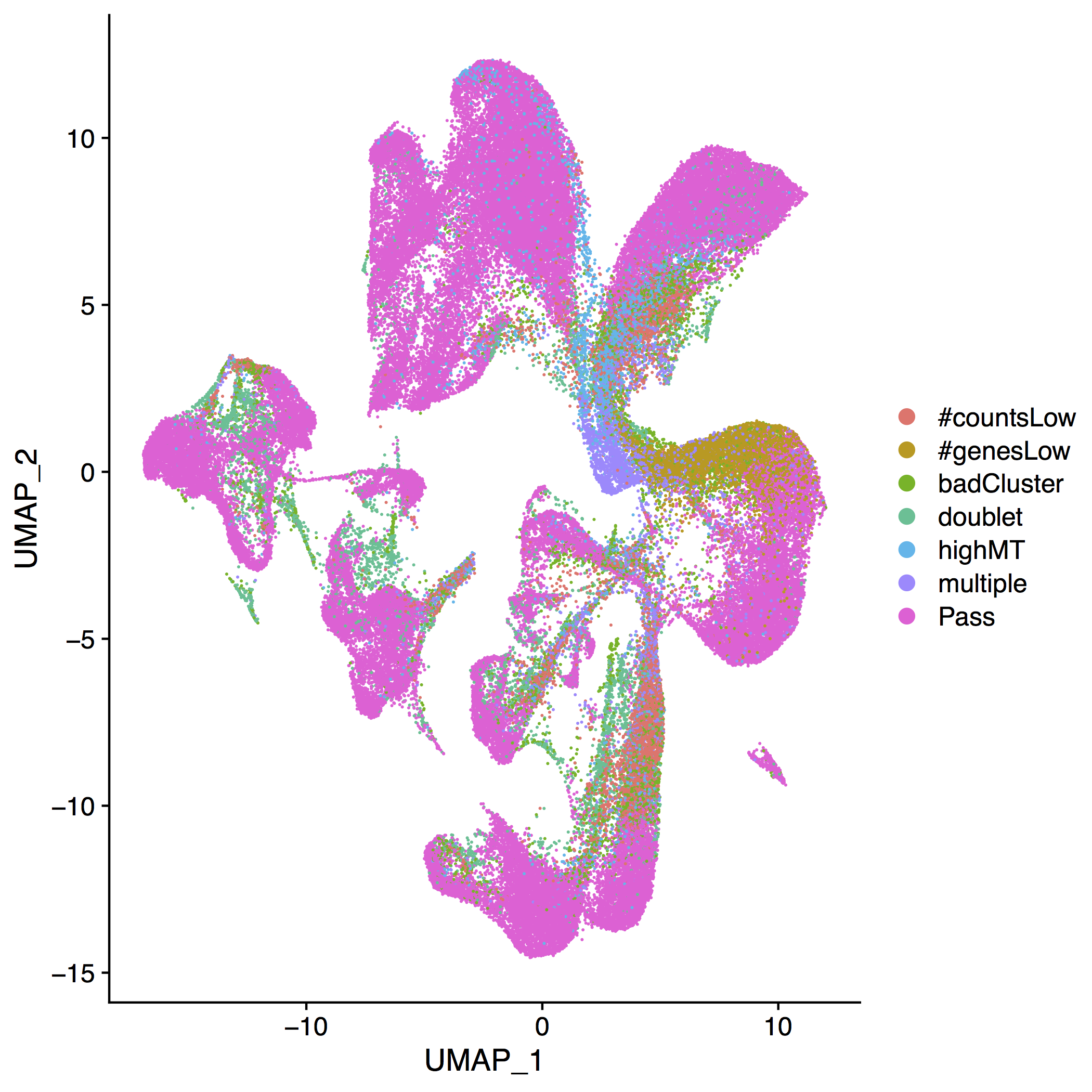

**Figure S15. - Which cells pass which QC filters for fetal adrenal**

UMAP representation of the fetal adrenal cells before QC filters showing which cells pass which QC filters. Cells are coloured by the reason they have failed (or passed) QC. Those cells marked as “badCluster” have passed all QC filters but belong to a cluster which has more than 50% failed cells. Those that are marked as “multiple” have failed more than one QC filter.

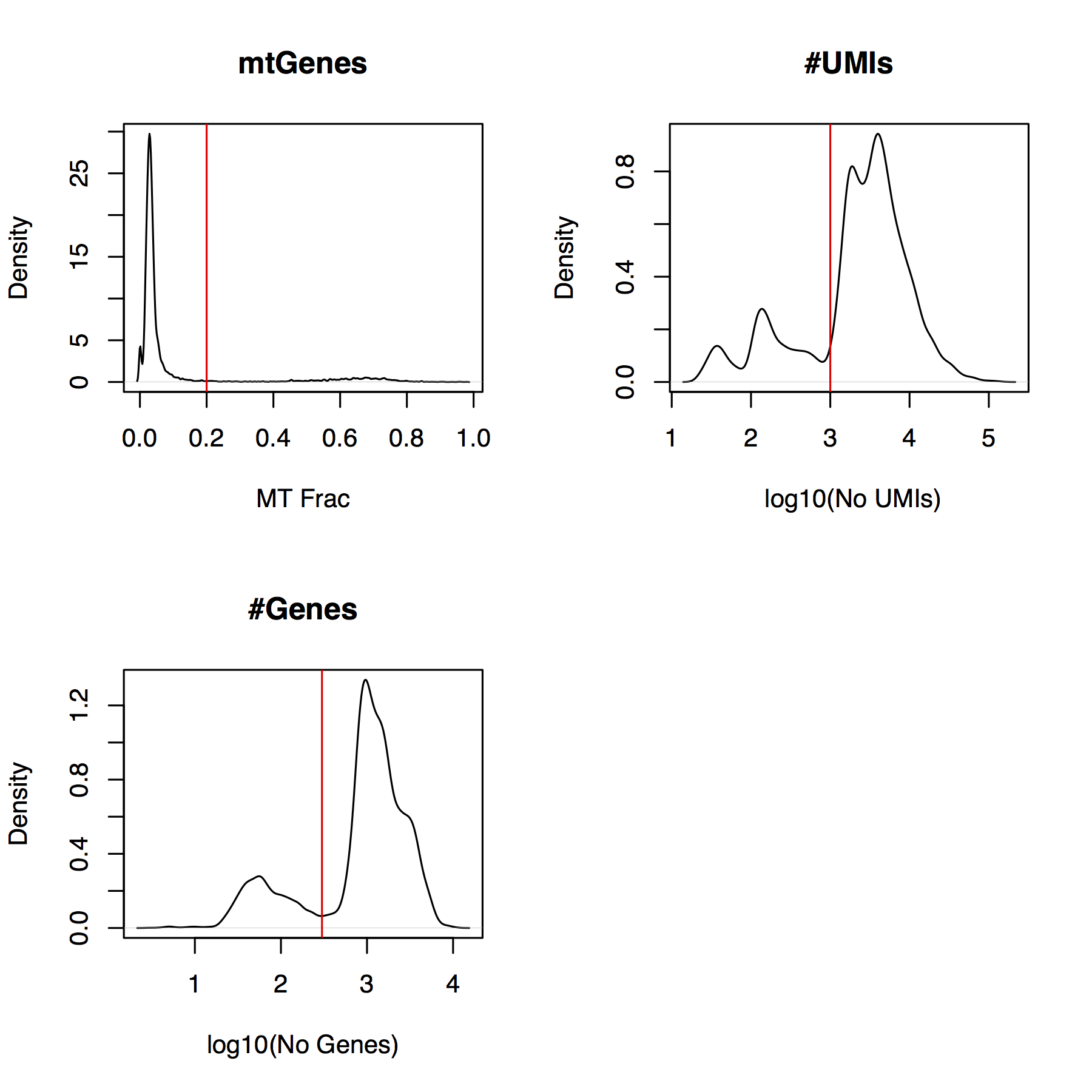

**Figure S16. QC plots for 10x tumour data**

Density plots of mitochondrial fraction, no. of Genes and no. of UMIs for all cells in the 10x tumour dataset. Red lines indicate the cut-off used to filter the data.

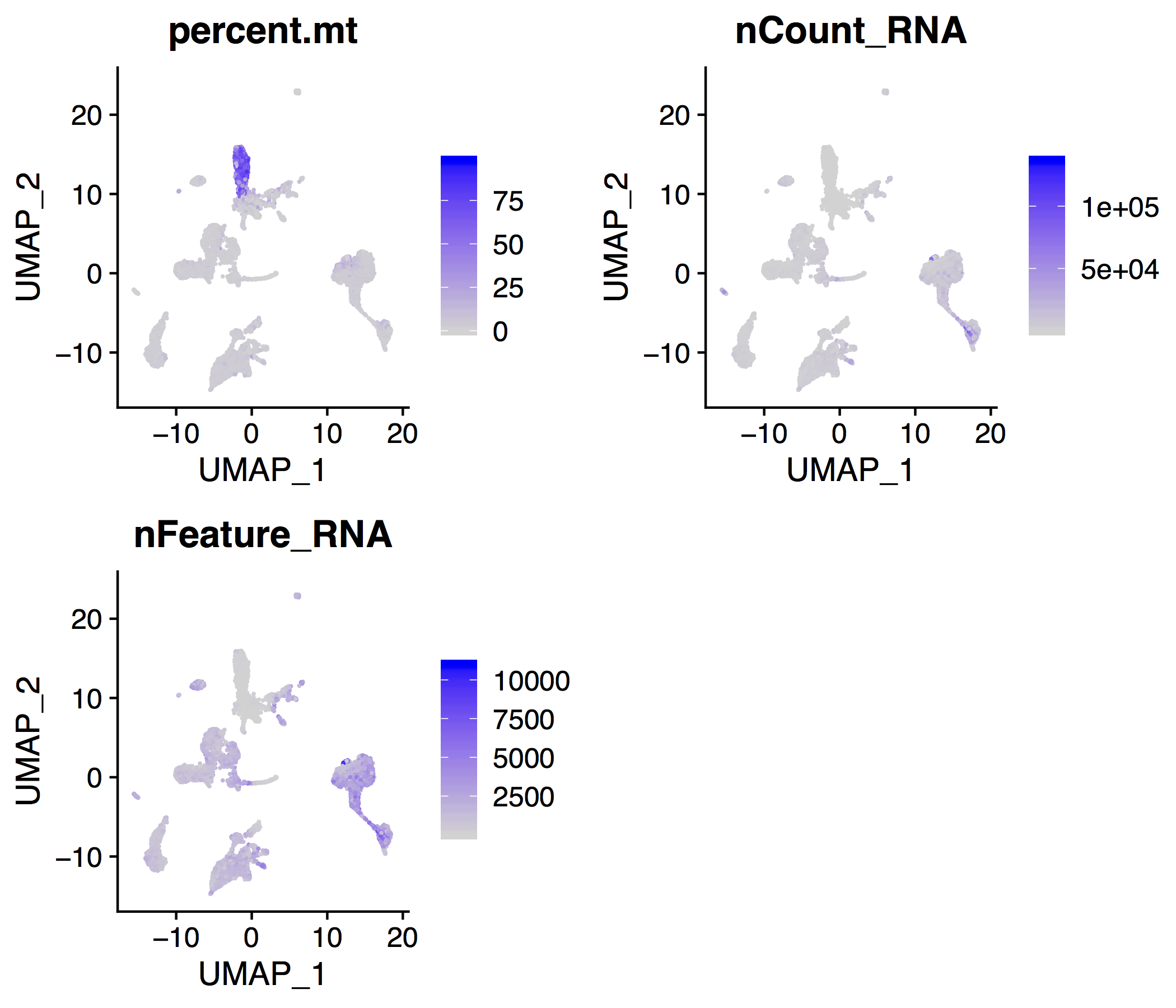

**Figure S17. Feature plots for 10x tumour data**

Plots showing percent of mitochondrial genes, number of UMIs and number of genes in each cell in the 10x tumour dataset.

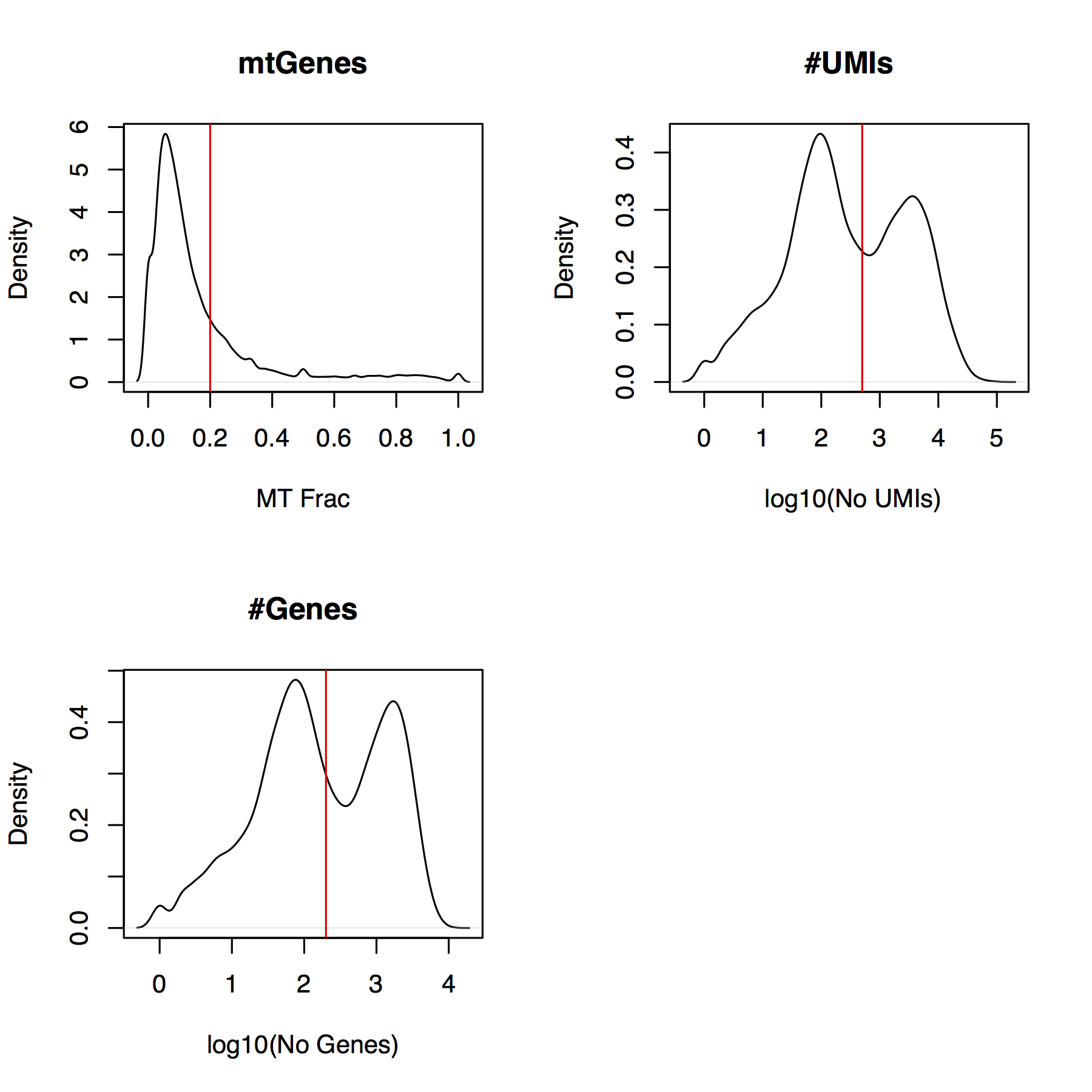

**Figure S18. QC plots for CEL-seq2 tumour data**

Density plots of mitochondrial fraction, no. of Genes and no. of UMIs for all cells in the CEL-seq2 tumour dataset. Red lines indicate the cut-off used to filter the data.

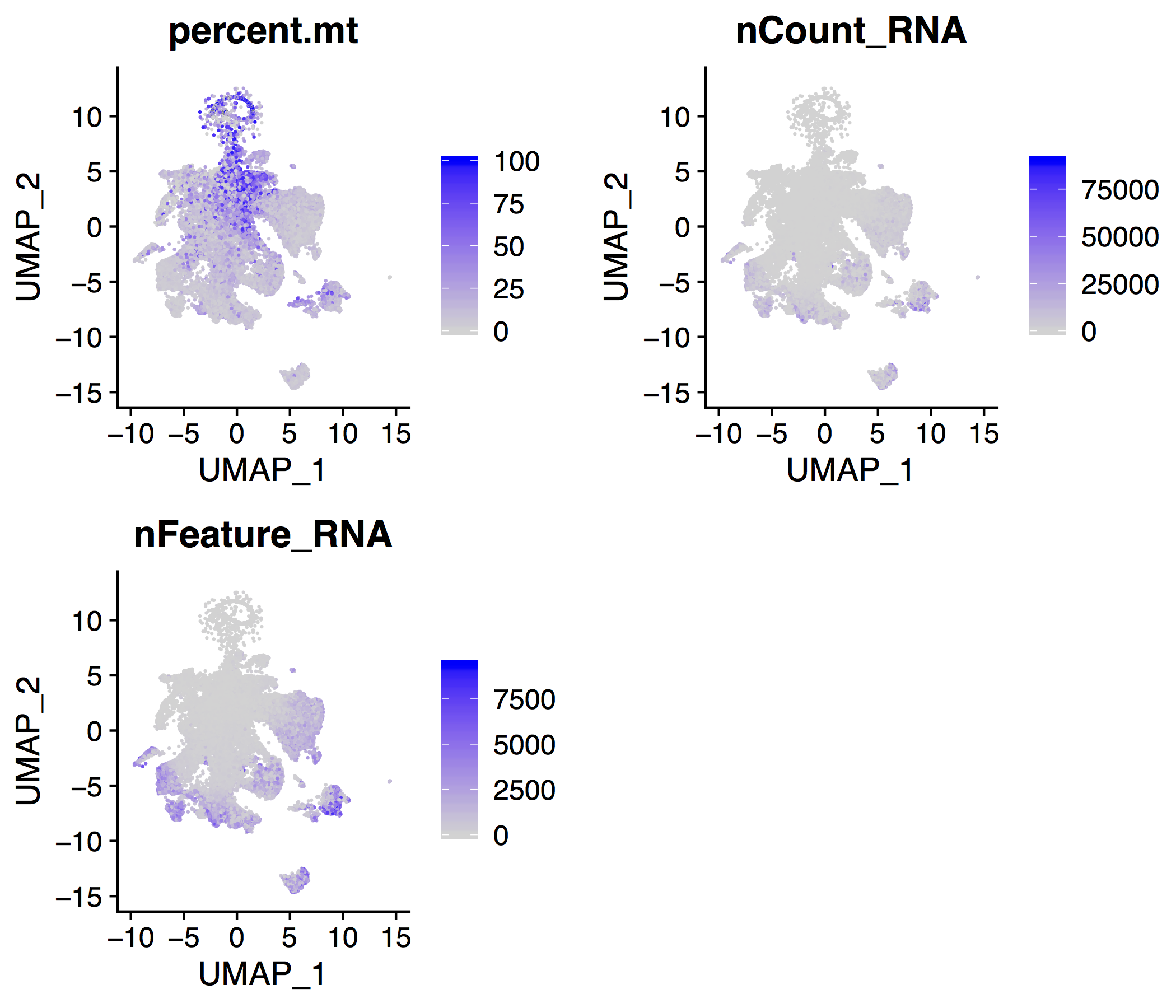

**Figure S19. Feature plots for CEL-seq2 tumour data**

Plots showing percent of mitochondrial genes, number of UMIs and number of genes in each cell in the CEL-seq2 tumour dataset.

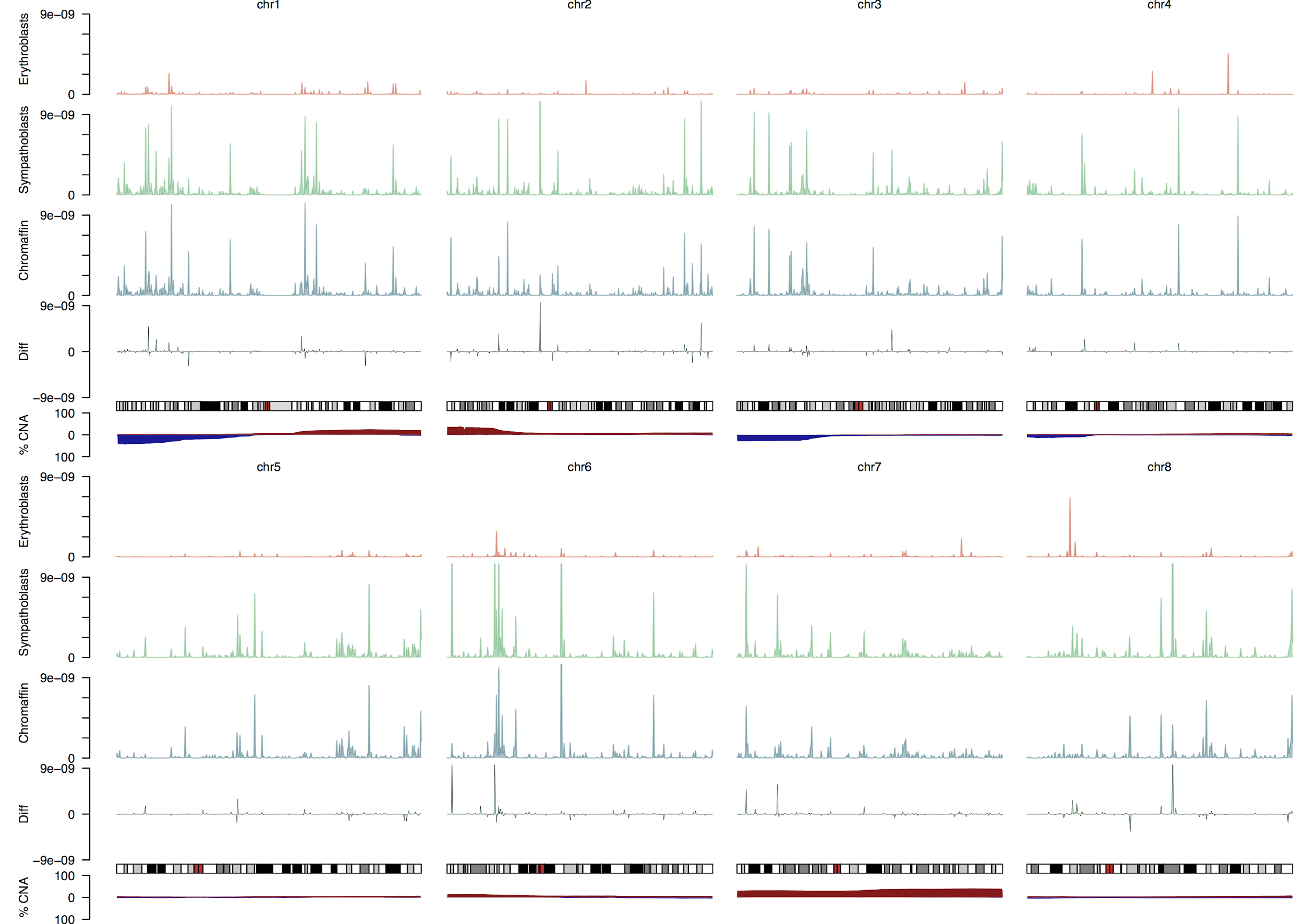

**Figure S20. - Average expression and recurrent CN changes chr 1-8**

Each panel shows the averaged expression (as in **Fig. 3F**) of Erythrocytes (top track), Sympathoblastic cells (second track), Chromaffin cells (third track), and the difference between sympathoblastic expression and chromaffin expression (fourth track). Below these is an ideogram of the chromosome and the bottom track shows the fraction of samples that can contain a copy number loss (blue area below axis) or gain (red area above axis). Expression values are calculated as described in the methods.

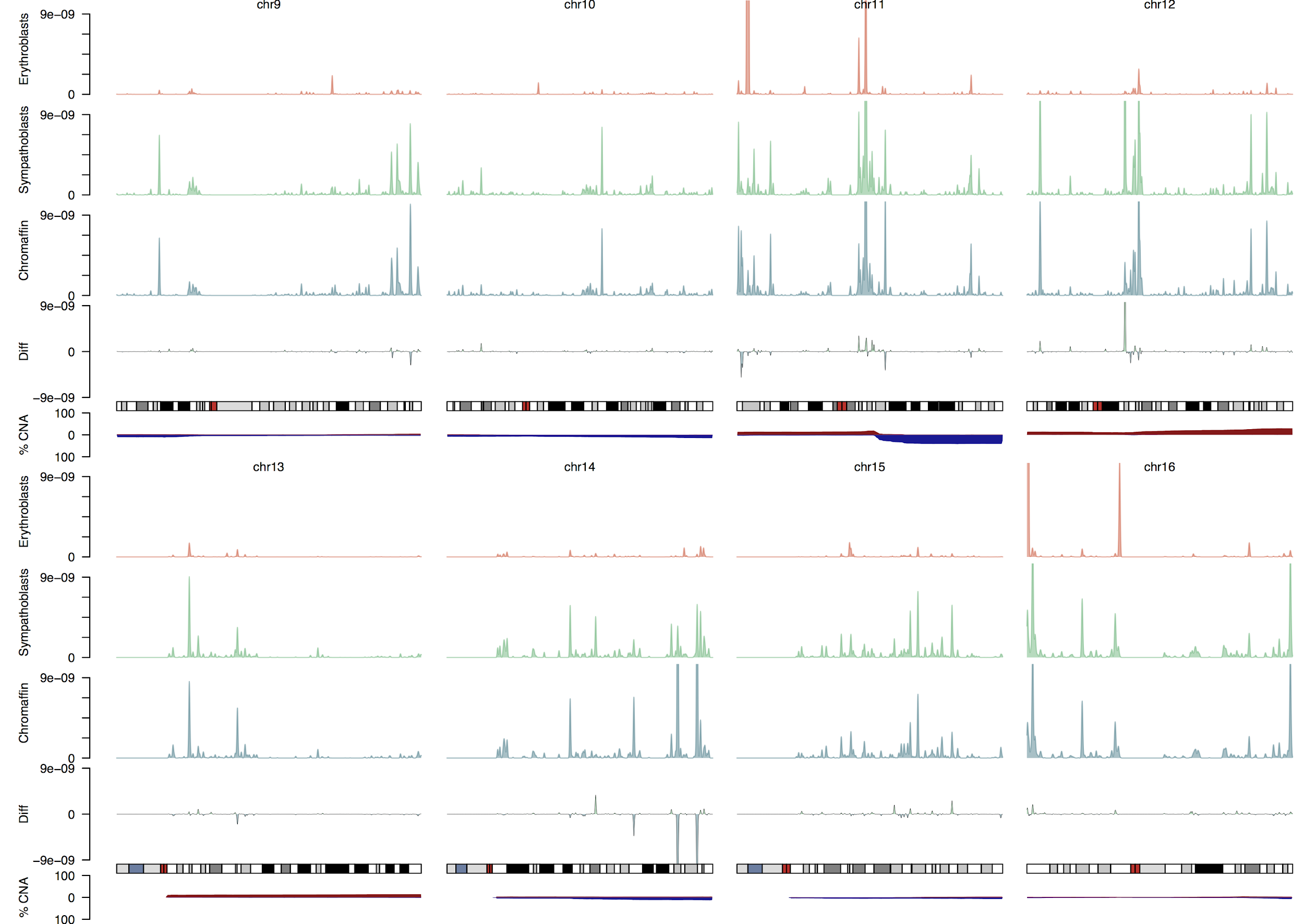

**Figure S21. - Average expression and recurrent CN changes chr 9-16**

Each panel shows the averaged expression (as in **Fig. 3F**) of Erythrocytes (top track), Sympathoblastic cells (second track), Chromaffin cells (third track), and the difference between sympathoblastic expression and chromaffin expression (fourth track). Below these is an ideogram of the chromosome and the bottom track shows the fraction of samples that can contain a copy number loss (blue area below axis) or gain (red area above axis). Expression values are calculated as described in the methods.

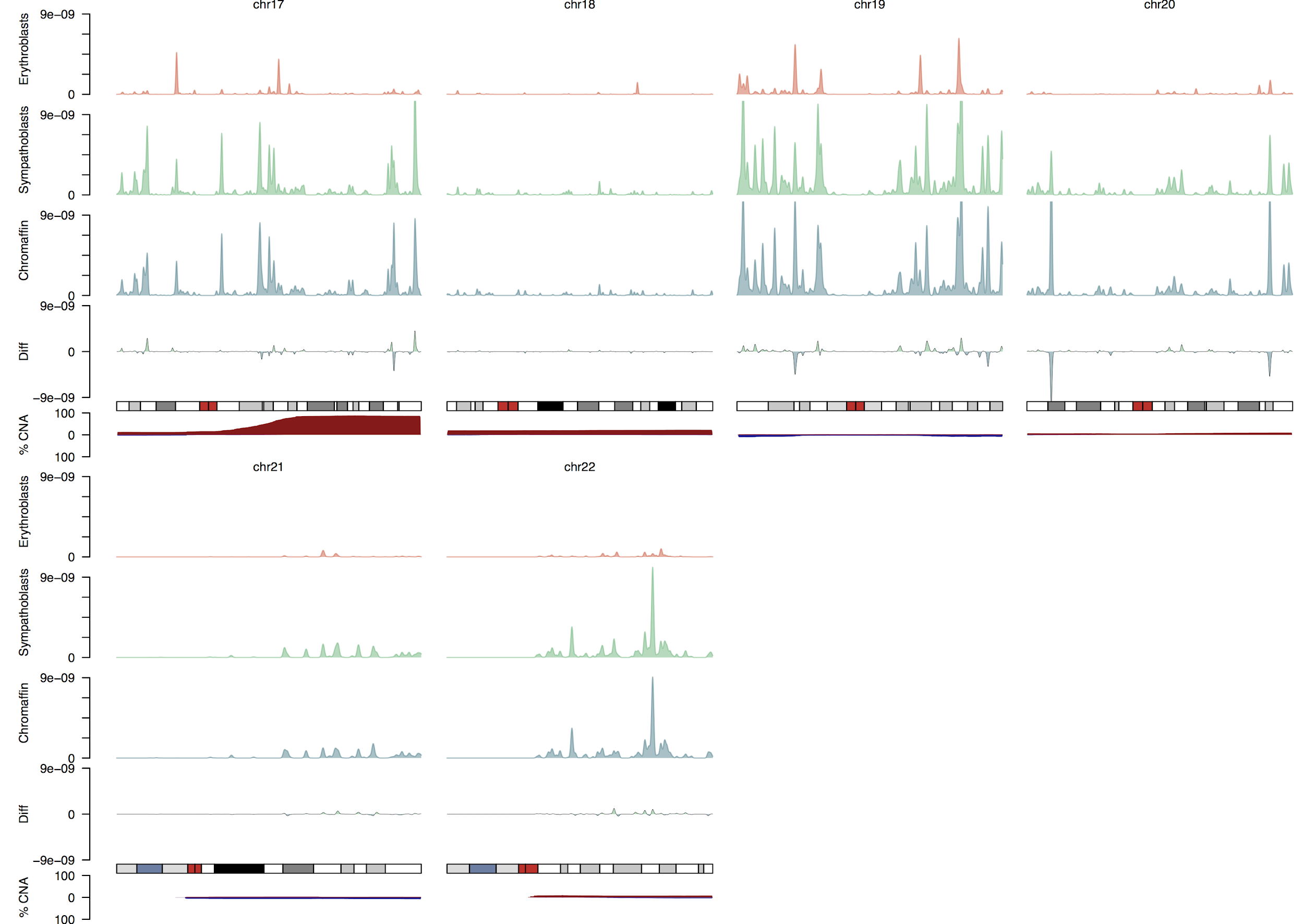

**Figure S22. - Average expression and recurrent CN changes chr 17-22**

Each panel shows the averaged expression (as in **Fig. 3F**) of Erythrocytes (top track), Sympathoblastic cells (second track), Chromaffin cells (third track), and the difference between sympathoblastic expression and chromaffin expression (fourth track). Below these is an ideogram of the chromosome and the bottom track shows the fraction of samples that can contain a copy number loss (blue area below axis) or gain (red area above axis). Expression values are calculated as described in the methods.

| **Age (Gestation weeks)** | **SCPs** | **Bridge** | **Chromaffin** | **Sympathoblastic** | **Medulla (total)** | **Cortex** | **Endothelium** | **Mesenchyme** | **Leukocytes** | **Erythroblasts** | **Other** | **Total** |
| --- | --- | --- | --- | --- | --- | --- | --- | --- | --- | --- | --- | --- |
| 8 | 3 | 81 | 8 | 40 | 132 | 2420 | 232 | 664 | 32 | 14 | 104 | 3598 |
| 8w6d | 6 | 495 | 23 | 4 | 528 | 1471 | 55 | 643 | 48 | 2 | 282 | 3029 |
| 10w5d | 112 | 32 | 187 | 412 | 743 | 7150 | 1771 | 1093 | 374 | 3245 | 130 | 14506 |
| 10w5d | 271 | 26 | 1774 | 323 | 2394 | 4263 | 1135 | 1165 | 351 | 2766 | 88 | 12162 |
| 11 | 194 | 126 | 890 | 1253 | 2463 | 2605 | 1351 | 664 | 452 | 190 | 47 | 7772 |
| 21 | 34 | 2 | 63 | 19 | 118 | 3731 | 1441 | 217 | 613 | 1291 | 43 | 7454 |
| 21 | 21 | 1 | 49 | 2 | 73 | 5912 | 2169 | 297 | 568 | 412 | 20 | 9451 |
| **Total** | 641 | 763 | 2994 | 2053 | 6451 | 27552 | 8154 | 4743 | 2438 | 7920 | 714 | 57972 |

**Table S1. List of fetal adrenal samples and cell type contribution**

Age is given in gestational weeks. Cell numbers are shown for each cell type within the medulla as well as the total number of medullary cells and all other cell types present in the data. The final row gives the total count of each cell type across all samples.

| **Gene** | **Marker of** | **Reference** |
| --- | --- | --- |
| PLVAP | Vascular | [(*44*)](http://f1000.com/work/citation?ids=686092&pre=&suf=&sa=0) |
| KDR | Vascular | [(*44*)](http://f1000.com/work/citation?ids=686092&pre=&suf=&sa=0) |
| PTPRB | Vascular | [(*45*)](http://f1000.com/work/citation?ids=7214781&pre=&suf=&sa=0) |
| PECAM1 | Vascular | [(*44*)](http://f1000.com/work/citation?ids=686092&pre=&suf=&sa=0) |
| HBG1 | Erythroblast | [(*46*)](http://f1000.com/work/citation?ids=926225&pre=&suf=&sa=0) |
| HBG2 | Erythroblast | [(*46*)](http://f1000.com/work/citation?ids=926225&pre=&suf=&sa=0) |
| HBB | Erythroblast | [(*46*)](http://f1000.com/work/citation?ids=926225&pre=&suf=&sa=0) |
| PTPRC | Leukocyte | [(*47*)](http://f1000.com/work/citation?ids=1117813&pre=&suf=&sa=0) |
| STAR | Adrenal cortex | [(*48*)](http://f1000.com/work/citation?ids=1546306&pre=&suf=&sa=0) |
| MC2R | Adrenal cortex | [(*48*)](http://f1000.com/work/citation?ids=1546306&pre=&suf=&sa=0) |
| TCF21 | Mesenchymal | [(*49*)](http://f1000.com/work/citation?ids=8700387&pre=&suf=&sa=0) |
| PDGFRB | Mesenchymal | [(*50*)](http://f1000.com/work/citation?ids=5370164&pre=&suf=&sa=0) |
| SOX10 | Schwann cell precursor | [(*9*, *51*)](http://f1000.com/work/citation?ids=3914331,5336106&pre=&pre=&suf=&suf=&sa=0,0) |
| MPZ | Schwann cell precursor | [(*51*)](http://f1000.com/work/citation?ids=5336106&pre=&suf=&sa=0) |
| PLP1 | Schwann cell precursor | [(*9*, *52*)](http://f1000.com/work/citation?ids=8243074,3914331&pre=&pre=&suf=&suf=&sa=0,0) |
| ERBB3 | Schwann cell precursor | [(*9*, *51*)](http://f1000.com/work/citation?ids=3914331,5336106&pre=&pre=&suf=&suf=&sa=0,0) |
| DLL3 | Bridge | [(*9*)](http://f1000.com/work/citation?ids=3914331&pre=&suf=&sa=0) |
| TLX2 | Bridge | [(*9*)](http://f1000.com/work/citation?ids=3914331&pre=&suf=&sa=0) |
| TH | Sympathoblast/chromaffin | [(*3*, *9*)](http://f1000.com/work/citation?ids=3914331,1219493&pre=&pre=&suf=&suf=&sa=0,0) |
| DBH | Sympathoblast/chromaffin | [(*3*, *9*)](http://f1000.com/work/citation?ids=3914331,1219493&pre=&pre=&suf=&suf=&sa=0,0) |
| PHOX2B | Sympathoblast/chromaffin | [(*9*)](http://f1000.com/work/citation?ids=3914331&pre=&suf=&sa=0) |
| CHGB | Sympathoblast/chromaffin | [(*3*, *9*)](http://f1000.com/work/citation?ids=3914331,1219493&pre=&pre=&suf=&suf=&sa=0,0) |
| GAP43 | Sympathoblast | [(*3*)](http://f1000.com/work/citation?ids=1219493&pre=&suf=&sa=0) |
| BCL2 | Sympathoblast | [(*3*)](http://f1000.com/work/citation?ids=1219493&pre=&suf=&sa=0) |
| NPY | Sympathoblast | [(*3*)](http://f1000.com/work/citation?ids=1219493&pre=&suf=&sa=0) |
| PNMT | Chromafin | [(*3*)](http://f1000.com/work/citation?ids=1219493&pre=&suf=&sa=0) |
| PHOX2B | Neuroblastoma | [(*53*)](http://f1000.com/work/citation?ids=8703026&pre=&suf=&sa=0), [(*54*)](http://f1000.com/work/citation?ids=1219227&pre=&suf=&sa=0) |
| PHOX2A | Neuroblastoma | [(*54*)](http://f1000.com/work/citation?ids=1219227&pre=&suf=&sa=0) |
| MYCN | Neuroblastoma | [(*55*, *56*)](http://f1000.com/work/citation?ids=8703042,4547231&pre=&pre=&suf=&suf=&sa=0,0) |

**Table S2. Markers curated from literature**

Genes used as markers of different cell types and references that justify their use.

| **Sample** | **Time point in treatment** | **Risk** | **INSS stage** | **MYCN amplification status at diagnosis** |
| --- | --- | --- | --- | --- |
| NB125 | Pre-treatment | High risk | III | Yes |
| GOSH021 | Pre-treatment | High risk | IV | No |
| NB060 | Pretreated (resection primary) | High risk | IV | Yes |
| NB098 | Pretreated (resection primary) | High risk | IV | No |
| NB107 | Pretreated (resection primary) | High risk | IV | Yes |
| NB123 | Pretreated (resection primary) | High risk | IV | No |
| NB124 | Pretreated (resection primary) | High risk | IV | Yes |
| NB130 | Pretreated (resection primary) | High risk | IV | Yes |
| NB132 | Pretreated (resection primary) | High risk | IV | Yes |
| NB086 | Pretreated (biopsy metastasis) | High risk | IV | No |
| NB106 | Pretreated (resection primary) | Intermediate risk | III | Yes |
| NB151 | Pretreated (resection primary) | Intermediate risk | IV | No |
| NB152 | Pretreated (resection primary) | Intermediate risk | IV | No |
| GOSH014 | Pretreated (resection primary) | Intermediate risk | IV | No |
| GOSH023 | Pretreated (resection primary) | Intermediate risk | IIA | No |
| GOSH019 | Pre-treatment | Low risk | IIA | No |
| GOSH025 | Pre-treatment | Low risk | III | No |
| NB138 | Pretreated (resection primary) | Low risk | IV | No |

**Table S3. Tumour sample manifest**

Clinical features of all single cell tumour samples.

| **Sample** | **Tumour** | **Mesenchyme** | **Leukocyte** | **Endothelium** | **Schwannian stroma** |
| --- | --- | --- | --- | --- | --- |
| NB125 | 425 | 68 | 694 | 17 | 14 |
| GOSH021 | 425 | 0 | 373 | 0 | 0 |
| NB060 | 67 | 97 | 209 | 16 | 3 |
| NB098 | 0 | 165 | 12 | 1 | 1 |
| NB107 | 0 | 155 | 46 | 44 | 17 |
| NB123 | 0 | 0 | 18 | 0 | 0 |
| NB124 | 8 | 436 | 1433 | 130 | 3 |
| NB130 | 11 | 16 | 634 | 9 | 2 |
| NB132 | 3 | 981 | 262 | 25 | 4 |
| NB086 | 5 | 102 | 57 | 5 | 2 |
| NB106 | 0 | 171 | 533 | 51 | 30 |
| NB151 | 0 | 443 | 508 | 48 | 149 |
| NB152 | 11 | 164 | 294 | 63 | 22 |
| GOSH014 | 1 | 125 | 3481 | 114 | 0 |
| GOSH023 | 21 | 0 | 84 | 35 | 0 |
| GOSH019 | 24 | 1 | 44 | 4 | 0 |
| GOSH025 | 1295 | 43 | 350 | 22 | 0 |
| NB138 | 0 | 1106 | 309 | 46 | 4 |
| Total | 2296 | 4073 | 9341 | 630 | 251 |

**Table S4. Tumour sample cell type contribution**

Number of cells of each cell type from each tumour sample.

**Table too large to embed, see supplementary_tables.xlsx**

**Table S5. Genes differentially expressed between cells with and without CN changes**

Comparison of cells from samples with and without copy number loss on chromosomes 1 and 11. For each gene, the log fold change (logFC) between genes with and without loss of 1 and 11 are given. logCPM gives the average counts per million, F gives the F-statistic, PValue the raw p-value, FDR the multiple hypothesis corrected false discovery rate. Finally, the genomic coordinates of each gene is given by its chromosome and transcription start site (TSS).

**Table too large to embed, see supplementary_tables.xlsx**

**Table S6. Algorithmically defined marker genes in fetal adrenal gland**

Genes specific to different clusters of cells and the evidence for their specificity in fetal adrenal gland. Genes listed here have a FDR (qval column) <0.01 from a hypergeometric test (see Methods). Within each cluster, genes are sorted by their tf-idf value (see Methods). The cluster ID is given in the “cluster” column. Other columns give the frequency with which a gene occurs within the target cluster, outside the cluster, in the second highest cluster, and globally.
**Table too large to embed, see supplementary_tables.xlsx**

**Table S7. Genes differentially expressed between fetal medulla and tumour cells**

Genes specific to tumour cells relative to fetal adrenal gland in both 10X and CEL-seq2 tumour datasets. For each dataset, genes are filtered so that they have FDR < 0.01 and less than 20% expression in the second highest cluster. These two lists are then combined (with NAs introduced when a gene is absent from one list) and genes with mean tf-idf > 0.85 are retained (see Methods). Those columns that end with .PMC (.GOSH) are derived from the comparison between the CEL-seq2 (10X) tumour data and the fetal adrenal data.

**Table too large to embed, see supplementary_tables.xlsx**

**Table S8. Algorithmically defined marker genes in 10x tumour data**

Genes specific to different clusters of cells and the evidence for their specificity in 10x tumour data. Genes listed here have a FDR (qval column) <0.01 from a hypergeometric test (see Methods). Within each cluster, genes are sorted by their tf-idf value (see Methods). The cluster ID is given in the “cluster” column. Other columns give the frequency with which a gene occurs within the target cluster, outside the cluster, in the second highest cluster, and globally.

**Table too large to embed, see supplementary_tables.xlsx**

**Table S9. Algorithmically defined marker genes in CEL-seq2 tumour data**

Genes specific to different clusters of cells and the evidence for their specificity in CEL-seq2 tumour data. Genes listed here have a FDR (qval column) <0.01 from a hypergeometric test (see Methods). Within each cluster, genes are sorted by their tf-idf value (see Methods). The cluster ID is given in the “cluster” column. Other columns give the frequency with which a gene occurs within the target cluster, outside the cluster, in the second highest cluster, and globally.

**Table too large to embed, see supplementary_tables.xlsx**

**Table S10. Transcription factors important to medulla development**

The transcription factors that change significantly along branches in the fetal adrenal medulla. Genes listed here have FDR < 0.001 from Moran’s I test (see Methods).

**Table too large to embed, see supplementary_tables.xlsx**

**Table S11. Copy number changes tumour samples from 10x data**

Table of copy number changes identified in bulk DNA for each of the tumor samples in the 10x dataset. Copy number segments are defined by their chromosome start and end, and the total and minimum (i.e. the number of copies of the minor allele) copy number are given for tumour and normal.

**Table too large to embed, see supplementary_tables.xlsx**

**Table S12. Copy number changes in DNA of tumours in CEL-seq2 data**

Table of copy number changes identified in bulk DNA for each of the tumor samples in the CEL-seq2 dataset. Copy number segments are defined by their chromosome start and end, and the total and minimum (i.e. the number of copies of the minor allele) copy number are given for tumour and normal.

**Table too large to embed, see supplementary_tables.xlsx**

**Table S13. Stringent algorithmically defined marker genes in fetal adrenal gland with podocytes included**

Genes specific to different clusters of cells and the evidence for their specificity in fetal adrenal gland, with fetal podocytes . Genes listed here have a FDR <0.01 from a hypergeometric test, tf-idf >1 and geneFrequencySecondBest <0.2(see Methods). Within each cluster, genes are sorted by their tf-idf value (see Methods). The cluster ID is given in the “cluster” column.

**Table too large to embed, see supplementary_tables.xlsx**

**Table S14. Differentially expressed genes between low risk and high risk neuroblastomas in TARGET**

Comparison of the risk group dependence between risk groups, controlling for age and MYCN status (see Methods) for genes identified as specific to tumour cells compared to the fetal medulla (**table S7**).

**see supplementary_scripts.gz**

**Data S1. - Scripts used to perform analysis and generate figures and tables**

Tar archive containing various scripts used to perform the analyses described in this manuscript and generate the results presented. This code is not presented as software intended for reuse but as extended documentation and precise description of the methodology used.
