## supplementary scripts for "Tumor to normal single cell mRNA comparisons reveal a pan-neuroblastoma cancer cell": CNsegExpression.R

```
setwd("~/trueHome/Projects/SCP")
#Want to look at how the genes that are expressed at different parts of the developmental trajectories are distributed along the genome and how this correlates with regions with recurrent CN change in Neurblastoma

###########
# Library #
###########
library(Seurat)
library(GenomicRanges)
library(SoupX)
#These should not be loaded, but are needed
required = c('rtracklayer')
if(!all(required %in% rownames(installed.packages())))
  stop("Required libraries not found.")
import('Code/plotParts.R',as='pl')


##############
# Parameters #
##############
srcGTF = 'Data/gtf10X_GRCh38_120.gtf'
srcSrat = 'Results/preProcess/fAdrenal/processedSeurat.RDS'
outDir = 'Results/'
binSize=1e6
cellsPerClust=100
collapseByCluster=TRUE


#############
# Functions #
#############
#' Nicked from GOseq
reversemapping=function(map){
	tmp=unlist(map,use.names=FALSE)
	names(tmp)=rep(names(map),times=as.numeric(summary(map)[,1]))
	return(split(names(tmp),as.vector(tmp)))
}

##################
# Standard plots #
##################
#Define annotation
clMap =  c('13'='Left',
           '25'='SCP',
           '24'='Bridge',
           '19'='Right',
           '20'='Right',
           '11'='Mes',
           '18'='Mes',
           '23'='Mes',
           '0'='Cortex',
           '1'='Cortex',
           '2'='Cortex',
           '4'='RBC',
           '7'='RBC')
cMap = list(Medulla=c(25,24,13,19,20),
            Cortex=c(5,22,28,2,1,0,10,8,6,9,21),
            Mesenchyme=c(11,18,23),
            Immune=c(16,26,29,31),
            RBC=c(7,4,12,30,27),
            Endo=c(3,15,14,17))
#Get markers of different parts
srat = readRDS(srcSrat)
marks = srat@meta.data$seurat_clusters
marks = ifelse(marks %in% names(clMap),clMap[marks],marks)
marks = quickMarkers(srat@assays$RNA@counts,marks,N=Inf)
marks = marks[marks$tfidf>0.8,]
chrLens = rtracklayer::SeqinfoForUCSCGenome('hg38')
chrLens = setNames(seqlengths(chrLens),gsub('chr','',seqnames(chrLens)))
ann = rtracklayer::import(srcGTF)
gns = ann[ann$type=='gene']
gns$TSS = ifelse(strand(gns)=='+',start(gns),end(gns))
#Reverse it
cMapRev = unlist(reversemapping(cMap))
#Get the binarised table of counts
cnts = srat@assays$RNA@counts
#cnts@x = rep(1,length(cnts@x))
#Make dense and transpose
cnts = as.matrix(t(cnts))
#Subset to matchable genes (it's basically all of them)
cnts = cnts[,colnames(cnts) %in% gns$gene_name]
#Order by chromosome, then position
m= match(colnames(cnts),gns$gene_name)
cnts = cnts[,order(as.character(seqnames(gns[m])),pmin(start(gns[m]),end(gns[m])))]
###########################
# Get recurrent CN changes
#Define genomic bins
gBins = lapply(1:22,function(e) data.frame(chr=e,binNum=seq(0,floor(chrLens[match(e,names(chrLens))]/binSize))))
gBins = do.call(rbind,gBins)
gBins = GRanges(gBins$chr,IRanges(gBins$binNum*binSize+1,(gBins$binNum+1)*binSize),nom=paste0(gBins$chr,'___',gBins$binNum))
#Get recurrent CN segments
cn = read.table('Data/cnSegsNB.txt',sep='\t',header=TRUE)
cn = read.table('Data/cnSegsNBlifted.txt',sep='\t',header=TRUE)
cn$chromosome = gsub('chr','',cn$chromosome)
cnSegs = GRanges(cn$chromosome,IRanges(cn$min,cn$max),cnType=cn$annotation)
#Get the pileup of gains and losses
gains = cn[cn$annotation=='gain',]
gains = GRanges(gains$chromosome,IRanges(gains$min,gains$max))
gains = setNames(countOverlaps(gBins,gains),gBins$nom)
losses = cn[cn$annotation=='loss',]
losses = GRanges(losses$chromosome,IRanges(losses$min,losses$max))
losses = setNames(countOverlaps(gBins,losses),gBins$nom)
#Get density of breakpoitns for gains and losses
bpGains = cn[cn$annotation=='gain',]
bpGains = GRanges(rep(bpGains$chromosome,2),IRanges(c(bpGains$min,bpGains$max),width=1))
#Drop ones near start and end of chromosomes
bpGains = bpGains[start(bpGains)>5e5]
bpGains = bpGains[which(start(bpGains)<chrLens[as.character(seqnames(bpGains))]-5e5)]
#Finally, get counts in bins
bpGains = setNames(countOverlaps(gBins,bpGains),gBins$nom)
#Repeat for losses
bpLosses = cn[cn$annotation=='loss',]
bpLosses = GRanges(rep(bpLosses$chromosome,2),IRanges(c(bpLosses$min,bpLosses$max),width=1))
bpLosses = bpLosses[start(bpLosses)>5e5]
bpLosses = bpLosses[which(start(bpLosses)<chrLens[as.character(seqnames(bpLosses))]-5e5)]
bpLosses = setNames(countOverlaps(gBins,bpLosses),gBins$nom)
####################################################
# This is too big to work, start bashing away at it
#We're only interested in the changes within Medulla, focus on that.
pCnts = cnts[which(clMap[as.character(srat@meta.data[rownames(cnts),'seurat_clusters'])] %in% c('RBC','Left','Right')),]# %in% c(25,24,13,19,20,11,18,23,0,1,2,4,7),]
#Group things into clusters
clusts = as.character(srat@meta.data[rownames(pCnts),'seurat_clusters'])
clusts = clMap[clusts]
if(collapseByCluster){
  #Collapse by cluster
  s = split(rownames(pCnts),clusts)
  pCnts = do.call(rbind,lapply(s,function(e) colSums(pCnts[e,,drop=FALSE])))
  #Give it a name that corresponds to one cell for consistency
  rownames(pCnts) = sapply(s,head,n=1)[rownames(pCnts)]
}else{
  #Randomly select X of each celltype
  w = unlist(lapply(split(seq_along(clusts),clusts),function(e) if(length(e)<cellsPerClust){e}else{sample(e,size=cellsPerClust)}))
  pCnts = pCnts[w,]
}
#Normalise to sum to 1
pCnts = pCnts/rowSums(pCnts)
#Forget binning, get the weighted density for each gene
tgts = marks[marks$cluster %in% c('Left','Right','RBC'),]
nTgts = min(table(tgts$cluster))
tgts = lapply(unique(tgts$cluster),function(e) {tmp = tgts[tgts$cluster==e,]
              tmp = tmp[order(-tmp$tfidf),]
              return(tmp$gene[seq(nTgts)])})
tgts = unique(unlist(tgts))
tgts = colnames(pCnts)
dds = list()
#Get the groupings of cells
m= match(colnames(pCnts),gns$gene_name)
for(chr in as.character(1:22)){
  message(sprintf("Averaging expression for chromosome %s",chr))
  dds[[chr]]=list()
  for(nom in rownames(pCnts)){
    w = which(as.character(seqnames(gns[m]))==chr & colnames(pCnts) %in% tgts)
    if(length(w)==0)
      next
    weights = pCnts[nom,w]
    if(sum(weights)==0)
      next
    #weights = weights/sum(weights)
    dds[[chr]][[nom]] = suppressWarnings(
                                         density(gns$TSS[m[w]],
                                                 weights=weights,
                                                 bw=2e5,
                                                 from=1,
                                                 to=chrLens[chr],
                                                 n=2048)
                                         )
  }
}
##################################
# Chromosome by chromosome plots #
##################################
doPlot = function(colScale=FALSE,topVal=200,tgtChr=1:22,diffOnly=FALSE,trans = I,expQuant=0.7,nCol=4,nRow=2){
  #Get range for colour scale
  colMax = lapply(dds,function(e) lapply(e,function(ee) max(ee$y)))
  colMax = quantile(unlist(colMax),expQuant)
  colMax = as.numeric(sprintf('%.01g',colMax))
  #Create layout
  plotBlock = seq(ifelse(diffOnly,3,6))
  heightBlock = c(rep(2,ifelse(diffOnly,0,3)),2,0.2,1)
  blockLen=length(plotBlock)
  layoutMat = matrix(NA,nrow=blockLen*nRow,ncol=nCol)
  offset=0
  for(j in seq(nRow)){
    for(i in seq(nCol)){
      layoutMat[(blockLen*(j-1)+1):(blockLen*j),i] = plotBlock+offset
      offset=offset+blockLen
    }
  }
  heights = rep(heightBlock,nRow)
  layout(layoutMat,heights=heights,widths=c(1.5,rep(1,nCol-1)))
  plotNum=0
  for(chr in as.character(tgtChr)){
    print(chr)
    #Decide if this is the leftmost block of plots
    plotLabs = plotNum%%nCol ==0
    if(plotLabs){
      leftMar = 7
    }else{
      leftMar = 0
    }
    mat = do.call(rbind,lapply(dds[[chr]],function(e) e$y))
    rowSplit = clMap[as.character(srat@meta.data[rownames(mat),'seurat_clusters'])]
    rowSplit = factor(rowSplit,levels=c('Left','Right','RBC'))
    #Get locations to sample at
    sampLoc = dds[[chr]][[1]]$x
    sampLoc = GRanges(chr,IRanges(sampLoc,width=1))
    #Alternative visualisation
    x=start(sampLoc)
    par(mar=c(0,leftMar,0,0)+0.1)
    ccs = list(RBC=pl$colAlpha('#E96245',0.5),
               Left = pl$colAlpha('#26828E',0.5),
               Right = pl$colAlpha('#35B779',0.5)
               )
    #yCon = mat[rowSplit=='RBC',]
    ##################
    # The raw tracks #
    ##################
    if(!diffOnly){
      for(i in seq(3)){
        if(i==1){
          tgt='RBC'
        }else if(i==2){ 
          tgt='Right'
        }else{
          tgt='Left'
        }
        lab =c(Left='Chromaffin',
               Right='Sympathoblasts',
               RBC='Erythroblasts')[tgt]
        y=trans(mat[rowSplit==tgt,])
        cc = ccs[[tgt]]
        #ccCon = ccs[['RBC']]
        #Plot control first
        plot(x,y,
             type='l',
             xaxt='n',
             yaxt='n',
             frame.plot=FALSE,
             ylim=c(0,trans(colMax)),
             xlim=c(1,chrLens[chr]),
             xlab='',
             ylab='',
             col=cc)
        if(plotLabs){
          tickLoc = axTicks(side=2)
          axis(side=2,
               las=1,
               line=1,
               at = tickLoc,
               labels=c(0,rep(NA,max(0,length(tickLoc)-2)),sprintf('%0.1g',trans(colMax)))
               )
          title(ylab=lab,line=5)
        }
        polygon(c(1,x,max(x),1),c(0,y,0,0),col=cc,border=NA)
        #Now the actual thing
        #polygon(c(x,1),c(yCon,0),col=ccCon,border=pl$colAlpha('black',1.0))
        if(i==1)
          text(chrLens[chr]/2,trans(colMax),paste0('chr',chr),xpd=NA)
      }
    }
    #############
    # Diff track
    par(mar=c(-0.1,leftMar,0,0)+0.1)
    y=mat[rowSplit=='Right',]-mat[rowSplit=='Left',]
    y = sign(y) * trans(abs(y))
    plot(0,
         type='n',
         xaxt='n',
         yaxt='n',
         frame.plot=FALSE,
         ylim=c(-trans(colMax),trans(colMax)),
         xlim=c(1,chrLens[chr]),
         xlab='',
         ylab=''
         )
    if(plotLabs){
      axis(side=2,
           las=1,
           line=1,
           at=c(-trans(colMax),0,trans(colMax)),
           labels=sprintf('%0.1g',c(-trans(colMax),0,trans(colMax)))
           )
      #title(ylab='Average expression of \nSympathoblasts - Chromaffin',line=5)
      title(ylab='Diff',line=5)
    }
    #The diff track
    polygon(c(1,x,max(x),1),c(0,ifelse(y>0,y,0),0,0),col=ccs[['Right']],border=NA)
    polygon(c(1,x,max(x),1),c(0,ifelse(y<0,y,0),0,0),col=ccs[['Left']],border=NA)
    lines(x,y,lwd=0.01)
    ###############
    # The ideogram
    par(mar=c(-.1,leftMar,-.1,0)+0.1)
    cyto = karyoploteR:::data.cache$cytobands$hg38
    cyto = cyto[as.character(seqnames(cyto)) == paste0('chr',chr)]
    plot(0,
         xlim=c(1,chrLens[chr]),
         ylim=c(0,1),
         type='n',
         xaxt='n',
         yaxt='n',
         xlab='',
         ylab='',
         frame.plot=FALSE
         )
    #Make thing, nicked from the internals
    color.table <- karyoploteR::getCytobandColors(color.table = NULL, color.schema = 'circos')
    border = ifelse("border" %in% names(color.table), color.table["border"], "black")
    ybottom = 0
    ytop = 1
    xleft = start(cyto)
    xright = end(cyto)
    col = color.table[as.character(cyto$gieStain)]
    rect(xleft = xleft, xright = xright, ybottom = ybottom, ytop = ytop, col = col, border = border)
    #Add cytoband labels. Only plotted if they fit.  Ripped from library again
    labs = cyto$name
    bands.chr = as.character(seqnames(cyto))
    xmids = (xleft + (xright - xleft)/2)
    label.width = graphics::strwidth(labs, cex = 0.5)
    do.fit = label.width < (xright - xleft)
    if (FALSE && any(do.fit)) {
      graphics::text(x = xmids[do.fit], y = 0.5, labels = labs[do.fit], cex = 0.5,xpd=NA)
    }
    ###########################
    # Finally add the CN track
    par(mar=c(0.1,leftMar,-.1,0)+0.1)
    loss = countOverlaps(sampLoc,cnSegs[cnSegs$cnType=='loss'])/556*100
    gain = countOverlaps(sampLoc,cnSegs[cnSegs$cnType=='gain'])/556*100
    topVal=100
    plot(0,
         xlim=c(1,chrLens[chr]),
         ylim=c(-topVal,topVal),
         type='n',
         xaxt='n',
         yaxt='n',
         ylab='',
         xlab='',
         frame.plot=FALSE
         )
    if(plotLabs){
      axis(side=2,
           las=1,
           line=1,
           at = c(-topVal,0,topVal),
           labels=c(topVal,0,topVal),
           )
      #title(ylab='% Samples\nwith CNA',line=5)
      title(ylab='% CNA',line=5)
    }
    polygon(c(1,x,max(x),1),c(0,-loss,0,0),col=pl$colAlpha('darkblue',0.9),border=NA)
    polygon(c(1,x,max(x),1),c(0,gain,0,0),col=pl$colAlpha('darkred',0.9),border=NA)
    ####Use this to make annotation track
    ###botAnno = HeatmapAnnotation(gains = countOverlaps(sampLoc,cnSegs[cnSegs$cnType=='gain']),
    ###                            losses = countOverlaps(sampLoc,cnSegs[cnSegs$cnType=='loss']),
    ###                            col = list(gains = circlize::colorRamp2(c(0,topVal),c('White','Red')),
    ###                                       losses = circlize::colorRamp2(c(0,topVal),c('White','Blue'))),
    ###                            which='column'
    ###                            )
    ###if(colScale)
    ###  mat = scale(mat)
    ####Set column names selectively to fake ticks
    ###ticks=start(sampLoc)/1e6
    ###ticks = round(ticks)
    ###fin=max(ticks)
    ###ticks = unlist(lapply(split(ticks,cut(seq_along(ticks),10)),function(e) rep(min(e),length(e))))
    ###ticks[length(ticks)]=fin
    ###ticks = ifelse(duplicated(ticks),'',as.character(ticks))
    ###colnames(mat) = ticks
    ####Define colour scheme
    ###col = suppressWarnings(brewer.pal(100,'PuRd'))
    ###col = circlize::colorRamp2(seq(0,colMax,length.out=length(col)),col)
    ###hm = Heatmap(mat,
    ###             col=col,
    ###             name='Smoothed Expression',
    ###             column_title=paste0('chr',chr),
    ###             row_split = rowSplit,
    ###             bottom_annotation = botAnno,
    ###             show_column_names=TRUE,
    ###             cluster_columns = FALSE,
    ###             cluster_row_slices = FALSE,
    ###             show_row_names=FALSE,
    ###             show_row_dend = FALSE)
    ###print(hm)
    plotNum=plotNum+1
  }
}
pdf('Results/cnRaw.pdf',width=11.7,height=8.3)
doPlot()
dev.off()
#pdf('Results/cnScaled.pdf',width=10,height=7)
#doPlot(TRUE)
#dev.off()
pdf('Results/chr1p.pdf',width=4,height=3)
doPlot(tgtChr='1',diffOnly=TRUE,expQuant=0.5)
dev.off()
pdf('Results/chr11q.pdf',width=4,height=3)
doPlot(tgtChr='11',diffOnly=TRUE,expQuant=0.5)
dev.off()


################################
# Statistical tests of regions #
################################
#First the loss on chr1
chr='1'
sampLoc = dds[[chr]][[1]]$x
sampLoc = GRanges(chr,IRanges(sampLoc,width=1))
x = countOverlaps(sampLoc,cnSegs[cnSegs$cnType=='loss'])
#200 is equavilent to 36%, total sample number is 556
testSegs = GRanges(chr,IRanges(min(start(sampLoc)[x>200]),max(start(sampLoc)[x>200])))
#And the boundary point around inflection on chr11
chr='11'
sampLoc = dds[[chr]][[1]]$x
sampLoc = GRanges(chr,IRanges(sampLoc,width=1))
x = countOverlaps(sampLoc,cnSegs[cnSegs$cnType=='loss'])
y = countOverlaps(sampLoc,cnSegs[cnSegs$cnType=='gain'])
tgt = min(start(sampLoc)[x>70])
tgtWid = 2e6
testSegs = c(testSegs,GRanges(chr,IRanges(tgt-tgtWid,tgt+tgtWid)))
#Make a plot to justify this choice
pdf('Results/boundaryDef.pdf')
plot(start(sampLoc),x,
     type='l',
     frame.plot=FALSE,
     main='chr11',
     xlab='Position',
     ylab='Reccurance',
     col='blue'
)
lines(start(sampLoc),y,col='red')
abline(v=c(tgt-tgtWid,tgt+tgtWid),lty=2)
legend(x='topleft',
       legend=c('Gain','Loss'),
       col=c('red','blue'),
       lty=1,
       bty='n')
dev.off()
#Add other test segments to prevent them being used in random sampling
#testSegs = c(testSegs,
#             GRanges('1',IRanges(1,chrLens['1'])),
#             GRanges('2',IRanges(1,floor(chrLens['2']*0.8))),
#             GRanges('3',IRanges(1,floor(chrLens['3']*0.4))),
#             GRanges('6',IRanges(1,floor(chrLens['6']*0.5))),
#             GRanges('7',IRanges(1,floor(chrLens['7']*0.4))),
#             GRanges('11',IRanges(1,floor(chrLens['11']*1.0))),
#             GRanges('12',IRanges(1,floor(chrLens['12']*1.0))),
#             GRanges('17',IRanges(1,floor(chrLens['17']*1.0))),
#             GRanges('18',IRanges(1,floor(chrLens['18']*1.0))))
#Get the max per chromosome
maxCnt = sapply(as.character(1:22),function(chr){
                  sampLoc = dds[[chr]][[1]]$x
                  sampLoc = GRanges(chr,IRanges(sampLoc,width=1))
                  x = countOverlaps(sampLoc,cnSegs[cnSegs$cnType=='loss'])
                  y = countOverlaps(sampLoc,cnSegs[cnSegs$cnType=='gain'])
                  return(max(x,y))})
tgts = names(maxCnt[maxCnt>100])
testSegs = c(testSegs,GRanges(tgts,IRanges(1,chrLens[tgts])))
#Now get the difference in average expression between left and right in bins of this size throughout the genome.
mat = lapply(1:22,function(chr) {
               tmp = do.call(rbind,lapply(dds[[chr]],function(e) e$y))
               colnames(tmp) = dds[[chr]][[1]]$x
               tmp})
names(mat) = as.character(1:22)
calcSegDiff = function(mat,seg,tgtA='Right',tgtB='Left'){
  tmp = mat[[as.character(seqnames(seg))]]
  rowSplit = clMap[as.character(srat@meta.data[rownames(tmp),'seurat_clusters'])]
  x = as.numeric(colnames(tmp))
  w = which(x>=start(seg) & x<=end(seg))
  #The average expression in both, then take the difference
  mean(tmp[rowSplit==tgtA,w])-mean(tmp[rowSplit==tgtB,w])
}
set.seed(1137)
randScores=list()
nRand=1000
for(i in seq_along(testSegs[1:2])){
  #Calculate a comparative difference thing across random genomic bins of same size
  segLen = width(testSegs[i])
  tt=NULL
  while(length(tt)<nRand){
    #Get a random chromosem
    cc = sample(names(mat),1)
    #Get a random start point in the valid range
    ss = sample(chrLens[cc]-segLen-1,1)
    tgt = GRanges(cc,IRanges(ss,width=segLen))
    if(length(suppressWarnings(findOverlaps(tgt,testSegs)))>0)
      next
    tgt$score =calcSegDiff(mat,tgt) 
    if(is.null(tt)){
      tt=tgt
    }else{
      tt = suppressWarnings(c(tt,tgt))
    }
    print(length(tt))
  }
  randScores[[i]]=tt
}
#Chr1 
pdf('Results/expAssocWithCN_chr1.pdf')
i=1
obs = calcSegDiff(mat,testSegs[i])
plot(density(randScores[[i]]$score),
     frame.plot=FALSE,
     main='chr1p Loss',
     xlab='Average expression difference\nRight - Left',
     ylab='Density'
)
pVal = sum(randScores[[i]]$score>=obs)/nRand
pVal2 = sum(abs(randScores[[i]]$score)>=obs)/nRand
legend(x='topleft',
       legend=c('Random CN Neutral Segments',sprintf('chr1p (p=%g)',pVal)),
       col=c('black','red'),
       lty=1,
       bty='n')
abline(v=obs,col='red')
dev.off()
message(sprintf('pVal=%g (one-sided) %g (two-sided)',pVal,pVal2))
#Chr11
pdf('Results/expAssocWithCN_chr11.pdf')
i=2
obs = calcSegDiff(mat,testSegs[i])
plot(density(randScores[[i]]$score),
     frame.plot=FALSE,
     main='chr11 breakpoint',
     xlab='Average expression difference\nRight - Left',
     ylab='Density'
)
pVal = sum(randScores[[i]]$score>=obs)/nRand
pVal2 = sum(abs(randScores[[i]]$score)>=obs)/nRand
legend(x='topleft',
       legend=c('Random CN Neutral Segments',sprintf('chr11 (p=%g)',pVal)),
       col=c('black','red'),
       lty=1,
       bty='n')
abline(v=obs,col='red')
dev.off()
message(sprintf('pVal=%g (one-sided) %g (two-sided)',pVal,pVal2))
```
