## supplementary scripts for "Tumor to normal single cell mRNA comparisons reveal a pan-neuroblastoma cancer cell": combinedAdrenal.R

```
setwd('~/trueHome/Projects/SCP/')

#############
# Libraries #
#############
library(Seurat)
library(SoupX)
library(ggplot2)
library(Matrix)
source('https://raw.githubusercontent.com/constantAmateur/MCVR/master/code.R')
import('Code/scQC.R')
import('Code/geneSets.R')

##############
# Parameters #
##############
#Output directory
outDir = 'Results/preProcess/fAdrenal'
srcs = 'Data/scData/fAdrenal'
srcs = file.path(srcs,list.files(srcs))
noms = basename(srcs)
srcs = lapply(srcs,function(e) setNames(file.path(e,list.files(e)),list.files(e)))
names(srcs) = noms
#Final cleanup
#Drop the bad bilateral adrenal sample
srcs$bilateralAdrenals = grep('7457$',srcs$bilateralAdrenals,invert=TRUE,value=TRUE)
srcs$bilateralAdrenals = setNames(file.path(srcs$bilateralAdrenals,'outs'),names(srcs$bilateralAdrenals))
#Drop the non-folders
srcs$lateAdrenals = grep('\\.RDS$',srcs$lateAdrenals,invert=TRUE,value=TRUE)
#Create a list of cleaned paths
strained=list()
for(ex in names(srcs)){
  strained[[ex]] = setNames(file.path(outDir,paste0(names(srcs[[ex]]),'_strainedCounts')),names(srcs[[ex]]))
}
#MT cut-off
mtCut = 0.3
#nGene Cut
minNumGenes = 300
#nUMI cut
minNumUMI = 1000
#How many doublets in a cluster before you throw the entire cluster
maxScrubFrac = 0.5
#UMAP parameters to try in grid
minDists = c(0,.01,.1,.3,.5)
NNs = c(5,10,20,50,100)
#Final parameters
finalMinDist = 0.5
finalNN = 50
#Clustering resolution
clusterRes=1.0
#Populations of genes to exclude from the analysis
excludeGenes = unlist(as.list(geneSets)[c('mtGenes','hspGenes','riboGenes')],use.name=FALSE)
excludeGenes = sort(unique(excludeGenes))
#Number of PCs to use for quick analysis during QC
defNumPCs=75

#############
# Functions #
#############
#Copy/pasted from dropletutils.  A useful function, but the package has an absurd number of dependencies
write10xCounts = function (path, x, barcodes = colnames(x), gene.id = rownames(x),
    gene.symbol = gene.id, overwrite = FALSE)
{
    temp.path <- tempfile(tmpdir = dirname(path))
    dir.create(temp.path, showWarnings = FALSE)
    on.exit({
        if (file.exists(temp.path)) {
            unlink(temp.path, recursive = TRUE)
        }
    })
    writeMM(x, file = file.path(temp.path, "matrix.mtx"))
    if (ncol(x) != length(barcodes)) {
        stop("'barcodes' must of of the same length as 'ncol(x)'")
    }
    write(barcodes, file = file.path(temp.path, "barcodes.tsv"))
    if (length(gene.id) != length(gene.symbol) || length(gene.id) !=
        nrow(x)) {
        stop("lengths of 'gene.id' and 'gene.symbol' must be equal to 'nrow(x)'")
    }
    write.table(data.frame(gene.id, gene.symbol), file = file.path(temp.path,
        "genes.tsv"), row.names = FALSE, col.names = FALSE, quote = FALSE,
        sep = "\t")
    if (overwrite) {
        unlink(path, recursive = TRUE)
    }
    else if (file.exists(path)) {
        stop("specified 'path' already exists")
    }
    file.rename(temp.path, path)
    return(invisible(TRUE))
}
#' Fine clustering for SoupX and other things
#'
#' @param dat Raw filtered count matrix.
#' @param numPCs How many PCs to use.
#' @param clusteringRes Clustering parameter to use.
#' @return Seurat object.
quickCluster = function(dat,numPCs=75,clusteringRes=10.0){
  ss = CreateSeuratObject(dat)
  ss = NormalizeData(ss,verbose=FALSE)
  ss = FindVariableFeatures(ss,verbose=FALSE)
  ss = ScaleData(ss,verbose=FALSE)
  ss = RunPCA(ss,npcs=numPCs,approx=FALSE,verbose=FALSE)
  ss = FindNeighbors(ss,dims=seq(numPCs),verbose=FALSE)
  ss = FindClusters(ss,res=clusteringRes,verbose=FALSE)
  ss = RunUMAP(ss,dims=seq(numPCs),verbose=FALSE)
  return(ss)
}
#' Standard processing
stdProcess = function(dat,...){
  ss = CreateSeuratObject(dat)
  ss = NormalizeData(ss,verbose=FALSE)
  #ss = CellCycleScoring(ss, s.features=geneSets[['S']][geneSets[['S']] %in% rownames(dat)],g2m.features=geneSets[['G2M']][geneSets[['G2M']] %in% rownames(dat)],seed=sample(1e9,1))
  #ss = ScaleData(ss,vars.to.regress=c('S.Score','G2M.Score'),verbose=FALSE,...)
  ss = ScaleData(ss,verbose=FALSE,...)
  return(ss@assays$RNA@scale.data)
}


#########
# SoupX #
#########
nonExpressedList = list(HG=c('HBB','HBG1','HBG2','HBA1','HBA2','HBM','HBD','HBE1','HBZ','HBQ1'))
#check if it's already been done
if(!all(file.exists(unlist(strained)))){
  for(ex in names(srcs)){
    pdf(file.path(outDir,sprintf('%s_SoupX_processing.pdf',ex)))
    for(i in seq_along(srcs[[ex]])){
      message(sprintf("Running SoupX on channel %d of %d in experiment %s",i,length(srcs[[ex]]),ex))
      sc = load10X(srcs[[ex]][i])
      #Do fine grained clustering
      cc = quickCluster(sc$toc,clusteringRes=1.0)
      sc = setClusters(sc,cc@active.ident)
      #Calculate which ones to exclude
      useToEst = estimateNonExpressingCells(sc,nonExpressedGeneList=nonExpressedList)
      #If this doesn't work, just use individual cells
      if(sum(useToEst)==0){
        #stop('Oh no!')
        useToEst = estimateNonExpressingCells(sc,nonExpressedGeneList=nonExpressedList,clusters=FALSE,maximumContamination=0.2,FDR=0.05)
      }
      gg = plotMarkerMap(sc,nonExpressedList[[1]],DR=cc@reductions$umap@cell.embeddings,useToEst=useToEst) + ggtitle(sprintf('HB expression and cell use in %s',names(srcs[[ex]][i])))
      plot(gg)
      sc = calculateContaminationFraction(sc,nonExpressedGeneList=nonExpressedList,useToEst=useToEst)
      out = adjustCounts(sc,verbose=2)
      #Save it
      write10xCounts(strained[[ex]][i],out,overwrite=TRUE)
    }
    dev.off()
  }
}

############
# Basic QC #
############
tgtFile = file.path(outDir,'baseSeurat.RDS')
if(file.exists(tgtFile)){
  srat = readRDS(tgtFile)
}else{
  #Create the basic Seurat object for each experiment
  message(sprintf("QC filters on everything at once"))
  pdf(file.path(outDir,sprintf('QC.pdf')))
    srat = scQC$basicQC(dataDirs=setNames(unlist(strained),unlist(lapply(strained,names))),
                        excludeGenes = excludeGenes,
                        geneSets = geneSets,
                        maxMT = mtCut,
                        minGenes = minNumGenes,
                        minUMIs = minNumUMI,
                        maxBadFrac = maxScrubFrac,
                        scrubScoreMax = 0.2,
                        numPCs = defNumPCs)
  dev.off()
  saveRDS(srat,tgtFile)
}

################
# Drop cycling #
################
tgtFile = file.path(outDir,'seuratG0.RDS')
if(file.exists(tgtFile)){
  srat = readRDS(tgtFile)
}else{
  #Aggressively filter out things
  w = which(srat@meta.data$S.Score<0 & srat@meta.data$G2M.Score<0)
  mDat = srat@meta.data[w,]
  srat = CreateSeuratObject(srat@assays$RNA@counts[,w])
  srat@meta.data = cbind(srat@meta.data,mDat[,!(colnames(mDat) %in% colnames(srat@meta.data)),drop=FALSE])
  srat = NormalizeData(srat,verbose=FALSE)
  srat = FindVariableFeatures(srat,verbose=FALSE)
  saveRDS(srat,tgtFile)
}


################
# Optimal nPCs #
################
#Work out what the best number of PCs is
tgtFile=file.path(outDir,'MCV.RDS')
if(file.exists(tgtFile)){
  mcv = readRDS(tgtFile)
}else{
  mcv = molecularCrossValidation(roundToInt(srat@assays$RNA@counts),VariableFeatures(srat),normalisation=stdProcess,features=VariableFeatures(srat),tFracs=c(1,0.9,0.8,0.5),nSplits=5,nCores=24)
  pdf(file.path(outDir,paste0(ex,'_mcv.pdf')))
    plotMCV(mcv)
  dev.off()
  saveRDS(mcv,tgtFile)
}
nPCs = max(mcv$mins1sd[[1]])
#nPCs = sapply(mcvList,function(e) max(e$mins1sd[[1]]))

####################
# Final processing #
####################
tgtFile = file.path(outDir,'processedSeurat.RDS')
pdf(file.path(outDir,paste0('standardPlots.pdf')),width=24,height=24)
  srat = ScaleData(srat)
  srat = RunPCA(srat,npcs=nPCs,approx=FALSE)
  #Run tSNE as well in case that turns out to be useful.
  #srat = RunTSNE(srat)
  #The old python based method seems to segfault every time...
  srat = FindNeighbors(srat,dims=seq(nPCs))
  srat = FindClusters(srat,res=clusterRes)
  #Make a range of UMAP plots
  umaps=list()
  for(minDist in minDists){
    for(NN in NNs){
      message(sprintf("Calculating UMAP for minDist=%g, NN=%d",minDist,NN))
      srat = RunUMAP(srat,dims=seq(nPCs[ex]),min.dist=minDist,n.neighbors=NN,verbose=FALSE)
      umaps[[paste0(minDist,'_',NN)]] = DimPlot(srat,label=TRUE) +
        guides(colour=FALSE) +
        ggtitle(sprintf("UMAP with minDist=%g, NN=%d",minDist,NN))
    }
  }
  plot(plot_grid(plotlist=umaps,
            nrow=length(minDists),
            ncol=length(NNs)))
  #Final UMAP
  srat = RunUMAP(srat,dims=seq(nPCs),min.dist=finalMinDist,n.neighbors=finalNN)
  #Clustering
  plot(DimPlot(srat,label=TRUE))
  #My ones
  plot(FeaturePlot(srat,c('PTPRC','PDGFRB','EPCAM','PECAM1','HBA1','HBB')))
  #Source
  plot(DimPlot(srat,group.by='orig.ident'))
  #Cell cycle
  plot(DimPlot(srat,group.by='Phase'))
  #QC things
  plot(FeaturePlot(srat,c('scrubScore','nFeature_RNA','nCount_RNA',gsub('Genes$','Frac',names(geneSets)))))
  #Visualise cell cycle
  #plot(FeaturePlot(srat,c('PCNA','MKI67','MCM4','CDKN1C')))
  #Genes specific to Steroidogenisis
  plot(FeaturePlot(srat,c('MC2R','CYP17A1','MRAP','STAR','CYP11B2','CYP11B1','INHA','GNRHR','HSD3B2')))
  #Medulla markers
  plot(FeaturePlot(srat,c('PHOX2A','ISL1','CHRNA3','SOX2','SOX10','ERBB3','DBH','CHGB','TH')))
  #Others
  plot(FeaturePlot(srat,c('NR5A1','NOV','HSD3B2','SULT2A1','FDX1')))
dev.off()
```
