## supplementary scripts for "Tumor to normal single cell mRNA comparisons reveal a pan-neuroblastoma cancer cell": Fig1_A-D_sub.R

```
library(Seurat)
library(monocle3)
library(ComplexHeatmap)
library(circlize)
library(RColorBrewer)
library(SoupX)
library(viridis)

#literature markers in adrenal gland
adr_all=readRDS("fAdrenal/processedSeurat.RDS")

FeaturePlot(adr_all, c("SOX10", "MPZ", "PLP1", "ERBB3", "DLL3", "TLX2", "TH", "DBH", "PHOX2B", 
  "CHGB","GAP43", "BCL2", "NPY", "PNMT", "PDGFRB", "TCF21", "PLVAP", "PECAM1", "KDR",
  "PTPRB","HBG1", "HBG2", "HBB","STAR", "MC2R", "PTPRC"))


#annotate louvain clusters
adr_all@meta.data$new_clust=as.character(adr_all@meta.data$seurat_clusters)
adr_all@meta.data$new_clust[which(adr_all@meta.data$new_clust%in%c("25"))]="SCPs"
adr_all@meta.data$new_clust[which(adr_all@meta.data$new_clust%in%c("13"))]="Chromaffin"
adr_all@meta.data$new_clust[which(adr_all@meta.data$new_clust%in%c("24"))]="Bridge"
adr_all@meta.data$new_clust[which(adr_all@meta.data$new_clust%in%c("19", "20"))]="Sympathoblastic"
adr_all@meta.data$new_clust[which(adr_all@meta.data$new_clust%in%c("3", "15", "14","17"))]="Endothelium"
adr_all@meta.data$new_clust[which(adr_all@meta.data$new_clust%in%c("23", "11", "18"))]="Mesenchyme"
adr_all@meta.data$new_clust[which(adr_all@meta.data$new_clust%in%c("22", "5", "28", "0", "1", "2", "10", "6", "8", "9","21"))]="Cortex"
adr_all@meta.data$new_clust[which(adr_all@meta.data$new_clust%in%c("16", "26"))]="Leukocytes"
adr_all@meta.data$new_clust[which(adr_all@meta.data$new_clust%in%c("30", "7", "4", "12"))]="Erythroblasts"
adr_all@meta.data$new_clust[which(adr_all@meta.data$new_clust%in%c("27", "31", "29"))]="Other"

adr_all@meta.data$new_clust=factor(adr_all@meta.data$new_clust, levels = c("SCPs","Bridge","Sympathoblastic",
                                                                           "Chromaffin","Endothelium",
                                                                           "Mesenchyme", "Cortex",
                                                                          "Leukocytes", "Erythroblasts", "Other"))
#make figure 1A
pdf("adrenal_gland.pdf", height = 3, width=3,useDingbats = F)
gg=DimPlot(adr_all, group.by = "new_clust", label=T,label.size = 3,cols=c(viridis(10)[c(1,3,5,7)],c("#CC6677", "#7E481C","#DDAA77","#DDCC77", "#E96245", "grey"))) +
  ggtitle("") + theme(text = element_text(size=12), axis.text.x = element_blank(),axis.text.y = element_blank(),
                      axis.ticks = element_blank(), legend.position = "none")+ylab("UMAP 2") +xlab("UMAP 1")
plot(gg)
dev.off()

#Cluster average expression of marker genes (fig S1)
#assign annotation as active.ident
adr_all_supp = adr_all
adr_names=names(adr_all_supp@active.ident)
adr_all_supp@active.ident=factor(adr_all_supp@meta.data$new_clust)
names(adr_all_supp@active.ident)=adr_names
adr_all_supp=ScaleData(adr_all_supp, features = rownames(adr_all_supp))
cluster.averages <- AverageExpression(adr_all_supp, return.seurat = T)
for_HM=cluster.averages@assays$RNA@scale.data[c("SOX10", "MPZ", "PLP1", "ERBB3", "DLL3", "TLX2", "TH", "DBH", "PHOX2B", 
                                                "CHGB","GAP43", "BCL2", "NPY", "PNMT", "PDGFRB", "TCF21", "PLVAP", "PECAM1", "KDR",
                                                "PTPRB","HBG1", "HBG2", "HBB","STAR", "MC2R", "PTPRC"),]
#literature marker cluster average heatmap
col_ord=c("SCPs","Bridge","Sympathoblastic","Chromaffin","Mesenchyme", "Endothelium",
          "Erythroblasts" ,"Cortex", "Leukocytes", "Other")
col_fun = colorRamp2(c(-3, 0, 3), c("blue", "white", "red"))

ht1=Heatmap(for_HM, cluster_rows = F, col=col_fun, column_order = col_ord, name = "Normalized expression", row_names_gp = gpar(fontsize = 8))

levels(adr_all_supp)=c("SCPs","Bridge","Sympathoblastic","Chromaffin","Mesenchyme", "Endothelium",
                       "Erythroblasts" ,"Cortex", "Leukocytes", "Other")
#top 5 markers of each cluster in adrenal gland
top_5_all_clust=quickMarkers(adr_all_supp@assays$RNA@counts, adr_all_supp@active.ident, N=5)
for_hm_inm=cluster.averages@assays$RNA@scale.data[as.character(top_5_all_clust$gene),]
#algorithmic marker heatmap
ht2=Heatmap(for_hm_inm, col=col_fun, cluster_rows=F, column_order = col_ord, name = "Normalized expression", row_names_gp = gpar(fontsize = 8))

ht_list = ht1 %v% ht2 

draw(ht_list)


#make figure 1B
adr_all_med=subset(adr_all, idents=c(25,24,13,19,20))
pdf("just_medulla.pdf", height = 3, width=3,useDingbats = F)
gg=DimPlot(adr_all_med, group.by = "new_clust", label=T,label.size = 3,cols=c(viridis(10)[c(1,3,5,7)])) +
  ggtitle("") + theme(text = element_text(size=12), axis.text.x = element_blank(),axis.text.y = element_blank(),
                      axis.ticks = element_blank(), legend.position = "none")+ylab("UMAP 2") +xlab("UMAP 1")
plot(gg)
dev.off()

#make figure 1C 
#get relevant data from Seurat medulla object
counts.data <-GetAssayData(object = adr_all_med, slot = "counts", assay="RNA")
pheno.data <- as.data.frame(adr_all_med@meta.data, row.names = rownames(adr_all_med@meta.data))
feature.data <- as.data.frame(adr_all_med@assays$RNA@counts@Dimnames[[1]], row.names = adr_all_med@assays$RNA@counts@Dimnames[[1]])

#create a monocle3 object
cds <- new_cell_data_set(counts.data, cell_metadata = pheno.data, gene_metadata = feature.data)
recreate.partition <- c(rep(1, length(cds@colData@rownames)))
names(recreate.partition) <- cds@colData@rownames
recreate.partition <- as.factor(recreate.partition)

cds@clusters@listData[["UMAP"]][["partitions"]] <- recreate.partition

list_cluster <- cds@colData$new_clust
names(list_cluster) <- adr_all_med@assays[["RNA"]]@data@Dimnames[[2]]


cds@clusters@listData[["UMAP"]][["clusters"]] <- as.factor(list_cluster)


cds@clusters@listData[["UMAP"]][["louvain_res"]] <- "NA"


# Assign UMAP coordinate

cds@reducedDims@listData[["UMAP"]] <-Embeddings(adr_all_med, reduction =  'umap')
cells_embeddings=Embeddings(adr_all_med, reduction =  'umap')
#learn principal graph
cds <- learn_graph(cds,use_partition = F)
#order cells with SCP node in shiny app
cds <- order_cells(cds)
#if shiny app doesn't work, use these cells
cds <- order_cells(cds, root_cells = c("babyAdrenal2_GTACTCCAGTCAAGGC","5388STDY7717454_CCAATCCAGTACACCT",
                                       "5388STDY7717456_AACACGTTCGTTACGA","5388STDY7717456_ATCTACTAGTAAGTAC",
                                       "5388STDY7717456_GACGTGCCATGCTGGC","5388STDY7717458_AAAGATGCATTACCTT",
                                       "5388STDY7717458_ACGCCGATCAGCACAT","5388STDY7717458_ACGGGCTTCGTCTGAA",
                                       "5388STDY7717458_CCTACCAAGGACAGAA","5388STDY7717458_TTCTCAAGTCTAGCGC",
                                       "5388STDY7717459_AACTCAGAGACCACGA" ,"5388STDY7717459_TGAAAGACAATGGATA",
                                       "cellranger302_count_32644_WSSS_F_Adr8710635_GRCh38-1_2_0_TTGGGCGAGCGTGAAC",
                                       "5698STDY7839907_GAATAAGCAGGAACGT","5698STDY7839907_TTCTCAATCGCTTGTC",
                                       "5698STDY7839909_AACCTGATCTGGCTGG","5698STDY7839909_ACACCAATCTCATAGG",
                                       "5698STDY7839909_ATACCTTCAGGCTATT","5698STDY7839909_CATAAGCTCCCTCATG",
                                       "5698STDY7839909_CTGCCATCAACTCGAT","5698STDY7839909_CTGGACGTCGTTCTGC",
                                       "5698STDY7839909_CTTACCGAGGTTCATC","5698STDY7839909_GAGGCCTAGACTCTAC",
                                       "5698STDY7839909_GTCTCACGTCACTGAT","5698STDY7839909_TAGTGCAAGAGAACCC",
                                       "5698STDY7839909_TGGATGTTCTTTCCAA" ,"5698STDY7839909_TTCCACGGTACGTAGG",
                                       "5698STDY7839917_AGTAGTCTCATTTGGG","5698STDY7839917_CAAGAAACACTTGGAT",
                                       "5698STDY7839917_CTCGTCATCGCAAGCC","5698STDY7839917_CTCTGGTCACTTAACG",
                                       "5698STDY7839917_CTTAGGAAGGTGTTAA" ,"5698STDY7839917_GGAATAAGTACAGTGG",
                                       "5698STDY7839917_TACAGTGCATACTCTT","5698STDY7839917_TTCCCAGAGCACCGCT"))
pseudo_srat = pseudotime(cds, reduction_method = 'UMAP')
adr_all_med@meta.data$pseudotime=pseudo_srat

pdf("all_medulla_pseudotime.pdf", height = 3, width=3,useDingbats = F)
gg=FeaturePlot(adr_all_med, features = "pseudotime", pt.size = 2) + scale_color_viridis(option="plasma", direction = 1) +
  ggtitle("") + theme(text = element_text(size=12), axis.text.x = element_blank(),axis.text.y = element_blank(),
                      axis.ticks = element_blank()) +ylab("UMAP 2") +xlab("UMAP 1")
plot(gg)
dev.off()

pdf("all_medulla_pseudotime_no_leg.pdf", height = 3, width=3,useDingbats = F)
gg=FeaturePlot(adr_all_med, features = "pseudotime", pt.size = 2) + scale_color_viridis(option="plasma", direction = 1) +
  ggtitle("") + theme(text = element_text(size=12), axis.text.x = element_blank(),axis.text.y = element_blank(),
                      axis.ticks = element_blank(), legend.position = "none") +ylab("UMAP 2") +xlab("UMAP 1")
plot(gg)
dev.off()
#make figure 1D 

#subset object to human TFs
rowData(cds)$gene_short_name=rownames(adr_all_med)
tf_table=read.table("useful_files/Homo_sapiens_TF.txt", sep = "\t", header = T)
tfs=as.character(tf_table$Symbol)
cds_subset = cds[rowData(cds)$gene_short_name %in% tfs,]
subset_pr_test_res <- graph_test(cds_subset, neighbor_graph="principal_graph")
pr_deg_ids <- subset(subset_pr_test_res, q_value < 0.001)
pr_deg_ids = pr_deg_ids[order(pr_deg_ids$q_value),]
#save the table of TFs
write.table(pr_deg_ids, "medulla_de_tfs.txt", row.names = T, col.names = T, quote = F, sep = "\t")
# subset cds to cells in "left" and "right" trajectory

left_cells=WhichCells(adr_all_med, idents = c(25,24,13))
right_cells=WhichCells(adr_all_med, idents = c(25,24,19,20))

left_cds=cds_subset[,left_cells]
right_cds=cds_subset[,right_cells]

left_pseudot=pseudotime(left_cds)
right_pseudot=pseudotime(right_cds)
#make the "left" heatmap of top 100 TFs (sorted by q-value)
m_left = match(rownames(pr_deg_ids),rownames(left_cds))[seq(100)]

pt_group_left = data.frame(cells = names(left_pseudot),
                           group = cut_number(left_pseudot,100))
#aggregate gene expression in each of the 100 pseudotime bins
dat_left = aggregate_gene_expression(left_cds[m_left,],cell_group_df = pt_group_left,scale_agg_value=FALSE)


#make the "right" heatmap of top 100 TFs (sorted by q-value)

m_right = match(rownames(pr_deg_ids),rownames(right_cds))[seq(100)]

pt_group_right = data.frame(cells = names(right_pseudot),
                            group = cut_number(right_pseudot,100))
#aggregate gene expression in each of the 100 pseudotime bins
dat_right = aggregate_gene_expression(right_cds[m_right,],cell_group_df = pt_group_right,scale_agg_value=FALSE)

#merge unscaled data so that SCP parts meet in the middle

dat_left_reverse=dat_left[,order(ncol(dat_left):1)]

joined_unscaled=cbind(dat_left_reverse, dat_right)
#scale data
joined_scaled = t(scale(t(joined_unscaled)))
joined_scaled[abs(joined_scaled)>3] = sign(joined_scaled[abs(joined_scaled)>3]) *3

Heatmap(joined_scaled, cluster_columns = F, cluster_rows=F, row_split = split_genes, show_column_names =F)
#since ComplexHeatmaps doesn't allow reordering of custer levels, run kmeans,
#extract which cluster each gene belongs to, and rarrange the levels in a way that makes sense 
kmeans_1 =kmeans(joined_scaled, 10)
split_genes=kmeans_1$cluster

col_fun = colorRamp2(c(-3,-1,0,1,3), rev(brewer.pal(n = 5, name = "RdYlBu")))
Heatmap(joined_scaled,col=col_fun, cluster_columns = F, cluster_rows=F, row_order = names(sort(split_genes)), show_column_names =F)
```
