## supplementary scripts for "Tumor to normal single cell mRNA comparisons reveal a pan-neuroblastoma cancer cell": FIg2_A-B_sub.R

```
library(Seurat)
library(ComplexHeatmap)
library(circlize)
library(Matrix)
source("gene_sets.R")

s.genes = cc.genes.updated.2019$s.genes
g2m.genes = cc.genes.updated.2019$g2m.genes

#all 10x Neuroblastoma samples
#list of paths to gene_bc_matrices: all_paths = c()
names(all_paths) = c("STDY7685340","STDY7685341","STDY7685342","STDY7843576", "STDY7843577", "STDY7843578","STDY7733084",
                     "STDY7733085", "STDY7733086", "STDY8004894",
                     "STDY8004902", "STDY8004910", "STDY7787237", "STDY7787238", "STDY7787239")
inhouse_srats=Read10X(all_paths)
#processing function
process_inhouse =function(mtx){
  srat=CreateSeuratObject(mtx)
  srat[["percent.mt"]] = PercentageFeatureSet(srat, pattern = "^MT-")
  srat=subset(srat, subset = nFeature_RNA > 300 & nCount_RNA > 1000 & percent.mt < mtCut)
  srat[["percent.hspGenes"]] = PercentageFeatureSet(srat, features=hspGenes[which(hspGenes%in%rownames(srat))])
  srat[["percent.riboGenes"]]=PercentageFeatureSet(srat, features=riboGenes[which(riboGenes%in%rownames(srat))])
  exclude_genes=as.character(read.table("useful_files/excludeGenes.tsv", header = F, sep = "\t")$V1)
  keep_features=rownames(srat)[which(!rownames(srat)%in%exclude_genes)]
  srat=subset(srat, features=keep_features)
  srat = NormalizeData(srat)
  srat =FindVariableFeatures(srat, selection.method = "vst", nfeatures = 2000)
  srat = CellCycleScoring(srat, s.features = s.genes, g2m.features = g2m.genes, set.ident = TRUE)
  srat = ScaleData(srat, features = rownames(srat))
  srat = RunPCA(srat, npcs = 75)
  srat = FindNeighbors(srat, dims=1:75)
  srat = FindClusters(srat, resolution = clusterRes)
  srat = RunUMAP(srat, dims=1:75, min.dist = finalMinDist, n.neighbors = finalNN)
  return(srat)
}
# get 10x tumour Seurat object
srat_inhouse=process_inhouse(inhouse_srats)
#literature markers 

lit_genes=c("SOX10", "MPZ", "PLP1", "ERBB3", "DLL3", "TLX2", "TH", "DBH", "PHOX2B", 
            "CHGB","GAP43", "BCL2", "NPY", "PHOX2A", "PHOX2B","MYCN", "PNMT", "PDGFRB", "TCF21", "PLVAP", "PECAM1", "KDR",
            "PTPRB","HBG1", "HBG2", "HBB","STAR", "MC2R", "PTPRC")
FeaturePlot(srat_inhouse, lit_genes)

#annotate louvain clusters 

srat_inhouse@meta.data$new_idents="x"
srat_inhouse@meta.data$idents_for_plot="x"

srat_inhouse@meta.data$new_idents[which(srat_inhouse@meta.data$seurat_clusters%in%c(1,16,8,25,12,5))]="tumour"
srat_inhouse@meta.data$new_idents[which(srat_inhouse@meta.data$seurat_clusters%in%c(20,14,29,19,0,7,2,9,23,13,
                                                                                    17,15,18,3,6,4,21,22,24))]="leukocytes"
srat_inhouse@meta.data$new_idents[which(srat_inhouse@meta.data$seurat_clusters%in%c(10))]="endothelium"
srat_inhouse@meta.data$new_idents[which(srat_inhouse@meta.data$seurat_clusters%in%c(11))]="mesenchyme"

srat_inhouse@meta.data$idents_for_plot[which(srat_inhouse@meta.data$seurat_clusters%in%c(1,16,8,25))]="Tumour cluster 1"
srat_inhouse@meta.data$idents_for_plot[which(srat_inhouse@meta.data$seurat_clusters%in%c(12))]="Tumour cluster 2"
srat_inhouse@meta.data$idents_for_plot[which(srat_inhouse@meta.data$seurat_clusters%in%c(5))]="Tumour cluster 3"
srat_inhouse@meta.data$idents_for_plot[which(srat_inhouse@meta.data$seurat_clusters%in%c(20,14,29,19,0,7,2,9,23,13,
                                                                                         17,15,18,3,6,4,21,22,24))]="Leukocytes"
srat_inhouse@meta.data$idents_for_plot[which(srat_inhouse@meta.data$seurat_clusters%in%c(10))]="Endothelium"
srat_inhouse@meta.data$idents_for_plot[which(srat_inhouse@meta.data$seurat_clusters%in%c(11))]="Mesenchyme"

#make fig2A
colfunc <- colorRampPalette(c("#084594", "#C6DBEF"))

pdf("inhouse_umap.pdf", height = 3, width=4,useDingbats = F)
gg=DimPlot(srat_inhouse, group.by = "idents_for_plot", label=T,
           cols = c(colfunc(3),"#7E481C", "#DDCC77", "#CC6677"),
           pt.size = 0.1) +theme(text = element_text(size=12), axis.text.x = element_blank(),axis.text.y = element_blank(),
                                 axis.ticks = element_blank(), legend.position = "none")+ylab("UMAP 2") +xlab("UMAP 1")
plot(gg)
dev.off()
#Cluster average expression of marker genes (literature and algorithmic)
#assign annotation as active.ident
srat_inhouse2=srat_inhouse
inhouse_names=names(srat_inhouse@active.ident)
srat_inhouse2@active.ident=factor(srat_inhouse2@meta.data$idents_for_plot)
names(srat_inhouse2@active.ident)=inhouse_names
levels(srat_inhouse2)=c("Tumour cluster 1", "Tumour cluster 2",
                        "Tumour cluster 3","Mesenchyme",
                        "Leukocytes","Endothelium")

inhouse_avg=AverageExpression(srat_inhouse2, return.seurat = T)
inhouse_markers=quickMarkers(srat_inhouse2@assays$RNA@counts, srat_inhouse2@active.ident, N=5)
col_fun = colorRamp2(c(-3, 0, 3), c("blue", "white", "red"))
im1=Heatmap(inhouse_avg@assays$RNA@scale.data[lit_genes,], col = col_fun, cluster_rows = F, cluster_columns = F,
            row_names_gp = gpar(fontsize = 8), name = "Normalized expression")
im2=Heatmap(inhouse_avg@assays$RNA@scale.data[as.character(inhouse_markers$gene),], col=col_fun,cluster_rows = F,
            cluster_columns = F,  row_names_gp = gpar(fontsize = 8), name = "Normalized expression")
im_list = im1 %v% im2 
draw(im_list)

#############################
### process CEL-seq2 data ###
load("scRNAseq_NB_PMC_Full.RData")
Jan_nb = data
genes_10x = read.table("GRCh38/genes.tsv", sep = "\t", header = F)
#main matrix 
mtx = Jan_nb$data$counts
#column metadata 
meta_data = Jan_nb$col.annot
#calculate fraction of spikeins
meta_data$spikeIn_fraction<- colSums(mtx[grep("^ERCC-",rownames(mtx)),])/colSums(mtx)
#remove spikeins
mtx = mtx[-grep("^ERCC-",rownames(mtx)),]
#calculate fraction of reads lost by using 10X ensembl IDs, and fraction of reads lost from  MT genes that are NOT in 10X. 
mtx_10x =mtx[which(rownames(mtx)%in%genes_10x$V1),]
not_10x=mtx[which(!rownames(mtx)%in%genes_10x$V1),]
meta_data$reads_lost =(colSums(mtx)-colSums(mtx_10x))/colSums(mtx)
rownames(not_10x)=Jan_nb$row.annot$gene[match(rownames(not_10x),Jan_nb$row.annot$ensemblId)]
meta_data$mt_reads_lost =colSums(not_10x[grep("^MT-",rownames(not_10x)),])/colSums(mtx)

patient_ids =unique(Jan_nb$col.annot$PatientId)
cells_each_sample =sapply(patient_ids,function(x){
  sapply(unique(Jan_nb$col.annot[which(Jan_nb$col.annot$PatientId==x),]$SampleType), function(y){
    dplyr::filter(Jan_nb$col.annot, PatientId==x, SampleType==y)$CellId
  })
})

cells_list = list("NB039_Tumour_organoids_in_20perc_HP_3D"=cells_each_sample$NB039$`Tumour organoids in 20perc HP (3D)`,
                   "NB039_Tumour_organoids_in_OM_WNT_RSPO_TGFb_inh"=cells_each_sample$NB039$`Tumour organoids in OM_WNT_RSPO_TGFb inh`,
                   "NB039_Tumour_organoids_in_OM_WNT_RSPO"=cells_each_sample$NB039$`Tumour organoids in OM _WNT_RSPO`, 
                   "NB060_Fresh_biopsy"=as.character(cells_each_sample$NB060), 
                   "NB067_Tumour_organoids_in_OM"=as.character(cells_each_sample$NB067),
                   "NB086_Fresh_biopsy"=as.character(cells_each_sample$NB086),
                   "NB098_Fresh_biopsy"=cells_each_sample$NB098$`Fresh biopsy`,
                   "NB098_peripheral_blood_from_fresh_biopsy"=cells_each_sample$NB098$`peripheral blood from fresh biopsy`,
                   "NB098_TILs_from_fresh_biopsy"=cells_each_sample$NB098$`TILs from fresh biopsy`,
                   "NB098_Tumour_organoids"=cells_each_sample$NB098$`Tumour organoids`, 
                   "NB106_Fresh_biopsy"=as.character(cells_each_sample$NB106),
                   "NB107_Fresh_biopsy"=as.character(cells_each_sample$NB107),
                   "NB123_Fresh_biopsy"=as.character(cells_each_sample$NB123),
                   "NB124_Fresh_biopsy"=as.character(cells_each_sample$NB124),
                   "NB125_Fresh_biopsy"=as.character(cells_each_sample$NB125),
                   "NB130_Fresh_biopsy"=as.character(cells_each_sample$NB130),
                   "NB132_Fresh_biopsy"=as.character(cells_each_sample$NB132),
                   "NB138_Fresh_biopsy"=as.character(cells_each_sample$NB138),
                   "NB151_Fresh_biopsy"=as.character(cells_each_sample$NB151),
                   "NB152_Fresh_biopsy"=cells_each_sample$NB152$`Fresh biopsy`,
                   "NB152_Tumour_organoids_P1"=cells_each_sample$NB152$`Tumour organoids P1`)


#add the unique sample to metadata 

meta_data$unique_sample <- "sample"
for (x in 1:length(cells_list)) {
  meta_data$unique_sample[which(meta_data$CellId%in%cells_list[[x]])] <-names(cells_list[x])
}

# going to be working with only 10X from now on, so use mtx_10x,
# convert to gene names, and force uniqueness on each rowname. 

rownames(mtx_10x)<-genes_10x$V2[match(rownames(mtx_10x),genes_10x$V1)]
rownames(mtx_10x) <- make.unique(rownames(mtx_10x))

#function to process PMC data
process_10x =function(mtx, meta_data){
  srat=CreateSeuratObject(mtx, meta.data = meta_data)
  srat[["percent.mt"]] = PercentageFeatureSet(srat, pattern = "^MT-")
  srat=subset(srat, subset = nFeature_RNA > 200 & nCount_RNA > 500 & percent.mt < 20)
  srat[["percent.hspGenes"]] = PercentageFeatureSet(srat, features=hspGenes[which(hspGenes%in%rownames(srat))])
  srat[["percent.riboGenes"]]=PercentageFeatureSet(srat, features=riboGenes[which(riboGenes%in%rownames(srat))])
  exclude_genes=as.character(read.table("excludeGenes.tsv", header = F, sep = "\t")$V1)
  keep_features=rownames(srat)[which(!rownames(srat)%in%exclude_genes)]
  srat=subset(srat, features=keep_features)
  srat = NormalizeData(srat)
  srat =FindVariableFeatures(srat, selection.method = "vst", nfeatures = 2000)
  srat = CellCycleScoring(srat, s.features = s.genes, g2m.features = g2m.genes, set.ident = TRUE)
  srat = ScaleData(srat, features = rownames(srat))
  srat = RunPCA(srat, npcs = 50)
  srat = FindNeighbors(srat, dims=1:50)
  srat = FindClusters(srat, resolution = 1)
  srat = RunUMAP(srat, dims=1:50, min.dist = 0.5, n.neighbors = 50)
  return(srat)
}
#select just the primary tumour data
fresh_biopsies =unique(meta_data$unique_sample)[grep("Fresh",unique(meta_data$unique_sample))]
tumour_only_mtx =mtx_10x[,rownames(meta_data)[which(meta_data$unique_sample%in%fresh_biopsies)]]
tumour_only_meta =meta_data[which(meta_data$unique_sample%in%fresh_biopsies),]
#get processed Seurat object
srat_tumour =process_10x(tumour_only_mtx,tumour_only_meta)


FeaturePlot(srat_tumour, lit_genes)

#annotate louvain clusters

srat_tumour@meta.data$new_idents="x"
srat_tumour@meta.data$idents_for_plot="x"

srat_tumour@meta.data$new_idents[which(srat_tumour@meta.data$seurat_clusters%in%c(22,8))]="tumour"
srat_tumour@meta.data$new_idents[which(srat_tumour@meta.data$seurat_clusters%in%c(13))]="schwannian stroma"
srat_tumour@meta.data$new_idents[which(srat_tumour@meta.data$seurat_clusters%in%c(0,2,11,14,15,18,25,10,12,20))]="mesenchyme"
srat_tumour@meta.data$new_idents[which(srat_tumour@meta.data$seurat_clusters%in%c(1,17,19,21,16,5,9,4,6,26,23,3,24))]="leukocytes"
srat_tumour@meta.data$new_idents[which(srat_tumour@meta.data$seurat_clusters%in%c(7))]="endothelium"

srat_tumour@meta.data$idents_for_plot[which(srat_tumour@meta.data$seurat_clusters%in%c(8))]="Tumour cluster 1"
srat_tumour@meta.data$idents_for_plot[which(srat_tumour@meta.data$seurat_clusters%in%c(22))]="Tumour cluster 2"
srat_tumour@meta.data$idents_for_plot[which(srat_tumour@meta.data$seurat_clusters%in%c(13))]="Schwannian stroma"
srat_tumour@meta.data$idents_for_plot[which(srat_tumour@meta.data$seurat_clusters%in%c(0,2,11,14,15,18,25,10,12,20))]="Mesenchyme"
srat_tumour@meta.data$idents_for_plot[which(srat_tumour@meta.data$seurat_clusters%in%c(1,17,19,21,16,5,9,4,6,26,23,3,24))]="Leukocytes"
srat_tumour@meta.data$idents_for_plot[which(srat_tumour@meta.data$seurat_clusters%in%c(7))]="Endothelium"

#make fig2B

pdf("CEL-seq2_umap.pdf", height = 3, width=4,useDingbats = F)
gg=DimPlot(srat_tumour, group.by = "idents_for_plot",label=T,
           cols = c("#084594", "#4292C6","#7c5295", "#CC6677", "#7E481C","#DDCC77"),
           pt.size = 0.1) +theme(text = element_text(size=12), axis.text.x = element_blank(),axis.text.y = element_blank(),
                                 axis.ticks = element_blank(), 
                                 legend.position = "none")+ylab("UMAP 2") +xlab("UMAP 1")


plot(gg)
dev.off()
#Cluster average expression of marker genes (literature and algorithmic)
#assign annotation as active.ident
srat_tumour2=srat_tumour
dutch_names=names(srat_tumour@active.ident)
srat_tumour2@active.ident=factor(srat_tumour2@meta.data$idents_for_plot)
names(srat_tumour2@active.ident)=dutch_names
levels(srat_tumour2)=c("Tumour cluster 1", "Tumour cluster 2","Schwannian stroma",
                       "Endothelium","Mesenchyme", "Leukocytes")
dutch_avg=AverageExpression(srat_tumour2, return.seurat = T)
dutch_markers=quickMarkers(srat_tumour2@assays$RNA@counts, srat_tumour2@active.ident, N=5)

dm1=Heatmap(dutch_avg@assays$RNA@scale.data[lit_genes,], col = col_fun, cluster_rows = F, cluster_columns = F,
            row_names_gp = gpar(fontsize = 8), name = "Normalized expression")
dm2=Heatmap(dutch_avg@assays$RNA@scale.data[as.character(dutch_markers$gene),], col=col_fun,cluster_rows = F,
            cluster_columns = F,  row_names_gp = gpar(fontsize = 8), name = "Normalized expression")
dm_list= dm1%v% dm2
draw(dm_list)
```
