## supplementary scripts for "Tumor to normal single cell mRNA comparisons reveal a pan-neuroblastoma cancer cell": Fig2_E-G_sub.R

```
library(Seurat)
library(ComplexHeatmap)
library(circlize)
source("logisticRegression.R")
source("gene_sets.R")
adr_all=readRDS("fAdrenal/processedSeurat.RDS")
srat_inhouse=readRDS("srat_inhouse.rds")
srat_tumour=readRDS("srat_dutch.rds")

fetal_pods=readRDS("fetal_podocytes.RDS")
genes_10x = read.table("GRCh38/genes.tsv", sep = "\t", header = F)

#convert IDs in fetal pod data
fetal_pds_counts=fetal_pods@assays$RNA@counts
fetal_pds_counts=fetal_pds_counts[which(rownames(fetal_pds_counts)%in%genes_10x$V1),]
rownames(fetal_pds_counts)=as.character(genes_10x$V2)[match(rownames(fetal_pds_counts), genes_10x$V1)]
rownames(fetal_pds_counts)=make.unique(rownames(fetal_pds_counts))

adr_all@meta.data$new_clust=as.character(adr_all@meta.data$seurat_clusters)
adr_all@meta.data$new_clust[which(adr_all@meta.data$new_clust%in%c("25"))]="SCPs"
adr_all@meta.data$new_clust[which(adr_all@meta.data$new_clust%in%c("13"))]="Chromaffin"
adr_all@meta.data$new_clust[which(adr_all@meta.data$new_clust%in%c("24"))]="Bridge"
adr_all@meta.data$new_clust[which(adr_all@meta.data$new_clust%in%c("19", "20"))]="Sympathoblastic"
adr_all@meta.data$new_clust[which(adr_all@meta.data$new_clust%in%c("3", "15", "14","17"))]="Endothelium"
adr_all@meta.data$new_clust[which(adr_all@meta.data$new_clust%in%c("23", "11", "18"))]="Mesenchyme"
adr_all@meta.data$new_clust[which(adr_all@meta.data$new_clust%in%c("22", "5", "28", "0", "1", "2", "10", "6", "8", "9","21"))]="Cortex"
adr_all@meta.data$new_clust[which(adr_all@meta.data$new_clust%in%c("16", "26"))]="Leukocytes"
adr_all@meta.data$new_clust[which(adr_all@meta.data$new_clust%in%c("30", "7", "4", "12"))]="Erythroblasts"
adr_all@meta.data$new_clust[which(adr_all@meta.data$new_clust%in%c("27", "31", "29"))]="Other"
adr_all@meta.data$new_clust=as.character(adr_all@meta.data$seurat_clusters)

comm_genes=intersect(rownames(fetal_pds_counts), rownames(srat_tumour))
#make sure all datasets have the same genes in the same order
pods_comm_genes=fetal_pds_counts[comm_genes, ]
adr_comm_genes=adr_all@assays$RNA@counts[comm_genes,]
#make a matrix that contains fetal adrenal cells and podocytes
merged_cells=cbind(adr_comm_genes, pods_comm_genes)
cell_labs=c(adr_all@meta.data$new_clust, rep("Podocytes", 278))
#train logistic regression model
fit = trainModel(merged_cells, cell_labs)

ps_dutch = predictSimilarity(fit, srat_tumour@assays$RNA@counts[comm_genes,],srat_tumour@meta.data$idents_for_plot)
ps_inhouse=predictSimilarity(fit, srat_inhouse@assays$RNA@counts[comm_genes,], srat_inhouse@meta.data$idents_for_plot)
ps_self=predictSimilarity(fit, merged_cells, cell_labs)


lr_col_ord=c("SCPs","Bridge","Sympathoblastic","Chromaffin","Mesenchyme","Cortex","Leukocytes", "Endothelium", "Erythroblasts","Other", "Podocytes")
pdf("dutch_lr.pdf", height = 4, width = 7)
print(similarityHeatmap(ps_dutch, column_order=lr_col_ord,row_order=as.character(c("Tumour cluster 1", "Tumour cluster 2",
                                                                                   "Schwannian stroma",
                                                                                   "Mesenchyme", "Leukocytes", "Endothelium"))))
dev.off()

pdf("inhouse_lr.pdf", height = 4, width = 7)
print(similarityHeatmap(ps_inhouse, column_order=lr_col_ord, row_order=c("Tumour cluster 1", "Tumour cluster 2",
                                                                         "Tumour cluster 3","Mesenchyme",
                                                                         "Leukocytes", "Endothelium")))
dev.off()

pdf("self_lr.pdf", height = 4, width = 7)
similarityHeatmap(ps_self, column_order=lr_col_ord, row_order=lr_col_ord)
dev.off()
#get markers for fig3 A-C
quick_with_podos=quickMarkers(merged_cells, cell_labs, FDR=0.01)
quick_with_podos_rel=quick_with_podos[which(quick_with_podos$cluster%in%c("SCPs","Bridge","Sympathoblastic", "Chromaffin","Podocytes")),]
podo_marks_filt=dplyr::filter(quick_with_podos_rel, tfidf>1 & geneFrequencySecondBest <0.2)
```
