## supplementary scripts for "Tumor to normal single cell mRNA comparisons reveal a pan-neuroblastoma cancer cell": Fig3_A-C_sub.R

```
library(ggplot2)
library(reshape2)
#import target data
target_counts=read.table("TARGET-NBL_BulkCountsFrag.tsv",
                         header = T, sep= "\t")
target_genes=read.table("TARGET-NBL_BulkGenes.tsv", sep = "\t",
                        header = T)
target_meta=read.table("TARGET-NBL_BulkMetadata.tsv", sep = "\t")
target_clinical_data=read.table("TARGET_NBL_ClinicalData_withClusters.tsv",
                                sep = "\t", header = T)
target_gene_length= read.table("TARGET-NBL_BulkGeneLengths.tsv", sep = "\t")
#sort out clinical data NAs
names(which(table(sapply(strsplit(as.character(target_clinical_data$TARGET.USI), "-"), "[", 3))==2))

#repeat metadata where samples are from the same patient
target_clinical_data[39,2:ncol(target_clinical_data)]=target_clinical_data[40,2:ncol(target_clinical_data)]
target_clinical_data[42,2:ncol(target_clinical_data)]=target_clinical_data[78,2:ncol(target_clinical_data)]
target_clinical_data[76,2:ncol(target_clinical_data)]=target_clinical_data[125,2:ncol(target_clinical_data)]
target_clinical_data[96,2:ncol(target_clinical_data)]=target_clinical_data[41,2:ncol(target_clinical_data)]
target_clinical_data[102,2:ncol(target_clinical_data)]=target_clinical_data[80,2:ncol(target_clinical_data)]
target_clinical_data[142,2:ncol(target_clinical_data)]=target_clinical_data[1,2:ncol(target_clinical_data)]

#unify the IDs in some capacity
colnames(target_counts)=paste0(sapply(strsplit(colnames(target_counts), "\\."), "[", 4), "-", sapply(strsplit(colnames(target_counts),
                                                                                                              "\\."), "[", 5))
target_clinical_data$TARGET_SHORT=gsub("NA", "01A",paste0(sapply(strsplit(as.character(target_clinical_data$TARGET.USI), "-"), "[", 3),
                                                          "-", sapply(strsplit(as.character(target_clinical_data$TARGET.USI), "-"), "[", 4)))

#convert to log2(TPM+1)
target_length_norm =target_counts/(target_gene_length/1000)

scale_factor=colSums(target_length_norm)/1000000
target_tpm=target_length_norm/scale_factor
target_log2tpm=log2(target_tpm+1)

rownames(target_log2tpm)=make.unique(as.character(target_genes$external_gene_name))
rownames(target_tpm)=make.unique(as.character(target_genes$external_gene_name))
#plot density of mean gene expression
plot(density(rowMeans(target_log2tpm)))
#second peak at 5, make that the threshold
genes_above_threshold=names(rowMeans(target_log2tpm)[rowMeans(target_log2tpm)>5])
percent_above_threshold=apply(target_log2tpm, 1, function(x){
  sum(x>5)/length(x)*100
})
#define risk groups
#all TARGET low risk are stage 4S
low_risk_4s=dplyr::filter(target_clinical_data, target_clinical_data$COG.Risk.Group=="Low Risk")
intermediate_risk=dplyr::filter(target_clinical_data, COG.Risk.Group=="Intermediate Risk")
high_risk=dplyr::filter(target_clinical_data, COG.Risk.Group=="High Risk")

low_risk_4s$risk="low_risk_4s"
intermediate_risk$risk="intermediate_risk"
high_risk$risk="high_risk"
target_risks=rbind(low_risk_4s,intermediate_risk, high_risk)

#import SEQC data

seqc_counts=read.table("seqc/GSE49711_SEQC_NB_TUC_G_log2.txt", sep = "\t", header = T)
seqc_meta=read.csv("seqc/seqc_meta.csv")

rownames(seqc_counts)=seqc_counts$X00gene_id

seqc_counts=seqc_counts[,2:ncol(seqc_counts)]
colnames(seqc_counts)=sapply(strsplit(colnames(seqc_counts), "_"), "[", 2)


seqc_unlist=apply(seqc_counts, 1, function(x){unlist(x)})

seqc_unlist_t=t(seqc_unlist)


#transform from log2(FPKM+1)
untransf=(2^seqc_unlist_t)-1

#transform to log2(tpm+1)
seqc_tpm=apply(untransf,2,function(x){
  (x/sum(x))*1000000
})


seqc_tpm_log2=log2(seqc_tpm+1)

seqc_percent_above_threshold=apply(seqc_tpm_log2, 1, function(x){
  sum(x>5)/length(x)*100
})
#SEQC risks
seqc_lr=dplyr::filter(seqc_meta, Age<540 & MYCN.status=="1", INSS.Stage!="4S")
seqc_4s=dplyr::filter(seqc_meta, INSS.Stage=="4S")
seqc_hr=dplyr::filter(seqc_meta, Age>540 & MYCN.status=="Amp" &INSS.Stage!="4S")
seqpt1=rbind(seqc_lr, seqc_4s, seqc_hr)
seqc_intermediate=seqc_meta[-which(seqc_meta$NB.ID%in%seqpt1$NB.ID),]

seqc_lr$risk="low_risk"
seqc_4s$risk="low_risk_4s"
seqc_hr$risk="high_risk"
seqc_intermediate$risk="intermediate_risk"
seqc_risks=rbind(seqc_lr, seqc_4s, seqc_hr, seqc_intermediate)

seqc_risks$patient=seqc_risks$NB.ID
target_risks$patient=target_risks$TARGET_SHORT


scp_marks=as.character(podo_marks_filt[which(podo_marks_filt$cluster=="SCPs"),]$gene)
bridge_marks=as.character(podo_marks_filt[which(podo_marks_filt$cluster=="Bridge"),]$gene)
chromaffin_marks=as.character(podo_marks_filt[which(podo_marks_filt$cluster=="Chromaffin"),]$gene)
sympathoblastic_marks=as.character(podo_marks_filt[which(podo_marks_filt$cluster=="Sympathoblastic"),]$gene)
pod_marks=as.character(podo_marks_filt[which(podo_marks_filt$cluster=="Podocytes"),]$gene)

t_scp=as.data.frame(percent_above_threshold[scp_marks[which(scp_marks%in%names(percent_above_threshold))]])
t_bridge=as.data.frame(percent_above_threshold[bridge_marks[which(bridge_marks%in%names(percent_above_threshold))]])
t_sympathoblastic=as.data.frame(percent_above_threshold[sympathoblastic_marks[which(sympathoblastic_marks%in%names(percent_above_threshold))]])
t_chromaffin=as.data.frame(percent_above_threshold[chromaffin_marks[which(chromaffin_marks%in%names(percent_above_threshold))]])
t_pods=as.data.frame(percent_above_threshold[pod_marks[which(pod_marks%in%names(percent_above_threshold))]])

t_scp$marker="scp"
t_bridge$marker="bridge"
t_sympathoblastic$marker="sympathoblastic"
t_chromaffin$marker="chromaffin"
t_pods$marker="podocyte"

t_scp$gene=rownames(t_scp)
t_bridge$gene=rownames(t_bridge)
t_sympathoblastic$gene=rownames(t_sympathoblastic)
t_chromaffin$gene=rownames(t_chromaffin)
t_pods$gene=rownames(t_pods)

rownames(t_scp)=1:nrow(t_scp)
rownames(t_bridge)=1:nrow(t_bridge)
rownames(t_sympathoblastic)=1:nrow(t_sympathoblastic)
rownames(t_chromaffin)=1:nrow(t_chromaffin)
rownames(t_pods)=1:nrow(t_pods)

colnames(t_scp)=c("value", "marker", "gene")
colnames(t_bridge)=c("value", "marker", "gene")
colnames(t_sympathoblastic)=c("value", "marker", "gene")
colnames(t_chromaffin)=c("value", "marker","gene")
colnames(t_pods)=c("value", "marker","gene")

t_all_m=rbind(t_scp, t_bridge,t_sympathoblastic,t_chromaffin, t_pods)

seqc_scp=as.data.frame(seqc_percent_above_threshold[scp_marks[which(scp_marks%in%names(seqc_percent_above_threshold))]])
seqc_bridge=as.data.frame(seqc_percent_above_threshold[bridge_marks[which(bridge_marks%in%names(seqc_percent_above_threshold))]])
seqc_sympathoblastic=as.data.frame(seqc_percent_above_threshold[sympathoblastic_marks[which(sympathoblastic_marks%in%names(seqc_percent_above_threshold))]])
seqc_chromaffin=as.data.frame(seqc_percent_above_threshold[chromaffin_marks[which(chromaffin_marks%in%names(seqc_percent_above_threshold))]])
seqc_pods=as.data.frame(seqc_percent_above_threshold[pod_marks[which(pod_marks%in%names(seqc_percent_above_threshold))]])

seqc_scp$marker="scp"
seqc_bridge$marker="bridge"
seqc_sympathoblastic$marker="sympathoblastic"
seqc_chromaffin$marker="chromaffin"
seqc_pods$marker="podocyte"

seqc_scp$gene=rownames(seqc_scp)
seqc_bridge$gene=rownames(seqc_bridge)
seqc_sympathoblastic$gene=rownames(seqc_sympathoblastic)
seqc_chromaffin$gene=rownames(seqc_chromaffin)
seqc_pods$gene=rownames(seqc_pods)

rownames(seqc_scp)=1:nrow(seqc_scp)
rownames(seqc_bridge)=1:nrow(seqc_bridge)
rownames(seqc_sympathoblastic)=1:nrow(seqc_sympathoblastic)
rownames(seqc_chromaffin)=1:nrow(seqc_chromaffin)
rownames(seqc_pods)=1:nrow(seqc_pods)

colnames(seqc_scp)=c("value", "marker", "gene")
colnames(seqc_bridge)=c("value", "marker", "gene")
colnames(seqc_sympathoblastic)=c("value", "marker", "gene")
colnames(seqc_chromaffin)=c("value", "marker","gene")
colnames(seqc_pods)=c("value", "marker","gene")

seqc_all_m=rbind(seqc_scp, seqc_bridge,seqc_sympathoblastic,seqc_chromaffin, seqc_pods)
t_all_m$dataset="TARGET"
seqc_all_m$dataset="SEQC"
joined_3a=rbind(t_all_m,seqc_all_m)


joined_3a$marker=factor(joined_3a$marker, levels = c("podocyte","scp", "bridge", "sympathoblastic", "chromaffin"))
joined_3a$dataset=factor(joined_3a$dataset, levels = c("TARGET", "SEQC"))
#Figure 3A
pdf("marks_all.pdf", width = 4, height = 3, useDingbats = F)
gg=ggplot(data = joined_3a, aes(x=marker,y=value, group=dataset)) +
  geom_quasirandom(mapping = aes(x=marker,y=value, group=dataset), dodge.width=.8, shape=19, cex=0.5, alpha=0.2) +
  geom_pointrange(mapping = aes(x = marker, y = value, shape=dataset),
                  stat = "summary",
                  fun.ymin = function(z) {quantile(z,0.25)},
                  fun.ymax = function(z) {quantile(z,0.75)},
                  fun.y = median, fill="black", 
                  position=position_dodge(width=0.8)) +  theme_bw() +
  theme(panel.grid.major = element_blank(),
        panel.grid.minor = element_blank(),
        strip.background = element_blank(),
        panel.border = element_rect(colour = "black"),
        text=element_text(size = 16),
        axis.text.x = element_text(angle = 90, hjust = 1)) + labs(x="Markers", y = "% samples above expression threshold")
plot(gg)
dev.off()


#non-medullary samples (target)
#exclude adrenal or putatively adrrnal samples
leftover_ids=as.character(target_clinical_data$ICDO.Description)[-grep("adrenal|abdominal|kidney|abdomen|unknown|retroperitoneum|other", tolower(as.character(target_clinical_data$ICDO.Description)))]
#exclude empty strings
leftover_ids=leftover_ids[c(1:13,16:23)]

leftover_samples=target_clinical_data$TARGET_SHORT[which(target_clinical_data$ICDO.Description%in%leftover_ids)]

os_percent_above_threshold=apply(target_log2tpm[,leftover_samples], 1, function(y){
  sum(y>5)/length(y)*100
})


os_scp=as.data.frame(os_percent_above_threshold[scp_marks[which(scp_marks%in%names(os_percent_above_threshold))]])
os_bridge=as.data.frame(os_percent_above_threshold[bridge_marks[which(bridge_marks%in%names(os_percent_above_threshold))]])
os_sympathoblastic=as.data.frame(os_percent_above_threshold[sympathoblastic_marks[which(sympathoblastic_marks%in%names(os_percent_above_threshold))]])
os_chromaffin=as.data.frame(os_percent_above_threshold[chromaffin_marks[which(chromaffin_marks%in%names(os_percent_above_threshold))]])
os_pods=as.data.frame(os_percent_above_threshold[pod_marks[which(pod_marks%in%names(os_percent_above_threshold))]])

os_scp$marker="scp"
os_bridge$marker="bridge"
os_sympathoblastic$marker="sympathoblastic"
os_chromaffin$marker="chromaffin"
os_pods$marker="podocytes"

os_scp$gene=rownames(os_scp)
os_bridge$gene=rownames(os_bridge)
os_sympathoblastic$gene=rownames(os_sympathoblastic)
os_chromaffin$gene=rownames(os_chromaffin)
os_pods$gene=rownames(os_pods)

rownames(os_scp)=1:nrow(os_scp)
rownames(os_bridge)=1:nrow(os_bridge)
rownames(os_sympathoblastic)=1:nrow(os_sympathoblastic)
rownames(os_chromaffin)=1:nrow(os_chromaffin)
rownames(os_pods)=1:nrow(os_pods)

colnames(os_scp)=c("value", "marker", "gene")
colnames(os_bridge)=c("value", "marker", "gene")
colnames(os_sympathoblastic)=c("value", "marker", "gene")
colnames(os_chromaffin)=c("value", "marker","gene")
colnames(os_pods)=c("value", "marker","gene")

os_all_m=rbind(os_scp, os_bridge,os_sympathoblastic,os_chromaffin, os_pods)

os_all_m$marker=factor(os_all_m$marker, levels=c("podocytes" , "scp","bridge","sympathoblastic","chromaffin"))
#Figure 3B
pdf("marks_om.pdf", width = 3, height = 3, useDingbats = F)
ggplot(data = os_all_m, aes(x=marker,y=value)) +
  geom_quasirandom(mapping = aes(x=marker,y=value), alpha=0.2, cex=0.5, shape=19) +
  geom_pointrange(mapping = aes(x = marker, y = value),
                  stat = "summary", shape=22,
                  fun.ymin = function(z) {quantile(z,0.25)},
                  fun.ymax = function(z) {quantile(z,0.75)},
                  fun.y = median, fill="black") +  theme_bw() +
  theme(panel.grid.major = element_blank(),
        panel.grid.minor = element_blank(),
        strip.background = element_blank(),
        panel.border = element_rect(colour = "black"),
        text=element_text(size = 16),
        axis.text.x = element_text(angle = 90, hjust = 1))
dev.off()

#per risk group function

get_per_risk=function(count_tab, risk_tab,th){
  values_per_risk_group=sapply(unique(risk_tab$risk), function(x){
    pts=risk_tab$patient[which(risk_tab$risk==x)]
    tab=count_tab[,pts]
    apply(tab, 1, function(y){
      sum(y>th)/length(y)*100
    })
  })
  
  values_per_risk_group=data.frame(values_per_risk_group)
  values_per_risk_group$gene=rownames(values_per_risk_group)
  
  scp_markers_in_rg=values_per_risk_group[scp_marks[which(scp_marks%in%rownames(values_per_risk_group))],]
  bridge_markers_in_rg=values_per_risk_group[bridge_marks[which(bridge_marks%in%rownames(values_per_risk_group))],]
  sympathoblastic_markers_in_rg=values_per_risk_group[sympathoblastic_marks[which(sympathoblastic_marks%in%rownames(values_per_risk_group))],]
  chromaffin_markers_in_rg=values_per_risk_group[chromaffin_marks[which(chromaffin_marks%in%rownames(values_per_risk_group))],]
  podo_markers_in_rg=values_per_risk_group[pod_marks[which(pod_marks%in%rownames(values_per_risk_group))],]
  
  scp_markers_in_rg=melt(scp_markers_in_rg, id.vars = "gene", measure.vars = as.character(unique(risk_tab$risk)))
  bridge_markers_in_rg=melt(bridge_markers_in_rg, id.vars = "gene", measure.vars = as.character(unique(risk_tab$risk)))
  sympathoblastic_markers_in_rg=melt(sympathoblastic_markers_in_rg, id.vars = "gene", measure.vars = as.character(unique(risk_tab$risk)))
  chromaffin_markers_in_rg=melt(chromaffin_markers_in_rg, id.vars = "gene", measure.vars = as.character(unique(risk_tab$risk)))
  podo_markers_in_rg=melt(podo_markers_in_rg, id.vars = "gene", measure.vars = as.character(unique(risk_tab$risk)))
  
  scp_markers_in_rg$marker="scp"
  bridge_markers_in_rg$marker="bridge"
  sympathoblastic_markers_in_rg$marker="sympathoblastic"
  chromaffin_markers_in_rg$marker="chromaffin"
  podo_markers_in_rg$marker="podocyte"
  
  all_in_groups=rbind(scp_markers_in_rg, bridge_markers_in_rg, sympathoblastic_markers_in_rg, chromaffin_markers_in_rg, podo_markers_in_rg)
  
  return(all_in_groups)
  
}

target_per_risk=get_per_risk(target_log2tpm, target_risks, 5)
seqc_per_risk=get_per_risk(seqc_tpm_log2, seqc_risks, 5)

target_per_risk$cohort="TARGET"
seqc_per_risk$cohort="SEQC"

combined_per_risk=rbind(target_per_risk, seqc_per_risk)

colnames(combined_per_risk)=c("gene", "risk", "value", "marker", "cohort")

combined_per_risk$risk=factor(combined_per_risk$risk, levels=c("low_risk","low_risk_4s", "intermediate_risk", "high_risk"))

combined_per_risk$marker=factor(combined_per_risk$marker, levels=c("podocyte" , "scp","bridge","sympathoblastic","chromaffin"))

combined_per_risk$cohort=factor(combined_per_risk$cohort, levels = c("TARGET", "SEQC"))
#Figure 3C
ggplot(data = combined_per_risk, aes(x=marker,y=value)) +
  geom_quasirandom(shape=19, alpha=0.2, cex=0.5)+facet_grid(vars(risk), vars(cohort))+
  geom_pointrange(mapping = aes(x = marker, y = value),
                  stat = "summary",
                  fun.ymin = function(z) {quantile(z,0.25)},
                  fun.ymax = function(z) {quantile(z,0.75)},
                  fun.y = median, shape=22, fill="black") +
  theme_bw() +
  theme(panel.grid.major = element_blank(),
        panel.grid.minor = element_blank(),
        strip.background = element_blank(),
        panel.border = element_rect(colour = "black"),
        text=element_text(size = 14),
        axis.text.x = element_text(angle = 90, hjust = 1))
```
