## supplementary scripts for "Tumor to normal single cell mRNA comparisons reveal a pan-neuroblastoma cancer cell": Fig4_A-B_sub.R

```
# comparison of cancer cells and normal medulla 
#get medullary cells 
adr_all=readRDS("fAdrenal/processedSeurat.RDS")
adr_all_med=subset(adr_all, idents=c(25,24,13,19,20))

tum_cells_inhouse=subset(srat_inhouse, idents = c(1,5,8,12,16,25))
tum_cells_dutch=subset(srat_tumour, idents = c(22,8))

#assign all tumour cells the same identity  
tum_cells_inhouse@meta.data$new_idents="tumour"
tum_cells_dutch@meta.data$new_idents="tumour"
adr_all_med@meta.data$new_idents=as.character(adr_all_med@meta.data$seurat_clusters) 
#combine each dataset with medulla cells and then find markers
combined_inhouse=merge(adr_all_med, tum_cells_inhouse)
inhouse_markers=quickMarkers(combined_inhouse@assays$RNA@counts, combined_inhouse@meta.data$new_idents, FDR = 0.01, N=Inf)
combined_dutch=merge(adr_all_med, tum_cells_dutch)
dutch_markers=quickMarkers(combined_dutch@assays$RNA@counts, combined_dutch@meta.data$new_idents, FDR = 0.01, N=Inf)
#filter by frequency in 2nd best cluster
rg_tc=org_markers[which(org_markers$cluster=="tumour"),]
org_tc_filt=org_tc[which(org_tc$geneFrequencySecondBest<0.2),]


inhouse_tc=inhouse_markers[which(inhouse_markers$cluster=="tumour"),]
inhouse_tc_filt=inhouse_tc[which(inhouse_tc$geneFrequencySecondBest<0.2),]

#calculate mean tfidf and sort by it
no_org_lj=full_join(inhouse_tc_filt, dutch_tc_filt, by='gene')
no_org_lj$mean_tfidf=rowMeans(no_org_lj[,c(8,17)], na.rm = T)
no_org_final=no_org_lj[no_org_lj$mean_tfidf>0.85,]
no_org_final_sorted=no_org_final[order(no_org_final$mean_tfidf, decreasing = T),]


#exclude genes absent from TARGET as we use it for validation
no_org_final=no_org_final[which(no_org_final$gene%in%rownames(target_log2tpm)),]

#exclude genes expressed in more than 25% of leukocyte cells in either sample as these are likely contaminants
inhouse_immune_cells=rownames(srat_inhouse@meta.data)[which(srat_inhouse@meta.data$idents_for_plot=="Leukocytes")]
inhouse_immune_cellexp=apply(srat_inhouse@assays$RNA@counts[,inhouse_immune_cells],1,function(y){
  sum(y>0)/length(y)*100
})

dutch_immune_cells=rownames(srat_tumour@meta.data)[which(srat_tumour@meta.data$idents_for_plot=="Leukocytes")]
dutch_immune_cellexp=apply(srat_tumour@assays$RNA@counts[,dutch_immune_cells],1,function(y){
  sum(y>0)/length(y)*100
})

immune_inhouse_t25=tum_de_genes[which(tum_de_genes%in%names(inhouse_immune_cellexp[inhouse_immune_cellexp>25]))]
dutch_immune_t25=tum_de_genes[which(tum_de_genes%in%names(dutch_immune_cellexp[dutch_immune_cellexp>25]))]

immune_sc=unique(c(immune_inhouse_t50, dutch_immune_t50))
final_de=no_org_final[which(!no_org_final$gene%in%immune_sc),]

a=gsub("\\.x", ".GOSH", colnames(final_de))
b=gsub("\\.y", ".PMC", a)

colnames(final_de)=b
rownames(final_de)=1:nrow(final_de)

final_de_sorted=final_de[order(final_de$mean_tfidf, decreasing = T),]


write.table(final_de_sorted, "final_de_sorted.txt", sep = "\t", row.names = F, col.names = T, quote = F)

#differential expression in TARGET


target_counts=read.table("/lustre/scratch119/casm/team274sb/ek12/Neuroblastoma/bulk_NB/TARGET-NBL_BulkCountsFrag.tsv",
                         header = T, sep= "\t")
target_genes=read.table("/lustre/scratch119/casm/team274sb/ek12/Neuroblastoma/bulk_NB/TARGET-NBL_BulkGenes.tsv", sep = "\t",
                        header = T)
target_meta=read.table("/lustre/scratch119/casm/team274sb/ek12/Neuroblastoma/bulk_NB/TARGET-NBL_BulkMetadata.tsv", sep = "\t")
target_clinical_data=read.table("/lustre/scratch119/casm/team274sb/ek12/Neuroblastoma/bulk_NB/TARGET_NBL_ClinicalData_withClusters.tsv",
                                sep = "\t", header = T)
target_gene_length= read.table("/lustre/scratch119/casm/team274sb/ek12/Neuroblastoma/bulk_NB/TARGET-NBL_BulkGeneLengths.tsv", sep = "\t")
#sort out clinical data NAs
names(which(table(sapply(strsplit(as.character(target_clinical_data$TARGET.USI), "-"), "[", 3))==2))

df_bla=target_clinical_data[grep("PARBAJ|PARHAM|PASGAP|PASNPG|PATYIL|PAUDDK",target_clinical_data$TARGET.USI),]

target_clinical_data[39,2:ncol(target_clinical_data)]=target_clinical_data[40,2:ncol(target_clinical_data)]
target_clinical_data[42,2:ncol(target_clinical_data)]=target_clinical_data[78,2:ncol(target_clinical_data)]
target_clinical_data[76,2:ncol(target_clinical_data)]=target_clinical_data[125,2:ncol(target_clinical_data)]
target_clinical_data[96,2:ncol(target_clinical_data)]=target_clinical_data[41,2:ncol(target_clinical_data)]
target_clinical_data[102,2:ncol(target_clinical_data)]=target_clinical_data[80,2:ncol(target_clinical_data)]
target_clinical_data[142,2:ncol(target_clinical_data)]=target_clinical_data[1,2:ncol(target_clinical_data)]

#unify the IDs in some capacity
colnames(target_counts)=paste0(sapply(strsplit(colnames(target_counts), "\\."), "[", 4), "-", sapply(strsplit(colnames(target_counts),
                                                                                                              "\\."), "[", 5))
target_clinical_data$TARGET_SHORT=gsub("NA", "01A",paste0(sapply(strsplit(as.character(target_clinical_data$TARGET.USI), "-"), "[", 3),
                                                          "-", sapply(strsplit(as.character(target_clinical_data$TARGET.USI), "-"), "[", 4)))


#define risk groups
low_risk_4s=dplyr::filter(target_clinical_data, target_clinical_data$COG.Risk.Group=="Low Risk")
intermediate_risk=dplyr::filter(target_clinical_data, COG.Risk.Group=="Intermediate Risk")
high_risk=dplyr::filter(target_clinical_data, COG.Risk.Group=="High Risk")

low_risk_4s$risk="low_risk_4s"
intermediate_risk$risk="intermediate_risk"
high_risk$risk="high_risk"
all=rbind(low_risk_4s,intermediate_risk, high_risk, high_risk_relapse)
extremes=rbind(low_risk_4s, high_risk, high_risk_relapse)

#subset to only include low and high risk samples
extreme_count_tab=target_counts[,extremes$TARGET_SHORT]

#make an EdgeR object 

edg_file=DGEList(counts=extreme_count_tab, samples = extremes, genes = target_genes, group=extremes$risk)

o = order(rowSums(edg_file$counts), decreasing=TRUE)
edg_file = edg_file[o,]
d = duplicated(edg_file$genes$external_gene_name)
edg_file = edg_file[!d,]

rownames(edg_file)=edg_file$genes$external_gene_name

keep = filterByExpr(edg_file)
table(keep)

edg_file = edg_file[keep, , keep.lib.sizes=FALSE]

edg_file=calcNormFactors(edg_file)

edg_file$samples$group=factor(edg_file$samples$group, levels = c("low_risk_4s", "high_risk"))
#design a model matrix. age at diagnosis and mycn status included as covariates
design=model.matrix(~group+Age.at.Diagnosis.in.Days+MYCN.status,edg_file$samples)
fit=glmQLFit(edg_file, design, robust=TRUE, dispersion = 0.4^2)
qlf=glmQLFTest(fit,coef=2)

de_of_de=qlf$table[as.character(final_de_sorted$gene)[which(as.character(final_de_sorted$gene)%in%rownames(qlf$table))],]
de_of_de_sorted=de_of_de[order(abs(de_of_de$logFC), decreasing = T),]
de_of_de_sorted$qval=p.adjust(de_of_de_sorted$PValue, method = "fdr")


de_of_de_pat=as.data.frame(percent_above_threshold[rownames(de_of_de_sorted)])
de_of_de_pat$gene=rownames(de_of_de_pat)

colnames(de_of_de_pat)=c("value", "gene")
de_of_de_pat=de_of_de_pat[order(de_of_de_pat$value, decreasing = T),]
de_of_de_pat$gene=factor(de_of_de_pat$gene, levels = de_of_de_pat$gene)

#% TARGET samples expressed above threshold 
ggplot(de_of_de_pat, aes(x=reorder(gene, value), y=value))+geom_bar(stat="identity", width = 0.6) +coord_flip() +
  theme_bw()+
  theme(axis.text.y = element_text(colour = "black", face = "italic", size = 8),
        panel.grid.major = element_blank(),
        panel.grid.minor = element_blank(),
        strip.background = element_blank(),
        panel.border = element_rect(colour = "black"),
        text=element_text(size = 12)) + labs( y ="% of samples above expression threshold", x = "Gene")


#plot some genes in SEQC and TARGET
genes_for_kap=c("ALDH1A2", "PHF24", "CCDC144NL-AS1", "NCAN", "PRAME", "CRYBB2", "SIX3", "CPEB1", "TMEM163", "LSMEM1", "EPB41L4B", "RAP1GAP2", "SLC9A7")

de_of_de_facet_tab=target_log2tpm[genes_for_kap, all$TARGET_SHORT]

de_tab=as.data.frame(t(de_of_de_facet_tab))  
de_tab$risk=all$risk  
de_tab$patient=rownames(de_tab)
#assign low risk 4s samples to low risk category to match with SEQC
de_tab_melted=melt(de_tab, id.vars=c("patient", "risk"))
de_tab_melted$risk[which(de_tab_melted$risk=="low_risk_4s")]="low_risk"
de_tab_melted$risk=factor(de_tab_melted$risk, levels = c("low_risk", "intermediate_risk","high_risk"))

de_tab_melted$variable=factor(de_tab_melted$variable, levels = c(rownames(de_of_de_sorted)))

genes_for_seqc=genes_for_kap[which(genes_for_kap%in%rownames(seqc_tpm_log2))]

seqc_facet_tab=seqc_tpm_log2[genes_for_seqc, seqc_risk$patient]
seqc_facet_tab=as.data.frame(t(seqc_facet_tab))

seqc_facet_tab$risk=seqc_risk$risk
seqc_facet_tab$patient=rownames(seqc_facet_tab)

seqc_facet_tab_melted=melt(seqc_facet_tab, id.vars=c("patient", "risk"))

seqc_facet_tab_melted$risk=factor(seqc_facet_tab_melted$risk, levels = c("low_risk_4s", "intermediate_risk","high_risk"))

de_tab_melted$dataset="TARGET"
seqc_facet_tab_melted$dataset="SEQC"

de_of_de_two_datasets=rbind(de_tab_melted, seqc_facet_tab_melted)

#make Figure 4
ggplot(data = de_of_de_two_datasets, aes(x=risk,y=value, group=dataset)) +
  geom_quasirandom(shape=1, alpha=0.3)+facet_wrap(~variable,scales="free_y")+
  geom_pointrange(mapping = aes(x = risk, y = value),
                  stat = "summary",
                  fun.ymin = function(z) {quantile(z,0.25)},
                  fun.ymax = function(z) {quantile(z,0.75)},
                  fun.y = median, shape=22, fill="black",
                  position=position_dodge(width=0.8)) +
  theme_bw() +
  theme(panel.grid.major = element_blank(),
        panel.grid.minor = element_blank(),
        strip.background = element_blank(),
        panel.border = element_rect(colour = "black"),
        text=element_text(size = 14),
        axis.text.x = element_text(angle = 90, hjust = 1))
```
