## supplementary scripts for "Tumor to normal single cell mRNA comparisons reveal a pan-neuroblastoma cancer cell": geneSets.R

```
#Housekeeping genes
#hkGenes = read.table('http://www.tau.ac.il/~elieis/HKG/HK_genes.txt',sep='\t')
hkGenes = read.table('~/trueHome/scratch/Common/HK_genes.txt',sep='\t')
hkGenes = gsub(' ','',hkGenes[,1])
#Taken from here Table S2
#http://journals.plos.org/plosone/article?id=10.1371/journal.pone.0012862
nuclearMTGenes = c("ABCB10", "ABCD3", "ACADM", "ACP6", "AGMAT", "AK2", "AKR7A2",
"ALDH4A1", "ALDH9A1", "APOA1BP", "ATAD3A", "ATAD3B", "ATP5F1",
"ATPAF1", "ATPIF1", "AURKAIP1", "BOLA1", "BRP44", "C1orf151",
"C1orf31", "C1orf69", "CABC1", "CCBL2", "CLIC4", "CPT2", "CYB5R1",
"DAP3", "DARS2", "DBT", "DDAH1", "DNAJC11", "FAM36A", "FDPS",
"FH", "GLRX2", "GNG5", "HAO2", "HBXIP", "HMGCL", "HMGCS2", "HSD3B1",
"HSPB7", "IARS2", "KIF1B", "MECR", "MFN2", "MMACHC", "MOSC1",
"MOSC2", "MRPL20", "MRPL24", "MRPL37", "MRPL55", "MRPL9", "MRPS14",
"MRPS15", "MRPS21", "MTX1", "MUL1", "MUTYH", "NDUFS2", "NDUFS5",
"NIT1", "NRD1", "OMA1", "PARK7", "PARS2", "PEX11B", "PINK1",
"PPOX", "PRDX1", "PRDX6", "SARS", "SCCPDH", "SCP2", "SDHB", "SDHC",
"SLC16A1", "SLC25A24", "SLC25A33", "SLC25A34", "SLC25A44", "TARS2",
"TATDN3", "TDRKH", "TFB2M", "THEM4", "THEM5", "TIMM17A", "TOMM20",
"TOMM40L", "TRIT1", "UQCRH", "WARS2", "ABCB6", "ACADL", "ACYP2",
"AGPS", "AGXT", "ATIC", "ATP5G3", "BCS1L", "C2orf47", "C2orf56",
"CASP8", "CCT7", "COQ10B", "COX5B", "COX7A2L", "CPS1", "CYP27A1",
"DGUOK", "DNAJC27", "EFHD1", "FAM136A", "FASTKD2", "FSIP2", "GLS",
"GPD2", "HADHA", "HADHB", "HIBCH", "HK2", "HSPD1", "HSPE1", "HTRA2",
"IDH1", "IMMT", "KYNU", "LIPT1", "LRPPRC", "MAP1D", "MARS2",
"MCEE", "MDH1", "MPV17", "MRPL19", "MRPL30", "MRPL33", "MRPL35",
"MRPL44", "MRPS9", "MTHFD2", "MTIF2", "MTX2", "NDUFA10", "NDUFB3",
"NDUFS1", "NFU1", "NIF3L1", "OSGEPL1", "PDK1", "PNKD", "PNPT1",
"PTCD3", "RAB11FIP5", "RDH14", "RNASEH1", "SFXN5", "SLC25A12",
"SLC30A6", "SPR", "SUCLG1", "UXS1", "VAMP8", "ABHD10", "ACAA1",
"ACAD11", "ACAD9", "ALAS1", "ALDH1L1", "AMT", "BDH1", "C3orf1",
"C3orf23", "C3orf31", "C3orf60", "CCDC51", "CCDC58", "CHCHD4",
"CHDH", "CMC1", "COX17", "CPOX", "EEFSEC", "EHHADH", "EXOG",
"FAM162A", "GFM1", "GMPPB", "GPX1", "GTPBP8", "HEMK1", "HIGD1A",
"LARS2", "MCCC1", "MFN1", "MOBP", "MRPL3", "MRPL47", "MRPS22",
"MRPS25", "NDUFB4", "NDUFB5", "NME6", "OGG1", "OPA1", "OXNAD1",
"OXSM", "PARL", "PCCB", "PDE12", "PDHB", "RG9MTD1", "RHOA", "RPL35A",
"SLC25A20", "SLC25A26", "SLC25A36", "SUCLG2", "TOMM70A", "TRNT1",
"UQCRC1", "AADAT", "ACOX3", "ACSL1", "ATP10D", "ATP5I", "C4orf14",
"CBR4", "COQ2", "COX18", "ETFDH", "GK2", "GRPEL1", "GRSF1", "GUF1",
"HADH", "LAP3", "LETM1", "LIAS", "MRPL1", "MRPS18C", "MTHFD2L",
"NFXL1", "OCIAD1", "PDHA2", "PET112L", "PIGY", "PPA2", "PPM1K",
"QDPR", "RPL34", "SLC25A31", "SLC25A4", "UCP1", "AGXT2", "AGXT2L2",
"ALDH7A1", "AMACR", "CKMT2", "DHX29", "DMGDH", "FTMT", "GFM2",
"GLRX", "GRPEL2", "HARS2", "HIGD2A", "HK3", "HSD17B4", "HSPA9",
"ISCA1L", "KIAA0141", "LOC153328", "LOC727947", "LYRM7", "MAT2B",
"MCCC2", "MRPL22", "MRPL36", "MRPS27", "MRPS30", "MRPS36", "NDUFA2",
"NDUFAF2", "NDUFS4", "NDUFS6", "NLN", "NNT", "OXCT1", "PCBD2",
"PRELID1", "PRELID2", "PTCD2", "RAB24", "RARS", "RPS14", "SDHA",
"SFXN1", "SLC22A4", "SLC25A2", "SLC25A46", "THG1L", "UQCRQ",
"VDAC1", "AARS2", "ACAT2", "ALDH5A1", "BAK1", "BCKDHB", "BPHL",
"BRP44L", "C6orf125", "C6orf203", "C6orf57", "C6orf66", "CCDC90A",
"COQ3", "COX7A2", "DACT2", "FARS2", "HDDC2", "HSD17B8", "LACE1",
"LOC643233", "LYRM4", "ME1", "MRPL14", "MRPL18", "MRPL2", "MRPS10",
"MRPS18A", "MRPS18B", "MRS2", "MTCH1", "MTHFD1L", "MTO1", "MTRF1L",
"MUT", "PACRG", "PDSS2", "PECI", "PRICKLE4", "PRSS35", "QRSL1",
"RAB32", "RARS2", "RMND1", "RPL10A", "RTN4IP1", "SIRT5", "SLC25A27",
"SOD2", "TAP1", "TFB1M", "THEM2", "TMEM14C", "VARS2", "AASS",
"ABCB8", "ABCF2", "ABHD11", "ACN9", "ADCK2", "AGK", "AGR2", "ARMC10",
"ATP5J2", "C7orf10", "C7orf55", "CCDC136", "CHCHD2", "CHCHD3",
"CROT", "CYCS", "DLD", "DNAJC30", "ELN", "FASTK", "FIS1", "FTSJ2",
"GARS", "GBAS", "HIBADH", "IMMP2L", "IQCE", "MDH2", "MRPL32",
"MRPS17", "MRPS24", "MRPS33", "MTERF", "MYLC2PL", "NDUFA4", "NDUFA5",
"NDUFB2", "NT5C3", "OGDH", "PDK4", "PMPCB", "SLC25A13", "SLC25A40",
"SND1", "SSBP1", "TBRG4", "tcag7.1260", "TOMM7", "ADCK5", "ADHFE1",
"AGPAT5", "ARMC1", "C8orf38", "CHCHD7", "COX6C", "CYC1", "CYP11B1",
"CYP11B2", "DECR1", "DUSP26", "FAM82B", "GSR", "HRSP12", "LACTB2",
"MRPL13", "MRPL15", "MRPS28", "MSRA", "MTERFD1", "MTFR1", "NDUFB9",
"OXR1", "PPM2C", "SLC25A32", "SLC25A37", "STAR", "TMEM65", "TMEM70",
"TOP1MT", "TTC35", "UQCRB", "VDAC3", "ACO1", "AK3", "ALDH1B1",
"AUH", "C9orf46", "COQ4", "CRAT", "ENDOG", "FPGS", "FXN", "GLDC",
"HINT2", "HSDL2", "ISCA1", "MCART1", "MRPL41", "MRPL50", "MRPS2",
"MRRF", "NDUFA8", "NDUFB6", "NIPSNAP3A", "NIPSNAP3B", "NUDT2",
"PMPCA", "PTGES2", "PTRH1", "RFK", "SARDH", "SECISBP2", "SLC25A25",
"STOML2", "SURF1", "TXN", "ACADSB", "ACSL5", "ADO", "AIFM2",
"ALDH18A1", "ANKRD26", "ARMC4", "AS3MT", "ASAH2", "ATAD1", "ATP5C1",
"C10orf10", "C10orf2", "C10orf58", "C10orf65", "CBARA1", "CCDC109A",
"CHAT", "CHCHD1", "CISD1", "COMTD1", "COX15", "DHTKD1", "ECHDC3",
"ECHS1", "GHITM", "GLUD1", "GPAM", "HK1", "IDE", "IDI1", "MRPL43",
"MRPS16", "MSRB2", "MTG1", "MTPAP", "NCOA4", "NUDT13", "OAT",
"OGDHL", "PDSS1", "PHYH", "PHYHIPL", "PITRM1", "PPIF", "PRDX3",
"PSTK", "SFXN2", "SFXN3", "SFXN4", "SLC25A16", "SLC25A28", "SUPV3L1",
"TFAM", "VDAC2", "YME1L1", "ACAD8", "ACAT1", "ATP5L", "BAD",
"BBOX1", "C11orf79", "CCDC90B", "CLPB", "COX8A", "CPT1A", "CYB5R2",
"DLAT", "DNAJC4", "FDX1", "FLJ14213", "FOXRED1", "FTH1", "FXC1",
"GLYAT", "LDHA", "LOC387787", "ME3", "MRPL16", "MRPL17", "MRPL21",
"MRPL23", "MRPL48", "MRPL49", "MTCH2", "NARS2", "NDUFC2", "NDUFS3",
"NDUFS8", "NDUFV1", "NLRX1", "NUDT8", "PC", "PDHX", "PRDX5",
"PTS", "REXO2", "RPUSD4", "SDHD", "SIRT3", "TCIRG1", "TIMM10",
"TIMM8B", "TMEM126A", "TMEM223", "UCP2", "UCP3", "ZBED5", "ABCD2",
"ACACB", "ACAD10", "ACADS", "ACSM4", "ALDH1L2", "ALDH2", "ATP5G2",
"BCAT1", "BLOC1S1", "C12orf62", "COQ10A", "COQ5", "COX6A1", "CRY1",
"CS", "CYP27B1", "DIABLO", "GATC", "GLS2", "ISCU", "KRT5", "LDHB",
"LETMD1", "LYRM5", "MARS", "MMAB", "MRPL42", "MRPL51", "MRPS35",
"MTERFD3", "NDUFA12", "NDUFA9", "PHB2", "PPTC7", "RAB35", "SDSL",
"SHMT2", "SLC25A3", "SPRYD4", "SUOX", "TMTC1", "TRIAP1", "TSFM",
"TXNRD1", "UNG", "VAMP1", "YARS2", "CARKD", "CARS2", "CLYBL",
"DNAJC15", "EFHA1", "hCG_1790806", "MIPEP", "MRP63", "MRPS31",
"MTIF3", "MTRF1", "PCCA", "SLC25A15", "SLC25A30", "SUCLA2", "ACOT2",
"ADCK1", "ALDH6A1", "ARG2", "ATP5S", "C14orf153", "C14orf156",
"C14orf159", "C14orf68", "COQ6", "COX16", "COX8C", "DHRS1", "DHRS4",
"DLST", "GALC", "GLRX5", "ISCA2", "L2HGDH", "METT11D1", "MRPL52",
"MTHFD1", "NUBPL", "OXA1L", "PCK2", "PTGR2", "SLC25A21", "SLC25A29",
"SPTLC2", "TIMM9", "WDR22", "C15orf48", "CKMT1A", "CKMT1B", "CLPX",
"COX5A", "CYP11A1", "DUT", "ETFA", "FAM82A2", "GATM", "IDH2",
"IDH3A", "IREB2", "IVD", "LACTB", "LDHAL6B", "MRPL46", "MRPS11",
"MTFMT", "MTHFS", "NDUFAF1", "POLG", "RAB11A", "RAB8B", "SQRDL",
"ABAT", "ABCC12", "ACSF3", "ACSM1", "ACSM2A", "ACSM3", "ACSM5",
"BCKDK", "CA5A", "COQ7", "COQ9", "COX4I1", "CYB5B", "DDX28",
"DHODH", "DNAJA3", "DUS2L", "EARS2", "FAHD1", "GFER", "GOT2",
"GPT2", "HAGH", "HSDL1", "KARS", "LDHD", "LYRM1", "Magmas", "MLYCD",
"MRPL28", "MRPS34", "NDUFAB1", "NDUFB10", "NME3", "NME4", "NTHL1",
"PDP2", "PDPR", "PLA2G15", "RHOT2", "SPG7", "TK2", "TMEM186",
"TRAP1", "TUFM", "UQCRC2", "ABCA9", "ACACA", "ACLY", "ACSF2",
"AKAP1", "AKAP10", "ALDH3A2", "ATP5G1", "ATP5H", "ATPAF2", "C17orf90",
"C1QBP", "CCDC44", "COX10", "COX11", "DCAKD", "DRG2", "ELAC2",
"FASN", "FDXR", "GLOD4", "ICT1", "MRM1", "MRPL10", "MRPL12",
"MRPL27", "MRPL45", "MRPS23", "MRPS7", "NAGS", "NME1", "NME1-NME2",
"NME2", "NT5C", "NT5M", "PDK2", "PGS1", "PHB", "POLDIP2", "POLG2",
"PTRH2", "RHOT1", "RILP", "RNMTL1", "RSAD1", "SCO1", "SEPT4",
"SHMT1", "SLC25A10", "SLC25A11", "SLC25A19", "SLC25A35", "SLC25A39",
"TIMM22", "TMEM11", "TTC19", "ACAA2", "AFG3L2", "ATP5A1", "BCL2",
"C18orf19", "C18orf55", "FECH", "ME2", "NARS", "NDUFV2", "SLMO1",
"ZADH2", "ADCK4", "ATP5SL", "ATPBD3", "BAX", "BCAT2", "BCKDHA",
"CLPP", "COX6B1", "COX7A1", "CPT1C", "ECH1", "ECSIT", "ETFB",
"ETHE1", "FDX1L", "FKBP8", "FKSG24", "GCDH", "GPX4", "GTPBP3",
"LONP1", "MRPL34", "MRPL4", "MRPL54", "MRPS12", "NDUFA13", "NDUFA3",
"NDUFA7", "NDUFB7", "NDUFS7", "NUDT19", "OPA3", "POLRMT", "PRODH2",
"RAB11B", "RAB3D", "RAB4B", "RDH13", "SARS2", "SLC25A23", "SLC25A42",
"TIMM44", "TMEM143", "TOMM40", "TSHZ3", "UQCR", "UQCRFS1", "ACSS1",
"ATP5E", "C20orf142", "C20orf24", "C20orf72", "COX4I2", "CRLS1",
"CYP24A1", "FAM65C", "GTPBP5", "IDH3B", "MRPS26", "NFS1", "PAK7",
"PANK2", "ROMO1", "SLMO2", "SNPH", "TOMM34", "UQCC", "VISA",
"ATP5J", "ATP5O", "C21orf33", "C21orf57", "MRPL39", "MRPS6",
"NDUFV3", "SOD1", "ACO2", "ADSL", "AIFM3", "BCL2L13", "BID",
"C22orf32", "CECR5", "CERK", "CHCHD10", "COMT", "CPT1B", "GCAT",
"HSCB", "MCAT", "MPST", "MRPL40", "MTP18", "NDUFA6", "NIPSNAP1",
"PISD", "PRODH", "SAMM50", "SERHL2", "SLC25A1", "SLC25A18", "SMCR7L",
"TOMM22", "TRMU", "TSPO", "TST", "TXN2", "TXNRD2", "UCRC")
#Taken from HUGO "ribosomal protein" family
riboGenes = c('RPLP0','RPLP1','RPLP2','RPL3','RPL3L','RPL4','RPL5','RPL6','RPL7','RPL7A','RPL7L1','RPL8','RPL9','RPL10','RPL10A','RPL10L','RPL11','RPL12','RPL13A','RPL14','RPL15','RPL17','RPL18A','RPL19','RPL21','RPL22','RPL23','RPL23A','RPL24','RPL26','RPL26L','RPL27','RPL27A','RPL28','RPL29','RPL30','RPL31','RPL32','RPL34','RPL35','RPL35A','RPL36','RPL36A','RPL36A','RPL37','RPL37A','RPL38','RPL39','RPL39L','UBA52','RPL41','MRPL1','MRPL2','MRPL3','MRPL4','MRPL9','MRPL10','MRPL11','MRPL12','MRPL13','MRPL14','MRPL15','MRPL16','MRPL17','MRPL18','MRPL19','MRPL20','MRPL21','MRPL22','MRPL23','MRPL24','MRPL27','MRPL28','MRPL30','MRPL32','MRPL33','MRPL34','MRPL35','MRPL36','MRPL37','MRPL38','MRPL39','MRPL40','MRPL41','MRPL42','MRPL43','MRPL44','MRPL45','MRPL46','MRPL47','MRPL48','MRPL49','MRPL50','MRPL51','MRPL52','MRPL53','MRPL54','MRPL55','LACTB','MRPL57','MRPS2','MRPS5','MRPS6','MRPS7','MRPS9','MRPS10','MRPS11','MRPS12','MRPS14','MRPS15','MRPS16','MRPS17','MRPS18','MRPS18','MRPS18','MRPS21','MRPS22','MRPS23','MRPS24','MRPS25','MRPS26','MRPS27','MRPS28','DAP3','MRPS30','MRPS31','MRPS33','MRPS34','MRPS35','MRPS36','RPS2','RPS3','RPS3A','RPS4X','RPS4Y1','RPS4Y2','RPS5','RPS6','RPS7','RPS8','RPS9','RPS10','RPS11','RPS12','RPS13','RPS14','RPS15','RPS15A','RPS16','RPS17','RPS18','RPS19','RPS20','RPS21','RPS23','RPS24','RPS25','RPS26','RPS27','RPS27A','RPS27L','RPS28','RPS29','FAU')
#Taken from HUGO "ribosomal rnas"
riboRNAGenes = c('RNA5S1','RNA5S2','RNA5S3','RNA5S4','RNA5S5','RNA5S6','RNA5S7','RNA5S8','RNA5S9','RNA5S10','RNA5S11','RNA5S12','RNA5S13','RNA5S14','RNA5S15','RNA5S16','RNA5S17','RNA18S1','RNA18S2','RNA18S3','RNA18S4','RNA18S5','RNA28S1','RNA28S2','RNA28S3','RNA28S4','RNA28S5','RNA45S1','RNA45S2','RNA45S3','RNA45S4','RNA45S5','RNA5-8S1','RNA5-8S2','RNA5-8S3','RNA5-8S4','RNA5-8S5','MT-RNR1','MT-RNR2','RNR1RNA','RNR2RNA','RNR3RNA','RNR4RNA','RNR5RNA','RNA18SN1','RNA18SN2','RNA18SN3','RNA18SN4','RNA18SN5','RNA28SN1','RNA28SN2','RNA28SN3','RNA28SN4','RNA28SN5','RNA45SN1','RNA45SN2','RNA45SN3','RNA45SN4','RNA45SN5','RNA5-8SN1','RNA5-8SN2','RNA5-8SN3','RNA5-8SN4','RNA5-8SN5')
#Taken from HUGO "Heat shock genes"
hspGenes = c('HSPA1A','HSPA1B','HSPA1L','HSPA2','HSPA4','HSPA4L','HSPA5','HSPA6','HSPA7','HSPA8','HSPA9','HSPA12A','HSPA12B','HSPA13','HSPA14','HSPH1','HYOU1','DNAJA1','DNAJA2','DNAJA3','DNAJA4','DNAJB1','DNAJB2','DNAJB3','DNAJB4','DNAJB5','DNAJB6','DNAJB7','DNAJB8','DNAJB9','DNAJB11','DNAJB12','DNAJB13','DNAJB14','DNAJC1','DNAJC2','DNAJC3','DNAJC4','DNAJC5','DNAJC5B','DNAJC5G','DNAJC6','DNAJC7','DNAJC8','DNAJC9','DNAJC10','DNAJC11','DNAJC12','DNAJC13','DNAJC14','DNAJC15','DNAJC16','DNAJC17','DNAJC18','DNAJC19','HSCB','DNAJC21','DNAJC22','SEC63','DNAJC24','DNAJC25','GAK','DNAJC27','DNAJC28','SACS','DNAJC30','HSPB1','HSPB2','HSPB3','CRYAA','CRYAB','HSPB6','HSPB7','HSPB8','HSPB9','ODF1','HSPB11','HSP90AA1','HSP90AA3P','HSP90AB1','HSP90B1','TRAP1','BBS10','BBS12','TCP1','CCT2','CCT3','CCT4','CCT5','CCT6A','CCT6B','CCT7','CCT8',
'CLPB','HSPD1','HSPE1','MKKS')
#Usual 13 MT genes
mtGenes = c("MT-ND1", "MT-ND2", "MT-CO1", "MT-CO2", "MT-ATP8","MT-ATP6", "MT-CO3", "MT-ND3", "MT-ND4L", "MT-ND4", "MT-ND5","MT-ND6", "MT-CYB")
#MHC
mhcGenes = c('HLA-A','HLA-B','HLA-C','HLA-DMA','HLA-DMB','HLA-DOA','HLA-DOB','HLA-DPA1','HLA-DPA2','HLA-DPA3','HLA-DPB1','HLA-DPB2','HLA-DQA1','HLA-DQA2','HLA-DQB1','HLA-DQB2','HLA-DQB3','HLA-DRA','HLA-DRB1','HLA-DRB2','HLA-DRB3','HLA-DRB4','HLA-DRB5','HLA-DRB6','HLA-DRB7','HLA-DRB8','HLA-DRB9','HLA-E','HLA-F','HLA-G','HLA-H','HLA-J','HLA-K','HLA-L','HLA-N','HLA-P','HLA-S','HLA-T','HLA-U','HLA-V','HLA-W','HLA-X','HLA-Y','HLA-Z')
mhc1Genes = grep('HLA-[^D]',mhcGenes,value=TRUE)
mhc2Genes = grep('HLA-D',mhcGenes,value=TRUE)
#Haemoglobin genes from HUGO
hgGenes = c('HBA1','HBA2','HBB','HBD','HBE1','HBG1','HBG2','HBM','HBQ1','HBZ')
#Constant part of immunoglobulin from HUGO
igGenes = c('IGHA1','IGHA2','IGHG1','IGHG2','IGHG3','IGHG4','IGHD','IGHE','IGHM','IGLC1','IGLC2','IGLC3','IGLC4','IGLC5','IGLC6','IGLC7','IGKC')
#Antisense genes.  Defined by regex '-AS[0-9]+$'
antisenseGenes = c("TTLL10-AS1", "TP73-AS1", "ENO1-AS1", "PIK3CD-AS1", "PIK3CD-AS2", 
"MTOR-AS1", "NPPA-AS1", "TMEM51-AS1", "ARHGEF19-AS1", "UBXN10-AS1", 
"ZNF436-AS1", "TCEB3-AS1", "MATN1-AS1", "CSMD2-AS1", "ZMYM4-AS1", 
"NFYC-AS1", "SLFNL1-AS1", "SLC2A1-AS1", "SZT2-AS1", "HYI-AS1", 
"KDM4A-AS1", "MKNK1-AS1", "EFCAB14-AS1", "CYP4A22-AS1", "FOXD2-AS1", 
"TTC39A-AS1", "TXNDC12-AS1", "SSBP3-AS1", "DAB1-AS1", "NFIA-AS2", 
"FOXD3-AS1", "ROR1-AS1", "GNG12-AS1", "DEPDC1-AS1", "ZRANB2-AS1", 
"ERICH3-AS1", "NEXN-AS1", "PKN2-AS1", "DPYD-AS1", "DPYD-AS2", 
"RTCA-AS1", "VAV3-AS1", "KCNC4-AS1", "FAM212B-AS1", "KCND3-AS1", 
"SLC16A1-AS1", "AP4B1-AS1", "HIPK1-AS1", "SRGAP2-AS1", "FLG-AS1", 
"UBE2Q1-AS1", "RUSC1-AS1", "ASH1L-AS1", "CADM3-AS1", "GAS5-AS1", 
"RASAL2-AS1", "KIAA1614-AS1", "IPO9-AS1", "LEMD1-AS1", "SERTAD4-AS1", 
"FLVCR1-AS1", "PROX1-AS1", "SPATA17-AS1", "TGFB2-AS1", "LYPLAL1-AS1", 
"HLX-AS1", "TAF1A-AS1", "ITPKB-AS1", "SNAP47-AS1", "IBA57-AS1", 
"LYST-AS1", "LGALS8-AS1", "CHRM3-AS2", "CHRM3-AS1", "HNRNPU-AS1", 
"GCSAML-AS1", "MYT1L-AS1", "TRAPPC12-AS1", "RNASEH1-AS1", "ID2-AS1", 
"DNAJC27-AS1", "AGBL5-AS1", "GTF3C2-AS1", "BIRC6-AS1", "BIRC6-AS2", 
"RMDN2-AS1", "CYP1B1-AS1", "SLC8A1-AS1", "SIX3-AS1", "MEIS1-AS3", 
"MEIS1-AS2", "ATP6V1B1-AS1", "DGUOK-AS1", "BOLA3-AS1", "DCTN1-AS1", 
"LBX2-AS1", "ST3GAL5-AS1", "RGPD4-AS1", "GCC2-AS1", "LIMS1-AS1", 
"SH3RF3-AS1", "PGM5P4-AS1", "DPP10-AS3", "DPP10-AS2", "DPP10-AS1", 
"STEAP3-AS1", "NIFK-AS1", "CCNT2-AS1", "DARS-AS1", "ZEB2-AS1", 
"CCDC148-AS1", "XIRP2-AS1", "DLX2-AS1", "RAPGEF4-AS1", "MLK7-AS1", 
"HOXD-AS2", "TTN-AS1", "FSIP2-AS1", "OSGEPL1-AS1", "SATB2-AS1", 
"CFLAR-AS1", "IDH1-AS1", "LANCL1-AS1", "DIRC3-AS1", "CATIP-AS2", 
"SPATA3-AS1", "IQCA1-AS1", "CAPN10-AS1", "BOK-AS1", "CHL1-AS1", 
"CNTN4-AS2", "CNTN4-AS1", "ITPR1-AS1", "BHLHE40-AS1", "GRM7-AS3", 
"GRM7-AS2", "GRM7-AS1", "LMCD1-AS1", "SRGAP3-AS1", "SRGAP3-AS2", 
"SRGAP3-AS3", "SRGAP3-AS4", "THUMPD3-AS1", "PRRT3-AS1", "SLC6A1-AS1", 
"HDAC11-AS1", "FGD5-AS1", "SH3BP5-AS1", "EAF1-AS1", "PLCL2-AS1", 
"ZNF385D-AS1", "ZNF385D-AS2", "UBE2E2-AS1", "UBE2E1-AS1", "RBMS3-AS3", 
"RBMS3-AS2", "RBMS3-AS1", "OSBPL10-AS1", "CNOT10-AS1", "ARPP21-AS1", 
"ACVR2B-AS1", "ENTPD3-AS1", "VIPR1-AS1", "HHATL-AS1", "CCDC13-AS1", 
"KRBOX1-AS1", "SNRK-AS1", "ZNF197-AS1", "LARS2-AS1", "LIMD1-AS1", 
"LRRC2-AS1", "KIF9-AS1", "CELSR3-AS1", "PRKAR2A-AS1", "BSN-AS2", 
"BSN-AS1", "RBM5-AS1", "SEMA3F-AS1", "SEMA3B-AS1", "RASSF1-AS1", 
"ZMYND10-AS1", "IQCF5-AS1", "ITIH4-AS1", "CACNA2D3-AS1", "WNT5A-AS1", 
"ARHGEF3-AS1", "ARF4-AS1", "DENND6A-AS1", "FLNB-AS1", "FAM3D-AS1", 
"C3orf67-AS1", "SYNPR-AS1", "THOC7-AS1", "PSMD6-AS2", "PRICKLE2-AS1", 
"PRICKLE2-AS1", "PRICKLE2-AS2", "PRICKLE2-AS3", "ADAMTS9-AS1", 
"ADAMTS9-AS2", "MAGI1-AS1", "SUCLG2-AS1", "FOXP1-AS1", "PDZRN3-AS1", 
"CADM2-AS2", "CADM2-AS1", "ST3GAL6-AS1", "ZBTB11-AS1", "MORC1-AS1", 
"PVRL3-AS1", "PLCXD2-AS1", "CFAP44-AS1", "SIDT1-AS1", "ZBTB20-AS1", 
"ZBTB20-AS2", "ZBTB20-AS3", "ZBTB20-AS4", "LSAMP-AS1", "IGSF11-AS1", 
"B4GALT4-AS1", "ARHGAP31-AS1", "MYLK-AS1", "MYLK-AS2", "ITGB5-AS1", 
"ALDH1L1-AS1", "ALDH1L1-AS2", "CCDC37-AS1", "RUVBL1-AS1", "DNAJB8-AS1", 
"GATA2-AS1", "H1FX-AS1", "TMCC1-AS1", "NPHP3-AS1", "TMEM108-AS1", 
"BFSP2-AS1", "NCK1-AS1", "IL20RB-AS1", "CLSTN2-AS1", "ATP1B3-AS1", 
"PLS1-AS1", "PAQR9-AS1", "SLC9A9-AS1", "SLC9A9-AS2", "PLSCR5-AS1", 
"ZIC4-AS1", "HLTF-AS1", "TM4SF1-AS1", "WWTR1-AS1", "ERICH6-AS1", 
"SIAH2-AS1", "CLRN1-AS1", "AADACL2-AS1", "MBNL1-AS1", "MME-AS1", 
"PLCH1-AS1", "PLCH1-AS2", "KCNAB1-AS2", "KCNAB1-AS1", "TIPARP-AS1", 
"IQCJ-SCHIP1-AS1", "IL12A-AS1", "SEC62-AS1", "TMEM212-AS1", "NLGN1-AS1", 
"NAALADL2-AS2", "NAALADL2-AS1", "TBL1XR1-AS1", "KCNMB2-AS1", 
"PEX5L-AS1", "PEX5L-AS2", "CCDC39-AS1", "MCCC1-AS1", "KLHL6-AS1", 
"YEATS2-AS1", "ABCC5-AS1", "HTR3E-AS1", "EIF2B5-AS1", "EHHADH-AS1", 
"IGF2BP2-AS1", "ETV5-AS1", "ADIPOQ-AS1", "LPP-AS2", "LPP-AS1", 
"TPRG1-AS1", "TPRG1-AS2", "P3H2-AS1", "OSTN-AS1", "FGF12-AS1", 
"FGF12-AS2", "FGF12-AS3", "ATP13A5-AS1", "ATP13A4-AS1", "OPA1-AS1", 
"TMEM44-AS1", "XXYLT1-AS1", "XXYLT1-AS2", "TNK2-AS1", "TM4SF19-AS1", 
"UBXN7-AS1", "NCBP2-AS1", "NCBP2-AS2", "MFI2-AS1", "DLG1-AS1", 
"LMLN-AS1", "CTBP1-AS2", "NOP14-AS1", "STX18-AS1", "AFAP1-AS1", 
"CPEB2-AS1", "TAPT1-AS1", "SEPSECS-AS1", "UGDH-AS1", "UCHL1-AS1", 
"OCIAD1-AS1", "USP46-AS1", "LNX1-AS1", "IGFBP7-AS1", "ADGRL3-AS1", 
"EPHA5-AS1", "UBA6-AS1", "THAP9-AS1", "WDFY3-AS1", "FAM13A-AS1", 
"SNCA-AS1", "BMPR1B-AS1", "STPG2-AS1", "TET2-AS1", "RPL34-AS1", 
"SEC24B-AS1", "IL21-AS1", "SCOC-AS1", "SMARCA5-AS1", "HHIP-AS1", 
"SMAD1-AS2", "SMAD1-AS1", "HAND2-AS1", "WWC2-AS2", "WWC2-AS1", 
"F11-AS1", "EXOC3-AS1", "ROPN1L-AS1", "ANKRD33B-AS1", "NADK2-AS1", 
"GDNF-AS1", "EGFLAM-AS4", "EGFLAM-AS3", "EGFLAM-AS2", "EGFLAM-AS1", 
"LIFR-AS1", "OSMR-AS1", "NNT-AS1", "FGF10-AS1", "MAST4-AS1", 
"FOXD1-AS1", "ZBED3-AS1", "SCAMP1-AS1", "RASGRF2-AS1", "CKMT2-AS1", 
"ATG10-AS1", "VCAN-AS1", "TMEM161B-AS1", "MEF2C-AS1", "ARRDC3-AS1", 
"NR2F1-AS1", "RGMB-AS1", "STARD4-AS1", "NREP-AS1", "EPB41L4A-AS1", 
"EPB41L4A-AS2", "SEMA6A-AS1", "ADAMTS19-AS1", "P4HA2-AS1", "C5orf66-AS1", 
"C5orf66-AS2", "SMAD5-AS1", "TRPC7-AS1", "TRPC7-AS2", "ARHGAP26-AS1", 
"JAKMIP2-AS1", "GRPEL2-AS1", "SAP30L-AS1", "HMMR-AS1", "PRR7-AS1", 
"GMDS-AS1", "LY86-AS1", "TFAP2A-AS1", "ELOVL2-AS1", "GFOD1-AS1", 
"JARID2-AS1", "ZSCAN16-AS1", "ZNRD1-AS1", "TRIM31-AS1", "MDC1-AS1", 
"DDR1-AS1", "DDX39B-AS1", "C6orf47-AS1", "SAPCD1-AS1", "EHMT2-AS1", 
"C2-AS1", "C4A-AS1", "C4B-AS1", "HLA-DQB1-AS1", "PSMB8-AS1", 
"BTBD9-AS1", "FOXP4-AS1", "TRAM2-AS1", "MLIP-AS1", "KCNQ5-AS1", 
"OOEP-AS1", "HMGN3-AS1", "TBX18-AS1", "MANEA-AS1", "UFL1-AS1", 
"MCHR2-AS1", "BVES-AS1", "OSTM1-AS1", "ARMC2-AS1", "TRAF3IP2-AS1", 
"RNF217-AS1", "NCOA7-AS1", "MYB-AS1", "PHACTR2-AS1", "STXBP5-AS1", 
"UST-AS1", "RAET1E-AS1", "RAET1E-AS1", "SYNE1-AS1", "EZR-AS1", 
"PACRG-AS2", "PACRG-AS3", "PACRG-AS1", "RPS6KA2-AS1", "MLLT4-AS1", 
"KIF25-AS1", "PSMG3-AS1", "ELFN1-AS1", "MEOX2-AS1", "ISPD-AS1", 
"MACC1-AS1", "KLHL7-AS1", "HOXA-AS2", "HOXA-AS3", "JAZF1-AS1", 
"SEPT7-AS1", "ELMO1-AS1", "TRG-AS1", "POU6F2-AS2", "POU6F2-AS1", 
"INHBA-AS1", "DDC-AS1", "EGFR-AS1", "ABHD11-AS1", "MAGI2-AS1", 
"MAGI2-AS2", "CYP51A1-AS1", "DLX6-AS1", "PCOLCE-AS1", "KMT2E-AS1", 
"SLC26A4-AS1", "DOCK4-AS1", "ST7-AS1", "ST7-AS2", "FEZF1-AS1", 
"POT1-AS1", "JHDM1D-AS1", "NDUFB2-AS1", "WEE2-AS1", "EPHA1-AS1", 
"OR2A1-AS1", "ATP6V0E2-AS1", "PRKAG2-AS1", "PAXIP1-AS2", "PAXIP1-AS1", 
"HTR5A-AS1", "MNX1-AS2", "MNX1-AS1", "ASMTL-AS1", "GYG2-AS1", 
"ARSD-AS1", "PRKX-AS1", "WWC3-AS1", "FRMPD4-AS1", "TLR8-AS1", 
"NHS-AS1", "PPEF1-AS1", "PHKA2-AS1", "EIF1AX-AS1", "PHEX-AS1", 
"ZFX-AS1", "PCYT1B-AS1", "GK-AS1", "TAB3-AS1", "TAB3-AS2", "DMD-AS3", 
"MID1IP1-AS1", "CASK-AS1", "NDP-AS1", "ZNF674-AS1", "UXT-AS1", 
"ZNF630-AS1", "SYP-AS1", "USP27X-AS1", "SPIN4-AS1", "IGBP1-AS2", 
"IGBP1-AS1", "DLG3-AS1", "PHKA1-AS1", "PABPC1L2B-AS1", "PABPC1L2B-AS1", 
"PABPC5-AS1", "DIAPH2-AS1", "ARMCX3-AS1", "TCEAL3-AS1", "MORF4L2-AS1", 
"FRMPD3-AS1", "ALG13-AS1", "PLS3-AS1", "RHOXF1-AS1", "XIAP-AS1", 
"RAP2C-AS1", "HS6ST2-AS1", "DDX26B-AS1", "FGF13-AS1", "MAGEA8-AS1", 
"GPR50-AS1", "HCFC1-AS1", "TMLHE-AS1", "ERICH1-AS1", "DLGAP2-AS1", 
"MCPH1-AS1", "LZTS1-AS1", "EXTL3-AS1", "RBPMS-AS1", "YTHDF3-AS1", 
"RRS1-AS1", "C8orf34-AS1", "LACTB2-AS1", "MSC-AS1", "RDH10-AS1", 
"STAU2-AS1", "ZFHX4-AS1", "PKIA-AS1", "CA3-AS1", "OTUD6B-AS1", 
"C8orf37-AS1", "UBR5-AS1", "AZIN1-AS1", "BAALC-AS2", "BAALC-AS1", 
"RAD21-AS1", "SAMD12-AS1", "HAS2-AS1", "FAM83A-AS1", "FER1L6-AS1", 
"FER1L6-AS2", "RNF139-AS1", "KIAA0196-AS1", "ZFAT-AS1", "RHPN1-AS1", 
"MAFA-AS1", "FAM83H-AS1", "TONSL-AS1", "ZNF252P-AS1", "PGM5P3-AS1", 
"VLDLR-AS1", "RFX3-AS1", "GLIS3-AS1", "CDC37L1-AS1", "PTPRD-AS1", 
"PTPRD-AS2", "LURAP1L-AS1", "FOCAD-AS1", "CDKN2A-AS1", "CDKN2B-AS1", 
"IFT74-AS1", "TOPORS-AS1", "B4GALT1-AS1", "UBE2R2-AS1", "CNTFR-AS1", 
"DNAJB5-AS1", "PGM5-AS1", "PGM5-AS1", "C9orf135-AS1", "MAMDC2-AS1", 
"SMC5-AS1", "RORB-AS1", "C9orf41-AS1", "VPS13A-AS1", "GNA14-AS1", 
"BARX1-AS1", "STX17-AS1", "TMEM246-AS1", "SMC2-AS1", "PAPPA-AS2", 
"PAPPA-AS1", "ASTN2-AS1", "PSMD5-AS1", "GSN-AS1", "SLC25A25-AS1", 
"PTGES2-AS1", "ODF2-AS1", "PRRX2-AS1", "DBH-AS1", "COL5A1-AS1", 
"PPP1R26-AS1", "CCDC183-AS1", "MAN1B1-AS1", "C9orf173-AS1", "ARRDC1-AS1", 
"TOLLIP-AS1", "KRTAP5-AS1", "MRPL23-AS1", "CD81-AS1", "KCNQ1-AS1", 
"CARS-AS1", "MRGPRG-AS1", "RRM1-AS1", "TMEM9B-AS1", "SBF2-AS1", 
"MRVI1-AS1", "ZBED5-AS1", "SPTY2D1-AS1", "NAV2-AS5", "NAV2-AS4", 
"NAV2-AS3", "NAV2-AS2", "NAV2-AS1", "BBOX1-AS1", "CSTF3-AS1", 
"FBXO3-AS1", "ALKBH3-AS1", "LRP4-AS1", "PTPRJ-AS1", "OVOL1-AS1", 
"MRGPRF-AS1", "ANO1-AS2", "ANO1-AS1", "SHANK2-AS1", "SHANK2-AS2", 
"SHANK2-AS3", "ARAP1-AS1", "ARAP1-AS2", "KCTD21-AS1", "RAB30-AS1", 
"PCF11-AS1", "GRM5-AS1", "JRKL-AS1", "NCAM1-AS1", "TMPRSS4-AS1", 
"USP2-AS1", "PKNOX2-AS1", "STT3A-AS1", "ST3GAL4-AS1", "KIRREL3-AS1", 
"KIRREL3-AS2", "KIRREL3-AS3", "NTM-AS1", "IDI2-AS1", "ADARB2-AS1", 
"PITRM1-AS1", "CALML3-AS1", "GATA3-AS1", "CELF2-AS2", "CELF2-AS1", 
"PROSER2-AS1", "VIM-AS1", "ST8SIA6-AS1", "STAM-AS1", "SLC39A12-AS1", 
"NEBL-AS1", "GPR158-AS1", "MKX-AS1", "WAC-AS1", "SVIL-AS1", "ZEB1-AS1", 
"PARD3-AS1", "ZNF32-AS3", "ZNF32-AS1", "ZNF32-AS2", "TMEM72-AS1", 
"FAM170B-AS1", "C10orf71-AS1", "SGMS1-AS1", "PRKG1-AS1", "TMEM26-AS1", 
"JMJD1C-AS1", "UNC5B-AS1", "CDH23-AS1", "DNAJC9-AS1", "ZSWIM8-AS1", 
"ZNF503-AS1", "ZNF503-AS2", "KCNMA1-AS1", "KCNMA1-AS2", "KCNMA1-AS3", 
"ZMIZ1-AS1", "NUTM2B-AS1", "SFTPD-AS1", "TMEM254-AS1", "GRID1-AS1", 
"NUTM2A-AS1", "ACTA2-AS1", "SLC16A12-AS1", "TNKS2-AS1", "PLCE1-AS2", 
"PLCE1-AS1", "ENTPD1-AS1", "SLIT1-AS1", "DNMBP-AS1", "KCNIP2-AS1", 
"CYP17A1-AS1", "NEURL1-AS1", "SH3PXD2A-AS1", "CFAP58-AS1", "SORCS3-AS1", 
"PDCD4-AS1", "WDR11-AS1", "ATE1-AS1", "FAM53B-AS1", "TEX36-AS1", 
"EDRF1-AS1", "FANK1-AS1", "TCERG1L-AS1", "ADGRA1-AS1", "CACNA1C-AS4", 
"CACNA1C-AS3", "CACNA1C-AS2", "CACNA1C-AS1", "CCND2-AS1", "CCND2-AS2", 
"C1RL-AS1", "A2ML1-AS1", "A2ML1-AS2", "A2M-AS1", "PLBD1-AS1", 
"RERG-AS1", "RASSF8-AS1", "ARNTL2-AS1", "OVCH1-AS1", "DDX11-AS1", 
"DENND5B-AS1", "BCDIN3D-AS1", "KRT73-AS1", "HOXC-AS1", "AGAP2-AS1", 
"TMEM5-AS1", "LLPH-AS1", "IFNG-AS1", "CEP83-AS1", "TMPO-AS1", 
"USP30-AS1", "FAM222A-AS1", "MAPKAPK5-AS1", "LHX5-AS1", "TBX5-AS1", 
"TESC-AS1", "PXN-AS1", "CLIP1-AS1", "LATS2-AS1", "SACS-AS1", 
"C1QTNF9B-AS1", "SPATA13-AS1", "C1QTNF9-AS1", "C1QTNF9B-AS1", 
"WASF3-AS1", "USP12-AS1", "USP12-AS2", "PDX1-AS1", "PAN3-AS1", 
"MTUS2-AS2", "MTUS2-AS1", "TEX26-AS1", "FRY-AS1", "SPG20-AS1", 
"FREM2-AS1", "VWA8-AS1", "ENOX1-AS2", "ENOX1-AS1", "SMIM2-AS1", 
"TSC22D1-AS1", "TPT1-AS1", "SLC25A30-AS1", "CPB2-AS1", "HTR2A-AS1", 
"SUCLA2-AS1", "MED4-AS1", "DLEU1-AS1", "DLEU7-AS1", "RNASEH2B-AS1", 
"INTS6-AS1", "DIAPH3-AS1", "DIAPH3-AS2", "PCDH9-AS1", "PCDH9-AS2", 
"PCDH9-AS3", "PCDH9-AS4", "LMO7-AS1", "MYCBP2-AS1", "SCEL-AS1", 
"EDNRB-AS1", "RNF219-AS1", "RBM26-AS1", "NDFIP2-AS1", "GPC5-AS2", 
"GPC5-AS1", "GPC6-AS2", "GPC6-AS1", "SOX21-AS1", "CLDN10-AS1", 
"DNAJC3-AS1", "FARP1-AS1", "STK24-AS1", "DOCK9-AS1", "DOCK9-AS2", 
"UBAC2-AS1", "CLYBL-AS2", "CLYBL-AS1", "PCCA-AS1", "NALCN-AS1", 
"FGF14-AS1", "FGF14-AS2", "DAOA-AS1", "MYO16-AS2", "MYO16-AS1", 
"COL4A2-AS2", "COL4A2-AS1", "ARHGEF7-AS2", "ARHGEF7-AS1", "ATP11A-AS1", 
"MCF2L-AS1", "F10-AS1", "GRTP1-AS1", "GAS6-AS1", "GAS6-AS2", 
"PRMT5-AS1", "DHRS4-AS1", "NOVA1-AS1", "FOXG1-AS1", "ARHGAP5-AS1", 
"EGLN3-AS1", "NKX2-1-AS1", "SLC25A21-AS1", "FRMD6-AS2", "FRMD6-AS1", 
"KTN1-AS1", "OTX2-AS1", "PSMA3-AS1", "HIF1A-AS1", "HIF1A-AS2", 
"FUT8-AS1", "ACTN1-AS1", "DIO2-AS1", "FOXN3-AS1", "FOXN3-AS2", 
"ITPK1-AS1", "FAM181A-AS1", "DICER1-AS1", "MKRN3-AS1", "GABRG3-AS1", 
"DPH6-AS1", "SRP14-AS1", "PLCB2-AS1", "EHD4-AS1", "PLA2G4E-AS1", 
"EIF3J-AS1", "GATM-AS1", "GABPB1-AS1", "RORA-AS1", "RORA-AS2", 
"USP3-AS1", "IQCH-AS1", "THSD4-AS1", "HEXA-AS1", "ADPGK-AS1", 
"C15orf59-AS1", "LOXL1-AS1", "UBL7-AS1", "LINGO1-AS1", "LINGO1-AS2", 
"ANKRD34C-AS1", "ST20-AS1", "TMC3-AS1", "CPEB1-AS1", "AGBL1-AS1", 
"NTRK3-AS1", "CRTC3-AS1", "PRC1-AS1", "NR2F2-AS1", "SPATA8-AS1", 
"CERS3-AS1", "PCSK6-AS1", "LMF1-AS1", "SRRM2-AS1", "MMP25-AS1", 
"ZNF213-AS1", "GLIS2-AS1", "CRYM-AS1", "LCMT1-AS1", "LCMT1-AS2", 
"IL21R-AS1", "ATP2A1-AS1", "CDIPT-AS1", "FBXL19-AS1", "PYCARD-AS1", 
"ITFG1-AS1", "BEAN1-AS1", "VAC14-AS1", "TAT-AS1", "FOXC2-AS1", 
"SNAI3-AS1", "VPS9D1-AS1", "GAS8-AS1", "PITPNA-AS1", "ALOX12-AS1", 
"TMEM220-AS1", "MPRIP-AS1", "RAI1-AS1", "EPN2-AS1", "CCDC144NL-AS1", 
"ABHD15-AS1", "RARA-AS1", "RAMP2-AS1", "ASB16-AS1", "RUNDC3A-AS1", 
"MAP3K14-AS1", "MAPT-AS1", "KANSL1-AS1", "SP2-AS1", "HOXB-AS1", 
"HOXB-AS3", "HOXB-AS2", "HOXB-AS4", "TMEM92-AS1", "CACNA1G-AS1", 
"BZRAP1-AS1", "SEPT4-AS1", "TBX2-AS1", "PRR29-AS1", "PRKCA-AS1", 
"ABCA9-AS1", "KCNJ2-AS1", "HID1-AS1", "C1QTNF1-AS1", "BAIAP2-AS1", 
"AATK-AS1", "MAFG-AS1", "DLGAP1-AS1", "DLGAP1-AS2", "DLGAP1-AS3", 
"DLGAP1-AS4", "DLGAP1-AS5", "L3MBTL4-AS1", "NDUFV2-AS1", "PPP4R1-AS1", 
"LDLRAD4-AS1", "GATA6-AS1", "TTC39C-AS1", "AQP4-AS1", "DSG1-AS1", 
"DSG2-AS1", "SLC14A2-AS1", "TCF4-AS1", "TCF4-AS2", "PARD6G-AS1", 
"NRSN2-AS1", "SDCBP2-AS1", "SIRPG-AS1", "UBOX5-AS1", "MCM8-AS1", 
"LAMP5-AS1", "SNAP25-AS1", "ISM1-AS1", "MACROD2-AS1", "KIZ-AS1", 
"NKX2-2-AS1", "ZNF337-AS1", "HM13-AS1", "ZNF341-AS1", "RALY-AS1", 
"ITCH-AS1", "MMP24-AS1", "FAM83C-AS1", "DLGAP4-AS1", "PLCG1-AS1", 
"OSER1-AS1", "HNF4A-AS1", "KCNK15-AS1", "STK4-AS1", "CSE1L-AS1", 
"CEBPB-AS1", "ADNP-AS1", "BMP7-AS1", "GNAS-AS1", "LAMA5-AS1", 
"C20orf166-AS1", "SLCO4A1-AS1", "OGFR-AS1", "ZBTB46-AS1", "UCKL1-AS1", 
"CIRBP-AS1", "CSNK1G2-AS1", "CACTIN-AS1", "DPP9-AS1", "RAB11B-AS1", 
"ILF3-AS1", "FARSA-AS1", "CEBPA-AS1", "ZNF30-AS1", "HPN-AS1", 
"TMEM147-AS1", "UPK1A-AS1", "ZNF529-AS1", "ZNF790-AS1", "ZNF793-AS1", 
"ZNF571-AS1", "LIPE-AS1", "DACT3-AS1", "NAPA-AS1", "GLTSCR1-AS1", 
"GLTSCR2-AS1", "PLA2G4C-AS1", "CARD8-AS1", "NUCB1-AS1", "PTOV1-AS1", 
"PTOV1-AS2", "ZNF649-AS1", "ZNF350-AS1", "LENG8-AS1", "ZNF582-AS1", 
"ZNF667-AS1", "ZIM2-AS1", "A1BG-AS1", "MZF1-AS1", "ZFY-AS1", 
"NLGN4Y-AS1", "POTEH-AS1", "CECR5-AS1", "THAP7-AS1", "MIF-AS1", 
"ADORA2A-AS1", "ZNRF3-AS1", "HORMAD2-AS1", "MORC2-AS1", "RNF185-AS1", 
"PIK3IP1-AS1", "ISX-AS1", "APOBEC3B-AS1", "MGAT3-AS1", "EP300-AS1", 
"EFCAB6-AS1", "NUP50-AS1", "PRR34-AS1", "GTSE1-AS1", "CHKB-AS1", 
"SAMSN1-AS1", "CHODL-AS1", "CYYR1-AS1", "BACH1-AS1", "GRIK1-AS1", 
"HUNK-AS1", "MIS18A-AS1", "URB1-AS1", "PAXBP1-AS1", "C21orf62-AS1", 
"IL10RB-AS1", "TTC3-AS1", "BRWD1-AS2", "BRWD1-AS1", "B3GALT5-AS1", 
"DSCAM-AS1", "ZNF295-AS1", "LRRC3-AS1", "TSPEAR-AS1", "TSPEAR-AS2", 
"ITGB2-AS1", "COL18A1-AS2", "COL18A1-AS1", "FTCD-AS1", "MCM3AP-AS1"
)
```
