## supplementary scripts for "Tumor to normal single cell mRNA comparisons reveal a pan-neuroblastoma cancer cell": phaseMutations.R

```
#############
# Libraries #
#############

library(GenomicAlignments)
#library(Rsamtools)
library(VariantAnnotation)
#library(refGenome)
library(Matrix)
#For sample matching
library(gdsfmt)
library(SNPRelate)

###########
# Example #
###########

ref = "~/canpipe/ref/human/1000Genomes_hs37d5/genome.fa"
ref10X= "~/scratch/Common/refdata-cellranger-GRCh38-1.2.0/fasta/genome.fa"
gtf='~/trueHome/scratch/Common/refdata-cellranger-GRCh38-1.2.0/genes/genes.gtf'
gtf='~/trueHome/scratch/Common/refdata-cellranger-hg19-1.2.0/genes/genes.gtf'
#Wilms2
childBAM_Normal = "/nfs/users/nfs_m/my4/PipeLive/1711/PD37276g/PD37276g.sample.dupmarked.bam"
childBAM_Tumour = "/nfs/users/nfs_m/my4/PipeLive/1711/PD37276a/PD37276a.sample.dupmarked.bam"
mumBAM = "/nfs/users/nfs_m/my4/PipeLive/1711/PD37286b/PD37286b.sample.dupmarked.bam"
dadBAM = "/nfs/users/nfs_m/my4/PipeLive/1711/PD37287b/PD37287b.sample.dupmarked.bam"
mutations = '~/PipeLive/1711/PD37276a/PD37276a.caveman_c.flag.vcf.gz'
outDir = '~/scratch/Imprinting/Wilms2Test'
#Wilms3
childBAM_Normal = "/nfs/users/nfs_m/my4/PipeLive/1711/PD37272c/PD37272c.sample.dupmarked.bam"
childBAM_Tumour = "/nfs/users/nfs_m/my4/PipeLive/1711/PD37272a/PD37272a.sample.dupmarked.bam"
mumBAM = "/nfs/users/nfs_m/my4/PipeLive/1711/PD37288b/PD37288b.sample.dupmarked.bam"
dadBAM = "/nfs/users/nfs_m/my4/PipeLive/1711/PD37289b/PD37289b.sample.dupmarked.bam"
mutations = '~/PipeLive/1711/PD37272a/PD37272a.caveman_c.flag.vcf.gz'
outDir = '~/scratch/Imprinting/Wilms3Test'
###########
# Not run
#hSNPs = gt$informativeSNPsFromBAMs(childBAM_Normal,ref,mumBAM,dadBAM,outDir,nParallel=12)
#muts = gt$phaseMutations(childBAM_Tumour,hSNPs,mutations)
#scDat = gt$phaseExpression(hSNPs,unlist(scRNA),ref,gtf,labels=names(unlist(scRNA)),outDir=outDir,chain=NULL,nParallel=8)

#############
# Functions #
#############

#' Finds all phased heterozygous SNPs from family DNA
#'
#' Given BAM files from a child and at least one parent, this function finds all heterozygous SNPs and then uses the parents to phase those that can be phased with the maternal/paternal allele.
#'
#' @param childBAM_Normal BAM derived from normal (not tumour) DNA from the child.
#' @param ref Path to reference genome fasta file.
#' @param mumBAM Path to BAM derived from the mother's DNA.  One of mumBAM and dadBAM must be given.
#' @param dadBAM Path to BAM derived from the father's DNA.  One of mumBAM and dadBAM must be given.
#' @param outDir Directory where intermediate output files (VCF and allele counts) are stored.
#' @param skipIfExists If any of the intermediate files exist, load them rather than running the generation step again.
#' @param nParallel How many processors to use.
#' @param ... Extra parameters passed to findHetSNPs and findInformativeSNPs
#' @return A GRanges object giving the coordinates of the heterozygous SNPs in the child and information about which of these can be phased.
informativeSNPsFromBAMs = function(childBAM_Normal,ref,mumBAM=NULL,dadBAM=NULL,outDir=getwd(),skipIfExists=TRUE,nParallel=2,...){
  mum = !is.null(mumBAM)
  dad = !is.null(dadBAM)
  if(!mum & !dad)
    stop("At least one parent must be given")
  #####################################################
  # Find all Heterozygous SNPs in the child's normal.
  message("Calling heterozygous SNPs in child...")
  hSNPs = findHetSNPs(childBAM_Normal,ref,VCF_path=file.path(outDir,sprintf("%s_hetSNPs.vcf.gz",basename(childBAM_Normal))),skipIfExists=skipIfExists,nParallel=nParallel,...)
  ##################################
  # Run allelecounter if necessary
  message("Counting alleles at SNPs in parents...")
  labs=c()
  if(mum)
    labs=c(labs,basename(mumBAM))
  if(dad)
    labs=c(labs,basename(dadBAM))
  outputs = file.path(outDir,sprintf('%s_%s_alleleCounts.tsv',c('Mum','Dad')[c(mum,dad)],labs[c(mum,dad)]))
  #Do we really have to run anything?
  if(!skipIfExists | any(!file.exists(outputs))){
    #Yes we do, so create the loci file
    tgtLoci = file.path(outDir,sprintf("%s_hetSNPs.tsv",basename(childBAM_Normal)))
    write.table(cbind(as.character(seqnames(hSNPs)),start(hSNPs)),tgtLoci,row.names=FALSE,sep='\t',col.names=FALSE,quote=FALSE)
    #Now construct the inputs
    bams=c()
    if(mum)
      bams = c(bams,mumBAM)
    if(dad)
      bams = c(bams,dadBAM)
    #Now run allele counter with things sensible for bulk DNA.  
    parallelAlleleCounter(tgtLoci,outputs,bams,x=FALSE,f=3,F=3852,m=20,q=35,r=ref,skipIfExists=skipIfExists,d=TRUE,nParallel=nParallel)
  }else{
    message("Previous results found, skipping re-calculation")
  }
  #########################
  # Find informative SNPs
  message("Identifying informative SNPs...")
  mumCnts=NULL
  if(mum)
    mumCnts=outputs[1]
  dadCnts=NULL
  if(dad)
    dadCnts=outputs[length(outputs)]
  hSNPs = findInformativeSNPs(hSNPs,mumCnts,dadCnts,...)
  return(hSNPs)
}

#' Get 10X informative SNP counts
#'
#' Given a set of SNPs phased with parental DNA, calculates the single cell expression of the phased SNPs. Runs allele counter to do this, also optionally does liftOver from hg19 to GRCh38.
#'
#'
#' @param hSNPs GRanges object containing informative SNPs.
#' @param bams Cellranger output 10X bams to extract counts from.
#' @param ref Reference genome for allelecounter to use.
#' @param labels Each sample needs a unique name.  Give them here.
#' @param outDir A directory where allele counts and other intermediate files are stored.
#' @param chain Path to liftover file to use.  If NULL, no coordinate translation is done.
#' @param nParallel Number of processors to use.
#' @param skipIfExists If the output from allelecounter already exists, don't remake it.
#' @param matAlleleCol Name of the column in \code{hSNPs} containing the maternal allele.
#' @param patAlleleCol Name of the column in \code{hSNPs} containing the maternal allele.
#' @param postChrStatus Set to NULL leaves chromosome names alone.  If set to TRUE forces post-count chromosome names to have Chr, FALSE, forces them not to. 
#' @param ... Extra parameter passed to parallelAlleleCounter.
phaseExpression = function(hSNPs,bams,ref,gtf,labels=basename(dirname(bams)),outDir=getwd(),chain=NULL,nParallel=1,skipIfExists=TRUE,matAlleleCol='maternalAllele',patAlleleCol='paternalAllele',postChrStatus=NULL,...){
  #Translate coordinates if necessary
  if(!is.null(chain)){
    ch = rtracklayer::import.chain(chain)
    hSNPs = unlist(rtracklayer::liftOver(hSNPs,ch))
  }
  #Keep the non-phased ones, they might be useful
  #hSNPs = hSNPs[!is.na(hSNPs$altIsMum)]
  #Decide which we actually need to process
  outputs = file.path(outDir,sprintf("%s_alleleCounts.tsv",labels))
  if(skipIfExists){
    w = which(!file.exists(outputs))
  }else{
    w = seq_along(outputs)
  }
  #Are there any?
  if(length(w)>0){
    #Yes, so let's make a loci file.
    ####################################################
    # Should put routine to make loci file unique here
    tgtLoci = file.path(outDir,'hetSNPsForPhasingExpression.tsv')
    write.table(cbind(as.character(seqnames(hSNPs)),start(hSNPs)),tgtLoci,row.names=FALSE,sep='\t',col.names=FALSE,quote=FALSE)
    #Now run what needs running
    parallelAlleleCounter(tgtLoci,outputs[w],bams[w],r=ref,x=TRUE,nParallel=nParallel,skipIfExists=skipIfExists,...)
    system(sprintf('rm %s',tgtLoci))
  }else{
    message("Previous results found, skipping re-calcualtion")
  }
  #Now load the data in.
  out = list()
  for(i in seq_along(bams)){
    message(sprintf("Loading counts for %s",labels[i]))
    out[[i]] = read.table(outputs[i],sep='\t',header=TRUE,comment.char='')
    out[[i]] = with(out[[i]],GRanges(X.CHR,IRanges(POS,width=1),A=Count_A,C=Count_C,G=Count_G,T=Count_T,coverage=Good_depth,cellID=paste0(labels[i],'___',gsub('-[0-9]+$','',Barcode))))
  }
  out = do.call(c,out)
  bases = colnames(mcols(out))[seq(4)]
  #Match up with the SNPs
  m = match(out,hSNPs)
  #Create counts for ref,alt,maternal, and paternal
  tmp = as.matrix(mcols(out)[,bases])
  out$matCount = tmp[cbind(seq_along(out),match(mcols(hSNPs)[m,matAlleleCol],bases))]
  out$patCount = tmp[cbind(seq_along(out),match(mcols(hSNPs)[m,patAlleleCol],bases))]
  out$refCount = tmp[cbind(seq_along(out),match(mcols(hSNPs)[m,grep('^REF$',colnames(mcols(hSNPs)),ignore.case = TRUE)],bases))]
  out$altCount = tmp[cbind(seq_along(out),match(mcols(hSNPs)[m,grep('^ALT$',colnames(mcols(hSNPs)),ignore.case = TRUE)],bases))]
  if(!is.null(postChrStatus)){
    if(postChrStatus){
      out = renameSeqlevels(out,setNames(paste0('chr',gsub('^chr','',seqlevels(out))),seqlevels(out)))
    }else{
      out = renameSeqlevels(out,setNames(gsub('^chr','',seqlevels(out)),seqlevels(out)))
    }
  }
  ##Merge in any columns that we don't already have
  #for(newCol in colnames(mcols(hSNPs))){
  #  if(newCol %in% colnames(mcols(out)))
  #    next
  #  mcols(out)[,newCol] = mcols(hSNPs)[m,newCol]
  #}
  ##Convert ref/alt to lower case if not already
  #tmp = colnames(mcols(out))
  #tmp[tmp=='REF']='ref'
  #tmp[tmp=='ALT']='alt'
  #colnames(mcols(out)) = tmp
  ##Import which allele is mum and which is dad.  If phasing unknown, make "maternal" the major allele
  #out$maternalAllele = out$ref
  #out$maternalAllele[which(out$altIsMum)] = out$alt[which(out$altIsMum)]
  #out$paternalAllele = out$alt
  #out$paternalAllele[which(out$altIsMum)] = out$ref[which(out$altIsMum)]
  #bases=c('A','C','G','T')
  #out$refCount = as.matrix(mcols(out)[,bases])[cbind(seq_along(out),match(out$ref,bases))]
  #out$altCount = as.matrix(mcols(out)[,bases])[cbind(seq_along(out),match(out$alt,bases))]
  #out$otherCount = out$coverage-out$refCount-out$altCount
  #out$maternalCount = as.matrix(mcols(out)[,bases])[cbind(seq_along(out),match(out$maternalAllele,bases))]
  #out$paternalCount = as.matrix(mcols(out)[,bases])[cbind(seq_along(out),match(out$paternalAllele,bases))]
  ##Load the gtf
  #txdb = ensemblGenome()
  #tmp = getwd()
  #setwd(dirname(gtf))
  #read.gtf(txdb,basename(gtf))
  #setwd(tmp)
  ##Now extract the information we need
  #gns = getGeneTable(txdb)
  #gns = GRanges(gns$seqid,IRanges(gns$start,gns$end),strand=gns$strand,geneID=gns$gene_id,geneName=gns$gene_name,geneBiotype=gns$gene_biotype)
  #exns = getGtf(txdb)
  #exns = exns[exns$feature=='exon',]
  #exns = GRanges(exns$seqid,IRanges(exns$start,exns$end),strand=exns$strand)
  message("Loading reference GTF")
  ann = rtracklayer::import(gtf)
  gns = ann[ann$type=='gene']
  exns = ann[ann$type=='exon']
  #Annotate each SNP as belonging to exon,intron or nothing
  out$regionType='InterGenic'
  o =suppressWarnings(findOverlaps(out,gns,ignore.strand=TRUE))
  out$regionType[queryHits(o)]='Intronic'
  #Get geneID and name in an efficient way
  out$geneID=NA
  out$geneName=NA
  oo = data.frame(o)
  oo$geneID = gns$gene_id[oo$subjectHits]
  oo$geneName = gns$gene_name[oo$subjectHits]
  #Sort so that the duplicate ones are always in the same order
  oo = oo[order(oo$queryHits,oo$subjectHits),]
  tmp = aggregate(cbind(geneID,geneName) ~ queryHits,data=oo,FUN=c)
  out$geneID[tmp$queryHits] = tmp$geneID
  out$geneName[tmp$queryHits] = tmp$geneName
  #Get exons
  o =suppressWarnings(findOverlaps(out,exns,ignore.strand=TRUE))
  out$regionType[queryHits(o)]='Exonic'
  #Try and record cellranger pass cells
  #out$isValidCell = NA
  #cellBarcodes = list()
  #for(i in seq_along(bams)){
  #  cellFiles = list.files(file.path(dirname(bams[i]),'filtered_gene_bc_matrices'))
  #  if(length(cellFiles)==1){
  #    cellFiles = file.path(dirname(bams[i]),'filtered_gene_bc_matrices',cellFiles,'barcodes.tsv')
  #    if(file.exists(cellFiles)){
  #      cellFiles = read.table(cellFiles,sep='\t')
  #      cellBarcodes[[labels[i]]] = gsub('-[0-9]+$','',cellFiles[,1])
  #      #Call this samples cells one way or another
  #      out$isValidCell[startsWith(out$cellID,labels[i])]=FALSE
  #      out$isValidCell[out$cellID %in% paste0(labels[i],'___',cellBarcodes[[labels[i]]])] = TRUE
  #    }
  #  }
  #}
  return(out)
}


createPhasedExpressionArray = function(phasedExpression){
  #Arregate to make matricies
  dat = data.frame(phasedExpression)
  tmp = dat$geneName
  dat$geneName = NA
  dat$geneName[lengths(tmp)==1] = unlist(tmp[lengths(tmp)==1])
  if(any(lengths(tmp)>1))
    dat$geneName[lengths(tmp)>1] = sapply(tmp[lengths(tmp)>1],paste,collapse=',')
  dimNames = list(unique(dat$geneName[!is.na(dat$geneName)]),unique(dat$cellID))
  mumMat = aggregate(maternalCount ~ cellID + geneName,data=dat[!is.na(dat$geneName),],sum)
  mumMat = sparseMatrix(i=match(mumMat$geneName,dimNames[[1]]),
                        j=match(mumMat$cellID,dimNames[[2]]),
                        x=mumMat$maternalCount,
                        dim=lengths(dimNames),
                        dimnames=dimNames)
  dadMat = aggregate(paternalCount ~ cellID + geneName,data=dat,sum)
  dadMat = sparseMatrix(i=match(dadMat$geneName,dimNames[[1]]),
                        j=match(dadMat$cellID,dimNames[[2]]),
                        x=dadMat$paternalCount,
                        dim=lengths(dimNames),
                        dimnames=dimNames)
  return(list(mumMat=mumMat,dadMat=dadMat))
}

#' @param SNPs Full list of all SNPs.
#' @param gtf GTF to use to subset data.
#' @param includeIntrons Should SNPs in introns be included?
#' @param minAF Minimum population allele frequency to allow for SNPs.
#' @param chain A chain file to liftover coordinates from the GTF genome build to the SNP genome build.  If NULL, no conversion is done.
subsetSNPsForMaternalPhasing = function(SNPs,gtf,includeIntrons=TRUE,minAF=0.02,chain=NULL){
  #Load the gtf
  txdb = ensemblGenome()
  tmp = getwd()
  setwd(dirname(gtf))
  read.gtf(txdb,basename(gtf))
  setwd(tmp)
  #Now extract the information we need
  gns = getGeneTable(txdb)
  gns = GRanges(gns$seqid,IRanges(gns$start,gns$end),strand=gns$strand,geneID=gns$gene_id,geneName=gns$gene_name,geneBiotype=gns$gene_biotype)
  exns = getGtf(txdb)
  exns = exns[exns$feature=='exon',]
  exns = GRanges(exns$seqid,IRanges(exns$start,exns$end),strand=exns$strand)
  #Do liftover if required
  if(!is.null(chain)){
    ch = rtracklayer::import.chain(chain)
    gns = unlist(rtracklayer::liftOver(gns,ch))
    exns = unlist(rtracklayer::liftOver(exns,ch))
  }
  #Subset to genomic regions of interest
  if(includeIntrons){
    SNPs = subsetByOverlaps(SNPs,gns,ignore.strand=TRUE)
  }else{
    SNPs = subsetByOverlaps(SNPs,exns,ignore.strand=TRUE)
  }
  #Remove the ones that have a AF too low to be of use
  SNPs = SNPs[abs(SNPs$AF-0.5) < (0.5-minAF),]
  #Ensure we don't have a list of alts or anything ambiguous
  SNPs = SNPs[lengths(SNPs$ALT)==1,]
  SNPs$ALT = unlist(SNPs$ALT)
  return(SNPs)
}


#' Try and infer which cells are from the parent using only the child's DNA
#'
#' The aim of this function is to infer at the individual cell level if a cell is from an individual who's DNA has been sequenced, or from one of its parents.  This works by focusing on a panel of sites of common variation in the genome (i.e., SNPs) and identifying those that are homozygous in the child.  Some fraction of these sites will be heterozygous in the parent and so by comparing the number of non-matching bases in the RNA we can identify when a cell is derived from one of the parents instead of the child.
#' 
#' Usually we use something like the 1000 genomes SNP panel.  For computational efficiency it is useful to exclude any non-exonic SNPs (as they will likely not be used anyway) and possibly include only those with a population allele frequency close to 50% (as these are the sites with the highest apriori chance of being informative).
#'
#' Labels for each BAM that is allele counted are either taken from the names of the input vector or basename(BAM)
#'
#' @param childBAM The BAM file with DNA sequence for the child
#' @param RNA_BAMs The BAM files containing the 10X RNA sequencing data to infer cell information from.
#' @param SNPs A GRanges object with ALT and REF columns containing the locations of sites to interogate and AF column giving population allele frequency.
#' @param ref The reference genome to use.
#' @param gtf The gtf file used to quantify expression.
#' @param outDir Directory to store results.
#' @param nParallel How many processors to use.
#' @param ... Passed to parallelAlleleCounters and phaseExpression
inferMaternalFromSNPs = function(childBAM,RNA_BAMs,SNPs,ref,gtf,minAF=0.01,minCoverage=10,errorRate=0.2,error10X=0.05,outDir=getwd(),nParallel=1,...){
  ################
  # Input checks #
  ################
  #Label for the childBAM
  label = names(childBAM)
  if(is.null(label))
    label = basename(label)
  #Labels for the RNA
  labels = names(RNA_BAMs)
  if(is.null(labels))
    labels = basename(RNA_BAMs)
  if(any(duplicated(c(label,labels))))
    stop("Each BAM must have a unique label.")
  bases = c('A','C','G','T')
  ########################
  # Get most useful SNPs #
  ########################
  #Load the gtf
  txdb = ensemblGenome()
  tmp = getwd()
  setwd(dirname(gtf))
  read.gtf(txdb,basename(gtf))
  setwd(tmp)
  #Now extract the information we need
  #gns = getGeneTable(txdb)
  #gns = GRanges(gns$seqid,IRanges(gns$start,gns$end),strand=gns$strand,geneID=gns$gene_id,geneName=gns$gene_name,geneBiotype=gns$gene_biotype)
  exns = getGtf(txdb)
  exns = exns[exns$feature=='exon',]
  exns = GRanges(exns$seqid,IRanges(exns$start,exns$end),strand=exns$strand)
  #Keep only exonic SNPs that have an AF of at least 0.01%
  SNPs = subsetByOverlaps(SNPs[abs(SNPs$AF-0.5)<(0.5-minAF)],exns,ignore.strand=TRUE)
  #Ensure we don't have a list of alts or anything ambiguous
  SNPs = SNPs[lengths(SNPs$ALT)==1,]
  SNPs$ALT = unlist(SNPs$ALT)
  ##################################
  # Determine which are homozygous #
  ##################################
  #Make the target file
  tgtLoci = file.path(outDir,'lociFile.tsv')
  write.table(cbind(as.character(seqnames(SNPs)),start(SNPs)),tgtLoci,row.names=FALSE,sep='\t',col.names=FALSE,quote=FALSE)
  #Run allele counter
  output=file.path(outDir,sprintf('%s.SNPs_alleleCounts.tsv',label))
  parallelAlleleCounter(tgtLoci,output,childBAM,nParallel=nParallel,f=3,F=3852,m=20,q=35,r=ref,d=TRUE,x=FALSE,...)
  out = read.table(output,sep='\t',comment.char='',header=TRUE)
  out = with(out,GRanges(X.CHR,IRanges(POS,width=1),A=Count_A,C=Count_C,G=Count_G,T=Count_T,coverage=Good_depth))
  m = match(out,SNPs)
  out$ref = snps$REF[m]
  out$alt = snps$ALT[m]
  out$AF = snps$AF[m]
  out$altCount = as.matrix(mcols(out)[,bases])[cbind(seq_along(out),match(unlist(out$alt),bases))]
  out$BAF = out$altCount/out$coverage
  #Do the basic cuts
  out = out[out$coverage>minCoverage & abs(out$BAF-0.5)>0.5-errorRate,]
  #Now get the ones we're very sure are homozygous
  pVals = pmin(
               pbinom(out$altCount-1,out$coverage,0.5+errorRate,lower.tail=FALSE),
               pbinom(out$altCount,out$coverage,0.5-errorRate)
               )
  qVals = p.adjust(pVals,method='BH')
  out = out[qVals<.01,]
  ########################################
  # Count homozygous alleles in 10X data #
  ########################################
  #Do the 10X quantification
  dat = phaseExpression(out,RNA_BAMs,ref,gtf,labels=labels,outDir=outDir,nParallel=nParallel,...)
  #Now get the non-child expression count
  m = match(dat,out)
  dat$BAF = out$BAF[m]
  dat$alt = out$alt[m]
  dat$ref = out$ref[m]
  dat$AF = out$AF[m]
  dat$childBase = ifelse(dat$BAF>0.5,dat$alt,dat$ref)
  dat$notChildBase = ifelse(dat$BAF>0.5,dat$ref,dat$alt)
  #Get the expression of the childBase
  dat$childBaseCount = as.matrix(mcols(dat)[,bases])[cbind(seq_along(dat),match(dat$childBase,bases))]
  #And the expression of the alt/ref (whatever is not childBase)
  dat$nonChildBaseCount = as.matrix(mcols(dat)[,bases])[cbind(seq_along(dat),match(dat$notChildBase,bases))]
  dat$otherBaseCount = dat$coverage-dat$childBaseCount - dat$nonChildBaseCount
  #We need to do some subtle filtering now.  Mapping artefacts are more likely to support the REF base than the ALT.  For some reason, it seems that empirically mapping artefacts tend to occur more commonly in introns than exons (not just more in an absolute sense, a genuinely higher rate).  This screws up our estimates of non child base counts if we don't correct for it by excluding those SNPs where the child base is ALT.
  #Aggregate across all genes in a cell
  tst = aggregate(cbind(nonChildBaseCount=nonChildBaseCount/coverage>0.1,otherBaseCount=otherBaseCount/coverage>0.1,coverage=coverage>0) ~ cellID ,data=data.frame(dat[dat$regionType=='Exonic' | dat$childBase==dat$ref,]),FUN=sum)
  tst$frac = tst$nonChildBaseCount / tst$coverage
  m = match(tst$cellID,rownames(gold))
  tst$call = gold$call[m]
  #tst = tst[!is.na(tst$call) & tst$regionType!='InterGenic',]
  #The expected median deviation for maternal cells is then...
  expMean=mean(0.5*(0.5-abs(dat$AF[dat$childBase==dat$ref]-0.5)))
  #Make the call 
  tst$call = ifelse(tst$frac > max(error10X,expMean/2),'Mum','Child')
  return(list(cellOfOrigin=tst,dat=dat))
}


#' Infer non-reference cells
#'
#' Use the counts at SNPs with known population AFs to infer which cells differ from a known genotype.  This is assessed by a binomial model allowing for \code{errRate} error and calculating the posterior probability of each cell's origin.  The probability of non-reference reads for the non-reference cell is calculated from the population allele frequencies.  Counts for each allele at each SNP are collapsed to 0 or 1 so that the genotype is not biased by high expression of one SNP.  See \code{\link{inferCellOrigin}}
#'
#' @param phasedExpression The output of \code{\link{phaseExpression}} using SNPs with known AF.
#' @param useX Use the X chromosome if present?
#' @param testForParent If we assume the non-reference cell is derived from the parent the expected number of deviating counts is different by a factor of 2.
#' @param errRate The assumed error rate for the null model.
#' @param pThresh A posterior probability that exceeds this value is considered sufficient evidence to call a cell's origin.
#' @param doPlot Make a plot.
#' @return A table with a genotyping call for each cell or a ggplot object with this table in $df
inferNonRefCells = function(phasedExpression,useX=FALSE,testForParent=TRUE,errRate=0.02,pThresh=0.999,doPlot=FALSE,plotValidOnly=TRUE){
  bases=c('A','C','G','T')
  #First derive count information for the reference and non reference 
  phasedExpression$childBase = ifelse(phasedExpression$BAF>0.5,phasedExpression$alt,phasedExpression$ref)
  phasedExpression$notChildBase = ifelse(phasedExpression$BAF>0.5,phasedExpression$ref,phasedExpression$alt)
  #Get the expression of the childBase
  phasedExpression$childBaseCount = as.matrix(mcols(phasedExpression)[,bases])[cbind(seq_along(phasedExpression),match(phasedExpression$childBase,bases))]
  #And the expression of the alt/ref (whatever is not childBase)
  phasedExpression$nonChildBaseCount = as.matrix(mcols(phasedExpression)[,bases])[cbind(seq_along(phasedExpression),match(phasedExpression$notChildBase,bases))]
  phasedExpression$otherBaseCount = phasedExpression$coverage-phasedExpression$childBaseCount - phasedExpression$nonChildBaseCount
  #We need to do some subtle filtering now.  Mapping artefacts are more likely to support the REF base than the ALT.  For some reason, it seems that empirically mapping artefacts tend to occur more commonly in introns than exons (not just more in an absolute sense, a genuinely higher rate).  This screws up our estimates of non child base counts if we don't correct for it by excluding those SNPs where the child base is ALT.
  wFilt = phasedExpression$regionType=='Exonic' | phasedExpression$childBase==phasedExpression$ref
  if(!useX)
    wFilt = wFilt & as.character(seqnames(phasedExpression))!='X'
  wFilt[is.na(wFilt)]=FALSE
  #Aggregate across all genes in a cell.  Collapse if more than 10% of reads support the non child base.
  tst = aggregate(cbind(nonChildBaseCount=nonChildBaseCount/coverage>0.1,otherBaseCount=otherBaseCount/coverage>0.1,coverage=coverage>0) ~ cellID ,data=data.frame(phasedExpression[wFilt,]),FUN=sum)
  tst$frac = tst$nonChildBaseCount / tst$coverage
  #The expected median deviation for maternal cells is then...
  expMean=mean((0.5-abs(dat$AF[wFilt]-0.5)))
  if(testForParent)
    expMean = expMean/2
  if(expMean < errRate*2)
    warning("The expected mean is close to the stated error rate.  Consider filtering to use SNPs with higher AF.")
  #Now calculate the posterior probabilities
  pRef = with(tst,dbinom(nonChildBaseCount,coverage,errRate))
  pNonRef = with(tst,dbinom(nonChildBaseCount,coverage,expMean))
  #Now normalise
  pNorm = pRef+pNonRef
  tst$pRef = pRef/pNorm
  tst$pNonRef = pNonRef/pNorm
  tst$pMax = pmax(tst$pRef,tst$pNonRef)
  tst$call = with(tst,ifelse(pRef>pThresh,
                             'Reference',
                             ifelse(pNonRef>pThresh,
                                    'NonReference',
                                    'Unclear'
                                    )
                             ))
  tst = tst[order(tst$pMax,decreasing=TRUE),]
  #Add the isValidCell column if present
  if('isValidCell' %in% colnames(mcols(phasedExpression)))
    tst$isValidCell = phasedExpression$isValidCell[match(tst$cellID,phasedExpression$cellID)]
  if(doPlot){
    tmp = tst
    if(plotValidOnly && !is.null(tst$isValidCell))
      tmp = tmp[which(tmp$isValidCell),]
    gg = ggplot(tmp,aes(log10(coverage),frac,colour=pNonRef,shape=call)) +
      geom_point() +
      geom_hline(yintercept=c(error=errRate),colour='red') +
      geom_hline(yintercept=c(exp=expMean),colour='green')+
      geom_hline(yintercept=c(exp=expMean/2),colour='blue')+
      xlab('log10(# informative SNPs)')+
      ylab('Fraction non-reference SNPs')
    gg$df = tst
    return(gg)
  }
  return(tst)
}
 
 


#' Infer the cell of origin
#'
#' Use the counts at phased SNPs to infer if each cell is maternal, child or paternal (should be impossible).  This is assessed using a binomial model allowing for \code{errRate} error and calculating the posterior probability of each cell's origin.  Counts for each allele at each SNP are collapsed to 0 or 1 so that the genotype is not biased by high expression of one SNP. 
#' 
#' The logic is that we expect UMIs for a given SNP to be highly correlated owing to the bursty nature of transcription.  In fact, SNPs should be correlated within a gene due for the same reason and possibly even correlated across genes that are nearby.  We do not correct for this effect at present though.  
#'
#' We should also really exclude SNPs that fall in imprinted regions.
#' 
#' @param phasedExpression The output of \code{phaseExpression}.
#' @param errRate What error rate to tollerate.  This logically equates to the expected contamination for your 10X samples.
#' @param pThresh A posterior probability that exceeds this value is considered sufficient evidence to call a cell's origin.
inferCellOrigin = function(phasedExpression,errRate=0.1,pThresh=0.999){
  dat = data.frame(phasedExpression)
  cellCnts = aggregate(cbind(maternalCount,paternalCount) ~ cellID,data=dat,function(e) sum(e>0))
  rownames(cellCnts) = cellCnts$cellID
  cellCnts = cellCnts[,-match('cellID',colnames(cellCnts))]
  colnames(cellCnts) = c('mum','dad')
  cellCnts$tot = cellCnts$mum+cellCnts$dad
  cellCnts$mumFrac = cellCnts$mum/cellCnts$tot
  #Calculate the posterior probabilities of mum,dad and child
  pMum = with(cellCnts,dbinom(mum,tot,1-errRate))
  pDad = with(cellCnts,dbinom(mum,tot,errRate))
  pChild = with(cellCnts,dbinom(mum,tot,0.5))
  #Now normalise
  pNorm = pMum+pDad+pChild
  cellCnts$pMum = pMum/pNorm
  cellCnts$pDad = pDad/pNorm
  cellCnts$pChild = pChild/pNorm
  cellCnts$pMax = apply(cellCnts[,c('pMum','pDad','pChild')],1,max)
  cellCnts$call = with(cellCnts,ifelse(pMum>pThresh,
                                       'Mum',
                                       ifelse(pDad>pThresh,
                                              'Dad',
                                              ifelse(pChild>pThresh,
                                                     'Child',
                                                     'Unclear'
                                                     )
                                              )
                                       ))
  cellCnts = cellCnts[order(cellCnts$pMax,decreasing=TRUE),]
  #Add the isValidCell column if present
  if('isValidCell' %in% colnames(mcols(phasedExpression)))
    cellCnts$isValidCell = phasedExpression$isValidCell[match(rownames(cellCnts),phasedExpression$cellID)]
  return(cellCnts)
}


findASE = function(phasedExpression){
  out = phasedExpression
  w = which(out$isValidCell & lengths(out$geneName)==1 & !is.na(out$altIsMum) & out$regionType=='Exonic')
  dimNames=list(unique(unlist(out$geneName[w])),unique(out$cellID[w]))
  #Arregate to make matricies
  dat = data.frame(out[w])
  dat$geneName=unlist(dat$geneName)
  mumMat = aggregate(maternalCount ~ cellID + geneName,data=dat,sum)
  mumMat = sparseMatrix(i=match(mumMat$geneName,dimNames[[1]]),
                        j=match(mumMat$cellID,dimNames[[2]]),
                        x=mumMat$maternalCount,
                        dim=lengths(dimNames),
                        dimnames=dimNames)
  dadMat = aggregate(paternalCount ~ cellID + geneName,data=dat,sum)
  dadMat = sparseMatrix(i=match(dadMat$geneName,dimNames[[1]]),
                        j=match(dadMat$cellID,dimNames[[2]]),
                        x=dadMat$paternalCount,
                        dim=lengths(dimNames),
                        dimnames=dimNames)
  #Get the cell counts
  cellCnts = data.frame(mum=colSums(mumMat),dad=colSums(dadMat))
  cellCnts$tot = cellCnts$mum+cellCnts$dad
  cellCnts$mumFrac = cellCnts$mum/cellCnts$tot
  cellCnts$pVal = pbinom(pmax(cellCnts$mum,cellCnts$dad)-1,cellCnts$tot,0.5,lower.tail=FALSE)
  cellCnts$call = ifelse(abs(cellCnts$mumFrac-0.5)>0.3 & p.adjust(cellCnts$pVal,method='BH')<0.01,
                         ifelse(cellCnts$mumFrac>0.5,
                                'MaternalCell',
                                'PaternalCell'
                                ),
                         'ChildCell'
                         )
  #Exclude the non-child cells
  good = rownames(cellCnts)[cellCnts$call=='ChildCell']
  flat = data.frame(geneName = rownames(mumMat[,good]),mum=rowSums(mumMat[,good]),dad=rowSums(dadMat[,good]),mumBinary=rowSums(mumMat[,good]>0),dadBinary=rowSums(dadMat[,good]>0),row.names=seq_len(nrow(dadMat)))
  flat$tot = flat$mum+flat$dad
  flat$mumFrac = flat$mum/flat$tot
  flat$pVal = pbinom(pmax(flat$mum,flat$dad)-1,flat$tot,0.5,lower.tail=FALSE)
  flat = flat[!is.na(flat$geneName),]
  flat = flat[order(abs(flat$mumFrac-0.5),decreasing=TRUE),]
  flat[flat$tot>10 & p.adjust(flat$pVal,method='BH')<0.05 & abs(flat$mumFrac-0.5)>0.3,]
  #Split by sample
  s = split(colnames(mumMat),gsub('___.*','',colnames(mumMat)))
  sampMum = do.call(cbind,lapply(s,function(e) rowSums(mumMat[,e,drop=FALSE])))
  sampMumNonZero = do.call(cbind,lapply(s,function(e) rowSums(mumMat[,e,drop=FALSE]>0)))
  sampDad = do.call(cbind,lapply(s,function(e) rowSums(dadMat[,e,drop=FALSE])))
  sampDadNonZero = do.call(cbind,lapply(s,function(e) rowSums(dadMat[,e,drop=FALSE]>0)))
  sampTot = (sampMum+sampDad)
  sampTotNonZero = sampMumNonZero + sampDadNonZero
  sampMumFrac = sampMum/sampTot
  pVal = pbinom(pmax(sampMumNonZero,sampDadNonZero)-1,sampTotNonZero,0.5,lower.tail=FALSE)
  pVal = pbinom(pmax(sampMum,sampDad)-1,sampTot,0.5,lower.tail=FALSE)
  w = which(abs(sampMumFrac-0.5)>0.3 & p.adjust(pVal,method='BH')<0.01,arr.ind=TRUE)
  tmp=data.frame(sampMum[w[i],],sampDad[w[i],],tot=sampMum[w[i],]+sampDad[w[i],],pVal = pVal[w[i],],mumFrac=sampMumFrac[w[i],],mumCells=sampMumNonZero[w[i],],dadCells=sampDadNonZero[w[i],]);colnames(tmp)[1:2]=paste0(c('Mum','Dad'),'_',rownames(w)[i])
  #Get the cell counts
  cellCnts = data.frame(mum=colSums(mumMat),dad=colSums(dadMat))
  cellCnts$tot = cellCnts$mum+cellCnts$dad
  cellCnts$mumFrac = cellCnts$mum/cellCnts$tot
  cellCnts$pVal = pbinom(pmax(cellCnts$mum,cellCnts$dad)-1,cellCnts$tot,0.5,lower.tail=FALSE)
  cellCnts$call = ifelse(abs(cellCnts$mumFrac-0.5)>0.3 & p.adjust(cellCnts$pVal,method='BH')<0.01,
                         ifelse(cellCnts$mumFrac>0.5,
                                'MaternalCell',
                                'PaternalCell'
                                ),
                         'ChildCell'
                         )
}


#' Use BCFtools to find heterozygous SNPs in a sample
#'
#' Find any base with decent coverage and BAF close to 0.5 and save them a VCF.  The vcf is constructed with loose limits.  A binomial test is then used to determine which locations we can reject the NULL hypothesis of homozygosity with an assumed error rate.  By default the error rate is set high to avoid worst case scenarios. FDR cut of 0.01 is used.
#'
#' @param BAM Path to BAM file to call Het SNPs from.
#' @param ref Path to reference genome to use.
#' @param bin Path to BCF tools binary.
#' @param concatBin Path to vcf-concat binary.
#' @param minCoverage Any location with coverage below this will not be reported.
#' @param BAF_lim Any SNP with BAF outside this range will be discarded.
#' @param VCF_path The path of the VCF file containing SNPs to be created.
#' @param skipIfExists Skip running the SNP caller if the VCF already exists.
#' @param nParallel How many processors to use
#' @param chrsToProcess Which chromosomes to process?  If NULL, all chromosomes in the BAM header are used.  Any chromosomes not present in the BAM header will be dropped.
#' @param errRate Assumed error rate for hypothesis test.
#' @param ... Absorb unused parameters.
findHetSNPs = function(BAM,ref,bin='bcftools',concatBin='vcf-concat',minCoverage=10,BAF_lim=c(0.2,0.8),VCF_path=file.path(getwd(),sprintf('%s_hetSNPs.vcf.gz',basename(BAM))),skipIfExists=TRUE,nParallel=1,chrsToProcess=c('X',1:22),errRate=0.2,...){
  #Get list of chromosomes to process
  chrs = seqlevels(BamFile(BAM))
  if(!is.null(chrsToProcess))
    chrs = chrsToProcess[chrsToProcess %in% chrs]
  if(length(chrs)==0)
    stop("No chromosomes found to process!")
  #Parallelise by piping into xargs.  Inspired by this lovely code https://www.biostars.org/p/48781/#48815
  cmd = sprintf("echo \"%s\" | xargs -I {} -n 1 -P %d sh -c ",paste(chrs,collapse='\n'),nParallel)
  #Construct the command to run
  tmp = sprintf("%s mpileup -Ou -f %s -r {} %s | %s call -cv -V indels | %s filter -Oz -o %s.{} -e '%%SUM(DP4)<%d || (DP4[2]+DP4[3])/(DP4[0]+DP4[1]+DP4[2]+DP4[3]) < %g || (DP4[2]+DP4[3])/(DP4[0]+DP4[1]+DP4[2]+DP4[3]) > %g'",bin,ref,BAM,bin,bin,VCF_path,minCoverage,BAF_lim[1],BAF_lim[2])
  #Find homozygous ALT
  #tmp = sprintf("%s mpileup -Ou -f %s -r {} %s | %s call -cv -V indels | %s filter -Oz -o %s.{} -e '%%SUM(DP4)<%d || (DP4[2]+DP4[3])/(DP4[0]+DP4[1]+DP4[2]+DP4[3]) < %g'",bin,ref,BAM,bin,bin,VCF_path,minCoverage,0.8)
  cmd = paste0(cmd,'"',tmp,'"')
  #Actually run it, or skip if exists
  if(!skipIfExists | !file.exists(VCF_path)){
    system(cmd)
    #Now merge everything back
    system(sprintf('%s -p %s | gzip > %s',concatBin,paste(sprintf('%s.%s',VCF_path,chrs),collapse=' '),VCF_path))
    #Remove intermediate files
    system(sprintf('rm %s',paste(sprintf('%s.%s',VCF_path,chrs),collapse=' ')))
    #Hack because VariantAnnotaion is a poo
    system(sprintf('gunzip %s',VCF_path))
    system(sprintf('bgzip %s',gsub('\\.gz$','',VCF_path)))
  }else{
    message("Previous results found, skipping re-calculation")
  }
  #Read the VCF back in
  vcf = readVcf(VCF_path)
  out = rowRanges(vcf)
  tmp = as.data.frame(as.matrix(info(vcf)$DP4))
  colnames(tmp) = c('fREF','rREF','fALT','rALT')
  mcols(out) = cbind(mcols(out)[,c('REF','ALT','QUAL')],tmp)
  #Filter out anything that has more than one alt
  tmp = tmp[lengths(out$ALT)==1,]
  out = out[lengths(out$ALT)==1,]
  #And do a stronger filter based on the null hypothesis of homozygous+error rate
  tot=rowSums(tmp)
  BAF = (tmp[,3]+tmp[,4])/tot
  pVals = pbinom(ifelse(BAF<0.5,(tmp[,3]+tmp[,4])-1,(tmp[,1]+tmp[,2])-1),
                 tot,
                 errRate,
                 lower.tail=FALSE)
  qVals = p.adjust(pVals,method='BH')
  out = out[qVals<0.01,]
  #Convert REF and ALT to characters
  out$REF = as.character(out$REF)
  out$ALT = as.character(unlist(out$ALT))
  return(out)
}

#' Identifies which Heterozygous SNPs can be phased.
#'
#' Given a list of heterozygous SNPs in a child and allele counts at each of these locations in one or both parents, identifies locations where each allele of the heterozygous SNP can be unambiguously determined to originate from one or other parent.
#'
#' @param hSNPs GRanges object containing heterozygous SNPs in child.  Must have REF and ALT columns.
#' @param mumCnts Output of AlleleCounter for the SNPs in hSNPs on the BAM file from Mum.  Only one parent is necessary, but having both is preferable.
#' @param dadCnts Output of AlleleCounter for the SNPs in hSNPs on the BAM file from Dad.  Only one parent is necessary, but having both is preferable.
#' @param minCoverage A SNP is only assigned if there is coverage exceeding this in at least one parent.
#' @param NullBAF BAF below this value is considered as evidence for absence (together with the minCoverage requirement).
#' @param NotNullBAF BAF above this value is considered as evidence for presence (together with the minCoverage requirement).
#' @param ... Absorb unused parameters.
#' @return A modified hSNPs with extra columns indicating if the SNP is informative and which parent each allele is.
findInformativeSNPs = function(hSNPs,mumCnts=NULL,dadCnts=NULL,minCoverage=10,NullBAF=0.05,NotNullBAF=0.3,...){
  #Do we have Mum and Dad?
  mum=!is.null(mumCnts)
  dad=!is.null(dadCnts)
  if(!mum & !dad)
    stop("Information from at least one parent is required.")
  bases=c('A','C','G','T')
  lab = c('Mum','Dad')
  #Load the allele count information 
  for(i in seq(2)){
    if(c(mum,dad)[i]){
      cnts = read.table(c(mumCnts,dadCnts)[i],sep='\t',header=FALSE)
      tmp = cnts[,3:6]
      colnames(tmp) = paste0(lab[i],'.',bases)
      cnts = GRanges(cnts[,1],IRanges(cnts[,2],width=1))
      mm = match(hSNPs,cnts)
      mcols(hSNPs) = cbind(mcols(hSNPs),tmp[mm,])
      mcols(hSNPs)[,sprintf('%s.REF',lab[i])] = as.matrix(tmp[mm,])[cbind(row=seq_len(length(hSNPs)),col=match(hSNPs$REF,bases))]
      mcols(hSNPs)[,sprintf('%s.ALT',lab[i])] = as.matrix(tmp[mm,])[cbind(row=seq_len(length(hSNPs)),col=match(hSNPs$ALT,bases))]
      mcols(hSNPs)[,sprintf('%s.Depth',lab[i])] = rowSums(tmp[mm,])
      mcols(hSNPs)[,sprintf('%s.BAF',lab[i])] = mcols(hSNPs)[,sprintf("%s.ALT",lab[i])] / mcols(hSNPs)[,sprintf("%s.Depth",lab[i])]
      mcols(hSNPs)[,sprintf('%s.AAF',lab[i])] = mcols(hSNPs)[,sprintf("%s.REF",lab[i])] / mcols(hSNPs)[,sprintf("%s.Depth",lab[i])]
    }
  }
  #Check each allele for absence in each parent.  Record which allele the ALT belongs to
  informative = rep(FALSE,length(hSNPs))
  altIsMum = rep(NA,length(hSNPs))
  sanityFail = rep(FALSE,length(hSNPs))
  for(i in seq(2)){
    if(c(mum,dad)[i]){
      noRef = mcols(hSNPs)[,sprintf('%s.Depth',lab[i])]>minCoverage & mcols(hSNPs)[,sprintf('%s.AAF',lab[i])] < NullBAF
      noAlt = mcols(hSNPs)[,sprintf('%s.Depth',lab[i])]>minCoverage & mcols(hSNPs)[,sprintf('%s.BAF',lab[i])] < NullBAF
      #Can't be missing both, if it is, we're in trouble...
      sanityFail[noRef & noAlt]=TRUE
      altIsMum[noRef] = c(TRUE,FALSE)[i]
      altIsMum[noAlt] = c(FALSE,TRUE)[i]
      informative = informative | noRef | noAlt
    }
  }
  #If we have mum and dad, we can do extra sanity checks
  #Do sanity check if we have both
  if(mum & dad){
    #Make sure that both REF and ALT are present
    hasBoth = pmax(hSNPs$Mum.BAF,hSNPs$Dad.BAF) > NotNullBAF & pmax(hSNPs$Mum.REF/hSNPs$Mum.Depth,hSNPs$Dad.REF/hSNPs$Dad.Depth) > NotNullBAF
    #Where data is missing from one, it's equivalent to not being able to check
    hasBoth[is.na(hasBoth)]=TRUE
    sanityFail[which(!hasBoth)] = TRUE
  }
  #Exclude those that fail the sanity checks
  informative[sanityFail]=FALSE
  #Now save the information and the phasing
  hSNPs$Informative = informative
  hSNPs$altIsMum = NA
  hSNPs$altIsMum[which(informative)] = altIsMum[which(informative)]
  return(hSNPs)
}

#' Phase mutations with SNPs
#'
#' Uses a collection of heterozygous SNPs with known phasing (usually derived from parental DNA) to try and identify which allele a mutation occurs on.  The phasing can be derived from either the reads that support the variant or those that do not.  Inference from the variant supporting reads is straight forward as these must all derive from tumour cells.  However, inference from the reads that do not support the variant may come from either the tumour or the normal.
#'
#' To make use of the reads that do not support the variant, we calculate the posterior probability of the variant being on each allele assuming a maximum likelihood estimate for the contamination (rho).
#'
#' @param BAM BAM file the mutations were called from.
#' @param hSNPs A GRanges object (usually produced by \code{findInformativeSNPs}) that contains all those SNPs that have been phased.
#' @param mutations A GRanges object containing the coordinates of mutated sequence and a "seqs" column containing what the mutated sequence is at this location.
#' @param rho The tumour cell fraction to use when calculating the probability of each allele using WT reads.  Lower values of rho (more contamination) results in more conservative calling.  If set to NULL the maximum likelihood estimator of rho is used for each mutation.
#' @param ref Extract tri-nucleotide context from reference.  Requires samtools
phaseMutations = function(BAM,hSNPs,mutations,rho=NULL,ref=NULL){
  #Keep only the useful SNPs
  hSNPs = hSNPs[which(!is.na(hSNPs$altIsMum)),]
  #Load the mutations from caveman if "mutations" is a caveman file
  if(!is(mutations,'GRanges')){
    if(!file.exists(mutations))
      stop("Could not find mutations caveman file.")
    message("Loading PASS mutations from caveman file")
    mutations = read.table(mutations,header=FALSE,sep='\t')
    colnames(mutations) = c('CHROM','POS','ID','REF','ALT','QUAL','FILTER','INFO','FORMAT','NORMAL','TUMOUR')
    mutations = mutations[mutations$FILTER=='PASS',]
    mutations = GRanges(mutations$CHROM,IRanges(mutations$POS,width=1),REF=mutations$REF,seqs=mutations$ALT)
    #Convert to pyramdine context as necessary for labelling
    toSwap = mutations$REF %in% c('A','G')
    alts = mutations$seqs
    refs = mutations$REF
    alts[toSwap] = as.character(reverseComplement(DNAStringSet(alts[toSwap])))
    refs[toSwap] = as.character(reverseComplement(DNAStringSet(refs[toSwap])))
    labs = sprintf("%s:%s_",seqnames(mutations),start(mutations))
    if(is.null(ref)){
      labs = paste0(labs,paste0(refs,'/',alts))
    }else{
      #Add tri-nucleotide context from reference
      triCon = system(sprintf("samtools faidx %s %s",ref,paste(paste0(seqnames(mutations),':',start(mutations)-1,'-',start(mutations)+1),collapse=' ')),intern=TRUE)
      triCon = triCon[seq(2,length(triCon),2)]
      triCon[toSwap] = as.character(reverseComplement(DNAStringSet(triCon[toSwap])))
      labs = paste0(labs,substr(tolower(triCon),1,1),refs,'/',alts,substr(tolower(triCon),3,3))
    }
    names(mutations)=labs
  }
  #Now do the phasing
  bf = BamFile(BAM,asMates=TRUE)
  muts = list()
  for(ii in seq_along(mutations)){
    tgtLoc = mutations[ii]
    seqlevels(tgtLoc) = as.character(seqnames(tgtLoc))
    param = ScanBamParam(which=tgtLoc,what=scanBamWhat())
    dat = readGAlignments(bf,param=param)
    #Drop mates off target chromosome
    dat = dat[seqnames(dat)%in% seqlevels(tgtLoc)]
    seqlevels(dat) = seqlevels(tgtLoc)
    #Test each location and find where we have a match
    varReads = unique(mcols(dat)$qname)
    for(i in seq_len(width(tgtLoc))){
      tmp = GRanges(seqlevels(tgtLoc),IRanges(start(tgtLoc)+i-1,width=1))
      onTgt = dat[subjectHits(findOverlaps(tmp,dat,type='within'))]
      #Get the reads that contain the variant
      pup = pileLettersAt(mcols(onTgt)$seq,seqnames(onTgt),start(onTgt),cigar(onTgt),tgtLoc)
      #Get the qnames of those that match expectations
      w = as.vector(gregexpr(substr(tgtLoc$seqs,i,i),pup)[[1]])
      if(any(w!=-1)){
        varReads = varReads[varReads %in% mcols(onTgt)$qname[w]]
      }else{
        varReads=c()
        break
      }
    }
    #Split into those that do and don't contain the variant
    mcols(dat)$supportsVar = mcols(dat)$qname %in% varReads
    #Only proceed if there are SNPs within range of the variant supporting reads
    mutSNPs = subsetByOverlaps(hSNPs,dat)
    message(sprintf("Interrogating mutation %d/%d at %s:%d with target sequence %s, found %d SNPs nearby",ii,length(mutations),seqnames(tgtLoc),start(tgtLoc),tgtLoc$seqs,length(mutSNPs)))
    #Get SNPs within range
    if(length(mutSNPs)==0){
      mutSNPs$Var.A = numeric()
      mutSNPs$Var.C = numeric()
      mutSNPs$Var.G = numeric()
      mutSNPs$Var.T = numeric()
      mutSNPs$WT.A = numeric()
      mutSNPs$WT.C = numeric()
      mutSNPs$WT.G = numeric()
      mutSNPs$WT.T = numeric()
      mutSNPs$MumAllele=character()
      mutSNPs$DadAllele=character()
      mutSNPs$VarMumCount=numeric()
      mutSNPs$VarDadCount=numeric()
      mutSNPs$VarOtherCount=numeric()
      mutSNPs$WT_MumCount=numeric()
      mutSNPs$WT_DadCount=numeric()
      mutSNPs$WT_OtherCount=numeric()
    }else{
      seqlevels(mutSNPs) = seqlevels(dat)
      #Get the count of supporting reads
      out=list()
      for(i in seq_along(mutSNPs)){
        cnts = lapply(split(dat,mcols(dat)$supportsVar),function(e) pileLettersAt(mcols(e)$seq,seqnames(e),start(e),cigar(e),mutSNPs[i]))
        cnts = lapply(cnts,letterFrequency,letters=c('A','C','G','T'))
        out[[i]] = cnts
      }
      #Make ACGT for var and WT
      varCnts = data.frame(do.call(rbind,lapply(out,function(e) e$`TRUE`)))
      colnames(varCnts) = paste0('Var.',colnames(varCnts))
      WT_cnts = data.frame(do.call(rbind,lapply(out,function(e) e$`FALSE`)))
      colnames(WT_cnts) = paste0('WT.',colnames(WT_cnts))
      mcols(mutSNPs) = cbind(mcols(mutSNPs),Var=varCnts,WT=WT_cnts)
      #Get the fraction that are maternal and paternal
      mumAllele = data.frame(mcols(mutSNPs)[c('REF','ALT')])[cbind(seq(length(mutSNPs)),ifelse(mutSNPs$altIsMum,2,1))]
      dadAllele = data.frame(mcols(mutSNPs)[c('REF','ALT')])[cbind(seq(length(mutSNPs)),ifelse(mutSNPs$altIsMum,1,2))]
      mutSNPs$MumAllele = mumAllele
      mutSNPs$DadAllele = dadAllele
      mutSNPs$VarMumCount = varCnts[cbind(seq(length(mutSNPs)),match(mumAllele,c('A','C','G','T')))]
      mutSNPs$VarDadCount = varCnts[cbind(seq(length(mutSNPs)),match(dadAllele,c('A','C','G','T')))]
      mutSNPs$VarOtherCount = rowSums(varCnts)-mutSNPs$VarMumCount - mutSNPs$VarDadCount
      mutSNPs$WT_MumCount = WT_cnts[cbind(seq(length(mutSNPs)),match(mumAllele,c('A','C','G','T')))]
      mutSNPs$WT_DadCount = WT_cnts[cbind(seq(length(mutSNPs)),match(dadAllele,c('A','C','G','T')))]
      mutSNPs$WT_OtherCount = rowSums(WT_cnts)-mutSNPs$WT_MumCount - mutSNPs$WT_DadCount
    }
    muts[[ii]] = mutSNPs
  }
  names(muts) = names(mutations)
  #Now flatten it down and return the phased mutations
  mutations$NumPhasedSNPs = lengths(muts)
  #Record the coordinates of the SNPs that are informative
  mutations$SNP_locations = sapply(muts,function(e) paste(names(e),colapse=','))
  mutations$VarMumCount = sapply(muts,function(e) sum(e$VarMumCount))
  mutations$VarDadCount = sapply(muts,function(e) sum(e$VarDadCount))
  mutations$VarOtherCount = sapply(muts,function(e) sum(e$VarOtherCount))
  mutations$WT_MumCount = sapply(muts,function(e) sum(e$WT_MumCount))
  mutations$WT_DadCount = sapply(muts,function(e) sum(e$WT_DadCount))
  mutations$WT_OtherCount = sapply(muts,function(e) sum(e$WT_OtherCount))
  #We can also infer the mutated allele from the WT reads, but this is trickier as it depends on the contamination fraction.  The posterior probability (assuming a uniform prior) that the mutated allele is Mum is
  #dbinom(WT_MumCount,WT_MumCount+WT_DadCount+WT_OtherCount,P)/(dbinom(WT_MumCount,WT_MumCount+WT_DadCount+WT_OtherCount,P)+dbinom(WT_MumCount,WT_MumCount+WT_DadCount+WT_OtherCount,1-P))
  #Where P = (1-rho)/(2-rho) and rho is the tumour cell fraction
  #Of course the tumour cell fraction is unknown, so there are three options:
  #1. Fix rho and calculate posterior probabilities at fixed value (could use prior distribution on rho and marginalise over it, but eh)
  #2. Report the rho at which the posterior probability exceeds a cut-off
  #3. Evaluate at the maximum likelihood value of rho independently for each
  #MLE of rho and determine cut-off
  rhoML = rep(NA,length(mutations))
  rhoCut = rep(NA,length(mutations))
  tgtProb=0.95
  for(i in seq_along(mutations)){
    rhoML[i] = with(mcols(mutations)[i,],
                    ifelse(WT_MumCount==WT_DadCount,
                           0,
                           ifelse(WT_MumCount>WT_DadCount,
                                  optimise(function(rho) dbinom(WT_MumCount,WT_MumCount+WT_DadCount+WT_OtherCount,1/(2-rho),log=TRUE),lower=0,upper=1,maximum=TRUE)[[1]],
                                  optimise(function(rho) dbinom(WT_MumCount,WT_MumCount+WT_DadCount+WT_OtherCount,(1-rho)/(2-rho),log=TRUE),lower=0,upper=1,maximum=TRUE)[[1]]
                                  )
                           )
                    )
    #If you write out the posterior probability in full and solve for when it equals the cut-off you get this expression
    rhoCut[i] = with(mcols(mutations)[i,],
                     ifelse(WT_MumCount==WT_DadCount,
                            NA,
                            ifelse(WT_MumCount>WT_DadCount,
                                   1-(tgtProb/(1-tgtProb))**(1/(WT_DadCount+WT_OtherCount-WT_MumCount)),
                                   1-((1-tgtProb)/tgtProb)**(1/(WT_DadCount+WT_OtherCount-WT_MumCount))
                                   )
                            )
                     )
  }
  mutations$rhoML = rhoML
  #What value of rho to use estimating the posterior
  if(is.null(rho)){
    rho = rhoML
  }else{
    rho = rep(rho,length(rhoML))
  }
  #Calculate the posterior probability of the mutation falling on the maternal allele assuming the Maximum Likelihood value of rho
  OA = mutations$WT_MumCount
  O = OA+mutations$WT_DadCount+mutations$WT_OtherCount
  postMum = dbinom(OA,O,(1-rho)/(2-rho))
  postDad = dbinom(OA,O,1/(2-rho))
  mutations$MutatedAllelePosteriorProbabilityOfMaternal = postMum/(postMum+postDad)
  #Use a cut-off of 0.95 for confident declaration of which allele is which
  mutations$MutatedAlleleFromWT_reads = ifelse(abs(mutations$MutatedAllelePosteriorProbabilityOfMaternal-0.5)<0.45,
                                               'Ambiguous',
                                               ifelse(mutations$MutatedAllelePosteriorProbabilityOfMaternal>0.5,
                                                      'Maternal',
                                                      'Paternal'
                                                      )
                                               )
  #Set as ambiguous if fewer than 90% of reads come from one allele or we have no reads
  mutations$MutatedAlleleFromVarReads = with(mutations,ifelse(VarMumCount==VarDadCount | abs(VarMumCount/(VarMumCount+VarDadCount)-0.5)<0.4,
                                                  'Ambiguous',
                                                  ifelse(VarMumCount>VarDadCount,
                                                         'Maternal',
                                                         'Paternal'
                                                         )
                                                  ))
  #Get a consensus using the two bits of information, flagging any conflicting calls
  mutations$MutatedAlleleConsensus = with(mutations,
                                          ifelse(MutatedAlleleFromVarReads == 'Ambiguous',
                                                 MutatedAlleleFromWT_reads,
                                                 ifelse(MutatedAlleleFromVarReads != MutatedAlleleFromWT_reads & MutatedAlleleFromWT_reads!="Ambiguous",
                                                        'Conflicting',
                                                        MutatedAlleleFromVarReads
                                                        )
                                                 )
                                          )
  #Return mutations and individual SNP data
  return(list(mutations=mutations,phasedSNP_counts=muts))
}

#' Runs allele Counter in parallel over a set of input and output files
#'
#' Employs two methods to parallelise allele counter.  First one processor is assigned to each sample.  If there are processors left over, each sample is split by chromosome and chromosomes are processed in parallel.  For example given 8 processors and 4 samples, all samples are run in parallel with each sample processing 2 chromosomes at a time.  Note that if nParallel is not a multiple of the number of samples, some processors will not be used.  For example, given 10 processors and 4 samples, each sample still only processors 2 chromosomes at a time.
#'
#' @param inputs Path to target loci files (two column chr\tpos, no header) containing the SNPs to count.  Either one per sample or if length 1 the same loci are used for all samples.
#' @param outputs The name of the file to write allele counts for each sample.  Lengths of outputs and bams must match.
#' @param bams The BAM files to get allele counts from.  Lengths of outputs and bams must match.
#' @param nParallel Number of processors to use.  Should be a multiple of number of samples.
#' @param f Allelecounter -f param
#' @param F Allelecounter -F param
#' @param x Include the -x option?
#' @param d Include the -d option?
#' @param m Allelecounter -m param
#' @param q Allelecounter -q param
#' @param r The reference genome for all samples.
parallelAlleleCounter = function(inputs,outputs,bams,nParallel=8,f=0,F=0,x=TRUE,d=TRUE,m=20,q=200,r='~/canpipe/ref/human/1000Genomes_hs37d5/genome.fa',bin='alleleCounter',skipIfExists=TRUE){
  #Split the loci file into multiple files
  chrs=list()
  for(input in unique(inputs)){
    loci = read.table(input,sep='\t',header=FALSE)
    for(subLoci in split(loci,loci[,1])){
      write.table(subLoci,paste0(input,'.',unique(subLoci[,1])),row.names=FALSE,sep='\t',col.names=FALSE,quote=FALSE)
    }
    chrs[[input]] = unique(loci[,1])
  }
  #Construct the base string
  baseCommand = sprintf('%s -l %%s.{} -b %%s -o %%s.{} -r %s -f %d -F %d -m %d -q %d',bin,r,f,F,m,q)
  if(x)
    baseCommand = paste0(baseCommand,' -x')
  if(d)
    baseCommand = paste0(baseCommand,' -d')
  if(length(inputs)==1)
    inputs=rep(inputs,length(bams))
  if(length(inputs)!=length(outputs) | length(inputs)!=length(bams))
    stop("inputs,outputs, and bams must all have the same length.")
  toRun = sprintf(baseCommand,inputs,bams,outputs)
  #Filter out anything where the output file exists
  if(skipIfExists)
    toRun = toRun[!file.exists(outputs)]
  #Expand things out to do one command per chromosomes
  toRun = gsub('\\{\\}','%s',toRun)
  toRun = lapply(seq_along(toRun),function(e) sprintf(toRun[e],chrs[[inputs[e]]],chrs[[inputs[e]]]))
  toRun = unlist(toRun)
  write.table(toRun,paste0(input,'_toRun.tsv'),row.names=FALSE,col.names=FALSE,quote=FALSE)
  system(sprintf("parallel -j %d < %s",nParallel,paste0(input,'_toRun.tsv')))
  ####Split each command by chromosome.  Parallelise by sample first, then chromosome
  ###nPerSample = max(1,floor(nParallel/length(inputs)))
  ###for(i in seq_along(inputs)){
  ###  toRun[i] = sprintf("echo '%s' | xargs -I {} -n 1 -P %d sh -c '%s'",paste(chrs[[inputs[i]]],collapse='\n'),nPerSample,toRun[i])
  ###}
  ####Filter out anything where the output file exists
  ###if(skipIfExists)
  ###  toRun = toRun[!file.exists(outputs)]
  ####Run the commands...
  ####Have to make each command be on "one line"
  ###write.table(gsub('\\n','\\\\n',toRun),paste0(input,'_toRun.tsv'),row.names=FALSE,col.names=FALSE,quote=FALSE)
  ###system(sprintf("parallel -j %d < %s",floor(nParallel/nPerSample),paste0(input,'_toRun.tsv')))
  system(sprintf('rm %s',paste0(input,'_toRun.tsv')))
  #Merge the results.  This could be cleaned up to not call redundant bits, but it works for now.
  for(i in seq_along(outputs)){
    #Skip if it already exists and we're doing that
    if(skipIfExists & file.exists(outputs[i]))
      next
    #Create an output file with the header
    system(sprintf("head -n 1 %s > %s",sprintf("%s.%s",outputs[i],chrs[[inputs[i]]])[1],outputs[i]))
    #Add in the calls, excluding the header
    system(sprintf('cat %s|grep -v "^#" >> %s',paste(sprintf("%s.%s",outputs[i],chrs[[inputs[i]]]),collapse=' '),outputs[i]))
    #Clean up temp files
    system(sprintf("rm %s",paste(sprintf("%s.%s",outputs[i],chrs[[inputs[i]]]),collapse=' ')))
  }
  #Remove the split loci files.  
  system(sprintf("rm %s",paste(sprintf("%s.%s",inputs[i],chrs[[inputs[i]]]),collapse=' ')))
}

#' Phase using CN
#'
#' Given CN segments and a set of heterozygous SNPs, uses the observed allele frequencies in the tumour BAM file to phase the SNPs within each segment.  The allele with the higher copy number is designated maternal arbitrarily.
#'
#' @param hSNPs A set of heterozygous SNPs.
#' @param tumBAM The BAM file for the tumour DNA.
#' @param segs The copy number segments to use for phasing.
#' @param lab The base of the file to store results in.
#' @param FDR False discovery rate for binomial test for deviation for 0.5 used when phasing.
#' @param nParallel How many cores to use.
#' @param skipIfExists Skip recalculation if results exist.
#' @return A modified version of \code{hSNPs} that is subset to those SNPs that overlap with \code{segs} and contains guesses at the phasing for each.
phaseUsingCN = function(hSNPs,tumBAM,segs,lab,ref,nParallel=1,FDR=0.05,skipIfExists=TRUE){
  #Get just the SNPs we care about
  hSNPs = subsetByOverlaps(hSNPs,segs,ignore.strand=TRUE)
  ##################################
  # Run allelecounter if necessary
  message("Counting alleles at SNPs In tumour...")
  output = paste0(lab,'_alleleCounts.tsv')
  #Do we really have to run anything?
  if(!skipIfExists | !file.exists(output)){
    #Yes we do, so create the loci filE
    tgtLoci = paste0(lab,'_hetSNPs.tsv')
    write.table(cbind(as.character(seqnames(hSNPs)),start(hSNPs)),tgtLoci,row.names=FALSE,sep='\t',col.names=FALSE,quote=FALSE)
    #Now run allele counter with things sensible for bulk DNA.  
    parallelAlleleCounter(tgtLoci,output,tumBAM,x=FALSE,f=3,F=3852,m=20,q=35,r=ref,skipIfExists=skipIfExists,d=TRUE,nParallel=nParallel)
    system(sprintf('rm %s',tgtLoci))
  }else{
    message("Previous results found, skipping re-calculation")
  }
  #Now load the results
  cnts = read.table(output,sep='\t',header=TRUE,comment.char='')
  tmp = cnts[,3:6]
  colnames(tmp) = gsub('Count_','',colnames(tmp))
  bases = colnames(tmp)
  cnts = GRanges(cnts[,1],IRanges(cnts[,2],width=1))
  mm = match(hSNPs,cnts)
  mcols(hSNPs) = cbind(mcols(hSNPs),tmp[mm,])
  mcols(hSNPs)[,sprintf('cntsTumRef')] = as.matrix(tmp[mm,])[cbind(row=seq_len(length(hSNPs)),col=match(hSNPs$REF,bases))]
  mcols(hSNPs)[,sprintf('cntsTumAlt')] = as.matrix(tmp[mm,])[cbind(row=seq_len(length(hSNPs)),col=match(hSNPs$ALT,bases))]
  mcols(hSNPs)[,sprintf('cntsTumDepth')] = rowSums(tmp[mm,])
  mcols(hSNPs)[,sprintf('tumBAF')] = mcols(hSNPs)[,'cntsTumAlt'] / mcols(hSNPs)[,'cntsTumDepth']
  mcols(hSNPs)[,sprintf('tumAAF')] = mcols(hSNPs)[,'cntsTumRef'] / mcols(hSNPs)[,'cntsTumDepth']
  #Get those cnts that are significantly away from 0.5.
  pVal = pmin(pbinom(hSNPs$cntsTumAlt-1,hSNPs$cntsTumDepth,0.5,lower.tail=FALSE),pbinom(hSNPs$cntsTumAlt,hSNPs$cntsTumDepth,0.5,lower.tail = TRUE))
  qVal = p.adjust(pVal,method='BH')
  hSNPs$qPhase = qVal
  hSNPs$matAllele = NA
  hSNPs$patAllele = NA
  w = which(qVal<FDR)
  #Arbitrarily designate the allele with the higher BAF the maternal one
  hSNPs$matAllele[w] = ifelse(hSNPs$tumBAF[w]>0.5,hSNPs$ALT[w],hSNPs$REF[w])
  hSNPs$patAllele[w] = ifelse(hSNPs$tumBAF[w]>0.5,hSNPs$REF[w],hSNPs$ALT[w])
  return(hSNPs)
}

exonsFromGTF =function(gtf){
  ann = rtracklayer::import(gtf)
  exns = ann[ann$type=='exon']
  return(exns)
}


#' Checks if BAM files have matching genotype. 
#'
#' Looks for locations that support the ALT allele in the target regions in a reference sample.  Then counts at those locations in all BAMs to see if the genotypes match.  Intended use is with a \code{regions} parameter containing exons.
#'
#' @param BAMs The BAM files to genotype.  Names are used as labels.
#' @param refGenomes The FASTA file for the genome of each sample.
#' @param outBase The path and base of the output filename.
#' @param snps A GRanges object with SNPs to inspect.  Must contain REF and ALT columns.  If given, refSample and regions are ignored. 
#' @param refSample Which BAM is the reference sample.
#' @param regions Which regions to look for genotyping SNPs.  Ignored if \code{snps} given.
#' @param is10X If a BAM is 10X, set to TRUE so it can be counted properly.
#' @param bin bcftools binary.
#' @param concatBin vcf-concat binary.
#' @param minCoverage Minimum coverage to accept in reference sample.
#' @param BAF_lim Minimum BAF to accept in the reference sample.
#' @param errRate What is the maximum error rate.
#' @param FDR What False Discover Rate to use.
#' @param skipIfExists Skip calculations if output files exist.
#' @param liftOvers If one or more of the BAMs uses a different coordinate system to the reference, lift the coordinates over with these files.  Vector of length BAMs, with NAs for no-liftover.
#' @param tgtChrs Which chromosomes to use.
#' @param useChr Does each sample stick a 'chr' prefix on chromosome name?
#' @param nParallel How many parallel threads to use.
matchSamples = function(BAMs,refGenomes,outBase,snps,refSample,regions,liftOvers=NULL,is10X=FALSE,useChr=FALSE,bin='bcftools',concatBin='vcf-concat',minCoverage=10,BAF_lim=0.2,errRate=0.25,FDR=0.01,skipIfExists=TRUE,tgtChrs=1:22,nParallel=1){
  if(is.null(names(BAMs)))
    names(BAMs) = gsub('\\.gsub$','',basename(BAMs))
  if(is.null(liftOvers))
    liftOvers=NA
  if(length(liftOvers)==1)
    liftOvers = rep(liftOvers,length(BAMs))
  if(length(is10X)==1)
    is10X = rep(is10X,length(BAMs))
  if(length(refGenomes)==1)
    refGenomes = rep(refGenomes,length(BAMs))
  if(length(useChr)==1)
    useChr = rep(useChr,length(BAMs))
  if(!missing(snps)){
    #Check that everything is what it should be
    if(all(c('REF','ALT') %in% colnames(snps)))
      stop("snps must have REF and ALT")
    if(!is.character(snps$REF))
      stop("REF and ALT must be a character vector")
    if(!is.character(snps$ALT))
      stop("REF and ALT must be a character vector")
    refSNPs = snps[nchar(snps$REF)==1 & nchar(snps$ALT)==1]
  }else{
    if(!is.numeric(refSample))
      refSample = match(refSample,BAMs)
    #Calculate the SNPs in the one that is reference
    #Reference samples get their SNPs called directly from the data.
    BAM = BAMs[refSample]
    nom = names(BAMs)[refSample]
    VCF_path = paste0(outBase,'_SNPs_',nom,'.vcf.gz')
    #Get list of chromosomes to process
    chrs = seqlevels(BamFile(BAM))
    if(!is.null(tgtChrs))
      chrs = tgtChrs[tgtChrs %in% chrs]
    if(length(chrs)==0)
      stop("No chromosomes found to process!")
    tgtLoci = regions[as.character(seqnames(regions)) %in% chrs,]
    chrs =chrs[chrs %in% as.character(seqnames(tgtLoci))]
    #Parallelise by piping into xargs.  Inspired by this lovely code https://www.biostars.org/p/48781/#48815
    tgtLoci = data.frame(chr=as.character(seqnames(tgtLoci)),
                         start=start(tgtLoci),
                         stop=end(tgtLoci))
    tDir = tempdir()
    tFile = tempfile(tmpdir=tDir)
    for(chr in chrs)
      write.table(tgtLoci[tgtLoci$chr==chr,],paste0(tFile,'.',chr),row.names=FALSE,col.names=FALSE,sep='\t',quote=FALSE)
    cmd = sprintf("echo \"%s\" | xargs -I {} -n 1 -P %d sh -c ",paste(chrs,collapse='\n'),nParallel)
    #Construct the command to run
    tmp = sprintf("%s mpileup -Ou -f %s -T %s.{} %s | %s call -cv -V indels | %s filter -Oz -o %s.{} -e '%%SUM(DP4)<%d || (DP4[2]+DP4[3])/(DP4[0]+DP4[1]+DP4[2]+DP4[3]) < %g'",bin,refGenomes[refSample],tFile,BAM,bin,bin,VCF_path,minCoverage,BAF_lim)
    cmd = paste0(cmd,'"',tmp,'"')
    #Actually run it, or skip if exists
    if(!skipIfExists || !file.exists(VCF_path)){
      system(cmd)
      #Now merge everything back
      system(sprintf('%s -p %s | gzip > %s',concatBin,paste(sprintf('%s.%s',VCF_path,chrs),collapse=' '),VCF_path))
      #Remove intermediate files
      system(sprintf('rm %s',paste(sprintf('%s.%s',VCF_path,chrs),collapse=' ')))
      system(sprintf('rm %s.*',tFile))
      #Hack because VariantAnnotaion is a poo
      system(sprintf('gunzip %s',VCF_path))
      system(sprintf('bgzip %s',gsub('\\.gz$','',VCF_path)))
    }else{
      message("Previous results found, skipping re-calculation")
    }
    #Read the VCF back in
    vcf = readVcf(VCF_path)
    out = rowRanges(vcf)
    tmp = as.data.frame(as.matrix(info(vcf)$DP4))
    colnames(tmp) = c('fREF','rREF','fALT','rALT')
    mcols(out) = cbind(mcols(out)[,c('REF','ALT','QUAL')],tmp)
    #Filter out anything that has more than one alt
    tmp = tmp[lengths(out$ALT)==1,]
    out = out[lengths(out$ALT)==1,]
    #And do a stronger filter based on the null hypothesis of homozygous for the reference +error rate
    tot=rowSums(tmp)
    BAF = (tmp[,3]+tmp[,4])/tot
    pVals = pbinom((tmp[,3]+tmp[,4])-1,
                   tot,
                   errRate,
                   lower.tail=FALSE)
    qVals = p.adjust(pVals,method='BH')
    out = out[BAF>=0.5 | qVals<FDR,]
    #Convert REF and ALT to characters
    out$REF = as.character(out$REF)
    out$ALT = as.character(unlist(out$ALT))
    refSNPs = out
  }
  #Obliterate chr if it's there
  refSNPs = renameSeqlevels(refSNPs,setNames(gsub('^chr','',seqlevels(refSNPs)),seqlevels(refSNPs)))
  #Useful for matching when using liftover
  refSNPs$snpID = paste0('SNP',seq_along(refSNPs))
  #Add default AF if not given
  if(!'AF' %in% colnames(refSNPs))
    refSNPs$AF=0.5
  #Convert NAs to string NA so we can work with them in a standard way
  liftOvers[is.na(liftOvers)]='NA'
  #Do each split differently
  outputs = paste0(outBase,'_alleleCounts_',names(BAMs),'.tsv')
  snpCnts = list()
  for(tgtLiftOver in unique(liftOvers)){
    for(tgt10X in c(TRUE,FALSE)){
      for(tgtUseChr in c(TRUE,FALSE)){
        for(refGenome in unique(refGenomes)){
          #Check if there are any matches for this combinatio
          w = liftOvers==tgtLiftOver & is10X==tgt10X & useChr==tgtUseChr & refGenomes==refGenome
          if(any(w)){
            message(sprintf("Processing %d samples",sum(w)))
            #What are the outputs
            tgtOuts = outputs[w]
            #Make the loci file
            tFile = tempfile()
            #Do we need to do the liftover?
            if(tgtLiftOver!='NA'){
              ch = rtracklayer::import.chain(tgtLiftOver)
              tgtSNPs = unlist(rtracklayer::liftOver(refSNPs,ch))
            }else{
              tgtSNPs = refSNPs
            }
            if(tgtUseChr){
              tgtSNPs = renameSeqlevels(tgtSNPs,setNames(paste0('chr',seqlevels(tgtSNPs)),gsub('chr','',seqlevels(tgtSNPs))))
            }
            write.table(cbind(as.character(seqnames(tgtSNPs)),start(tgtSNPs)),tFile,row.names=FALSE,sep='\t',col.names=FALSE,quote=FALSE)
            if(!skipIfExists || any(!file.exists(tgtOuts))){
              if(tgt10X){
                parallelAlleleCounter(tFile,tgtOuts,BAMs[w],r=refGenome,x=TRUE,nParallel=nParallel,skipIfExists=skipIfExists)
              }else{
                parallelAlleleCounter(tFile,tgtOuts,BAMs[w],x=FALSE,f=3,F=3852,m=20,q=35,r=refGenome,skipIfExists=skipIfExists,d=TRUE,nParallel=nParallel)
              }
            }
            system(sprintf('rm %s',tFile))
            #Read them back in
            out = lapply(tgtOuts,read.table,sep='\t',header=TRUE,comment.char='')
            if(tgt10X){
              #We don't actually care about the cellID column for 10X, so aggregate over it
              out = lapply(out,function(e) {
                             tmp = aggregate(cbind(Count_A,Count_C,Count_G,Count_T,Good_depth) ~ paste0(X.CHR,'___',POS),data=e,FUN=sum)
                             data.frame(X.CHR = gsub('___.*','',tmp[,1]),
                                        POS = gsub('.*___','',tmp[,1]),
                                        Count_A = tmp$Count_A,
                                        Count_C = tmp$Count_C,
                                        Count_G = tmp$Count_G,
                                        Count_T = tmp$Count_T,
                                        Good_depth = tmp$Good_depth
                                        )
                 })
            }
            #Add the SNP ID back in
            out = lapply(out,function(e) {
                           m = match(paste0(e[,1],':',e[,2]),as.character(tgtSNPs))
                           e$snpID = tgtSNPs$snpID[m]
                           e})
            #Put them where they should go
            for(i in seq_along(out)){
              snpCnts[[(names(BAMs)[w])[i]]] = out[[i]]
            }
          }
        }
      }
    }
  }
  #Make into a giant comparison matrix
  cnts = matrix(NA,
                nrow=length(refSNPs),
                ncol=length(BAMs),
                dimnames = list(refSNPs$snpID,names(BAMs))
                )
  altCnts = cnts
  #Fill in each
  for(i in seq_along(snpCnts)){
    jj = match(names(snpCnts)[i],colnames(cnts))
    ii = match(snpCnts[[i]]$snpID,refSNPs$snpID)
    #Get the Alt allele frequency for each
    bases=c('A','C','G','T')
    tmp = snpCnts[[i]]
    colnames(tmp) = gsub('Count_','',colnames(tmp))
    tmp = as.matrix(tmp[,bases])
    m = match(snpCnts[[i]]$snpID,refSNPs$snpID)
    cnts[ii,jj] = rowSums(tmp)
    altCnts[ii,jj] = tmp[cbind(seq_along(ii),match(refSNPs$ALT[ii],bases))]
  }
  #What is the evidence for AA,Aa,aa?
  pAA = dbinom(altCnts,cnts,0.1,log=TRUE)
  pAa = dbinom(altCnts,cnts,0.5,log=TRUE)
  paa = dbinom(altCnts,cnts,0.9,log=TRUE)
  #Work out the normalised prob of aa, the most interesting one
  pp = data.frame(pAA=as.vector(pAA),
                  pAa=as.vector(pAa),
                  paa=as.vector(paa))
  pp = exp(pp)/rowSums(exp(pp),na.rm=TRUE)
  pp$sampleID = rep(colnames(cnts),each=nrow(cnts))
  pp$snpID = rep(rownames(cnts),ncol(cnts))
  pp$snpAF = rep(refSNPs$AF,ncol(cnts))
  #Make call of the number of Alt alleles
  pp$numAlt = rep(3,nrow(pp))
  pp$numAlt[pp$paa>0.9]=2
  pp$numAlt[pp$pAa>0.9]=1
  pp$numAlt[pp$pAA>0.9]=0
  #Make this into a genotype matrix
  gtMat = matrix(3,
                 nrow=length(refSNPs),
                 ncol=length(BAMs),
                 dimnames = list(refSNPs$snpID,names(BAMs))
                 )
  #Fill it in
  gtMat[cbind(match(pp$snpID,rownames(gtMat)),match(pp$sampleID,colnames(gtMat)))]=pp$numAlt
  #Make the necessary object for relatdness
  gdsFile = tempfile()
  snpgdsCreateGeno(gdsFile,
                   genmat = gtMat,
                   sample.id=colnames(gtMat),
                   snp.id=rownames(gtMat),
                   snp.chromosome = as.character(seqnames(refSNPs)),
                   snp.position = start(refSNPs),
                   snp.allele = paste0(refSNPs$ALT,'/',refSNPs$REF),
                   snpfirstdim=TRUE,
                   )
  gds = snpgdsOpen(gdsFile)
  gdsSub = snpgdsLDpruning(gds,ld.threshold=0.2)
  ibs = snpgdsIBS(gds,num.thread=nParallel)
  #Return everything
  return(list(ibs=ibs,pp=pp,snpCnts=snpCnts,refSNPs))
}
```
