## supplementary scripts for "Tumor to normal single cell mRNA comparisons reveal a pan-neuroblastoma cancer cell": reformatDutchBAMs.py

```
import pysam
import re
from os.path import expanduser as eu
from os import listdir
from os.path import join,basename,exists

umi = re.compile('.*umi=([A-Z]+)')
cbc = re.compile('.*cbc=([A-Z]+)')
tgtDir = eu('~/trueHome/scratch/ProjectsExtras/SCP/Data/dutchTumourBAMs/RNA/')
tgtFiles = [join(tgtDir,x) for x in listdir(tgtDir) if x.find('.bam')!=-1]
tgtFiles = [x for x in tgtFiles if x.find('sanitised.bam')==-1]
pollFreq = 1000000
for tgtFile in tgtFiles:
  #Check if it's already done
  try:
    tgtOut = re.sub('\\.bam$','_sanitised.bam',tgtFile)
    if exists(tgtOut):
      print("Sample already processed.")
      continue
    print("Adding 10X tags to file " + basename(tgtFile))
    inFile = pysam.AlignmentFile(tgtFile,'rb')
    outFile = pysam.AlignmentFile(tgtOut,'wb',template=inFile)
    i=0
    for x in inFile:
      ub = umi.match(x.qname).group(1)
      cb = cbc.match(x.qname).group(1)
      if(len(ub)==0 or len(cb)==0):
        raise ValueError("Barcodes not found in "+x.qname)
      x.tags += [('CB',cb)]
      x.tags += [('UB',ub)]
      outFile.write(x)
      i= i+1
      if(i % pollFreq==0):
        print('Wrote %d million lines'%(i//pollFreq))
    outFile.close()
    inFile.close()
  except:
    print("Sample failed :(")
    next
      


#Run in bash after above has run
#Samtools sort
for dir in NB*
do
  echo ""
  echo "Processing patient ${dir}"
  echo ""
  mkdir -p Reprocessed/${dir}
  cd $dir
  for file in *sorted.bam
  do
    echo "Sorting ${file}"
    samtools sort -T /tmp/aligned.sorted -@ 4 -o ../Reprocessed/${dir}/${file} ../${file%%_sorted.bam}.bam
  done
  cd ../Reprocessed/${dir}
  for file in *sorted.bam
  do
    echo "Indexing ${file}"
    samtools index ${file}
  done
  cd ../..
done
```
