## supplementary scripts for "Tumor to normal single cell mRNA comparisons reveal a pan-neuroblastoma cancer cell": rnaVelocity.R

```
#Run RNA velocity on fAdrenal 
setwd('~/trueHome/Projects/SCP')

library(Seurat)
library(velocyto.R)

#Load the object
srat = readRDS('Results/preProcess/fAdrenal/processedSeurat.RDS')
#And one loom to test
ldats = 'Data/velocyto/bilateralAdrenals'
ldats = file.path(ldats,list.files(ldats))
ldats = c(ldats,
          'Data/velocyto/babyAdrenal1.loom',
          'Data/velocyto/babyAdrenal2.loom')
ldats = lapply(ldats,read.loom.matrices)
#Extract and merge the emat and nmat things
emat = do.call(cbind,lapply(ldats,function(e) e$spliced))
nmat = do.call(cbind,lapply(ldats,function(e) e$unspliced))
sratIDs = paste0(gsub('_',':',rownames(srat@meta.data)),'x')
emat = emat[,colnames(emat) %in% sratIDs]
nmat = nmat[,colnames(nmat) %in% sratIDs]
m = match(colnames(emat),sratIDs)
clustLab = as.character(srat@meta.data$seurat_clusters[m])
names(clustLab) = colnames(emat)
emb = srat@reductions$umap@cell.embeddings[m,]
rownames(emb) = colnames(emat)
clustLab[clustLab=='28'] = '27'
clustLab[clustLab=='21'] = '22'
emat = filter.genes.by.cluster.expression(emat,clustLab,min.max.cluster.average = 0.5)
nmat = filter.genes.by.cluster.expression(nmat,clustLab,min.max.cluster.average = 0.05)
length(intersect(rownames(emat),rownames(emat)))
fitQuant = 0.02
rVel = gene.relative.velocity.estimates(emat,nmat,deltaT=1,kCells=20,fit.quantile=fitQuant)
#Plot it on embedding for just the medulla
gg = show.velocity.on.embedding.cor(emb,rVel,
                                    n=300,
                                    scale='sqrt',
                                    cex=0.8,
                                    arrow.scale=5,
                                    show.grid.flow=TRUE,
                                    min.grid.cell.mass=0.5,
                                    grid.n=40,
                                    arrow.lwd=1,
                                    do.par=F,
                                    return.details=TRUE,
                                    cell.border.alpha = 0.1)
```
