## supplementary scripts for "Tumor to normal single cell mRNA comparisons reveal a pan-neuroblastoma cancer cell": scGenotyping.R

```
#Genotype single cell data using DNA
setwd('~/trueHome/Projects/SCP')

#############
# Libraries #
#############
library(GenomicRanges)
library(ComplexHeatmap)
library(HoneyBADGER)
library(TxDb.Hsapiens.UCSC.hg19.knownGene)
library(biomaRt)
library(Seurat)
library(VGAM)
import('Code/phaseMutations.R',as='ph')


##############
# Parameters #
##############
outDir = 'Results/Genotyping/'
#DNA reference genome
#hg19 reference
refDNA = 'Data/genome.fa'
#GRCh38 reference genome
ref10X = 'Data/genome10X_GRCh38_120.fa'
#And GTF
gtf = 'Data/gtf10X_GRCh38_120.gtf'
#And the liftover file
hg19to38 = 'Data/hg19ToHg38_noChr.over.chain'
#Minimum length of CN segment to use
minSegLen = 1e6
#How many cores
nParallel=16
#Which types of regions to use
regionsToUse = c('Exonic','Intronic')
#Error rate in sequencing
errRate = 0.01
#Do we need to liftover?
liftOverChain = hg19to38
#Store the transcript object
txdb = TxDb.Hsapiens.UCSC.hg19.knownGene
chrLens = seqlengths(seqinfo(txdb))
names(chrLens) = gsub('chr','',names(chrLens))
#Load gene annotation
ann = rtracklayer::import('Data/gtf10X_GRCh38_120.gtf')
gns = ann[ann$type=='gene']
gns$TSS = ifelse(strand(gns)=='+',start(gns),end(gns))

#############
# Functions #
#############

#' Determines posterior probability of each cell being a tumour cell
#' 
#' The assumption is everything is given using chrosome names that do **NOT** begin with 'chr'.  Where we rely on BAM files where this is not true, 'chr' is temporarily added then removed.
#' 
#' @param Tumour Path to tumour BAM file (or vcf if using array mode)
#' @param Normal Path to Normal BAM
#' @param CN Either a battenberg/ascat summary file or a series of regions in the format chr:start-stop,minCN,totCN.  start/stop can contain N which is replaced by the chromosome start or end.
#' @param scBAMs The BAM files containing scRNA-seq reads.
#' @param scData Either a Seurat object or cellranger output folders to make a Seurat object with.  Used to determine which cells to genotype.
#' @param outBase Base path to use for file names.
#' @param GTF The GTF used to create the quantification of the scBAMs.
#' @param refDNA fasta file for the reference genome used for Tumour/Normal BAMs.
#' @param refRNA As above, but for the scRNA-seq BAMs.
#' @param liftOverChain If the scRNA-seq data uses a difference reference build to the DNA, the lift over file needed to convert from the DNA reference to RNA.
#' @param errRate The assumed error rate for the genotyping model.  Setting this lower will make it harder for cells to be called Tumour.
#' @param minSegLen Drop CN segements shorter than this.
#' @param tgtChrs Drop all but these chromosomes.
#' @param regionsToUse Only use SNPs that fall in these types of regions.  Must be some combination of 'Exonic', 'Intronic', or 'InterGenic'.
#' @param odFiat If you want to force a certain over-dispersion 'rho' (see \code{\link{dbetabinom}}) set it here.
#' @param neutCells If you have prior information that certain cells are likely to be neutral, pass it here.
#' @param tumUsesChr Does the tumour BAM have 'chr' at the start of chromosome names?
#' @param scUsesChr Do the single cell BAMs have 'chr' at the start of chromosome names?
#' @param arrayMode Is the input genotyping arrays. 
#' @param skipIfExists Don't recalculate things from BAMs if output files exist.
#' @param nParallel How many threads to use.
callTumourNormal = function(Tumour,Normal,CN,scBAMs,scData,outBase,GTF,refDNA,refRNA=refDNA,liftOverChain=NULL,errRate=0.01,minSegLen=1e6,tgtChrs=1:22,regionsToUse=c('Exonic','Intronic'),odFiat=NULL,neutCells=NULL,tumUsesChr=FALSE,scUsesChr=FALSE,arrayMode=FALSE,skipIfExists=TRUE,nParallel=1){
  #Create or use Seurat object
  if(length(scData)==1 && !is.character(scData)){
    #Assuming we've been passed a Seurat object
    srat = scData
  }else{
    message(sprintf("Loading single cell count data"))
    scSRC = file.path(scData,'filtered_gene_bc_matrices','GRCh38')
    names(scSRC) = names(scData)
    if(is.null(names(scData)))
      names(scSRC) = basename(dirname(scData))
    srat = Read10X(scSRC)
    srat = CreateSeuratObject(srat)
    srat = NormalizeData(srat,verbose=FALSE)
    srat = FindVariableFeatures(srat,verbose=FALSE)
    srat = ScaleData(srat,verbose=FALSE)
    srat = RunPCA(srat,npcs = 75,approx=FALSE,verbose=FALSE)
    srat = FindNeighbors(srat,dims=seq(75),verbose=FALSE)
    srat = FindClusters(srat,verbose=FALSE)
    srat = RunUMAP(srat,dims=seq(75),verbose=FALSE)
  }
  cellWhitelist = rownames(srat@meta.data)
  ##################
  # Get CN segments
  message(sprintf("Loading copy number segments"))
  if(length(CN)>1 || !file.exists(CN)){
    #It's just a list of segments
    chrs = gsub('[:,].*','',CN)
    starts=rep(1,length(CN))
    ends = chrLens[chrs]
    #If we have the coordinates, extract them
    w = grepl(':',CN)
    if(length(w)>0){
      coords = gsub(',.*','',CN[w]) 
      tmp = gsub('.*:([^-]+)-([^-]+)','\\1',coords)
      tmp[tmp=='N']='1'
      starts[w] = as.numeric(tmp)
      tmp = gsub('.*:([^-]+)-([^-]+)','\\2',coords)
      tmp[tmp=='N']=as.character(chrLens[(chrs[w])[tmp=='N']])
      ends[w] = as.numeric(tmp)
    }
    #Get the segment numbers
    tmp = strsplit(CN,',')
    tumMin = as.numeric(sapply(tmp,`[`,2))
    tumTot = as.numeric(sapply(tmp,`[`,3))
    segs = GRanges(chrs,IRanges(starts,ends),totCN=tumTot,minCN=tumMin)
  }else{
    segs = read.table(CN,
                      sep=',',
                      header=FALSE,
                      col.names = c('Idx','Chr','Start','Stop','normTot','normMin','tumTot','tumMin')
                      )
    #Keep only informative ones
    segs = segs[segs$Stop-segs$Start>minSegLen & segs$tumMin!=segs$tumTot/2,]
    segs = GRanges(segs$Chr,IRanges(segs$Start,segs$Stop),totCN=segs$tumTot,minCN=segs$tumMin)
    segs = segs[seqnames(segs) %in% tgtChrs]
  }
  ###############
  # Get het SNPs
  #Do we just have an array?  In which case, need to do some extra work.
  if(arrayMode){
    message("Getting SNPs from arrays")
    #First get all the potential het SNPs.
    hSNPs = readVcf(Tumour)
    tmp = rowRanges(hSNPs)
    tmp$BAF = geno(hSNPs)$BAF[,1]
    tmp$qual = geno(hSNPs)$IGC[,1]
    tmp$logR = geno(hSNPs)$LRR[,1]
    tmp$GT = geno(hSNPs)$GT[,1]
    tmp$GQ = geno(hSNPs)$GQ[,1]
    tmp$alleleA = info(hSNPs)$ALLELE_A
    tmp$alleleB = info(hSNPs)$ALLELE_B
    #Correct the BAF to ensure B allele refers to the Alt allele
    #w = which(tmp$Allele_B=='0')
    #tmp$BAF[w] = 1-tmp$BAF[w]
    #Ensure character and  biallelic
    tmp$REF = as.character(tmp$REF)
    tmp = tmp[lengths(tmp$ALT)==1]
    tmp$ALT = as.character(unlist(tmp$ALT))
    tmp = tmp[nchar(tmp$ALT)==1]
    #Exclude any location we don't have BAF for
    hSNPs = tmp[!is.na(tmp$BAF) & !is.na(tmp$logR)]
    hSNPs = hSNPs[seqnames(hSNPs) %in% tgtChrs]
    message(sprintf("Getting allele counts in sc data"))
    #Lift them over
    snps = hSNPs
    if(!is.na(liftOverChain)){
      ch = rtracklayer::import.chain(liftOverChain)
      snps = unlist(rtracklayer::liftOver(snps,ch))
    }
    if(scUsesChr){
      tmp = renameSeqlevels(snps,setNames(paste0('chr',seqlevels(snps)),seqlevels(snps)))
    }else{
      tmp=snps
    }
    phCnts = ph$phaseExpression(hSNPs = tmp,
                                bams = scBAMs,
                                ref = refRNA,
                                gtf = GTF,
                                labels = paste0(basename(outBase),'_RNA_',gsub('\\.bam$','',basename(scBAMs))),
                                skipIfExists=skipIfExists,
                                outDir = dirname(outBase),
                                matAlleleCol = 'REF',
                                patAlleleCol = 'ALT',
                                postChrStatus=FALSE,
                                nParallel = nParallel)
    #Get the BAF for each location
    m = match(phCnts,snps)
    phCnts$BAF = ifelse(snps$alleleB=='1',snps$BAF,1-snps$BAF)[m]
    phCnts$GT = snps$GT[m]
    phCnts$GQ = snps$GQ[m]
    phCnts$mark = as.character(phCnts)
    #Fix up cellID
    phCnts$cellID = gsub(paste0(basename(outBase),'_RNA_'),'',phCnts$cellID)
    phCnts$cellID = gsub('___','_',phCnts$cellID)
    #Remove 'chr' if present
    if('chr' %in% seqlevels(phCnts))
      phCnts = renameSeqlevels(phCnts,setNames(gsub('chr','',seqlevels(phCnts)),seqlevels(phCnts)))
    #Get just the neutral cells for determining which SNPs are het
    tmp = phCnts
    if(!is.null(neutCells))
      tmp = tmp[tmp$cellID %in% neutCells]
    #Do a sanity check that the conversion has worked
    tt = aggregate(cbind(altCount,refCount,coverage) ~ cut(BAF,seq(0,1,0.1)),data=mcols(tmp),FUN=sum)
    colnames(tt) = c('BAF','alt','ref','cov')
    tt$alt=round(tt$alt/tt$cov*100,1)
    tt$ref=round(tt$ref/tt$cov*100,1)
    plot(tt$BAF,tt$alt,
         las=2,
         xlab='BAF',
         ylab='%AltInSC')
    #Aggregate to the SNP level
    tmp = aggregate(cbind(refCount,altCount,coverage) ~ mark,FUN=sum,data=mcols(tmp))
    #Combine info
    tmp = cbind(tmp,mcols(phCnts[match(tmp$mark,phCnts$mark)])[,c('BAF','GT','GQ')])
    #Get evidence of ALT
    tmp$pAlt = p.adjust(pbinom(tmp$altCount-1,tmp$coverage,errRate,lower.tail=FALSE),method='BH')
    tmp$pRef = p.adjust(pbinom(tmp$refCount-1,tmp$coverage,errRate,lower.tail=FALSE),method='BH')
    tmp$isHet = (tmp$pAlt<0.05 & tmp$pRef<0.05) |  #Evidence of both from the scRNA
                (tmp$pAlt<0.05 & grepl('0',tmp$GT) & tmp$GQ>5) | #Evidence of REF from array and ALT from scRNA
                (tmp$pRef<0.05 & grepl('1',tmp$GT) & tmp$GQ>5)   #Evidence of ALT from array and REF from scRNA
    #Now we can finally declare which are heterozygous
    hetCalls = tmp
    #Label the ones in CN regions
    tmp$fCN = 0.5
    tmp$isCN = FALSE
    oo = findOverlaps(phCnts[match(tmp$mark,phCnts$mark)],segs)
    tmp$fCN[queryHits(oo)]=(segs$minCN/segs$totCN)[subjectHits(oo)]
    tmp$isCN[queryHits(oo)]=TRUE
    #Two possibilities, minor allele is REF or ALT.  Calaculate posterior probability of both.
    #If minor is ALT, the only counts should be from REF
    tmp$minorIsAlt = dnorm(tmp$BAF,tmp$fCN,0.02)
    tmp$minorIsRef = dnorm(tmp$BAF,1-tmp$fCN,0.02)
    normFac = tmp$minorIsAlt+tmp$minorIsRef
    tmp$minorIsAlt = tmp$minorIsAlt/normFac
    tmp$minorIsRef = tmp$minorIsRef/normFac
    tmp$pos = as.numeric(gsub('.*:','',tmp$mark))
    tmp$chr = gsub(':.*','',tmp$mark)
    #Now we can adjust the "patCount" column to give the count for the minor allele
    m = match(phCnts$mark,tmp$mark)
    phCnts$patCount = ifelse(tmp$minorIsAlt[m]>0.99,
                             phCnts$altCount,
                             ifelse(tmp$minorIsRef[m]>0.99,
                                    phCnts$refCount,
                                    NA
                                    )
                             )
    phCnts$matCount = ifelse(tmp$minorIsAlt[m]>0.99,
                             phCnts$refCount,
                             ifelse(tmp$minorIsRef[m]>0.99,
                                    phCnts$altCount,
                                    NA
                                    )
                             )
    #Make a schematic plot
    w=which(tmp$chr=='1')
    cc = ifelse(tmp$isHet[w],
                'Red',
                '#00000033'
                )
    plot(tmp$pos[w],tmp$BAF[w],
         frame.plot=FALSE,
         xlab='Position',
         ylab='BAF',
         main='Chr1',
         las=1,
         col=cc,
         pch=19,
         cex=0.1
         )
    legend(x='topright',
           legend=c('Hom','Het'),
           col=c('#00000033','red'),
           bty='n',
           pch=19)
    #Drop the ones that aren't HET
    phCnts = phCnts[which(tmp$isHet[m])]
  }else{
    ###################
    # Get all het SNPs
    message(sprintf("Finding heterozygous SNPs"))
    noNorm = missing(Normal) || is.null(Normal)
    if(noNorm){
      #Assume a certain amount of normal contamination and call Het from Tumour
      hSNPs = ph$findHetSNPs(BAM=Tumour,
                             ref=refDNA,
                             VCF_path = paste0(outBase,'_DNA_hetSNPs.vcf.gz'),
                             skipIfExists=skipIfExists,
                             BAF_lim=c(0.05,0.95),
                             nParallel=nParallel)
    }else{
      hSNPs = ph$findHetSNPs(BAM=Normal,
                             ref=refDNA,
                             VCF_path = paste0(outBase,'_DNA_hetSNPs.vcf.gz'),
                             skipIfExists=skipIfExists,
                             nParallel=nParallel)
    }
    #Strip 'chr' if it's there
    if('chr' %in% seqlevels(hSNPs))
      hSNPs = renameSeqlevels(hSNPs,setNames(gsub('chr','',seqlevels(hSNPs)),seqlevels(hSNPs)))
    ############################
    # Phase SNPs in CN segments
    message(sprintf("Phasing SNPs"))
    #Do we need to add 'chr' to get this to work?
    if(tumUsesChr){
      tmp = renameSeqlevels(hSNPs,setNames(paste0('chr',seqlevels(hSNPs)),seqlevels(hSNPs)))
    }else{
      tmp = hSNPs
    }
    phasedSNPs = ph$phaseUsingCN(tmp,
                                 tumBAM=Tumour,
                                 segs=segs,
                                 lab= paste0(outBase,'_tumourDNA'),
                                 ref=refDNA,
                                 skipIfExists=skipIfExists,
                                 nParallel = nParallel)
    if('chr' %in% seqlevels(phasedSNPs))
      phasedSNPs = renameSeqlevels(phasedSNPs,setNames(gsub('chr','',seqlevels(phasedSNPs)),seqlevels(phasedSNPs)))
    #########################
    # Get counts in 10X data
    message(sprintf("Getting allele counts in sc data"))
    #Lift them over
    ch = rtracklayer::import.chain(liftOverChain)
    snps = unlist(rtracklayer::liftOver(phasedSNPs,ch))
    #Get counts at each allele in the data
    snps = subsetByOverlaps(snps,segs,ignore.strand=TRUE)
    if(scUsesChr){
      tmp = renameSeqlevels(snps,setNames(paste0('chr',seqlevels(snps)),seqlevels(snps)))
    }else{
      tmp=snps
    }
    phCnts = ph$phaseExpression(hSNPs = tmp,
                                bams = scBAMs,
                                ref = refRNA,
                                gtf = GTF,
                                labels = paste0(basename(outBase),'_RNA_',gsub('\\.bam$','',basename(scBAMs))),
                                skipIfExists=skipIfExists,
                                outDir = dirname(outBase),
                                matAlleleCol = 'matAllele',
                                patAlleleCol = 'patAllele',
                                nParallel = nParallel)
    ####Work out the CN changes
    ###lab='/tmp/asoteuh_c'
    ###output = paste0(lab,'_alleleCounts.tsv')
    ###tgtLoci = paste0(lab,'_hetSNPs.tsv')
    ###write.table(cbind(as.character(seqnames(snps_DNA)),start(snps_DNA)),tgtLoci,row.names=FALSE,sep='\t',col.names=FALSE,quote=FALSE)
    ####Now run allele counter with things sensible for bulk DNA.  
    ###ph$parallelAlleleCounter(tgtLoci,output,Tumour,x=FALSE,f=3,F=3852,m=20,q=35,r=refDNA,skipIfExists=FALSE,d=TRUE,nParallel=30)
    ###cnts = read.table(output,sep='\t',header=TRUE,comment.char='')
    ###tmp = cnts[,3:6]
    ###colnames(tmp) = gsub('Count_','',colnames(tmp))
    ###bases = colnames(tmp)
    ###cnts = GRanges(cnts[,1],IRanges(cnts[,2],width=1))
    ###mm = match(hSNPs,cnts)
    ###mcols(hSNPs) = cbind(mcols(hSNPs),tmp[mm,])
    ###mcols(hSNPs)[,sprintf('cntsTumRef')] = as.matrix(tmp[mm,])[cbind(row=seq_len(length(hSNPs)),col=match(hSNPs$REF,bases))]
    ###mcols(hSNPs)[,sprintf('cntsTumAlt')] = as.matrix(tmp[mm,])[cbind(row=seq_len(length(hSNPs)),col=match(hSNPs$ALT,bases))]
    ###mcols(hSNPs)[,sprintf('cntsTumDepth')] = rowSums(tmp[mm,])
    ###mcols(hSNPs)[,sprintf('tumBAF')] = mcols(hSNPs)[,'cntsTumAlt'] / mcols(hSNPs)[,'cntsTumDepth']
    ###mcols(hSNPs)[,sprintf('tumAAF')] = mcols(hSNPs)[,'cntsTumRef'] / mcols(hSNPs)[,'cntsTumDepth']
    #Fix up cellID
    phCnts$cellID = gsub(paste0(basename(outBase),'_RNA_'),'',phCnts$cellID)
    phCnts$cellID = gsub('___','_',phCnts$cellID)
    #Remove 'chr' if present
    if('chr' %in% seqlevels(phCnts))
      phCnts = renameSeqlevels(phCnts,setNames(gsub('chr','',seqlevels(phCnts)),seqlevels(phCnts)))
    #If we don't have normal, need to do post-filtering to get the ones that are heterozygous
    if(is.null(Normal)){
    }
  }
  ##################
  # Sanitise counts
  message(sprintf("Sanitising counts..."))
  #Drop ones where we have no phasing
  goodCnts = phCnts[!is.na(phCnts$patCount),]
  #Drop those not on the whitelist
  goodCnts = goodCnts[goodCnts$cellID %in% cellWhitelist]
  #And those that are not exonic
  goodCnts = goodCnts[goodCnts$regionType %in% regionsToUse]
  #Give each SNP a "gene" label.  If inter-geneic, each SNP a separate "gene"
  goodCnts$geneLab = gsub(' ','',gsub('\\)','',gsub('\\"','',gsub('c\\(','',as.character(goodCnts$geneName)))))
  naGenes = unique(as.character(goodCnts)[goodCnts$geneLab=='NA'])
  naGenes = setNames(paste0('genelessSNP',seq_along(naGenes)),naGenes)
  w = which(goodCnts$geneLab=='NA')
  goodCnts$geneLab[w] = naGenes[as.character(goodCnts)[w]]
  if(length(goodCnts)==0){
    stop("No counts found for cells on whitelist")
  }
 ##########################
  # Get the over-dispersion
  #The maximum over-dispersion is...
  cellSum = aggregate(cbind(patCount,coverage) ~ cellID,data=mcols(goodCnts),FUN=sum)
  fit = suppressWarnings(vglm(cbind(patCount,coverage-patCount) ~ 1,betabinomial,data=cellSum,trace=FALSE))
  maxOD = 1/(1+exp(-coef(fit)[2]))
  if(is.null(odFiat)){
    #Can we use a guess at which cells are neutral to guess it?
    if(!is.null(neutCells)){
      #Get just the neutral cnts
      neutCnts = goodCnts[gsub(paste0(basename(outBase),'_RNA_'),'',goodCnts$cellID) %in% neutCells]
      cellSum = aggregate(cbind(patCount,coverage) ~ cellID,data=mcols(neutCnts),FUN=sum)
      fit = suppressWarnings(vglm(cbind(patCount,coverage-patCount) ~ 1,betabinomial,data=cellSum,trace=FALSE))
      od = 1/(1+exp(-coef(fit)[2]))
    }else{
      #If not, let's just set it to the maximum
      od=maxOD
    }
  }else{
    #It's being fixed from on high.  Make sure it's sane
    if(odFiat>maxOD)
      stop("Provided over-dispersion exceeds maximum from data!")
    od=odFiat
  }
  message(sprintf('Over-dispersion set to %g',od))
  outProbs = list()
  toDrop = c()
  for(chr in unique(as.character(seqnames(segs)))){
    message(sprintf("Procesing chromosome %s",chr))
    regions = segs[as.character(seqnames(segs))==chr]
    regions$fCN = regions$minCN/regions$totCN
    #Get the different configs
    #configs = paste0(regions$minCN,'___',regions$totCN)
    #At the moment, the only thing that matters is the ratio
    for(fCN in unique(regions$fCN)){
      message(sprintf("Processing regions with allele ratio %g",fCN))
      ###############
      # Loading data
      #Assume they're all the one allele then
      tgt = regions[regions$fCN==fCN]
      subCnts = subsetByOverlaps(goodCnts,tgt,ignore.strand=FALSE)
      if(length(subCnts)==0)
        next
      #Construct the r object
      jj = match(subCnts$cellID,cellWhitelist)
      ii = match(subCnts,snps)
      w = which(!is.na(jj))
      r = sparseMatrix(i=ii[w],
                       j=jj[w],
                       x = subCnts$patCount[w],
                       dim=c(length(snps),length(cellWhitelist)),
                       dimnames=list(as.character(snps),cellWhitelist))
      #And the coverage
      nCov = sparseMatrix(i=ii[w],
                       j=jj[w],
                       x = subCnts$coverage[w],
                       dim=c(length(snps),length(cellWhitelist)),
                       dimnames=list(as.character(snps),cellWhitelist))
      #Keep only those that have coverage
      w = which(rowSums(nCov)>0)
      r=r[w,,drop=FALSE]
      nCov=nCov[w,,drop=FALSE]
      #Get the mapping between snps and genes
      m = match(rownames(r),as.character(subCnts))
      geneMap = setNames(subCnts$geneLab[m],rownames(r))
      geneToSNP = split(names(geneMap),geneMap)
      #Now store some more stuff
      Ij = unlist(lapply(geneToSNP,length))
      numGenes = length(Ij)
      numCells = ncol(r)
      maxNumSNPs = max(Ij)
      #Make the data in long format
      dd = mcols(subCnts)
      dd$snpID = as.character(subCnts)
      colnames(dd) = gsub('patCount','obs',colnames(dd))
      colnames(dd) = gsub('coverage','tot',colnames(dd))
      dd$pp = dd$obs/dd$tot
      dd$pCN = ifelse(is.na(dd$pp),
                      fCN,
                      ifelse(dd$pp <= fCN,
                             fCN, #It's below the error range
                             ifelse(dd$pp > errRate+(1-2*errRate)*fCN,
                                    errRate+(1-2*errRate)*fCN, #It's above the error range
                                    dd$pp #It's inside the error range
                                    )
                             )
                      )
      dd$pNeut = 0.5
      #Diagnostic plot
      x=colSums(r)
      y=colSums(nCov)
      plot(y,x/y)
      abline(h=c(0.5,fCN),col=c(1,2))
      ##########
      # Fitting
      #Find genes to exclude.  Anything that is above can be tested globally
      #Actually this is a dumb idea as it systematically favours excluding SNPs that are high
      #geneSum = aggregate(cbind(obs,tot) ~ geneLab,data=dd,FUN=sum)
      #qVals = p.adjust(1-pbetabinom(geneSum$obs-1,geneSum$tot,0.5,0),method='BH')
      ##plot(geneSum$tot,geneSum$obs/geneSum$tot,
      ##     col=ifelse(qVals<0.01,'red','black'),
      ##     log='x')
      ##abline(h=0.5)
      #message(sprintf("Removing %d genes with evidence of strong ASE.",sum(qVals<0.01)))
      #toDrop = c(toDrop,geneSum$geneLab[qVals<0.01])
      #Start with no over-dispersion
      postCN = function(dd,od,toDrop){
        toUse = dd[!dd$geneLab %in% toDrop,]
        toUse$llCN = dbetabinom(toUse$obs,toUse$tot,prob=toUse$pCN,rho=od+1e-6,log=TRUE)
        toUse$llNeut = dbetabinom(toUse$obs,toUse$tot,prob=toUse$pNeut,rho=od+1e-6,log=TRUE)
        #Get summary probabilities
        tmp = aggregate(cbind(llCN,llNeut) ~ cellID,data=toUse,FUN=sum)
        #Make a table with the likelihoods and the posterior probs
        out = data.frame(cellID = cellWhitelist,
                         llCN = 0,
                         llNeut = 0,
                         postProb = 0)
        m = match(tmp$cellID,out$cellID)
        out$llCN[m] = tmp[,2]
        out$llNeut[m] = tmp[,3]
        #Use difference to avoid rounding errors
        out$postProb = exp(out$llCN-out$llNeut)
        out$postProb = out$postProb/(out$postProb+1)
        return(out)
      }
      #Iteratively find drop genes and fit things
      while(TRUE){
        #First find neutral cells
        post = postCN(dd,od,toDrop)
        #post = postCN(dd,od,c())
        neutCells = post$cellID[post$postProb<0.05]
        if(length(neutCells)==0)
          break
        #Find any genes that look like they have evidence of ASE
        geneSum = aggregate(cbind(obs,tot) ~ geneLab,data=dd[dd$cellID %in% neutCells & !(dd$geneLab %in% toDrop),],FUN=sum)
        qVals = p.adjust(pmin(pbetabinom(geneSum$obs,geneSum$tot,0.5,od),1-pbetabinom(geneSum$obs-1,geneSum$tot,0.5,od)),method='BH')
        w = qVals<0.01
        if(sum(w)==0)
          break
        message(sprintf("Removing %d genes with evidence of strong ASE.",sum(w)))
        #plot(geneSum$tot,geneSum$obs/geneSum$tot,
        #     col=ifelse(w,'red','black'),
        #     log='x')
        #abline(h=0.5)
        toDrop = unique(c(toDrop,geneSum$geneLab[w]))
      }
      #Save the result
      post$chr=chr
      post$fCN=fCN
      outProbs[[length(outProbs)+1]]=post
    }
  }
  outAll= do.call(rbind,outProbs)
  out = aggregate(cbind(llCN,llNeut) ~ cellID,data=outAll,FUN=sum)
  out$post = out$llCN-out$llNeut
  out$post = exp(out$post)/(1+exp(out$post))
  ############################
  # Make a demonstration plot
  #Designed to run on GOSH021
  #tt = aggregate(cbind(llCN,llNeut) ~ cellID,data=outAll[outAll$fCN==0 & outAll$chr=='1',],FUN=sum)
  #tt2 = aggregate(cbind(matCount,patCount,coverage) ~ cellID,data=subsetByOverlaps(goodCnts,segs[segs$minCN==0 & as.character(seqnames(segs))=='1']),FUN=sum,na.rm=TRUE)
  #m = match(tt$cellID,tt2$cellID)
  #tt$patCount = tt2$patCount[m]
  #tt$matCount = tt2$matCount[m]
  #tt$coverage = tt2$coverage[m]
  #tt$BAF = tt$patCount/(tt$patCount+tt$matCount)
  #tt$ll = tt$llCN-tt$llNeut
  #tt$pos = tt$llCN-tt$llNeut
  #tt$pos = 1/(1+exp(-out$pos))
  #cc = 'OrRd'
  #cc = suppressWarnings(brewer.pal(100,cc))
  #cFun = circlize::colorRamp2(seq(0,1,length.out=length(cc)),cc)
  #plot(tt$coverage,
  #     tt$BAF,
  #     las=1,
  #     frame.plot=FALSE,
  #     xlab='scCoverage',
  #     ylab='fracLostAllele',
  #     col='black',
  #     bg=cFun(tt$ll),
  #     pch=21,
  #     log='x'
  #     )
  #pl$addColorBar('topright',
  #               col=cc,
  #               breaks=seq(0,1,length.out=length(cc)),
  #               title='Prob\nTumour',
  #               ticks=c(0,1),
  #               )
  srat@meta.data = cbind(srat@meta.data,srat@reductions$umap@cell.embeddings)
  srat@meta.data[gsub('___','_',out$cellID),'post']=out$post
  srat@meta.data$call = ifelse(srat@meta.data$post>0.99,
                               'Tumour',
                               ifelse(srat@meta.data$post<0.01,
                                      'Normal',
                                      '???'
                                      )
                               )
  #Add calls to everything
  outAll$lab = sprintf('%s:%.02g',outAll$chr,outAll$fCN)
  segs$lab = sprintf('%s:%.02g',as.character(seqnames(segs)),segs$minCN/segs$totCN)
  for(lab in unique(outAll$lab)){
    tmp = outAll[outAll$lab==lab,]
    tmp$ll = tmp$llCN-tmp$llNeut
    srat@meta.data[,lab] = tmp$ll[match(rownames(srat@meta.data),tmp$cellID)]
  }
  #Add posterior prob to counts
  goodCnts$postTumProb = srat@meta.data[goodCnts$cellID,'post']
  #Add the marker of whether it was dropped for using ASE
  goodCnts$strongASE = goodCnts$geneLab %in% toDrop
  sumStats=NULL
  #Calculate some summary statistics
  #For each cell, how many informative, genes, SNPs, reads
  sumStats = data.frame(cellID = cellWhitelist,
                        genes = 0,
                        SNPs = 0,
                        reads = 0,
                        lostGenes = 0,
                        lostSNPs = 0,
                        lostReads = 0,
                        exonFrac = 0,
                        intronFrac = 0,
                        interFrac = 0,
                        row.names=cellWhitelist
                        )
  x = aggregate(geneLab ~ cellID,FUN=function(e) length(unique(e)),data=goodCnts[!goodCnts$strongASE])
  sumStats[x$cellID,'genes']=x$geneLab
  x = aggregate(coverage ~ cellID,FUN=length,data=goodCnts[!goodCnts$strongASE])
  sumStats[x$cellID,'SNPs']=x$coverage
  x = aggregate(coverage ~ cellID,FUN=sum,data=goodCnts[!goodCnts$strongASE])
  sumStats[x$cellID,'reads']=x$coverage
  if(!is.null(toDrop)){
    x = aggregate(geneLab ~ cellID,FUN=function(e) length(unique(e)),data=goodCnts[goodCnts$strongASE])
    sumStats[x$cellID,'lostGenes']=x$geneLab
    x = aggregate(coverage ~ cellID,FUN=length,data=goodCnts[goodCnts$strongASE])
    sumStats[x$cellID,'lostSNPs']=x$coverage
    x = aggregate(coverage ~ cellID,FUN=sum,data=goodCnts[goodCnts$strongASE])
    sumStats[x$cellID,'lostReads']=x$coverage
  }
  if(any('Exonic' %in% goodCnts$regionType)){
    x = aggregate(coverage ~ cellID,FUN=sum,data=goodCnts[!goodCnts$strongASE & goodCnts$regionType=='Exonic'])
    sumStats[x$cellID,'exonFrac']=x$coverage/sumStats[x$cellID,'reads']
  }  
  if(any('Intronic' %in% goodCnts$regionType)){
    x = aggregate(coverage ~ cellID,FUN=sum,data=goodCnts[!goodCnts$strongASE & goodCnts$regionType=='Intronic'])
    sumStats[x$cellID,'intronFrac']=x$coverage/sumStats[x$cellID,'reads']
  }  
  if(any('InterGenic' %in% goodCnts$regionType)){
    x = aggregate(coverage ~ cellID,FUN=sum,data=goodCnts[!goodCnts$strongASE & goodCnts$regionType=='InterGenic'])
    sumStats[x$cellID,'interFrac']=x$coverage/sumStats[x$cellID,'reads']
  }  
  sumStats$readsPerMB = sumStats$reads/sum(width(segs)/1e6)
  return(list(out=out,srat=srat,cnts=goodCnts,segs=segs,outAll=outAll,snps=snps,toDrop=toDrop,sumStats=sumStats))
}

#' Plots the evidence for one cell in detail.
#'
#' Plots summary of evidence for an individual cell.  SNPs are ordered such that those that provide the most weight are on the right.  Shape shows if the SNP is in an intron,exon, or intergenic.  The red/black line shows the tumour/normal BAF for each chrommosome.  The dots and error bars show the estimate and confidence interval for the BAF of this SNP aggregating across cells marked as normal.  The dotted line shows the cumulative BAF aggregating across SNPs from the left.
#'
#' @param fit Output of callTumourNormal.
#' @param cellID The ID of the cell to plot evidence for.
#' @param chr Plot only the evidence from these chromosomes.  If NULL, does all.
#' @param ncol Number of columns to use in array of chromosomes.
#' @param nrow Number of columns to use in array of chromosomes.
#' @param colMarks Shape points by either allele specific expression or Ref/Alt.
plotCell = function(fit,cellID,chr=NULL,ncol=NULL,nrow=NULL,colMarks=c('ASE','REF')){
  colMarks = match.arg(colMarks)
  cnts = fit$cnts
  cnts$patIsRef = (fit$snps$patAllele==fit$snps$REF)[match(cnts,fit$snps)]
  #Decide what we'll colour
  if(colMarks=='ASE'){
    cnts$toCol = cnts$strongASE
  }else if(colMarks=='REF'){
    cnts$toCol = cnts$patIsRef
  }
  if(!cellID %in% cnts$cellID)
    stop("Invalide cell ID")
  #Loop over segments
  segs = fit$segs
  segs$fCN = segs$minCN/segs$totCN
  #Split them into groups
  s = split(segs,segs$lab)
  s = s[lengths(s)>0]
  #Include only groups that have counts for our cell
  xx = cnts[cnts$cellID==cellID]
  s = s[sapply(s,function(e) length(findOverlaps(e,xx))>0)]
  #And include only groups we have selected
  if(!is.null(chr))
    s = s[sapply(s,function(e) as.character(seqnames(e)) %in% chr)]
  #Decide how to plot things
  nPlots = length(s)
  if(is.null(ncol) & is.null(nrow)){
    #Go with 4/3 ratio
    r=4/3
    ncol = sqrt(nPlots*r)
    nrow = sqrt(nPlots/r)
    #Now we need to make an integer...
    ncol = ceiling(ncol)
    nrow = ceiling(nPlots/ncol)
  }
  if(is.null(ncol))
    nrow = ceiling(nPlots/ncol)
  if(is.null(nrow))
    ncol = ceiling(nPlots/nrow)
  if(nrow==1)
    ncol=nPlots
  if(ncol==1)
    nrow=nPlots
  par(mfrow=c(nrow,ncol))
  for(ii in seq_along(s)){
    tgt = s[[ii]]
    subCnts = subsetByOverlaps(cnts,tgt)
    fCN=(tgt$minCN/tgt$totCN)[1]
    #Make the label
    lab=sprintf("%s\n%s p=%.02g ",
                cellID,
                tgt$lab[1],
                fit$outAll$postProb[fit$outAll$cellID==cellID & fit$outAll$lab==tgt$lab[1]])
    #Get the local values
    xx = subCnts[subCnts$cellID==cellID]
    alt =xx$patCount
    tot = xx$coverage
    #Aggregate across all normal cells
    tmp = subCnts[as.character(subCnts) %in% as.character(xx)]
    tmp$adjAlt = (1-tmp$postTumProb)*tmp$patCount
    tmp$adjTot = (1-tmp$postTumProb)*tmp$coverage
    tmp$mark = as.character(tmp)
    normRange = aggregate(cbind(adjAlt,adjTot) ~ mark,data=mcols(tmp),FUN=sum)
    normRange$low = ifelse(normRange$adjAlt==0,0,qbeta(.025,normRange$adjAlt,normRange$adjTot-normRange$adjAlt+1))
    normRange$high = ifelse(normRange$adjAlt==normRange$adjTot,1,qbeta(.975,normRange$adjAlt+1,normRange$adjTot-normRange$adjAlt))
    normRange$est = normRange$adjAlt/normRange$adjTot
    #Make this into a data-frame
    dd = data.frame(alt=alt,tot=tot,type=xx$regionType,row.names=as.character(xx))
    dd$ratio = dd$alt/dd$tot
    dd$totGlob = normRange$adjTot[match(rownames(dd),normRange$mark)]
    dd$toCol = xx$toCol
    #Define the order
    dd = dd[order(dd$tot,dd$totGlob),]
    #Cumulative contribution to estimate
    dd$x = cumsum(dd$tot/sum(dd$tot))
    #Make the plot
    plot(0,0,
         type='n',
         xlim=c(0,1),
         ylim=c(0,1),
         xaxt='n',
         xlab='',
         ylab='BAF',
         las=2,
         main=lab,
         frame.plot=FALSE)
    #Normal fraction
    abline(h=0.5)
    #Put box and whisker range for normals (assume binomial)
    lwd = .01
    m = match(rownames(dd),normRange$mark)
    #Top
    segments(dd$x-lwd/2,normRange$high[m],dd$x+lwd/2,normRange$high[m],col='darkblue')
    #Join
    segments(dd$x,normRange$high[m],dd$x,normRange$low[m],col='darkblue',lwd=0.3)
    #Bottom
    segments(dd$x-lwd/2,normRange$low[m],dd$x+lwd/2,normRange$low[m],col='darkblue')
    points(dd$x,normRange$est[m],pch=18,cex=1.0,col='darkblue')
    #Tumour fraction
    abline(h=fCN,col='darkred')
    #The observations for this cell
    #segments(dd$x,0.5,dd$x,dd$ratio,lwd=dd$tot/sum(dd$tot)*5)
    pchVal = as.numeric(factor(dd$type,levels=c('Exonic','Intronic','InterGenic')))
    points(dd$x,dd$ratio,pch=pchVal,cex=2,col=ifelse(dd$toCol,'Orange','Black'))
    #The progressive estimate
    lines(dd$x,cumsum(dd$alt)/cumsum(dd$tot),lty=2,col='darkgreen')
  }
}

#' Plots frequency of minor allele across all SNPs and cells
#'
#' Make a plot summarising the evidence for all cells across all chromosomes.  SNPs that contribute little information are not shown (as determined by minCells).
#'
#' @param fit Output of callTumourNormal.
#' @param chr Which Chromosomes to include.  Null includes all.
#' @param minCells Don't show a SNP unless it has coverage in at least this fraction of cells.
#' @param colPalMain What colour scheme to use for main heatmap.
#' @param colPalAnn What colour scheme to use for track showing Tumour probability.
plotAllData = function(fit,chr=NULL,minCells=0.01,colPalMain='RdBu',colPalAnn='Greys'){
  cnts = fit$cnts
  if(!is.null(chr))
    cnts = cnts[as.character(seqnames(cnts)) %in% chr]
  #Calculate the main things
  cnts$mark = as.character(cnts)
  cnts$ratio = cnts$patCount/cnts$coverage
  #Get the total counts in each SNP
  snpCnts = sapply(split(cnts$coverage,as.character(cnts)),sum)
  #Turn into a matrix.
  marks = unique(cnts$mark)
  cells = unique(cnts$cellID)
  mat = matrix(NA,
               nrow=length(cells),
               ncol=length(marks),
               dimnames = list(cells,marks))
  mat[cbind(match(cnts$cellID,cells),match(cnts$mark,marks))] = cnts$ratio
  #Drop columns with not many entries
  mat = mat[,colSums(!is.na(mat))/ncol(mat)>=minCells]
  #Order by posterior probability
  pp = fit$out$post[match(rownames(mat),fit$out$cellID)]
  mat = mat[order(pp),]
  pp = fit$out$post[match(rownames(mat),fit$out$cellID)]
  colFun = suppressWarnings(brewer.pal(100,colPalAnn))
  colFun = circlize::colorRamp2(seq(0,1,length.out=length(colFun)),colFun)
  leftAnn = HeatmapAnnotation(probTum=pp,which='row',col=list(probTum=colFun))
  #And by the total counts
  mat = mat[,order(as.character(seqnames(cnts))[match(colnames(mat),as.character(cnts))],snpCnts[colnames(mat)])]
  #Define colour scheme
  colFun = suppressWarnings(brewer.pal(100,colPalMain))
  colFun = circlize::colorRamp2(seq(0,1,length.out=length(colFun)),colFun)
  hm = Heatmap(mat,
               col=colFun,
               name='BAF',
               show_row_names=FALSE,
               show_column_names=FALSE,
               cluster_rows=FALSE,
               cluster_columns=FALSE,
               left_annotation=leftAnn,
               column_split = as.character(seqnames(cnts))[match(colnames(mat),as.character(cnts))]
               #row_split = ifelse(pp>0.99,'Tumour',ifelse(pp<0.01,'Normal','???')),
               #column_split=cnts$geneLab[match(colnames(mat),as.character(cnts))],
               #row_order = pp
               )
  return(hm)
}


#' Plots the posterior probability by segment
#'
#' Plot the posterior probability of each cell being tumour independently for each chromosome.  Usefull for finding sub-clonal populations.
#' 
#' @param fit Output of callTumourNormal.
#' @param chr Chromosomes to show, NULL shows all.
#' @param zCut Don't show probabilities above this threshold (logit scale).
#' @param colPal Colour scheme to use.
#' @param linearScole Plot on linear scale (probabilities) rather than logistic scale.
plotBySegment = function(fit,chr=NULL,zCut=5,colPal='BrBG',linearScale=FALSE){
  dat = fit$outAll
  if(!is.null(chr))
    dat = dat[dat$chr %in% chr,]
  cells = unique(dat$cellID)
  legs = unique(dat$lab)
  mat = matrix(0,
               nrow=length(cells),
               ncol=length(legs),
               dimnames=list(cells,legs))
  mat[cbind(match(dat$cellID,cells),match(dat$lab,legs))]=dat$llCN-dat$llNeut
  if(linearScale)
    mat = exp(mat)/(1+exp(mat))
  colFun = suppressWarnings(brewer.pal(100,colPal))
  zLims = c(-zCut,zCut)
  if(linearScale)
    zLims = exp(zLims)/(1+exp(zLims))
  colFun = circlize::colorRamp2(seq(zLims[1],zLims[2],length.out=length(colFun)),colFun)
  llTot =fit$out$post[match(rownames(mat),fit$out$cellID)]
  if(!linearScale)
    llTot = log(llTot/(1-llTot))
  leftAnn = HeatmapAnnotation(totProb = llTot,
                              which='row',
                              show_legend=FALSE,
                              col=list(totProb=colFun))
  hm = Heatmap(mat,
               col=colFun,
               name=ifelse(linearScale,'Prob','LL'),
               show_row_names=FALSE,
               left_annotation = leftAnn
               )
  return(hm)
}


###############
# Load Seurat #
###############
sratRef = readRDS('Data/fAdRef.RDS')
sratPM = readRDS('Data/srat_dutch_primary.rds')
#Fix up cell names
tmp = read.table('Data/celseq2_bc384-v4.csv',sep='\t',header=FALSE)
tmp$V2 = toupper(tmp$V2)
sratPM = RenameCells(sratPM,new.names=with(sratPM@meta.data,paste0(PlateId,'_sanitised_sorted_',tmp$V2[match(gsub('.*-','',CellId),tmp$V1)])))
sratOrg = readRDS('Data/srat_dutch_organoids.rds')
sratOrg = RenameCells(sratOrg,new.names=with(sratOrg@meta.data,paste0(PlateId,'_sanitised_sorted_',tmp$V2[match(gsub('.*-','',CellId),tmp$V1)])))
sratSI = readRDS('Data/srat_inhouse.rds')


###################
# Tree of samples #
###################
srcData = list(GOSH025 = list(Normal = 'Data/BAMs/GOSH025/PD46693b.sample.dupmarked.bam',
                              Tumour = 'Data/BAMs/GOSH025/PD46693a.sample.dupmarked.bam',
                              #CN = 'Data/BAMs/GOSH025/PD46693a.battenberg.summary.csv',
                              CN = c('10,0,2','11,0,2','21,0,2','4,0,2','1,1,3','12,1,3','6,1,3','21,0,2'),
                              scBAMs = file.path('Data/BAMs/GOSH025/',c('4602STDY8004894.bam','4602STDY8004902.bam','4602STDY8004910.bam')),
                              scData = file.path('Data/scData/Neuroblastoma/',c('4602STDY8004894','4602STDY8004902','4602STDY8004910'),'outs')
                              ),
               GOSH021 = list(Normal = 'Data/BAMs/GOSH021/PD43255b.sample.dupmarked.bam',
                              Tumour = 'Data/BAMs/GOSH021/PD43255a.sample.dupmarked.bam',
                              CN = 'Data/BAMs/GOSH021/PD43255a.ascat_ngs.summary.csv',
                              scBAMs = file.path('Data/BAMs/GOSH021/',c('4602STDY7787237.bam','4602STDY7787238.bam','4602STDY7787239.bam')),
                              scData = file.path('Data/scData/Neuroblastoma/',c('4602STDY7787237','4602STDY7787238','4602STDY7787239'),'outs')
                              ),
               GOSH019 = list(Normal = 'Data/BAMs/GOSH019/PD42752b.sample.dupmarked.bam',
                              Tumour = 'Data/BAMs/GOSH019/PD42752a.sample.dupmarked.bam',
                              #CN = 'Data/BAMs/GOSH019/PD42752a.battenberg.summary.csv',
                              CN = c('11,0,2'),
                              scBAMs = file.path('Data/BAMs/GOSH019/',c('4602STDY7733084.bam','4602STDY7733085.bam','4602STDY7733086.bam')),
                              scData = file.path('Data/scData/Neuroblastoma/',c('4602STDY7733084','4602STDY7733085','4602STDY7733086'),'outs')
                              ),
               GOSH023 = list(Normal = 'Data/BAMs/GOSH023/PD42752c.sample.dupmarked.bam',
                              Tumour = 'Data/BAMs/GOSH023/PD42752d.sample.dupmarked.bam',
                              #CN = 'Data/BAMs/GOSH023/PD42752d.ascat_ngs.summary.csv',
                              CN = c('3,0,2','11,0,2','14,0,2'),
                              scBAMs = file.path('Data/BAMs/GOSH023/',c('4602STDY7843576.bam','4602STDY7843577.bam','4602STDY7843578.bam')),
                              scData = file.path('Data/scData/Neuroblastoma/',c('4602STDY7843576','4602STDY7843577','4602STDY7843578'),'outs')
                              ),
               GOSH014 = list(Normal = NULL,
                              Tumour = 'Data/BAMs/GOSH014/PD42184a.bam',
                              CN = c(),
                              scBAMs = file.path('Data/BAMs/GOSH014/',c('4602STDY7685340.bam','4602STDY7685341.bam','4602STDY7685342.bam')),
                              scData = file.path('Data/scData/Neuroblastoma/',c('4602STDY7685340','4602STDY7685341','4602STDY7685342'),'outs')
                              ),
               #GOSH037 = list('DataNotBack'),
               NB060 = list(Tumour = 'Data/BAMs/dutchTumourBAMs/CNarrays/Conv1.1/201208660036_R03C01.vcf',
                            CN=c('1:1-11e7,0,1',
                                 '3:1-5e7,0,1',
                                 '4:12.5e7-N,0,1',
                                 '5:12.5e7-N,1,3'
                                 ),
                            scBAMs = grep('_sorted.bam$',list.files('Data/BAMs/dutchTumourBAMs/RNA/Reprocessed/NB060/',full.names=TRUE),value=TRUE),
                            scUsesChr=TRUE,
                            neutCells = rownames(sratPM@meta.data)[!sratPM@meta.data$seurat_clusters %in% c(9,23,15,12)],
                            arrayMode=TRUE,
                            scData=sratPM),
               NB098 = list(Tumour = 'Data/BAMs/dutchTumourBAMs/CNarrays/Conv1.1/202250710023_R08C01.vcf',
                            CN=c('9,0,1',
                                 '2:12e7-N,1,3',
                                 '3,1,3',
                                 '5,1,3'
                                 ),
                            scBAMs = grep('_sorted.bam$',list.files('Data/BAMs/dutchTumourBAMs/RNA/Reprocessed/NB098/',full.names=TRUE),value=TRUE),
                            scUsesChr=TRUE,
                            neutCells = rownames(sratPM@meta.data)[!sratPM@meta.data$seurat_clusters %in% c(9,23,15,12)],
                            arrayMode=TRUE,
                            scData=sratPM),
               NB106 = list(Tumour = 'Data/BAMs/dutchTumourBAMs/CNarrays/Conv1.1/201874100206_R07C01.vcf',
                            CN=c('1,1,3',
                                 '2,1,3',
                                 '5,1,3',
                                 '6,1,3',
                                 '8,1,3',
                                 '11,1,3',
                                 '12,1,3',
                                 '18,1,3',
                                 '19,0,1',
                                 '20,1,3'),
                            scBAMs = grep('_sorted.bam$',list.files('Data/BAMs/dutchTumourBAMs/RNA/Reprocessed/NB106/',full.names=TRUE),value=TRUE),
                            scUsesChr=TRUE,
                            neutCells = rownames(sratPM@meta.data)[!sratPM@meta.data$seurat_clusters %in% c(9,23,15,12)],
                            arrayMode=TRUE,
                            scData=sratPM),
               NB107 = list(Tumour = 'Data/BAMs/dutchTumourBAMs/CNarrays/Conv1.1/202105120074_R08C01.vcf',
                            CN=c('2:21e7-N,0,1',
                                 '3:1-6e7,0,1',
                                 '4,1,3',
                                 '10,1,3',
                                 '11:1-7e7,0,1',
                                 '12:10e7-N,0,1',
                                 '14:7.2e7-N,0,1'),
                            scBAMs = grep('_sorted.bam$',list.files('Data/BAMs/dutchTumourBAMs/RNA/Reprocessed/NB107/',full.names=TRUE),value=TRUE),
                            scUsesChr=TRUE,
                            neutCells = rownames(sratPM@meta.data)[!sratPM@meta.data$seurat_clusters %in% c(9,23,15,12)],
                            arrayMode=TRUE,
                            scData=sratPM),
               NB123 = list(Tumour = 'Data/BAMs/dutchTumourBAMs/CNarrays/Conv1.1/201874100176_R07C01.vcf',
                            CN=c('12,1,3'),
                            scBAMs = grep('_sorted.bam$',list.files('Data/BAMs/dutchTumourBAMs/RNA/Reprocessed/NB123/',full.names=TRUE),value=TRUE),
                            scUsesChr=TRUE,
                            neutCells = rownames(sratPM@meta.data)[!sratPM@meta.data$seurat_clusters %in% c(9,23,15,12)],
                            arrayMode=TRUE,
                            scData=sratPM),
               NB125 = list(Tumour = 'Data/BAMs/dutchTumourBAMs/CNarrays/Conv1.1/202884920022_R06C01.vcf',
                            CN=c('11:1-7e7,1,3',
                                 '11:7.2e7-N,0,1',
                                 '17:5e7-N,1,3'),
                            scBAMs = grep('_sorted.bam$',list.files('Data/BAMs/dutchTumourBAMs/RNA/Reprocessed/NB125/',full.names=TRUE),value=TRUE),
                            scUsesChr=TRUE,
                            neutCells = rownames(sratPM@meta.data)[!sratPM@meta.data$seurat_clusters %in% c(9,23,15,12)],
                            arrayMode=TRUE,
                            scData=sratPM),
               NB130 = list(Tumour = 'Data/BAMs/dutchTumourBAMs/CNarrays/Conv1.1/202329370152_R07C01.vcf',
                            CN=c('2:1.6e7-7e7,1,3',
                                 '4:1-3.5e7,0,1',
                                 '4:16e7-N,0,1',
                                 '11:7.2e7-N,0,1',
                                 '12:12e7-N,1,3',
                                 '18,1,3'
                                 ),
                            scBAMs = grep('_sorted.bam$',list.files('Data/BAMs/dutchTumourBAMs/RNA/Reprocessed/NB130/',full.names=TRUE),value=TRUE),
                            scUsesChr=TRUE,
                            neutCells = rownames(sratPM@meta.data)[!sratPM@meta.data$seurat_clusters %in% c(9,23,15,12)],
                            arrayMode=TRUE,
                            scData=sratPM),
               NB124 = list(Tumour = 'Data/BAMs/dutchTumourBAMs/CNarrays/Conv1.1/202329370209_R08C01.vcf',
                            CN=c('1:1-7e7,0,1',
                                 '2:1-3e7,1,3',
                                 '6:11.2e7-15.5e7,0,1',
                                 '11:7.2e7-N,0,1',
                                 '18,1,3'
                                 ),
                            scBAMs = grep('_sorted.bam$',list.files('Data/BAMs/dutchTumourBAMs/RNA/Reprocessed/NB124/',full.names=TRUE),value=TRUE),
                            scUsesChr=TRUE,
                            neutCells = rownames(sratPM@meta.data)[!sratPM@meta.data$seurat_clusters %in% c(9,23,15,12)],
                            arrayMode=TRUE,
                            scData=sratPM),
               NB132 = list(Tumour = 'Data/BAMs/dutchTumourBAMs/CNarrays/Conv1.1/201874140217_R08C01.vcf',
                            CN=c('1:15e7-N,1,3',
                                 '2,1,3',
                                 '4,1,3',
                                 '6,1,3',
                                 '7,1,3',
                                 '9,0,1',
                                 '12:1-2.5e7,0,1',
                                 '17,1,3',
                                 '19,1,3',
                                 '20,1,3'),
                            scBAMs = grep('_sorted.bam$',list.files('Data/BAMs/dutchTumourBAMs/RNA/Reprocessed/NB132/',full.names=TRUE),value=TRUE),
                            scUsesChr=TRUE,
                            neutCells = rownames(sratPM@meta.data)[!sratPM@meta.data$seurat_clusters %in% c(9,23,15,12)],
                            arrayMode=TRUE,
                            scData=sratPM),
               NB152 = list(Tumour = 'Data/BAMs/dutchTumourBAMs/CNarrays/Conv1.1/202884920173_R08C01.vcf',
                            CN=c('1,1,3',
                                 '7,1,3',
                                 '11:7.5e7-N,0,1',
                                 '16,0,1',
                                 '17:4.5e7-N,1,3',
                                 '20,1,3'),
                            scBAMs = grep('_sorted.bam$',list.files('Data/BAMs/dutchTumourBAMs/RNA/Reprocessed/NB152/',full.names=TRUE),value=TRUE),
                            scUsesChr=TRUE,
                            neutCells = rownames(sratPM@meta.data)[!sratPM@meta.data$seurat_clusters %in% c(9,23,15,12)],
                            arrayMode=TRUE,
                            scData=sratPM),
               NB151 = list(Tumour = 'Data/BAMs/dutchTumourBAMs/CNarrays/Conv1.1/202105120136_R04C01.vcf',
                            CN = c('3:1-6e7,0,1',
                                   '7:9e7-N,1,3',
                                   '9:2.5e7-3e7,0,1',
                                   '10:1-3e7,0,1',
                                   '11:8e7-N,0,1',
                                   '12:1e8-N,1,3',
                                   '14:6.3e7-7.8e7,0,1',
                                   '17:5e7-N,1,3',
                                   '19:4.8e7-N,0,1'),
                            scBAMs = grep('_sorted.bam$',list.files('Data/BAMs/dutchTumourBAMs/RNA/Reprocessed/NB151/',full.names=TRUE),value=TRUE),
                            scUsesChr=TRUE,
                            neutCells = rownames(sratPM@meta.data)[!sratPM@meta.data$seurat_clusters %in% c(9,23,15,12)],
                            arrayMode=TRUE,
                            scData = sratPM),
               NB152Org = list(Tumour = 'Data/BAMs/dutchTumourBAMs/CNarrays/Conv1.1/202884920173_R08C01.vcf',
                            CN=c('1,1,3',
                                 '7,1,3',
                                 '11:7.5e7-N,0,1',
                                 '16,0,1',
                                 '17:4.5e7-N,1,3',
                                 '20,1,3'),
                            scBAMs = grep('_sorted.bam$',list.files('Data/BAMs/dutchTumourBAMs/RNA/Reprocessed/NB152Org/',full.names=TRUE),value=TRUE),
                            scUsesChr=TRUE,
                            #neutCells = rownames(sratOrg@meta.data)[sratOrg@meta.data$seurat_clusters %in% c(5,10,11)],
                            neutCells = NULL,
                            arrayMode=TRUE,
                            scData=sratOrg),
               NB067 = list(Tumour = 'Data/BAMs/dutchTumourBAMs/CNarrays/Conv1.1/201874140219_R08C01.vcf',
                            CN=c('1:1-6e7,0,1',
                                 '2:1-5.5e7,1,3',
                                 '10:1-N,0,1'
                                 ),
                            scBAMs = grep('_sorted.bam$',list.files('Data/BAMs/dutchTumourBAMs/RNA/Reprocessed/NB067/',full.names=TRUE),value=TRUE),
                            scUsesChr=TRUE,
                            #neutCells = rownames(sratOrg@meta.data)[sratOrg@meta.data$seurat_clusters %in% c(5,10,11)],
                            neutCells = NULL,
                            arrayMode=TRUE,
                            scData=sratOrg),
               NB098Org = list(Tumour = 'Data/BAMs/dutchTumourBAMs/CNarrays/Conv1.1/202250710023_R08C01.vcf',
                            CN=c('9,0,1',
                                 '2:12e7-N,1,3',
                                 '3,1,3',
                                 '5,1,3'
                                 ),
                            scBAMs = grep('_sorted.bam$',list.files('Data/BAMs/dutchTumourBAMs/RNA/Reprocessed/NB098Org/',full.names=TRUE),value=TRUE),
                            scUsesChr=TRUE,
                            #neutCells = rownames(sratOrg@meta.data)[sratOrg@meta.data$seurat_clusters %in% c(5,10,11)],
                            neutCells = NULL,
                            arrayMode=TRUE,
                            scData=sratOrg),
               NB039 = list(Tumour = 'Data/BAMs/dutchTumourBAMs/CNarrays/Conv1.1/201558000107_R08C01.vcf',
                            CN=c('1:1-1e8,0,1',
                                 '1:17e7-N,1,3',
                                 '4:6e7-10e7,0,1',
                                 '6:14.5e7-N,0,1',
                                 '11:9e7-N,0,1',
                                 '19:4e7-N,0,1'
                                 ),
                            scBAMs = grep('_sorted.bam$',list.files('Data/BAMs/dutchTumourBAMs/RNA/Reprocessed/NB039/',full.names=TRUE),value=TRUE),
                            scUsesChr=TRUE,
                            #neutCells = rownames(sratOrg@meta.data)[sratOrg@meta.data$seurat_clusters %in% c(5,10,11)],
                            neutCells = NULL,
                            arrayMode=TRUE,
                            scData=sratOrg)
               )


##################
# Run genotyping #
##################
fits=list()
for(nom in names(srcData)){
  #Skip NB152 organoids as we're excluding it
  if(grepl('NB152Org',nom))
    next
  #The tumour DNA sequencing contained no tumour here
  if(grepl('GOSH014',nom))
    next
  if(!grepl('GOSH',nom))
    next
  message(sprintf("Genotyping sample %s",nom))
  params = srcData[[nom]]
  params$outBase = file.path('Results/Genotyping/',nom)
  params$GTF = gtf
  params$refDNA = refDNA
  params$refRNA = ref10X
  params$liftOverChain = liftOverChain
  params$errRate = errRate
  params$regionsToUse = c('Intronic','Exonic')
  params$nParallel=nParallel
  params$skipIfExists=TRUE
  fit = do.call(callTumourNormal,params)
  fits[[nom]]=fit
}
saveRDS(fits,'Results/Genotyping/fits.RDS')

#############################
# Custom method for GOSH014 #
#############################
nom='GOSH014'
errRate = 0.01
hetCut=0.95
params = srcData[[nom]]
params$outBase = file.path('Results/Genotyping/',nom)
params$GTF = gtf
params$refDNA = refDNA
params$refRNA = ref10X
params$liftOverChain = liftOverChain
params$errRate = errRate
params$regionsToUse = c('Intronic','Exonic')
params$nParallel=nParallel
params$skipIfExists=FALSE
#Clinical report says there is a loss of 11p.  Define this region
cyto = karyoploteR:::data.cache$cytobands$hg38
cyto = cyto[as.character(seqnames(cyto))=='chr11']
cyto = cyto[grep('p',cyto$name)]
segs = GRanges(11,IRanges(1,max(end(cyto))),minor=0,total=1)
#Now do the standard clustering on it
scData=params$scData
scSRC = file.path(scData,'filtered_gene_bc_matrices','GRCh38')
names(scSRC) = names(scData)
if(is.null(names(scData)))
  names(scSRC) = basename(dirname(scData))
srat = Read10X(scSRC)
srat = CreateSeuratObject(srat)
srat = NormalizeData(srat,verbose=FALSE)
srat = FindVariableFeatures(srat,verbose=FALSE)
srat = ScaleData(srat,verbose=FALSE)
srat = RunPCA(srat,npcs = 75,approx=FALSE,verbose=FALSE)
srat = FindNeighbors(srat,dims=seq(75),verbose=FALSE)
#Make it more broad to help aggregation
srat = FindClusters(srat,verbose=FALSE,resolution=0.3)
srat = RunUMAP(srat,dims=seq(75),verbose=FALSE)
#Use ExAC SNPs in target region 
exac = readRDS('Data/ExAC_hg19.RDS')
snps = exac$snps
snps$AF = unlist(exac$info$AF)
snps$REF = as.character(snps$REF)
snps$ALT = as.character(unlist(snps$ALT))
snps = snps[nchar(snps$ALT)==1]
#Liftover to hg38
ch = rtracklayer::import.chain(params$liftOverChain)
snps = unlist(rtracklayer::liftOver(snps,ch))
#Get the ones in our target region
snps = subsetByOverlaps(snps,segs)
#Now get counts at these locations
phCnts = ph$phaseExpression(hSNPs = snps,
                           bams = params$scBAMs,
                           ref = params$refRNA,
                           gtf = params$GTF,
                           labels = paste0(basename(params$outBase),'_RNA_',gsub('\\.bam$','',basename(params$scBAMs))),
                           skipIfExists=TRUE,
                           outDir = dirname(params$outBase),
                           matAlleleCol = 'REF',
                           patAlleleCol = 'ALT',
                           postChrStatus=FALSE,
                           nParallel = nParallel)
phCnts$mark = as.character(phCnts)
phCnts$cellID = gsub('___','_',gsub('GOSH014_RNA_','',phCnts$cellID))
#Declare something heterozygous on global aggregation
gCnts = aggregate(cbind(refCount,altCount,coverage) ~ mark,FUN=sum,data=phCnts)
gCnts$tot = gCnts$refCount+gCnts$altCount
gCnts$nDiff = pmin(gCnts$refCount,gCnts$altCount)
#For each SNP calculate a probability of it being heterozygous under two models: normality and errors
gCnts$pHet = dbinom(gCnts$nDiff,gCnts$tot,0.5)
gCnts$pHom = dbinom(gCnts$nDiff,gCnts$tot,errRate)
#Normalise and decide which model works best for each
norm = gCnts$pHet+gCnts$pHom
gCnts$pHet = gCnts$pHet/norm
gCnts$pHom = gCnts$pHom/norm
#Decide which are heteroyzgous
hetSNPs = gCnts$mark[gCnts$pHet>hetCut]
#Now aggregate reads in each cluster.
nHets = list()
for(cl in levels(srat@meta.data$seurat_clusters)){
  tgts = rownames(srat@meta.data)[srat@meta.data$seurat_clusters==cl]
  cnts = aggregate(cbind(refCount,altCount,coverage) ~ mark,FUN=sum,data=phCnts[phCnts$cellID %in% tgts & phCnts$mark %in% hetSNPs])
  cnts$tot = cnts$refCount+cnts$altCount
  cnts$nDiff = pmin(cnts$refCount,cnts$altCount)
  #For each SNP calculate a probability of it being heterozygous under two models: normality and errors
  cnts$pHet = dbinom(cnts$nDiff,cnts$tot,0.5)
  cnts$pHom = dbinom(cnts$nDiff,cnts$tot,errRate)
  #Normalise and decide which model works best for each
  norm = cnts$pHet+cnts$pHom
  cnts$pHet = cnts$pHet/norm
  cnts$pHom = cnts$pHom/norm
  #Finally get an aggregate count of het SNPs across region
  nHets[[cl]] = cnts
}
#Aggregate into a final score of normal versus tumour
pTumClust = sapply(nHets,function(e) {
                    pTum = exp(sum(log(e$pHom)))
                    pNorm = exp(sum(log(e$pHet)))
                    norm = pTum+pNorm
                    pTum = pTum/norm
                    pNorm = pNorm/norm
                    return(pTum)})
fits[[nom]] = setNames(pTumClust[as.character(srat@meta.data$seurat_clusters)],rownames(srat@meta.data))
 


#######################
# Fine tuning cleanup #
#######################
srat = sratSI
srat@meta.data$call=NA
srat@meta.data$cellID = paste0('4602',rownames(srat@meta.data))
srat@meta.data$scID = NA
for(nom in names(fits)){
  if(nom=='GOSH014'){
    w = which(srat@meta.data$cellID %in% names(fits[[nom]]))
  }else{
    w = which(srat@meta.data$cellID %in% fits[[nom]]$outAll$cellID)
  }
  srat@meta.data$scID[w] = nom
}
isTumour = c()
for(nom in names(fits)){
  fit = fits[[nom]]
  if(nom=='GOSH025'){
    cut=0.99
    cut=log((1-cut)/cut)
    tmp = aggregate(llCN-llNeut~cellID,data=fit$outAll[fit$outAll$chr!='4',],FUN=sum)
    tmp$call = ifelse(tmp[,2]>cut,'Tumour',ifelse(tmp[,2] <  -cut,'Normal','???'))
    #Add a sub-call based on chr4
    tt = aggregate(llCN-llNeut~cellID,data=fit$outAll[fit$outAll$chr=='4',],FUN=sum)
    tt = tt[match(tmp$cellID,tt$cellID),]
    tmp$call = ifelse(tmp$call=='Tumour',
                      ifelse(tt[,2]>cut,
                             'SubClone',
                             ifelse(tt[,2] < -cut,
                                    'MainClone',
                                    '???'
                                    )
                             ),
                      tmp$call
                      )
    rownames(tmp) = tmp[,1]
  }else if(nom=='GOSH014'){
    tmp = data.frame(row.names=names(fit),
                     call = ifelse(fit>0.99,
                                   'Tumour',
                                   ifelse(fit<0.01,
                                          'Normal',
                                          '???'
                                          )
                                   ),
                     pTum = fit)
  }else{
    tmp = fit$srat@meta.data
  }
  m= match(gsub('^[0-9]+','',rownames(tmp)),rownames(srat@meta.data))
  srat@meta.data$call[m[!is.na(m)]] = tmp$call[!is.na(m)]
  message(nom)
  print(table(is.na(m)))
}
sratSI = srat
#Do the same for PMC data
srat = sratPM
srat@meta.data$call=NA
for(nom in names(fits)){
  fit = fits[[nom]]
  if(nom=='GOSH014')
    next
  tmp = fit$srat@meta.data
  #Work out which ones to update
  w = fit$out$cellID[fit$out$llCN!=0 | fit$out$llNeut!=0] 
  m= match(gsub('^[0-9]+','',rownames(tmp[w,])),rownames(srat@meta.data))
  srat@meta.data$call[m[!is.na(m)]] = (tmp[w,'call'])[!is.na(m)]
  message(nom)
  print(table(is.na(m)))
}
#Default the ones for which we don't have information to ambiguous
srat@meta.data$call[is.na(srat@meta.data$call)] = '???'
sratPM = srat
#Organoids
srat = sratOrg
srat@meta.data$call=NA
for(nom in names(fits)){
  fit = fits[[nom]]
  if(nom=='GOSH014')
    next
  tmp = fit$srat@meta.data
  #Work out which ones to update
  w = fit$out$cellID[fit$out$llCN!=0 | fit$out$llNeut!=0] 
  m= match(gsub('^[0-9]+','',rownames(tmp[w,])),rownames(srat@meta.data))
  srat@meta.data$call[m[!is.na(m)]] = (tmp[w,'call'])[!is.na(m)]
  message(nom)
  print(table(is.na(m)))
}
#Default the ones for which we don't have information to ambiguous
srat@meta.data$call[is.na(srat@meta.data$call)] = '???'
sratOrg = srat

############################################
# DE of samples with prognostic CN changes #
############################################
srat = sratSI
#Which cells are tumour
isTum = srat@meta.data$seurat_clusters %in% c(1,8,16,5,25) & srat@meta.data$Phase=='G1'
#GOSH021 has 1p loss and 17q gain.
hasChange = srat@meta.data$scID %in% 'GOSH021'
#Do the differential expression based on this
avg = srat@assays$RNA@data[,isTum]
avg = do.call(cbind,lapply(split(colnames(avg),hasChange[isTum]),function(e) rowMeans(avg[,e])))
avg = as.data.frame(avg)
colnames(avg) = c('noChange','loss')
avg$logFC = log10(avg$loss/avg$noChange)
library(edgeR)
dge = DGEList(srat@assays$RNA@counts[,isTum],group=ifelse(hasChange[isTum],'hasChange','noChange'))
mm = model.matrix(~group,data=dge$samples)
dge = estimateDisp(dge,mm,prior.df=0)
deGenes = glmQLFit(dge,mm)
deGenes = glmQLFTest(deGenes,coef=2)
deGenes = topTags(deGenes,n=Inf)$table
m = match(rownames(deGenes),gns$gene_name)
deGenes$chr=as.character(seqnames(gns))[m]
deGenes$TSS=gns$TSS[m]
#Work out if the DE genes are over-represented for chromafin marker genes.
ann = list(SCP=25,
           Bridge=24,
           Left=13,
           Right=c(19,20),
           Cortex=c(5,22,28,10,0,1,2,6,8,9,21),
           Mesenchyme=c(11,18,23),
           Immune=c(16,26,29,31,27),
           Blood=c(30,4,7,12),
           Vasc=c(3,15,14,17)
           )
ann = reversemapping(ann)
ann = unlist(ann)
ann = ann[as.character(sratRef@meta.data$seurat_clusters)]
marks = quickMarkers(sratRef@assays$RNA@data,ann,N=Inf)
#The precise question is, are chromaffin markers over-represented amongst the list of DE genes
de = rownames(deGenes)[deGenes$FDR<0.05]
#Are chromaffin markers over-represented?
cMarks = marks[marks$cluster=='Right',]
cc = cMarks$gene[cMarks$tfidf>1]
pVal = phyper(q=sum(cc %in% de)-1,
              m=length(cc),
              n=nrow(deGenes)-length(cc),
              k=length(de),
              lower.tail=FALSE
              )
#Plot showing the average expression change for chromaffin genes
x = avg[cc,'logFC']
plot(density(x[!is.na(x)]),
     xlab='log10(CN/WT)',
     ylab='',
     main='Chromaffin gene marker expression change',
     frame.plot=FALSE)
abline(v=0,col='red',lty=2)


###############
# Final plots #
###############
colPal = c('Normal'='#9ACE8E',
           'Tumour'='#FF0000',
           '???'='#B3B3B3')
for(nom in c('Sanger','PMC','Organoids')){
  srat = list(Sanger=sratSI,PMC=sratPM,Organoids=sratOrg)[[nom]]
  dd = cbind(srat@meta.data,srat@reductions$umap@cell.embeddings)
  dd$call[grep('Clone',dd$call)]='Tumour'
  lab = c(Sanger='Sanger',PMC='PMC\nPrimary',Organoids='PMC\nOrganoids')[nom]
  pdf(sprintf('Results/Genotyping/genotyping_umap_%s.pdf',nom),width=4,height=3,useDingbats=FALSE)
    par(mar=c(2,2.5,2,0)+0.1)
    plot(0,
         frame.plot=FALSE,
         las=1,
         xlab='',
         ylab='',
         xlim=range(dd$UMAP_1),
         ylim=range(dd$UMAP_2),
         main = lab,
         type='n')
    #Plot ambiguous ones first
    isAmb = dd$call=='???'
    points(dd$UMAP_1[isAmb],dd$UMAP_2[isAmb],
           col = colPal[dd$call[isAmb]],
           pch=19,
           cex=0.1
           )
    points(dd$UMAP_1[!isAmb],dd$UMAP_2[!isAmb],
           col = colPal[dd$call[!isAmb]],
           pch=19,
           cex=0.1
           )
    #legend(x='topright',
    #       bty='n',
    #       legend=c('Tumour','Normal','Ambiguous'),
    #       col = colPal[c('Tumour','Normal','???')],
    #       pch=19,
    #       cex=1
    #       )
  dev.off()
  pdf('Results/Genotyping/genotyping_umap_legend.pdf',width=4,height=3,useDingbats=FALSE)
    par(mar=c(2,2.5,2,0)+0.1)
    plot(0,
         frame.plot=FALSE,
         las=1,
         xlab='',
         ylab='',
         xlim=range(dd$UMAP_1),
         ylim=range(dd$UMAP_2),
         main = lab,
         type='n')
    legend(x='topright',
           bty='n',
           legend=c('Tumour','Normal','Ambiguous'),
           col = colPal[c('Tumour','Normal','???')],
           pch=19,
           cex=1
           )
  dev.off()
}
```
