## supplementary scripts for "Tumor to normal single cell mRNA comparisons reveal a pan-neuroblastoma cancer cell": scQC.R

```
#' Basic QC and filtering functions.  Depends on scrublet being installed.

#' Initial filtering and metadata
#'
#' Calculates metadata, drop cells based on QC metrics.  Assumes Seurat V3.
#'
#' @param dataDirs Data directories to load, passed to Read10X.
#' @param excludeGenes Genes that will be dropped from the count matrix.
#' @param geneSets Sets of genes to aggregate counts for and store as metadata.  Name by list element names.  Must include an entry named "mtGenes".
#' @param sPhaseGenes Genes used to identify S phase genes.  If NULL, will be loaded from Seurat.
#' @param g2mPhaseGenes Genes used to identify G2M phase genes.  If NULL, will be loaded from Seurat.
#' @param maxMT Maximum fraction of expression from Mitochondrial genes allow per cell.
#' @param minGenes Minimum number of genes for a cell to express and pass QC.
#' @param minUMIs Minimum number of UMIs for a cell to have and pass QC.
#' @param maxBadFrac Maximum fraction of cells not passing QC in a cluster before we dump the whole cluster.
#' @param scrubScoreMax Mark a cell as a doublet if its scrublet score exceeds this value, even if scrublet does not call it as such.
#' @param numPCs Number of PCs to use for the rough clustering and doublet identification.  This is just used for generating very fine grained clustering and it shouldn't matter much what you set it to.
#' @param clusteringRes Resolution for fine clustering.  Should be very large.
#' @param doPlot Make some QC summary plots.
#' @param ... Parameters passed to \code{Read10X}
#' @return A Seurat object with cells dropped that fail the QC filters.
basicQC = function(dataDirs,excludeGenes = c(),geneSets=list(mtGenes=c("MT-ND1", "MT-ND2", "MT-CO1", "MT-CO2", "MT-ATP8", "MT-ATP6","MT-CO3", "MT-ND3", "MT-ND4L", "MT-ND4", "MT-ND5", "MT-ND6","MT-CYB")),sPhaseGenes=NULL,g2mPhaseGenes=NULL,maxMT=0.2,minGenes=300,minUMIs=1000,maxBadFrac=0.5,scrubScoreMax = 0.25,numPCs=75,clusteringRes=10.0,doPlot=TRUE,...){
  if(is.null(dataDirs)){
    message("Data directories not named, generating automatic names.")
    names(dataDirs) = paste0('DataSource',seq_along(dataDirs))
  }
  srat = Read10X(dataDirs,...)
  srat = CreateSeuratObject(srat)
  if(is.null(names(geneSets))){
    warning("No names given for gene sets.  Setting to uninformative auto-generated values.")
    names(geneSets) = paste0('geneSet',seq_along(geneSets))
  }
  if(is.null(sPhaseGenes) | is.null(g2mPhaseGenes)){
    data('cc.genes.updated.2019',package='Seurat')
  }
  if(is.null(sPhaseGenes))
    sPhaseGenes = cc.genes.updated.2019$s.genes
  if(is.null(g2mPhaseGenes))
    g2mPhaseGenes = cc.genes.updated.2019$g2m.genes
  #Get extra meta-data things
  for(nom in names(geneSets)){
    tmp = geneSets[[nom]]
    tmp = tmp[tmp %in% rownames(srat@assays$RNA@data)]
    if(length(tmp)==0)
      next
    srat@meta.data[,nom] = colSums(srat@assays$RNA@counts[tmp,,drop=FALSE])/srat@meta.data$nCount_RNA
  }
  #Get cell cycle
  srat = NormalizeData(srat,verbose=FALSE)
  #The seeed parameter magic is needed because the Seurat authors are dicks
  srat = CellCycleScoring(srat, s.features=sPhaseGenes,g2m.features=g2mPhaseGenes,seed=sample(1e9,1))
  #Call doublets
  srat@meta.data$scrubScore=NA
  srat@meta.data$scrubCall=NA
  srat@meta.data$scrubSim1=NA
  srat@meta.data$scrubSim2=NA
  for(i in seq_along(dataDirs)){
    scrubScores = runScrublet(dataDirs[i],nPCs=numPCs)
    #Make barcodes like with the load function
    if(all(gsub('.*-','',scrubScores$barcode)=='1'))
      scrubScores$barcode = gsub('-1','',scrubScores$barcode)
    #Save the relevant information in meta.data
    m = match(paste0(names(dataDirs[i]),'_',scrubScores$barcode),rownames(srat@meta.data))
    o = !is.na(m)
    srat@meta.data$scrubScore[m[o]] = scrubScores$score[o]
    srat@meta.data$scrubCall[m[o]] = scrubScores$call[o]
    srat@meta.data$scrubSim1[m[o]] = scrubScores$simScore1[o]
    srat@meta.data$scrubSim2[m[o]] = scrubScores$simScore2[o]
  }
  #Do basic processing and clustering
  srat = FindVariableFeatures(srat,verbose=FALSE)
  srat = ScaleData(srat,verbose=FALSE)
  srat = RunPCA(srat,npcs=numPCs,approx=FALSE,verbose=FALSE)
  srat = FindNeighbors(srat,dims=seq(numPCs),verbose=FALSE)
  srat = FindClusters(srat,res=clusteringRes,verbose=FALSE)
  srat = RunUMAP(srat,dims=seq(numPCs),verbose=FALSE)
  #Set the basic filters.  Default to TRUE if NA
  srat@meta.data$PASS_MT = !(srat@meta.data$mtGenes >= maxMT)
  srat@meta.data$PASS_nGenes = !(srat@meta.data$nFeature_RNA <= minGenes)
  srat@meta.data$PASS_nCounts = !(srat@meta.data$nCount_RNA <= minUMIs)
  srat@meta.data$PASS_doublet = !(!is.na(srat@meta.data$scrubCall) & (srat@meta.data$scrubCall | srat@meta.data$scrubScore > scrubScoreMax))
  #Work out which ones don't pass because of clustering
  tt = aggregate(!PASS_MT | !PASS_nGenes | !PASS_nCounts | !PASS_doublet ~ seurat_clusters,FUN=mean,data=srat@meta.data)
  srat@meta.data$fracBadClust = tt[match(srat@meta.data$seurat_clusters,tt[,1]),2]
  srat@meta.data$PASS_cluster = !(srat@meta.data$fracBadClust > maxBadFrac)
  #Work out the final filter
  srat@meta.data$PASS = with(srat@meta.data,PASS_MT & PASS_nGenes & PASS_nCounts & PASS_doublet & PASS_cluster)
  #Make a reason for fail variable
  nBasicFail = with(srat@meta.data,4 - (PASS_MT + PASS_nGenes + PASS_nCounts + PASS_doublet))
  tmp = rep('Pass',length(nBasicFail))
  tmp[!srat@meta.data$PASS_MT] = 'highMT'
  tmp[!srat@meta.data$PASS_nGenes] = '#genesLow'
  tmp[!srat@meta.data$PASS_nCounts] = '#countsLow'
  tmp[!srat@meta.data$PASS_doublet] = 'doublet'
  tmp[nBasicFail>1] = 'multiple'
  #Don't want "multiple" to encompass cluster otherwise there'll be heaps where cluster + bad obscures the true reason
  tmp[tmp=='Pass' & !srat@meta.data$PASS_cluster] = 'badCluster'
  srat@meta.data$reasonForFail = tmp
  #Make some QC plots before dropping things
  if(doPlot){
    #Plot distributions of basic things
    df = srat@meta.data
    par(mfrow=c(2,2))
    plot(density(df$mtGenes),main='MT',xlab='MT Frac')
    abline(v=maxMT,col='red')
    plot(density(log10(df$nFeature_RNA)),main='#Genes',xlab='log10(No Genes)')
    abline(v=log10(minGenes),col='red')
    plot(density(log10(df$nCount_RNA)),main='#UMIs',xlab='log10(No UMIs)')
    abline(v=log10(minUMIs),col='red')
    plot(density(df$scrubScore),main='Scrublet Score',xlab='Scrub Score')
    #Standard QC things
    plot(DimPlot(srat,group.by='seurat_clusters',label=TRUE,repel=TRUE)+guides(colour=FALSE))
    #A hack for another of Seurat's stupid decisions
    if('labels' %in% names(as.list(args(DimPlot)))){
      plot(DimPlot(srat,group.by=c('Phase','orig.ident'),labels=c('Phase','Source')))
    }else{
      gg = DimPlot(srat,group.by=c('Phase','orig.ident'),combine=FALSE)
      labs = c('Phase','Source')
      for(i in seq_along(labs)){
        gg[[i]] = gg[[i]]  + ggtitle(labs[i])
      }
      plot(patchwork::wrap_plots(gg))
    }
    plot(FeaturePlot(srat,c('mtGenes','nFeature_RNA','nCount_RNA','fracBadClust')))
    #Which cells pass based on what
    if('labels' %in% names(as.list(args(DimPlot)))){
      plot(DimPlot(srat,group.by=c('PASS_MT','PASS_nGenes','PASS_nCounts','PASS_doublet','PASS_cluster','PASS'),labels=c('Pass MT Frac?','Pass #Genes?','Pass #Counts?','Pass Doublet?','Pass cluster','PASS'),legend='none'))
    }else{
      gg = DimPlot(srat,group.by=c('PASS_MT','PASS_nGenes','PASS_nCounts','PASS_doublet','PASS_cluster','PASS'),combine=FALSE)
      labs =c('Pass MT Frac?','Pass #Genes?','Pass #Counts?','Pass Doublet?','Pass cluster','PASS')
      for(i in seq_along(labs)){
        gg[[i]] = gg[[i]] +guides(colour=FALSE) + ggtitle(labs[i])
      }
      plot(patchwork::wrap_plots(gg))
    }
    plot(DimPlot(srat,group.by=c('reasonForFail')))
   #srat@meta.data$isClustBad = ifelse(srat@meta.data$fracBadClust >maxBadFrac | !srat@meta.data$PASS,'Drop','Keep')
    #plot(DimPlot(srat,group.by='isClustBad',label=FALSE))
    #srat@meta.data$isClustBad=NULL
    #srat@meta.data$fracBadClust=NULL
  }
  #Filter
  w = which(srat@meta.data$PASS)
  #w = which(!(as.character(srat@meta.data$seurat_clusters) %in% as.character(tt[tt[,2]>maxBadFrac,1]) | !srat@meta.data$PASS))
  #srat@meta.data$clusterPASS = TRUE
  #srat@meta.data$clusterPASS[w] = FALSE
  #Make the new version, store the old one in it
  sratOld = srat
  #Restart without them
  mDat = srat@meta.data[w,]
  #Drop old clustering
  mDat = mDat[,!colnames(mDat) %in% c('seurat_clusters',grep('^RNA_snn_res',colnames(mDat),value=TRUE))]
  srat = CreateSeuratObject(srat@assays$RNA@counts[,w])
  #Merge in metadata
  srat@meta.data = cbind(srat@meta.data,mDat[,!(colnames(mDat) %in% colnames(srat@meta.data)),drop=FALSE])
  srat = NormalizeData(srat,verbose=FALSE)
  srat = FindVariableFeatures(srat,verbose=FALSE)
  srat@misc$preQC = sratOld
  return(srat)
}


#' Run scrublet
#' 
#' Uses a crude call to python to run scrublet and pull in the results.
#'
#' @param dat10X Directory containing 10X matrix and genes files.
#' @param nPCs Number of PCs to use.
#' @return A data.frame with the scrublet results
runScrublet = function(dat10X,nPCs){
  fNom = tempfile()
  cmd = sprintf(
  "import scrublet as scr
import scipy.io
import numpy as np
import os
import gzip
os.chdir('%s')
input_dir = os.path.expanduser('%s')
output_file = '%s'
try:
  counts_matrix = scipy.io.mmread(input_dir + '/matrix.mtx').T.tocsc()
  genes = np.array(scr.load_genes(input_dir + '/genes.tsv', delimiter='\\t', column=1))
except FileNotFoundError:
  counts_matrix = scipy.io.mmread(input_dir + '/matrix.mtx.gz').T.tocsc()
  #Copy relevant part of scr.load_genes
  gene_list = []
  gene_dict = {}
  with gzip.open(input_dir + '/features.tsv.gz','rt') as f:
    for l in f:
      gene = l.strip('\\n').split('\\t')[1]
      if gene in gene_dict:
        gene_dict[gene] += 1
        gene_list.append(gene + '__' + str(gene_dict[gene]))
        if gene_dict[gene] == 2:
          i = gene_list.index(gene)
          gene_list[i] = gene + '__1'
      else: 
        gene_dict[gene] = 1
        gene_list.append(gene)
  genes = np.array(gene_list)
scrub = scr.Scrublet(counts_matrix)
doublet_scores, predicted_doublets = scrub.scrub_doublets(min_counts=2, 
                                                        min_cells=3, 
                                                        min_gene_variability_pctl=85, 
                                                        n_prin_comps=%d)
with open(output_file,'w') as f:
  #Header row
  f.write('score\\tcall\\tsimScore1\\tsimScore2\\n')
  #As nSim = 2*nObs, we can split into two columns 
  offset = len(doublet_scores)
  f.write('\\n'.join([str(x)+'\\t'+str(predicted_doublets[i]).upper()+'\\t'+str(scrub.doublet_scores_sim_[i])+'\\t'+str(scrub.doublet_scores_sim_[offset+i]) for i,x in enumerate(doublet_scores)]))
  f.close()",getwd(),dat10X,fNom,nPCs)
  system(paste0('python -c "',cmd,'"'))
  out = read.table(fNom,header=TRUE,sep='\t')
  if(file.exists(file.path(dat10X,'barcodes.tsv'))){
    tmp = read.table(file.path(dat10X,'barcodes.tsv'),header=FALSE)
  }else{
    tmp = read.table(file.path(dat10X,'barcodes.tsv.gz'),header=FALSE)
  }
  out$barcode = tmp[,1]
  return(out)
}
```
