## supplementary scripts for "Tumor to normal single cell mRNA comparisons reveal a pan-neuroblastoma cancer cell": trajectoryExpr.R

```
setwd('~/trueHome/Projects/SCP/')

#############
# Libraries #
#############
library(Seurat)
library(Matrix)
import('Code/plotParts.R',as='pt')

##############
# Parameters #
##############
#Output directory
outDir = 'Results/'
srat = readRDS('Data/fAdRef.RDS')


########
# Code #
########
#Get mid point of annotation
dd = cbind(srat@meta.data,srat@reductions$umap@cell.embeddings)
dd$ann = as.character(dd$seurat_clusters)
dd$ann[dd$ann=='25']='SCP'
dd$ann[dd$ann=='24']='Bridge'
dd$ann[dd$ann=='13']='Chromaffin'
dd$ann[dd$ann%in%c('19','20')]='Sympathoblastic'
srat@meta.data$ann = dd$ann
#Get coordinates
mids = aggregate(cbind(UMAP_1,UMAP_2) ~ ann,FUN=mean,data=dd)
#Get average expression
plotGene = function(gene,tgts=c('SCP','Bridge','Chromaffin','Sympathoblastic'),plotLegend=TRUE,plotLabs=TRUE){
  #Get the average by annotation
  geneAvg = sapply(split(srat@assays$RNA@data[gene,],srat@meta.data$ann),mean)
  geneFrac = sapply(split(srat@assays$RNA@data[gene,]>0,srat@meta.data$ann),mean)
  #Define colours
  colPal = c('#ffffcc','#ffeda0','#fed976','#feb24c','#fd8d3c','#fc4e2a','#e31a1c','#bd0026','#800026')
  colBreaks = seq(min(geneAvg[tgts]),max(geneAvg[tgts]),length.out=length(colPal))
  colFun = circlize::colorRamp2(breaks=colBreaks,colPal)
  #Get the mids to use, coordinates etc.
  circPos = mids[match(tgts,mids$ann),]
  #Get the rough circle size
  x = srat@reductions$umap@cell.embeddings
  spread = lapply(tgts,function(e) {
                    y = as.numeric(circPos[circPos$ann==e,c(2,3)])
                    xx = x[srat@meta.data$ann==e,]
                    sqrt((xx[,1]-y[1])**2+(xx[,2]-y[2])**2)})
  circSize = min(sapply(spread,quantile,0.95))
  #Alternative manual specification
  circSize = 0.8
  circPos = data.frame(ann=c('SCP','Bridge','Chromaffin','Sympathoblastic'),
                       UMAP_1 = c(1,1,1-sqrt(2),1+sqrt(2)),
                       UMAP_2 = c(0,2,2+sqrt(2),2+sqrt(2)))
  #Now set plot and start
  par(mar=c(0,0,1,0))
  plot(0,
       xlim=c(-2,5),
       ylim=c(-2,5),
       xlab='',
       ylab='',
       xaxt='n',
       yaxt='n',
       type='n',
       main=gene,
       frame.plot=FALSE)
  symbols(circPos[,2],circPos[,3],
          circles=rep(circSize,length(tgts)),
          xpd=NA,
          add=TRUE,
          bg='grey',
          inches=FALSE)
  #points(circPos[,2],circPos[,3],cex=circSize/strwidth('o'),pch=1,xpd=NA)
  #Finally the fill and fill size
  symbols(circPos[,2],circPos[,3],
          circles=circSize*geneFrac[tgts],
          xpd=NA,
          add=TRUE,
          bg=colFun(geneAvg[tgts]),
          inches=FALSE)
  #points(circPos[,2],circPos[,3],
  #       cex=circSize*geneFrac[tgts]/strwidth('o'),
  #       pch=19,
  #       xpd=NA,
  #       col=colPal(geneAvg[tgts]))
  if(plotLegend){
    pt$addColorBar(x=4,
                   y=0,
                   title='Expression',
                   col=colPal,
                   breaks=colBreaks,
                   ticks=range(colBreaks),
                   tickLabs=c('Lowest','Highest'))
  }
  if(plotLabs){
    text(0,0,'SCP',pos=2)
    text(0,2,'Bridge',pos=2)
    text(circPos[3,2]-0.5,circPos[3,3]+1,'Chromaffin',pos=3)
    text(circPos[4,2],circPos[4,3]+1,'Sympathoblast',pos=3)
  }
}

plotGenes= function(genes,nRows=NULL,nCols=NULL){
  if(is.null(nRows) && is.null(nCols)){
    stop("Must specify rows or cols")
  }
  if(is.null(nRows))
    nRows=ceiling(length(genes)/nCols)
  if(is.null(nCols))
    nCols=ceiling(length(genes)/nRows)
  layout(matrix(seq(nRows*nCols),nrow=nRows,ncol=nCols,byrow=TRUE))
  first=TRUE
  for(gene in genes){
    plotGene(gene,plotLegend=first)
    first=FALSE
  }
}

#Make main figure plots
mainGenes = c('MYCN','PHOX2B','ALK','B4GALNT1','ST3GAL5','ST8SIA1')
for(gene in mainGenes){
  pdf(sprintf('Results/expnCircles%s.pdf',gene),width=2,height=2,useDingbats=FALSE)
    plotGene(gene,plotLabs=FALSE)
  dev.off()
}
#Supplementary genes
supGenes = c('SDHB','ATRX','ALK','PHOX2B','MYCN','KIF1B','EZH2','HRAS','NRAS','PTPN11','SMARCA4','BARD1','BRCA2','APC','CHEK2','PALB2','PINK1','PTPRD','SHANK2','B4GALNT1','ST3GAL5','ST8SIA1','RARA','RARB','RARG','RXRA','RXRB','RXRG')
pdf('Results/expCirclesSupplementary.pdf',width=8,height=10,useDingbats=FALSE)
  plotGenes(supGenes,nCols=5)
dev.off()


srcs = 'Data/scData/fAdrenal'
srcs = file.path(srcs,list.files(srcs))
noms = basename(srcs)
srcs = lapply(srcs,function(e) setNames(file.path(e,list.files(e)),list.files(e)))
names(srcs) = noms
#Final cleanup
#Drop the bad bilateral adrenal sample
srcs$bilateralAdrenals = grep('7457$',srcs$bilateralAdrenals,invert=TRUE,value=TRUE)
srcs$bilateralAdrenals = setNames(file.path(srcs$bilateralAdrenals,'outs'),names(srcs$bilateralAdrenals))
#Drop the non-folders
srcs$lateAdrenals = grep('\\.RDS$',srcs$lateAdrenals,invert=TRUE,value=TRUE)
#Create a list of cleaned paths
strained=list()
for(ex in names(srcs)){
  strained[[ex]] = setNames(file.path(outDir,paste0(names(srcs[[ex]]),'_strainedCounts')),names(srcs[[ex]]))
}
#MT cut-off
mtCut = 0.3
#nGene Cut
minNumGenes = 300
#nUMI cut
minNumUMI = 1000
#How many doublets in a cluster before you throw the entire cluster
maxScrubFrac = 0.5
#UMAP parameters to try in grid
minDists = c(0,.01,.1,.3,.5)
NNs = c(5,10,20,50,100)
#Final parameters
finalMinDist = 0.5
finalNN = 50
#Clustering resolution
clusterRes=1.0
#Populations of genes to exclude from the analysis
excludeGenes = unlist(as.list(geneSets)[c('mtGenes','hspGenes','riboGenes')],use.name=FALSE)
excludeGenes = sort(unique(excludeGenes))
#Number of PCs to use for quick analysis during QC
defNumPCs=75
```
